# Parallel Concerted Evolution of Ribosomal Protein Genes in Fungi and Its Adaptive Significance

**DOI:** 10.1101/751792

**Authors:** Alison Mullis, Zhaolian Lu, Yu Zhan, Tzi-Yuan Wang, Judith Rodriguez, Ahmad Rajeh, Ajay Chatrath, Zhenguo Lin

**Author notes:** These authors contributed equally to this work (Alison Mullis, Zhaolian Lu, Yu Zhan). To whom correspondence should be addressed Zhenguo Lin.

## Abstract

Ribosomal proteins (RPs) genes encode structure components of ribosomes, the cellular machinery for protein synthesis. A single functional copy has been maintained in most of 78-80 RP families in animals due to evolutionary constraints imposed by gene dosage balance. Some fungal species have maintained duplicate copies in most RP families. How the RP genes were duplicated and maintained in these fungal species, and their functional significance remains unresolved. To address these questions, we identified all RP genes from 295 fungi and inferred the timing and nature of gene duplication for all RP families. We found that massive duplications of RP genes have independently occurred by different mechanisms in three distantly related lineages. The RP duplicates in two of them, budding yeast and Mucoromycota, were mainly created by whole genome duplication (WGD) events. However, in fission yeasts, duplicate RP genes were likely generated by retroposition, which is unexpected considering their dosage sensitivity. The sequences of most RP paralogs in each species have been homogenized by repeated gene conversion, demonstrating parallel concerted evolution, which might have facilitated the retention of their duplicates. Transcriptomic data suggest that the duplication and retention of RP genes increased RP transcription abundance. Physiological data indicate that increased ribosome biogenesis allowed these organisms to rapidly consuming sugars through fermentation while maintaining high growth rates, providing selective advantages to these species in sugar-rich environments.

## INTRODUCTION

Gene duplication has served as a driving force for the evolution of new phenotypic traits and contributed to adaptation of organisms to their specific niches (Ohno 1970; Sidow 1996). Duplicate genes are mainly generated by chromosome or whole genome duplication (WGD), unequal crossing-over, and retroposition (Zhang 2013). Similar to other types of mutations, only a small portion of duplicate genes can be eventually fixed in a population, and the survivors are usually advantageous to the organisms (Zhang 2003; Kondrashov and Kondrashov 2006). Highly diverse retention patterns of duplicate genes have been observed among gene families (Hahn, et al. 2005). For instance, tens to hundreds of odorant receptors (ORs) genes can be found in metazoan genomes (Sanchez-Gracia, et al. 2009). In contrast, many genes have been maintained as a single copy since the divergence of eukaryotes, such as the DNA repair genes RAD51, MSH2, and MLH1 (Lin, et al. 2006; Lin, et al. 2007; Zeng, et al. 2014).

Another notable example is the gene families encoding for cytosolic ribosomal proteins (RPs), which are the structural components of ribosomes. Ribosomes carry out one of the most fundamental processes of living systems by translating genetic information from mRNA into proteins. In eukaryotes, each ribosome consists of two subunits, the small and large subunit, which consist of 78-80 different RPs and four types of ribosomal RNAs (rRNA) (Wool 1979; Wimberly, et al. 2000). RP genes are highly conserved in all domains of life (Korobeinikova, et al. 2012). Each RP was found to have unique amino acid sequences with very limited to none similarities between each other. For most animals studied, only a single functional copy of RP gene is maintained in each family, although many processed pseudogenes may be found (Dudov and Perry 1984; Kuzumaki, et al. 1987; Kenmochi, et al. 1998). As structural components of the highly expressed macromolecular complex, the evolutionary constraints on duplicate RP genes was believed to be imposed by gene dosage balance (Birchler and Veitia 2012). In plants, attributing to polyploidization or WGD events, multiple gene copies are usually present in each RP families in polyploid plants (Vision, et al. 2000; Barakat, et al. 2001). This is probably because all RP genes were duplicated simultaneously by WGD, allowing maintenance of balanced dosage among RPs (Birchler and Veitia 2012).

Similar to polyploid plants, most of RP families in the budding yeast *Saccharomyces cerevisiae* have duplicate copies due to the occurrence of a WGD (Wolfe and Shields 1997; Kellis, et al. 2004). Many RP paralogous genes in *S. cerevisiae* generated by WGD, or RP ohnologs, are more similar to each other than to their orthologous genes due to interlocus gene conversion (Evangelisti and Conant 2010; Casola, et al. 2012). During a interlocus gene conversion, one gene serves as a DNA donor that replaces the sequences of its paralogous gene (Chen, et al. 2007). As a result, the sequences of paralogous genes have been homogenized, resulted in the ancient duplicate events to appear much more recent, which was called “concerted evolution” (Brown, et al. 1972). One of the best-known examples of concert evolution is the genes encoding RNA component of ribosomes, the rRNA genes, in both prokaryotes and eukaryotes (Arnheim, et al. 1980; Schlotterer and Tautz 1994; Blattner, et al. 1997).

According to the Ribosomal Protein Gene Database (RPG) (Nakao, et al. 2004), three of ten fungal species listed have multiple gene copies in most RP families, including *S. cerevisiae*, a fission yeast *Schizosaccharomyces pombe*, and a pin mold *Rhizopus oryza*e. The duplicate RP genes in *R. oryzae* could be generated by a WGD event it is ancestor (Ma, et al. 2009), although it has not been systematically examined. In addition, most RP families have more than four gene copies, which cannot be explained by a single WGD. Unlike *S. cerevisiae* and *R. oryzae*, no WGD has been detected during the evolution of *Sch. pombe* (Rhind, et al. 2011), suggesting that each RP family might be duplicated independently by small scale duplication (SSDs) events. This observation is unexpected because the duplicate genes encoding macromolecules generated by SSDs are much less likely to survive because they are sensitive to gene dosage balance (Li, et al. 1996; Conant and Wolfe 2008). It remains an unexplored dimension about how RP genes have been duplicate and maintained in fungi, particularly in the fission yeast *Sch. pombe*.

The expression of RP genes has been thoroughly linked to growth and proliferation, reflecting their central role in the regulation of growth in yeast (Montagne, et al. 1999; Jorgensen, et al. 2002; Brauer, et al. 2008). In rapid growth yeast cells, ∼50% of RNA polymerase II (Pol II) transcription initiation events are devoted to RP expression (Warner 1999). Therefore, the duplication and retention of RP genes might have more functional impacts on these microorganisms than animals or plants. Like other types of mutations, the occurrence of gene duplication is largely due to scholastic events, but retention of duplicate genes have been mainly driven by natural selection (Panchy, et al. 2016). A better understanding of evolutionary fates of RP duplicate genes could offer new insights into how gene duplication produced adaptive solutions to microorganisms.

To better understand the evolutionary patterns of RP genes and their adaptive significance, we conducted systematic identification and evolutionary analyses of all RP families in all fungal species with well-annotated genomes. We searched for RP genes from 295 fungal species and identified independent duplications of most RP families in three distantly related fungal lineages. We inferred the timing and nature of gene duplication for each RP family in each fungal lineage. We found that a vast majority of RP paralogous genes have experienced repeated gene conversion events that have homogenized their sequences in each species. In aligning with integrative analyses of genomic, transcriptomic data and physiological data, we proposed that the massive duplication, retention and concerted evolution of RP genes have contributed to the evolution of fermentative lifestyle in fungal species. This study offers a classic example illustrating the mechanisms and adaptive significance of maintaining duplicate genes encoding macromolecules.

## RESULTS

### Massive duplications of RP genes found in three distantly related fungal lineages

To determine the prevalence of gene duplications in RP families in fungi, we first searched for RP homologous genes for all fungal species with NCBI Reference Sequence (RefSeq) protein data (Supplementary table 1). As of March 2019, 285 fungal species were annotated with RefSeq protein data, covering five of the seven fungal phyla. We conducted BLASTP searches against the 285 RefSeq protein datasets using amino acid sequences of RP genes from both *S. cerevisiae* and *Sch. pombe* as queries (see Methods and Materials). Based on BLASTP search results, we calculated the gene copy numbers in each RP family for every examined species, and the total number of RP families with duplicate copies (Supplementary table 1).

We considered a species with massive RP duplications if more than 50% (≥ 40) of RP families have duplicate copies. Among the 285 fungi examined, only ten species meet the criterion of massive RP duplication. The ten species distributes in three distantly related fungal lineages: three in the class of Saccharomycetes (budding yeast), four in the class of Schizosaccharomycetes (fission yeast), and three in the phylum of Mucoromycota (Supplementary table 1). Although multiple hits of most CRP families were found in a budding yeast *Candida viswanathii*, because the assembly type of its genome is diploid, the two hits found in most RP families are alleles instead of paralogous gene, it was not considered as massive RP duplication.

Because protein annotations of a genome could be incomplete or inaccurate, manual curation is required for a more accurate survey of RP repertoire. It is necessary to carry out a second-round identification of RP genes with manual curation, focusing on the three fungal lineages. We selected 24 species from the three fungal lineages, including the ten species with massive RP duplication. To provide a more even distribution of taxonomic groups in each lineage, we included ten other species whose genomic data are available in NCBI Whole Genome Shotgun (WGS), Yeast Gene Order Browser (YGOB) and JGI (Byrne and Wolfe 2005; Maguire, et al. 2013). In total, our second-round search examined 34 fungal species, which includes 23 Saccharomycetes species, five Taphrinomycotina species (including the four fission yeasts), and six Mucoromycota species (fig. 1, Supplementary table 2). The phylogenetic relationships of the 34 species were inferred using the amino acid sequences of the largest subunit of RNA Pol II proteins (Supplementary fig. 1). Including the 285 species analyzed in our first round analysis, we have examined a total number of 295 fungal genomes in the two rounds of RP gene searches, representing the largest scale of RP gene survey in fungi to our knowledge.

**Figure 1:**
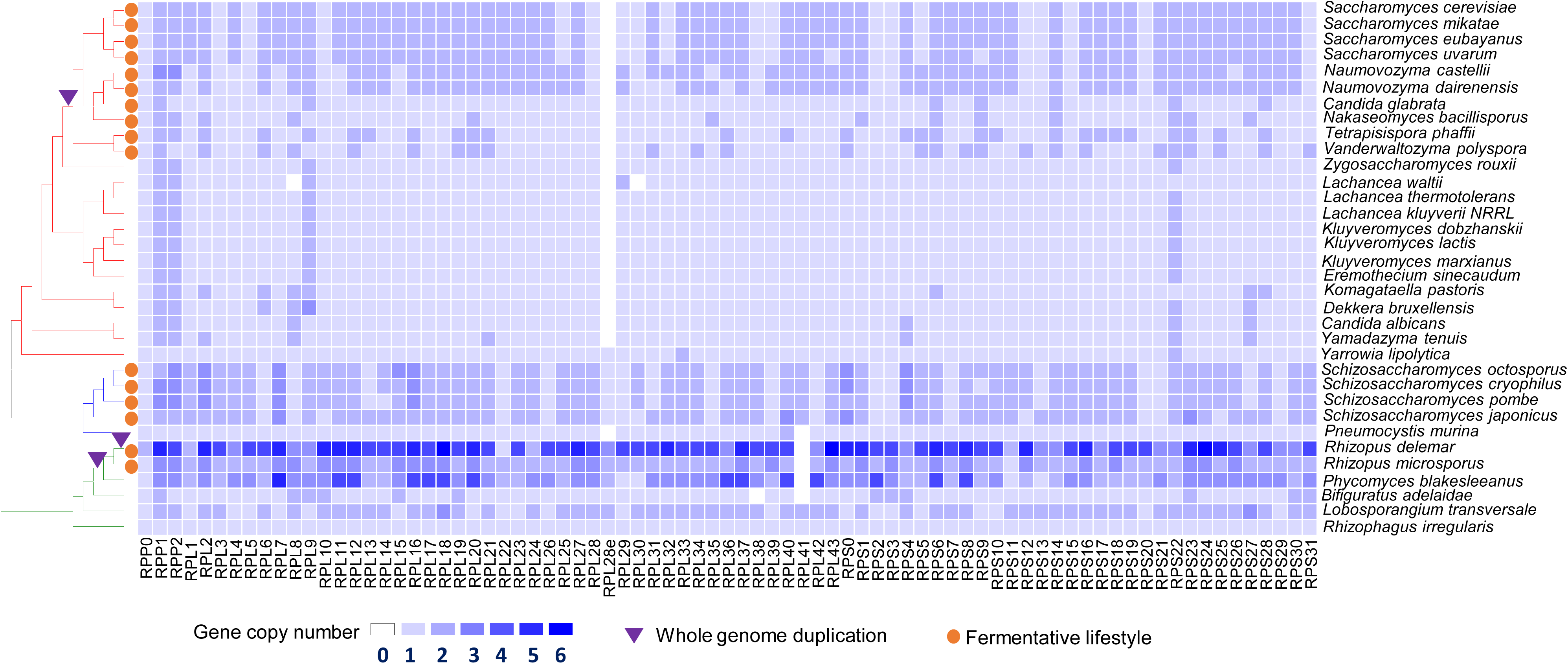
Schematic illustration of gene duplication patterns of 79 RP families in 34 fungal species. Each row represents a fungal species, and each column represents an RP gene family. The colors of a cell represent the numbers of gene copies identified in an RP family of a species. The evolutionary relationship for 34 species, inferred based on amino acid sequences of RNA polymerase II were shown to the left side of the matrix. The species names were provided to the right of the matrix.

To manually curate RP repertoire in a genome, we performed both BLASTP and TBLASTN searches for each of the 34 fungal species. By comparing BLASTP and TBLASTN search results, we identified discrepancies in the number of RP genes and aligned regions. We found that many TBLASTN hits were not present in BLASTP searches, indicating the presence of unannotated RP genes. Thus, we have manually predicted 259 novel RP genes from 32 of the 34 species. We also revised the annotations of open reading frame (ORF) for 95 RP genes. In total, we identified a total number of 3950 RP genes from the 34 fungal species (Supplementary tables 2 and 3).

We constructed maximum likelihood (ML) phylogenetic trees for each RP family (see Materials and Methods). Similar tree topologies were observed among RP families with duplicate copies (Supplementary File 1). For instances, two copies of RPL6 genes are present in all ten post-WGD budding yeasts and all four fission yeast species (fig. 2A). More copies of RPL6 genes are present in Mucoromycota species *Lobosporangium transversale, Phycomyces blakesleeanus, Rhizopus microspores*, and *Rhizopus delemar* (former name *R. oryzae*), which have 2, 4, 3, and 5 copies of RPL6 genes, respectively. According to the ML tree (fig. 2A), the RPL6 genes are more closely related to their paralogous genes in each species, instead of their orthologous genes. Similar patterns are present in the RPS19 genes, which encode a ribosomal small subunit protein (fig. 2B). These tree topologies suggest that the RP genes have been independently duplicated in each species after their divergence from each other. However, at least in the post-WGD budding yeasts, it has been documented that RPL6 and RPS19 were generated by the WGD occurred prior to the divergence of *S. cerevisiae* and *Vanderwaltozyma polyspora* (Conant and Wolfe 2006). Therefore, the phylogenetic trees do not accurately depict the evolutionary history of RPL6 and RPS19 families in budding yeasts. It has been shown that RPL6 and RPS19 genes have experienced gene conversion during the evolution of *S. cerevisiae*, which explains the discrepancy between the tree topology and their duplication history in the budding yeast (Evangelisti and Conant 2010; Casola, et al. 2012). However, it is not known whether it is the same case in the fission yeast and Mucoromycota species, which requires accurate timing of duplication events in the two lineages.

**Figure 2:**
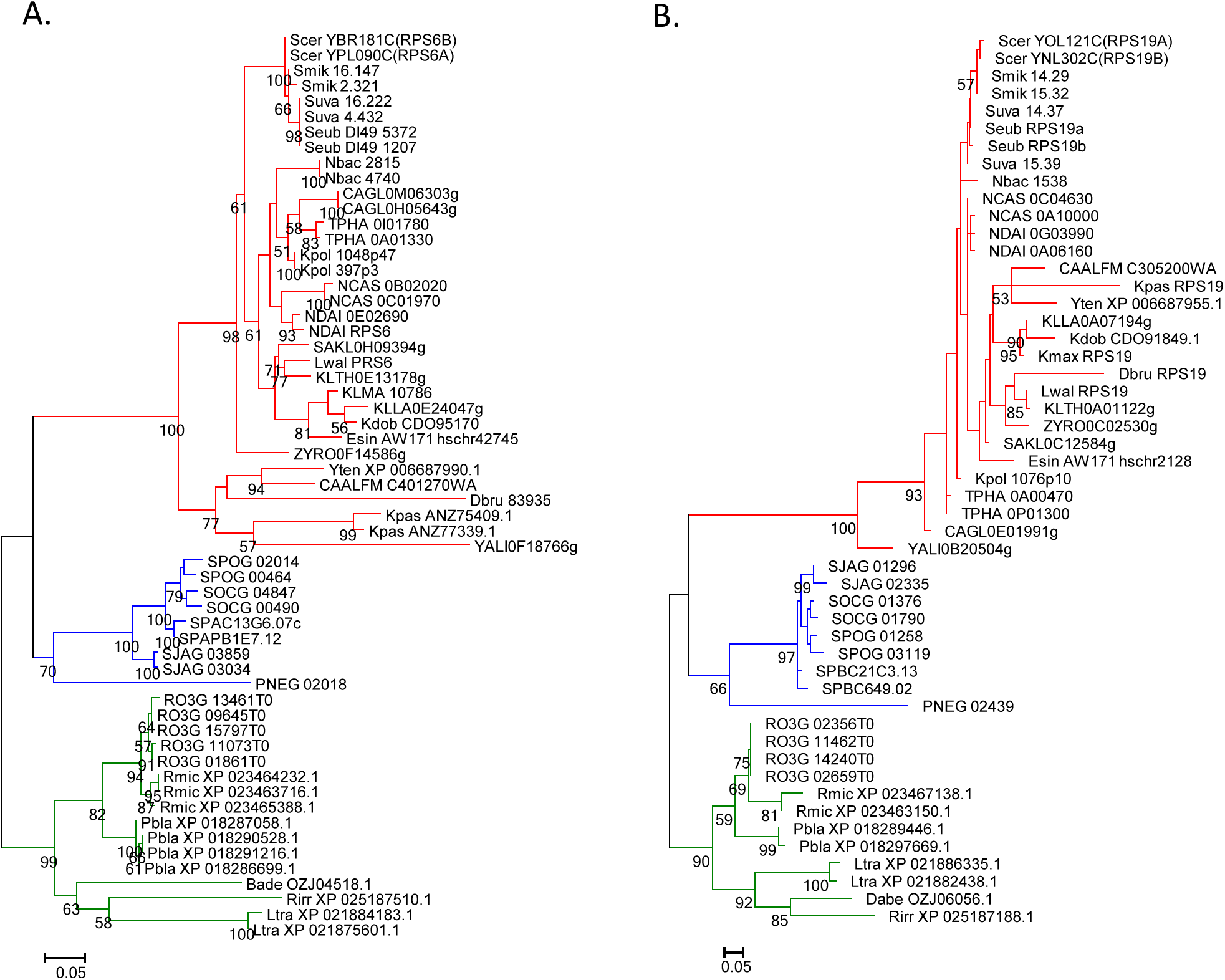
Phylogenetic trees of representative RP gene families. Phylogenetic trees of the RPS6 gene family (A) and the RPS19 (B) gene family in 34 fungal species. The phylogenetic trees were inferred by maximum likelihood (ML) method with 100 bootstrap tests. Only bootstrap values above 50 are shown next to each node. The branches of budding yeast, fission yeast, and Mucoromycota species are colored in red, blue and green respectively. The full species names in taxa were provided in Supplementary table 2.

Based on these cases, we cannot infer the evolutionary history of RP genes in fungi solely based on the topology of phylogenetic trees due to the possibility of gene conversion. It is necessary to carry out additional analyses to determine when gene duplication events have occurred. Because only a small number of species have experienced massive RP duplication in each of the three fungal lineages (fig. 1 and Supplementary table 2), the most parsimonious scenario is that the expansion of RP genes in each fungal lineage occurred independently. In our subsequent analyses, we separately inferred the timing and nature of gene duplications for each RP family and determined whether they have experienced gene conversion after gene duplication in each lineage.

### Duplication and concerted evolution of RP genes in the budding yeasts

We manually identified all RP genes for the 23 Saccharomycetes species (budding yeasts). A total number of 59 RP families have duplicate copies in most WGD species (fig. 1 and Supplementary table 2). 55 of them are ohnologs generated by an ancestral WGD (Conant and Wolfe 2006). The other four RP families, including RPP1, RPP2, RPL9, and RPS22, have duplicates in most budding yeasts, including these non-WGD species, suggesting that they have been duplicated before the divergence of all budding yeasts. Therefore, most post-WGD budding yeasts have a significant increase in RP gene number. 135-137 RP genes are present in the four species in the *Saccharomyces sensu stricto* group, *S. cerevisiae*, S. *mikatae, S. uvarum*, and *S. eubayanus*. The two *Naumovozyma* species have 134 and 137 RP genes, respectively. The numbers are relatively smaller (106 and 104) in the other two early-diverging post-WGD species, *Tetrapisispora phaffii* and *Vanderwaltozyma polyspora*. In contrast, the human opportunistic pathogen *Candida glabrata* (*Nakaseomyces glabrata*) and its closely related species, *Nakaseomyces bacillisporus*, have only 85 and 97 RP genes respectively, suggesting most RP duplicate genes have been lost in these species.

Because the timings of these RP duplications in budding yeast have already been determined, it is possible to infer which RP paralogs have experienced gene conversion by comparing their gene tree with their true duplication history. We can also use the tree topology to infer when concerted evolution had terminated, which is the time when the paralogous genes started to accumulate mutations independently in each paralogous gene. To simplify this process, we constructed phylogenetic trees for each duplicate RP family using five representative WGD species with different divergence times, including *S. cerevisiae, S. mikatae, S. eubayanus, N. castellii* and *T. phaffii* (fig. 3 and Supplementary File 2). We found that at least 52 RP duplicate pairs in *S. cerevisiae* have experienced gene conversion, including 50 pairs generated by WGD and two pairs (RPL9 and RPS22) generated by the ancient duplication events (fig. 3). Therefore, 88% (52 out of 59) of RP paralogous genes in *S. cerevisiae* have experienced gene conversion, which is more than that of previously identified (16 and 29) (Evangelisti and Conant 2010; Casola, et al. 2012), suggesting that concerted evolution of RP genes in the budding yeasts is more prevalent than previously recognized.

**Figure 3:**
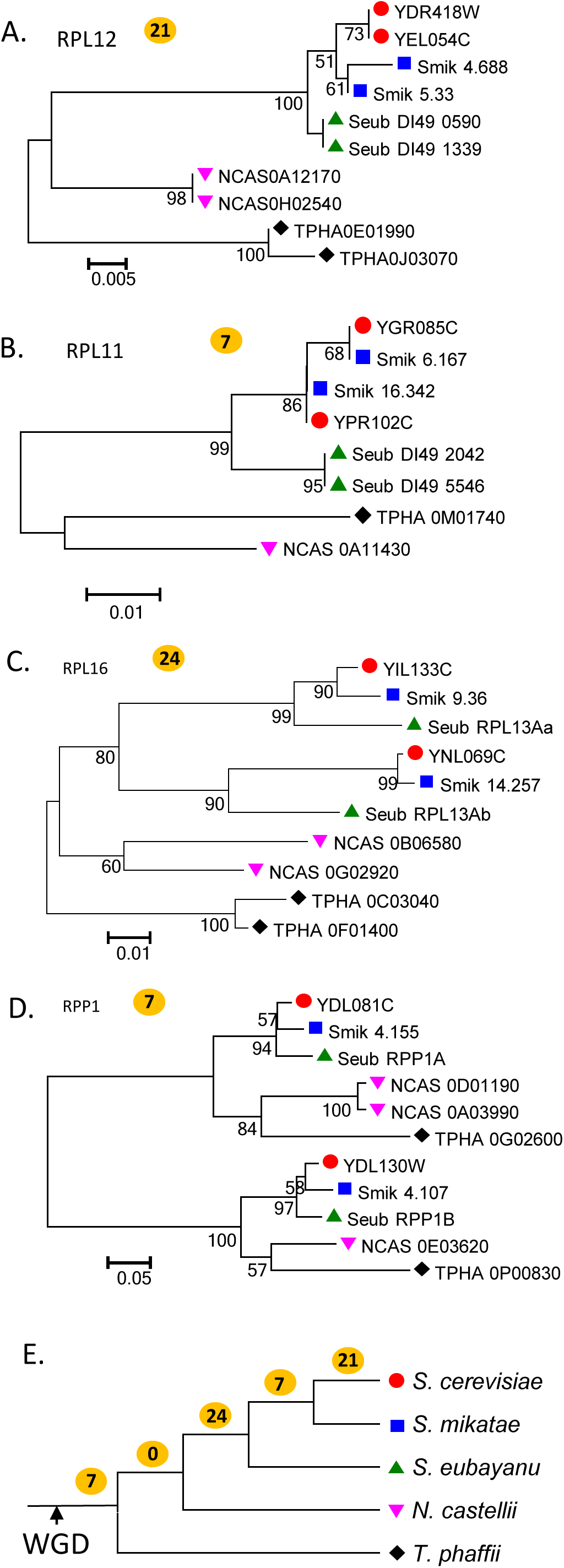
Major types of tree topologies observed from 55 RP families with duplicates in the budding yeasts. **(A)** Phylogenetic relationships of RPL12 genes from five representative budding yeast species. In this case, the two copies of RPL12 genes in *S. cerevisiae* for a species-specific clade. 21 RP gene families demonstrate similar tree topology, as indicated by number “21” in a yellow dot; (B) Phylogenetic relationships of RPL11 genes. The two copies of RPL11 genes in *S. cerevisiae* and *S. mikatae* form a well-supported clade, and the orthologous genes between *S. cerevisiae* and *S. mikatae* are more closely related to each other. Seven RP gene families have a similar tree topology; (C) Phylogenetic relationships of RPL16 genes. Each of RPL16 duplicate genes in *S. cerevisiae* is more closely related to their orthologous genes in *S. mikatae* and *S. eubayanus*. 24 RP gene families share a similar tree topology; (D) Phylogenetic relationships of RPP1 genes. Each of RPP1 duplicate genes in *S. cerevisiae* is more closely related to their orthologous genes in the five WGD species. Seven RP gene families demonstrate a similar tree topology. (E) The distribution of RP families with different termination points of concerted evolution based on evolutionary relationships of RP duplicate genes in *S. cerevisiae*. The numbers on each tree branch represent the numbers of RP families that have terminated concerted evolution at the indicated evolutionary stages.

Based on gene tree topologies, we inferred when converted evolution of RP genes had terminated during the evolution of budding yeasts. In 21 RP families, two copies of each RP family gene in *S. cerevisiae* form a species-specific clade in gene tree (fig. 3A and Supplementary File 2), suggesting that the concerted evolution is still ongoing in *S. cerevisiae* or has recently terminated after its divergence from *S. mikatae*. In seven RP families, termination of concerted evolution occurred before the split between *S. cerevisiae* and *S. mikatae* (fig. 3B). 24 RP families have ended their concerted evolution before the divergence of the *Saccharomyces sensu stricto* group, which including *S. cerevisiae, S. mikatae, S. eubayanus* (fig. 3C). Only seven RP gene pairs do not show strong evidence of gene conversion (fig. 3 D). A summary of the termination time of concerted evolution in 59 RP pairs in *S. cerevisiae* was provided in fig. 3E.

### Duplication and concerted evolution of RP genes in the fission yeasts

We identified all RP genes for five species in the subphylum of Taphrinomycotina, including the four fission yeasts and *Pneumocystis murina. P. murina* belongs to the class of Pneumocystidomycetes, which is probably the most closely related lineage to the fission yeasts, and it was used as an outgroup to infer the evolutionary history of RP genes. 142 to 145 RPs are present in the four fission yeast species. The number of RP families with duplicate copies range from 58 to 59 in the four species (fig. 1 and Supplementary table 2). Most of them have two gene copies, but three copies of RP genes are present in 6 RP families (fig. 1). Only 78 RP genes were identified in *P. murina*. Thus, it is reasonable to assume that the massive expansion of RPs genes in the fission yeasts occurred after their divergence with *P. murina*.

Similar to the budding yeasts, most paralogous RP genes in each fission yeast are more similar to each other than to their orthologous genes (fig. 2 and Supplementary File 3). The tree topologies indicate that these RP genes were duplicated independently in each fission yeast after their divergence. However, we should consider the possibility of gene duplication which resulted in underestimation the ages of duplicate genes. To infer when gene duplication occurred, we conducted gene collinearity (microsynteny) analysis for all duplicate RP genes in the four fission yeasts (see Methods and Materials). If an RP gene was duplicated independently in each species after their divergence, the daughter genes are expected to be found in different genomic regions among these species. Under this scenario, only the parental copy of RP genes share microsynteny by them, and it is extremely unlikely that they also share conserved regions of microsynteny around the daughter genes. On the other hand, if these fission yeasts share microsynteny in both copies of RP genes, the two copies should be created by a single gene duplication event in their common ancestor.

We first identified all orthologous groups in the four fission yeasts (Supplementary table 4). Based on gene order and ortholog group information, we analyzed microsynteny for each pair of RP genes. Herein, we defined a conserved region of microsynteny as a block containing three or more conserved homologs within five genes downstream and upstream of an RP gene (fig. 4A). In *Sch. pombe*, 58 RP families have at least two gene copies. In each of the RP families, the duplicate genes share microsynteny in *Sch. pombe, Sch. cryophilus*, and *Sch. octosporus* (Supplementary table 5). This result suggests that duplicate pairs in the 58 RP families were generated at least before the divergence of the three fission yeasts, which have occurred ∼119 million years ago (mya) (Rhind, et al. 2011). We then inferred how many of gene duplication events occurred even before the split of *Sch. japonicas*, approximately 220 mya (Rhind, et al. 2011). Thanks to the highly conserved gene order in fission yeasts (Rajeh, et al. 2018), we were able to detect the presence of microsynteny in both RP duplicates in 49 families in *Sch. japonicus*, suggesting that gene duplication of these RP families have occurred before the divergence of the four fission yeasts (Supplementary table 5). For example, two copies of RPL11 genes are found in each *Schizosaccharomyces* species. Highly conserved regions of microsynteny surrounding RPL11A genes were found in all four fission yeasts, and so were the RPL11B genes (fig. 4A), supporting that RPL11 was duplicated in their common ancestor and both copies have been maintained in each fission yeast after their divergence.

**Figure 4:**
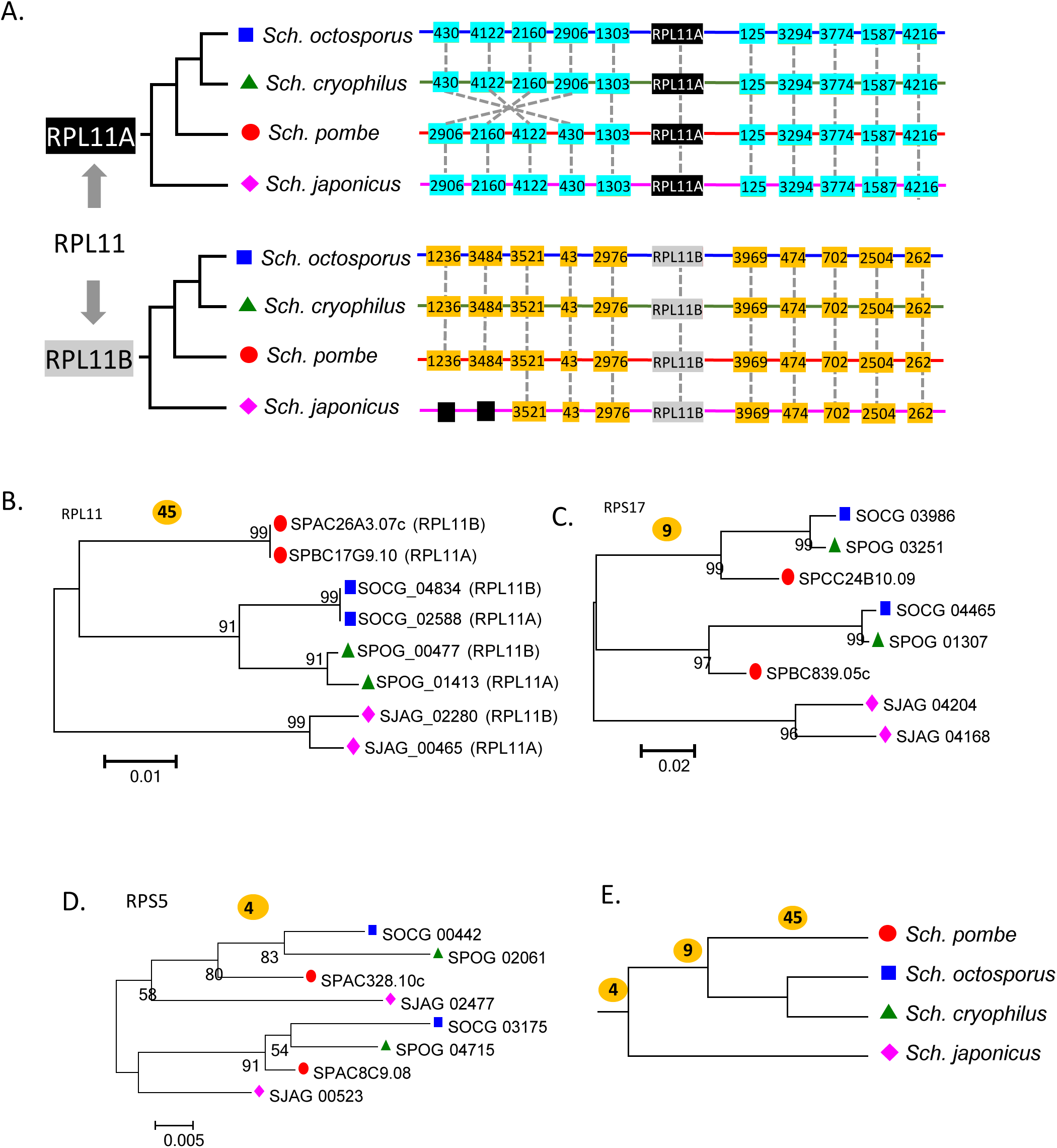
The origin and evolution of RP genes in the fission yeasts. (A) A schematic illustration of microsynteny structures of RPL11 genes in four budding yeasts. The microsynteny blacks of RPL11A are shared by all fission yeasts, so are the RPL11 genes, supporting that the duplication of RPL11 occurred prior to the divergence of the fission yeasts. The number in each box represents its orthologous group ID. (B) Phylogenetic relationships of RPL11 genes in four fission yeast species. The RPL11 duplicate genes in *Sch. pombe* are more closely related to each other than to any orthologous genes. 45 RP gene families demonstrate similar tree topology; (C) Phylogenetic relationships of RPS17 genes. Each of RPL17 duplicate genes in *Sch. pombe* are more closely related to their orthologous genes in *Sch. octosporus* and *Sch. cryophilus*. Nine RP gene families demonstrate similar tree topology; (D) Phylogenetic relationships of RPS5 genes. Each copy of RPS5 duplicates in *Sch. pombe* are more closely related to their orthologous genes. Four RP gene families demonstrate similar tree topology. (E) The distribution of RP families with different termination points of concerted evolution in *Sch. pombe*. The numbers on each tree branch indicate the numbers of RP families that have terminated concerted evolution at the indicated evolutionary stages.

For the rest 9 RP families, we did not obtain conclusive evidence to determine whether they were duplicated before the split of *Sch. japonicus*. Four of them (RPL10, RPL30, RPS12, and RPS25) has only a single RP copy present in *Sch. japonicus*. These genes could be duplicated in the common ancestor of fission yeasts, and a duplicate copy has subsequently lost in *Sch. japonicus*. Alternatively, the duplication events have occurred after the split of *Sch. japonicus* from the other species. In the other five RP families (RPL3, RPL17, RPL18, RPL21, and RPS19), only one RP copy in *Sch. japonicus* share microsynteny with the other three species. Similarly, the duplication events of these RP families could be predated to the divergence of fission yeasts, following by genome rearrangements in *Sch. japonicus* that resulted in the loss of its gene collinearity. However, we cannot exclude the possibility that they were generated by independent duplication events in *Sch. japonicus*.

To determine the number of RP families has an incompatibility between gene phylogenetic tree and duplication history, we constructed a phylogenetic tree for each RP family with duplicates in the fission yeasts. In the case of RPL11, contradict to the gene true duplication history as inferred by microsynteny analysis (fig. 4A), the phylogenetic tree shows that RPL11 paralogs from species-specific clades in each fission yeast (Fig. 4B). Such incompatibility suggests that gene conversion has occurred between RPL11 paralogous genes in each fission yeast after their divergence. A total number of 45 RP families (77.6%) in fission yeasts have a similar tree topology to RPL11 (Supplementary File 3). In other families, such as RPS17, the two copies of RP genes from *Sch. pombe, Sch. octosporus* and *Sch. cryophilus* form two clades and each clade have one gene copy from the three species. We observed nine RP families similar to RPS17, suggesting that concerted evolution of these RP genes might have been terminated before their divergence (fig. 4C). However, we did not find evidence of gene conversion in four RP families, including RPL30, RPS5, RPS12 and RPS28 (fig. 4D and E).

### Retroposition as a major mechanism for massive duplication of RP genes in the ancestral fission yeast

Because no WGD was detected during the evolution of *Sch. pombe* (Rhind, et al. 2011), we then inferred other mechanisms that resulted in massive duplication of RP genes in the fission yeasts, such as unequal crossing-over and retroposition. Unequal crossing-over typically generates segmental or tandem gene duplicates. If a pair of genes was generated by segmental duplication, we expect to observe microsynteny between regions of paralogous RP genes within a species. However, we did not find any case in these RP families (Supplementary table 5). Furthermore, we did not detect tandemly arranged RP paralogous genes, suggesting that unequal crossing-over is not a main contributor for RP duplications in the fission yeasts either.

Retroposition generates retroduplicates through random insertion of a retrotranscribed cDNA from parental source genes, resulting in intron-less retroduplicate genes (Kaessmann, et al. 2009). We examined the exon-intron structure for all RP paralogous genes in *Sch. pombe*. Among the 21 singleton RP families in *Sch. pombe*, only 7 of them (33.3%) are intron-less (Supplementary table 6). In contrast, 33 of 58 duplicate RP families (56.9%) have at least one copy of intron-less gene, which is significantly higher than the group of singleton RPs (*p* = 0.006, Fisher exact test). This ratio is also significantly higher than 27.3% of RP duplicates generated by WGD in *S. cerevisiae*. Thus, the enrichment of intron-less RP genes in the duplicate RP families in fission yeast suggests that they were likely generated by retroposition. For those RP duplicates with intron in both copies, the possibility that they may be created by retroposition following by insertion of intron cannot be excluded, because the locations and phases of introns between these paralogous RP genes in *Sch. pombe* are usually different.

### Duplication and concerted evolution of RP genes in the Mucoromycota species

Four Mucoromycota species examined demonstrate massive duplication of RP genes. Three of them belong to the order of Mucorales (pin molds) in subphylum of Mucoromycotina (311 RP genes in *R. delemar*, 182 in *R. microspores*, and 217 in *P. blakesleeanus*). In their distantly related species in the same subphylum, *Bifiguratus adelaidae*, only 89 RP genes were found. Massive duplication of RP genes (137 RP genes) was also observed in *Lobosporangium transversale*, which is a distantly related species belonging to another subphylum Mortierellomycotina. The earliest diverging species among all Mucoromycota species examined is *Rhizophagus irregularis*, which has only 78 RPs genes (fig. 1).

Based on RP gene copy numbers and the evolutionary relationships of these Mucoromycota species, it is most parsimonious to conclude that massive expansion of RP genes in the three pin mold species and *L. transversale* occurred independently. *L. transversal* is a rare species that having only been reported by a few isolations in North American (Benny and Blackwell 2004). The genomic studies and physiological characterizations *L. transversale* are scarce. Due to lack of genomic data from closely related species, we cannot provide a systematic inference of the timing and nature of massive RP duplication in *L. transversale*. Thus, our subsequent analysis only focused on the origin and evolution of RP duplicate genes in pin molds.

A WGD event has been proposed in ancestral *R. delemar* (Ma, et al. 2009). Another WGD was speculated to have occurred in *P. blakesleeanus* prior to its divergence from *R. microspores* and *R. delemar* (Corrochano, et al. 2016). Therefore, *R. delemar* might have experienced two rounds of WGDs, which correlates with the largest RP repertoire (311) identified in *R. delemar*, while *P. blakesleeanus* and *R. microspores* have 217 and 182 RP genes respectively. Based on RP gene numbers, it is reasonable to conclude that the second WGD occurred after the divergence of *R. delemar* from *R. microspores*.

We conducted microsynteny analysis to infer which RP gene pairs were generated by the WGDs in the pin molds. The estimated divergence time between *Phycomyces* and *Rhizopus* is over 750 mya (Mendoza, et al. 2014). Most, if not all, microsynteny blocks generated by the first WGD might have lost during the evolution of these pin molds. Even though we have used a less strict definition of microsynteny (a minimum of 3 shared homologs in a block of ± 10 neighboring genes surrounding RP), we only identified 3 and 10 pairs of microsynteny blocks between paralogous RP genes in *R. microspores* and *P. blakesleeanus* respectively. In contrast, in *R. delemar*, which has experienced a second round of WGD after its divergence from *R. microspores*, we detected microsynteny for 63 pairs of RP paralogous genes (Supplementary table 7), supporting the recent WGD as a major contributor to the expansion of RP genes in *R. delemar*.

We attempted to identify microsynteny for orthologous RP genes to infer the evolutionary history of each RP family in pin molds (Supplementary table 8). Due to the large divergence times between these species, most RP orthologous genes lack well-supported microsynteny (Supplementary table 7). The most well-supported example is probably the RPL3 family (Fig. 5A). Based on the shared gene orders between paralogous and orthologous RPL3 genes, it is reasonable to infer that RPL3 was duplicated before the divergence of the three pin mold species, probably due to the first WGD. In *R. delemar*, the two RPL3 genes have been further duplicated by the recent WGD, generating four copies. However, their gene tree (fig. 5B) demonstrates that the RPL3 paralogous genes in each species form a species-specific clade, suggesting the occurrence of gene conversion between paralogous RPL3 genes in each species. A total number of 57 RP families have a similar tree topology (Supplementary File 4). Although there is no conclusive microsynteny evidence to support that these RP families have the same evolutionary history as RPL3, we believed that it would be the most likely scenario. In some RP families, such as RPL38 (fig. 5C), the genes from *R. delemar* and *R. microspores* form two clades, and each clade has members from both species. There are 17 RP families have a similar tree topology to RPL38. If these RP genes were the product of the ancient WGD, their concerted evolution had terminated prior to the divergence of the two *Rhizopus* species. The last type of tree topology, such as RPS20 (fig. 5D), whose members form two clades, and each clade include genes from all the three pin mold species. Such tree topology does not support the occurrence of gene conversion. We observed two RP families belonging to this type. In summary, our results implied that most RP paralogous genes in pin molds might have also experienced gene conversion, similar to what happened in budding yeasts and fission yeasts.

**Figure 5:**
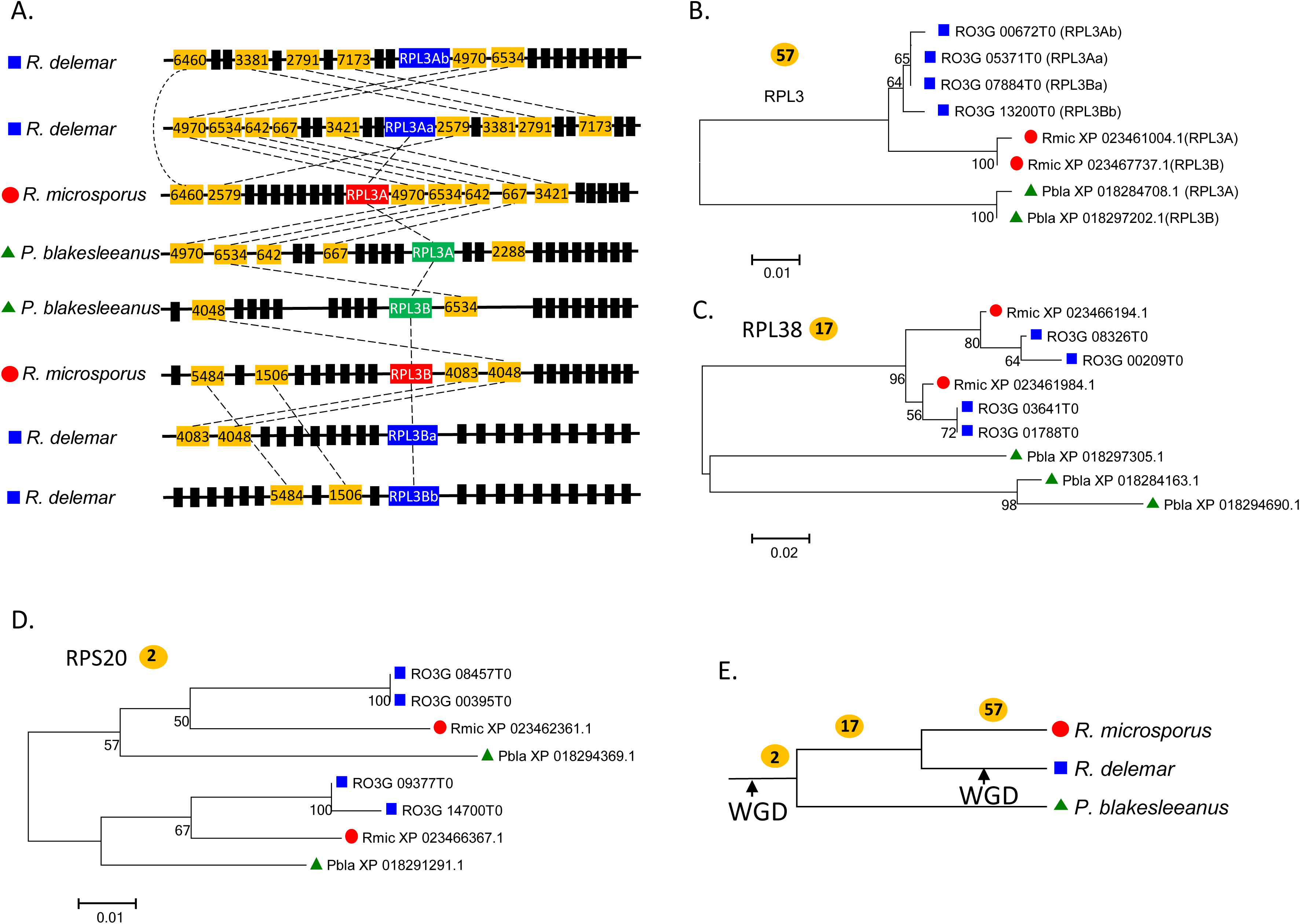
The origin and evolution of RP genes in pin molds. (A) A schematic illustration of microsynteny structures of RPL3 genes in three pin molds. The shared microsynteny structure suggested that the first duplication event of RPL3 genes have occurred prior to the divergence of the pin molds species. The two RPL3 genes have experienced a second round of duplication by WGD in *R. delemar.* (B) Phylogenetic relationships of RPL3 genes in three pin molds. The RPL3 paralogous genes in *R. microsporus* are more closely related to each other than to any orthologous genes. 57 RP gene families demonstrate similar tree topology; (C) Phylogenetic relationships of RPL38 genes in Mucorales. Each of RPL38 duplicate genes in *R. microsporus* is more closely related to their orthologous genes in *R. delemar*. 17 RP gene families demonstrate similar tree topology; (D) Phylogenetic relationships of RPS20 genes in three pin molds. Each RPS20 duplicate gene in *R. microsporus* is more closely related to their orthologous genes. Two RP gene families demonstrate similar tree topology. (E) The distribution of RP families with different termination points of concerted evolution during the evolution of *R. microsporus*. The numbers on each tree branch represent the estimated numbers of RP families that have terminated concerted evolution at the indicated evolutionary stage.

### cDNA as the probable donor for gene conversion between RP paralogous genes

During gene conversion, the genomic sequence of the ‘acceptor’ locus is replaced by a ‘donor’ sequence through recombination (Chen, et al. 2007). The donor can be genomic DNA or cDNA derived from an mRNA intermediate (Derr and Strathern 1993; Storici, et al. 2007). If genomic DNA is the donor, the sequences of both intron and exon can be homogenized. In contrast, if cDNA is the donor, only the exon sequences of the acceptor are replaced. Considering that synonymous mutations are largely free from natural selection, it is possible to determine the donor of gene conversion by comparing the substitution rates between intron and synonymous sites of exons. If the synonymous substitution rates (*d*_*S*_) are significantly lower than intron mutation rates (*μ*_*intron*_), supporting cDNA as a donor. We calculated *d*_*S*_ and *μ*_*intron*_ for all RP duplicate genes for presentative species from each fungal lineage: *S. cerevisiae, Sch. pombe* and *R. microspores* (Supplemental table 9). Overall, the *d*_*S*_ values of all paralogous RP genes are significantly lower than *μ*_*intron*_ in each species examined (fig. 6A-C, Student’s t-test, *p* < 0.01). Considering that different genomic regions might have different mutation rates, we then compared the *d*_*S*_ and *μ*_*intron*_ between each pair of RP duplicate genes (fig. 6D-E). Consistently, most of RP duplicate gene pairs have lower *d*_*S*_ values than *μ*_*intron*_. In a small number of cases, high *d*_*S*_ are observed, probably because the concerted evolution between a pair of orthologous genes have terminated long time ago, resulting accumulation of many synonymous mutations. These results suggest that, in most cases, only the coding sequences have been homogenized by gene conversion, supporting cDNA as the probable gene conversion donor.

**Figure 6:**
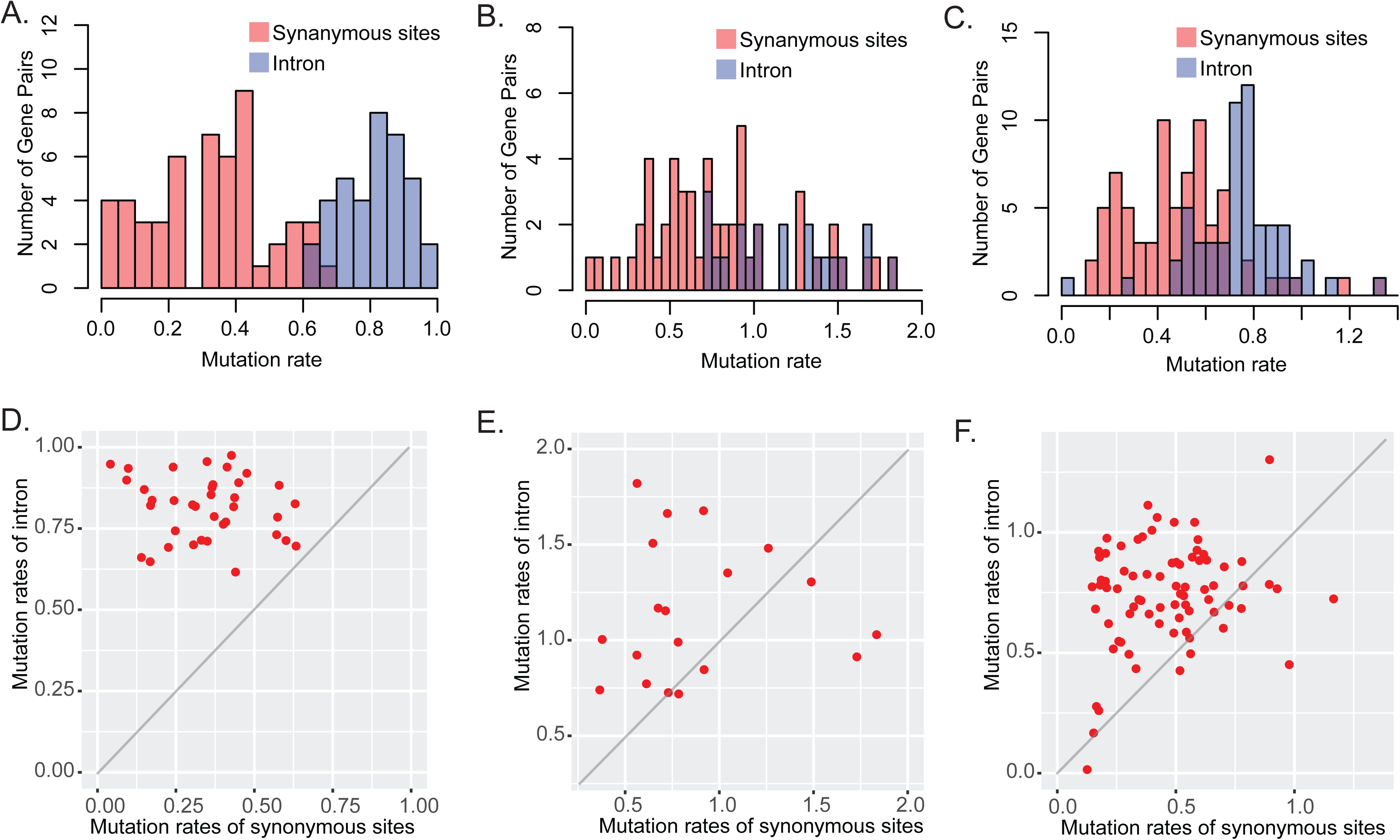
Distinct mutations rates in substitution sites and introns between RP paralogous genes. The distributions of mutation rates in intron and synonymous sites between RP paralogous genes in *S. cerevisiae* (A), *Sch. pombe* (B) and *R. microspores* (C). Scatter plots of mutation rates in intron against synonymous sites between RP paralogous genes in in *S. cerevisiae* (D), *Sch. pombe* (E) and *R. microspores* (F).

### The retention of RP gene duplicates was associated with the evolution to fermentative ability in fungi

Most eukaryotic species fully oxidize glucose, their primary carbon and energy source, through mitochondrial oxidative phosphorylation in the presence of oxygen for maximum energy production. In contrast, post-WGD budding yeasts and fission yeasts predominantly ferment sugar to ethanol in the presence of excess sugars, even under aerobic conditions, which was called aerobic fermentation (Alexander and Jeffries 1990; Lin and Li 2011a). Aerobic fermentation has independently evolved in the budding yeasts and fission yeasts (de Jong-Gubbels, et al. 1996). In addition, the domesticated form of *R. microspores* has been a widely used starter culture for the production of tempeh from fermented soybean (Hachmeister and Fung 1993). Its close relative, *R. delemar*, was also well known as efficient ethanol and fumaric acid producer by fermentation (Kito, et al. 2009; Straathof and van Gulik 2012). *P. blakesleeanus*h was known for capable of fermenting sugar into β-carotene at an industrial scale, which is derived from the end product of glycolysis (Kaessmann, et al. 2009).

We speculated that the massive duplication and retention of RP genes have contributed to the evolution of fermentative ability in these species. Increased gene dosage could lead to a quantitative increase in gene expression and production of protein. To determine the impact of gene duplication on the production of RP transcripts, we calculated the total transcription abundance of all RP genes using our transcriptomic data generated by Cap Analysis of Gene Expression (CAGE) (McMillan, et al. 2019). The CAGE technique captures and sequences the first 75 bp of transcripts, which quantifies the transcription abundance based on numbers of mapped reads (Murata, et al. 2014). Among the 34 species, 11 of them have available CAGE data, including nine budding yeasts and two fission yeasts (Supplementary table 10). As shown in fig. 7A, the RP copy numbers are positively corrected with total transcription abundance value of all RP genes (Supplemental table 10, Pearson correlation *r* = 0.72), supporting that the increased RP gene dosage might have increased ribosome biogenesis by generating more RP transcripts.

**Figure 7:**
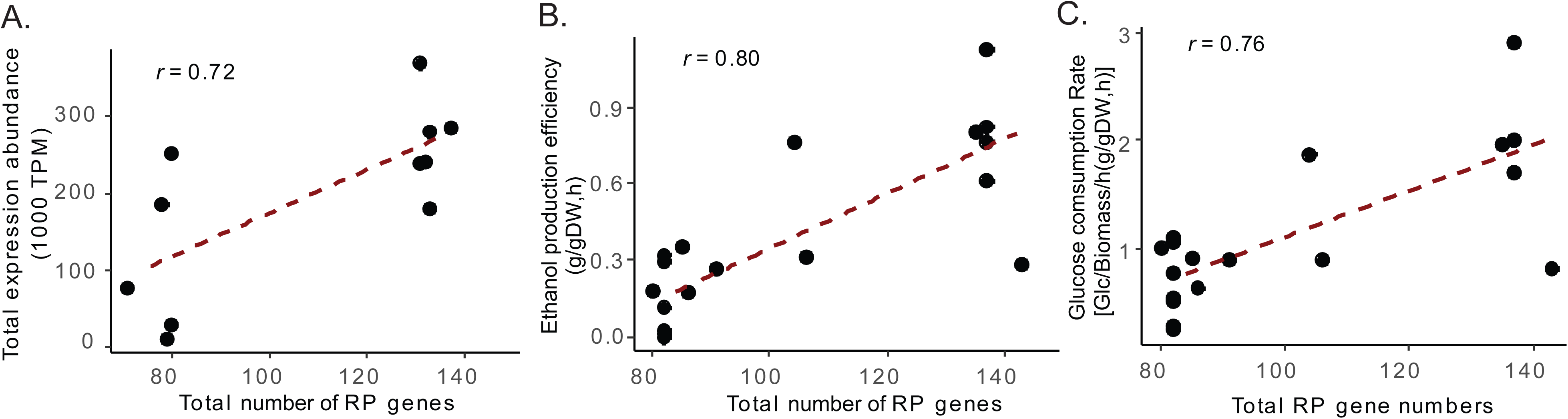
RP gene copy numbers are positively corrected with RP transcription abundance, ethanol production ability, and glucose consumption rates. (A) A scatter plot between the RP copy number and total RP transcription abundance in 11 yeast species. (B) A scatter plot between the RP copy number and ethanol production efficiency in 19 yeast species. (C) A scatter plot between the RP copy number and glucose consumption rate in 19 yeast species.

We then infer whether increased RP gene dosage is associated with better fermentative ability. A previous study has measured various physiological characteristics for over 40 yeast species (Hagman, et al. 2013), including 19 species examined in this study. We observed a positive correlation between RP gene number and ethanol production efficiency (*r* = 0.80), and glucose consumption rate (*r* = 0.76) (fig. 7B and C). We also observed a significant positive correlation between total RP expression and both ethanol production efficiency (r = 0.87) and glucose consumption rate (r = 0.88) (Supplementary fig. 2). These results suggest that the increased RP expression by gene duplication might have enhanced these organisms’ ability to rapidly consuming glucose through the fermentation pathway.

## Discussion

### The preferential retention of RP duplicate genes was selection-driven

Our survey of 295 fungal genomes revealed that massive duplications of RP genes are not prevalent. However, a significant increase in RP gene copy numbers had independently occurred a small number of species in three distantly related lineages in fungi. WGD events have played an important role in the expansion RP repertoire in the budding yeasts and pin molds. In the budding yeasts, only ∼10% of WGD ohnologs have survived, while 70.5% of RP duplicates generated by WGD have been maintained in *S. cerevisiae*. As indicated by previous studies, the survival rate of RP ohnologs is significantly higher than the other WGD ohnologs (Papp, et al. 2003).

Our results suggested that RP genes in the fission yeasts were likely individually duplicated by small-scale duplication events (SSD), such as retroposition. In general, the gene retention rate of SSDs is much lower than ohnologs (half-life of 4 million years vs 33 million years) (Hakes, et al. 2007), it is even lower in genes encoding macromolecular complexes due tp evolutionary constraints imposed by gene dosage balance (Li, et al. 1996; Conant and Wolfe 2008). Similar to the fission yeasts, 99.8% of RP duplicates in mammals were found to be generated by retroposition (Dharia, et al. 2014). However, almost all RP retroduplicates in mammals become pseudogenes (Dharia, et al. 2014). Therefore, the high retention rates of functional RP duplicates generated by SSDs in each fission yeasts are indeed unexpected. A reasonable explanation is that the increased RP gene dosage have provided selective advantages to these species, natural selection favored the retention of RP duplicate genes.

There is another line of evidence supporting that the retention of RP duplicate genes in fission yeasts was selection-driven. The fission yeasts have been known to maintain a single copy of genes in most gene families (Rhind, et al. 2011; Rajeh, et al. 2018). Based on our orthologous group data (Supplementary table 4), in 86% (4069/4734) of fission yeast ortholog groups, only a single copy gene is present in each of the four species (or 1:1:1:1 ortholog). Of the gene families with gene duplication or loss in at least one fission yeast species, ribosomal proteins account for 9.3% (62/665) of them, which is significantly overrepresented in this group (*p* < 10^-5^, Fisher exact test).

### How duplication and retention of RP genes contributed to the evolution of fermentative ability in fungi

Our data suggested that the retention of RP duplication genes might have been driven by their contributions to the evolution of strong fermentative ability in these organisms. The fermentative yeasts were believed to have gained a growth advantage through rapid glucose fermentation in the presence of excess sugars (Piskur, et al. 2006). It was found that *S. cerevisiae* often outgrew its non-fermentative competitors in co-culture experiments (Pérez-Nevado, et al. 2006; Williams, et al. 2015). Fermentation is a much less efficient way to generate energy. Through fermentation, each glucose molecule only yields 2 ATP from glycolysis, compared to 32 ATP through mitochondrial oxidative phosphorylation pathway. In sugar rich environments, fermentative organisms are able to produce more ATP per unit time by rapidly consuming sugars through fermentation, providing selective advantages (Pfeiffer, et al. 2001; Pfeiffer and Morley 2014). The rapid glucose consumption was facilitated by the increased glycolysis flux and more efficiently transporting glucose across cellular membranes. The enhanced glycolytic activity is also present in many tumor cells, known as the “Warburg effect” (Vander Heiden, et al. 2009; Diaz-Ruiz, et al. 2011). It has been shown that increased copy number of genes related to glycolysis (Conant and Wolfe 2007) and glucose transporters (Lin and Li 2011b) have played an important role in the switch of glucose metabolism. Similarly, it is reasonable to propose that the increased RP gene dosage increased their transcript abundance and ribosome biogenesis, resulting in increased biogenesis of glycolysis enzymes, glucose transporters, and other building blocks for cell growth and proliferation.

Ribosome biosynthesis, however, comes with the opportunity cost of higher expression of other cellular processes needed for cell viability and function. Maintaining one RP gene per family may be advantageous for most other species to allow for greater Pol II transcription potential for other genes. However, massive duplication of RP genes made it possible to rapidly consume glucose through the low-efficient fermentative pathway, and at the same time maintain a high growth rate. Such physiological chrematistic provided selective advantages to the organisms in sugar rich environments, which well explained the high retention rate of RP duplicates in fungal species.

### Gene duplication allows further increase of transcription abundance from highly expressed RP genes

One may argue that increased of ribosome biosynthesis can also be achieved by elevated transcription activities of RP genes. It is probably true because we also observed elevated expression level of RP genes in the two non-WGD yeasts with an intermediate level of ethanol fermentation ability: *Lachancea thermotolerans* and *Lachancea waltii* (Hagman, et al. 2013). Across all eukaryotic species, RP genes are the group of most abundantly transcribed genes in eukaryotic cells, accounting for 50% of RNA polymerase II (Pol II) transcription (Warner 1999). RP genes have the highest density of bound Pol II. 100 RP genes have on average >60% of the maximum Pol II occupancy, while a majority of the genome only has <5% of the maximum Pol II density (Venters and Pugh 2009). Therefore, there is limited room for further increase the transcription abundance of RP genes. However, duplication of RP genes provides an additional substrate on which Pol II can transcribe into RP mRNAs, reducing transcription as a rate-limiting step in the process of ribosome biogenesis.

### Gene conversion facilitated retentions of RP duplicate genes

The sequences of all RP families are highly conserved during the evolution of eukaryotes due to their vital roles in many cellular functions (Korobeinikova, et al. 2012). Because misfolding and misinteractions of highly abundant proteins can be more costly, proteins likely RPs should have been under more functional constraints and evolved even slower than other important proteins (Zhang and Yang 2015). It has shown that there is a strong evolutionary constraint posed on the duplicability of genes encoding core components of protein complexes (Li, et al. 2006). Accumulation of new mutations in duplicate genes could impact the stability of protein complexes, posing selective disadvantages. The other structure component of ribosomes, rRNA, is one of the best-known examples of concerted evolution (Liao 1999; Nei and Rooney 2005).

Our study confirms that gene conversion following the duplication of RP genes appears to be a universal path in fungal species. Through gene conversion, the sequences of paralogous genes are homogenized, easing the new mutations accumulated in one of the paralogous genes. It has been shown that highly-expressed genes are more likely to experience mRNA-mediated gene conversion (Weng, et al. 2000; Schildkraut, et al. 2006). Thus, as a group of most actively transcribed genes (Warner 1999), the repeated occurrence of gene conversion on RP genes might be due to the highly expressed nature of RP genes. It was also found that the promoter or flanking genomic sequences are much more divergence than coding sequences between paralogous RP genes (Evangelisti and Conant 2010), further supporting gene conversion in RPs was mediated by cDNA. Gene conversion could provide an additional layer of protection on top of purifying selection to remove newly accumulated mutations (Evangelisti and Conant 2010; Scienski, et al. 2015). Increased RP genes copies could be advantageous to fungal species in sugar-rich environments through fermentative growth. Therefore, the repeated occurrence of gene conversions between RP paralogous genes have contributed to the maintenance of functional RP duplicates in these organisms by removing newly accumulated mutations.

## Materials and Methods

### Data sources, identification and manual curation of RP repertoire

We obtained a list of RP genes from *S. cerevisiae* and *Sch. pombe* from the Ribosomal Protein Gene Database (RPG)(Nakao, et al. 2004). We downloaded RefSeq protein sequence data of 285 fungal genomes from NCBI (Table S1). In the first round of homologous sequences, we used RP sequences from *S. cerevisiae* and *Sch. pombe* as queries to run BLASTP search against the 285 proteomic data (Camacho, et al. 2009). For BLASTP search, we used e-value cutoff of 1e-10, and only hits with a minimum alignment length of 50% of query sequences were considered as homologous RP genes.

In the second round of homologous searches, we obtained the protein and genome sequences of 34 fungal species from NCBI WGS, JGI and Yeast Gene Order Browser (YGOB) (Byrne and Wolfe 2005; Maguire, et al. 2013) (Supplementary table 2). We first used 79 RP protein sequences from *S. cerevisiae* and *Sch. pombe* as queries to search for homologous sequences from the ten newly added fungal species using BLASTP. To identify RP sequences not predicted by existing genome annotations, we conducted TBLASTN searches against all 34 genomic sequences. Manual inspections were performed to compare the BLASTP and TBLASTN results to identify discrepant hits. For hits obtained by TBLASTN by not BLASTP, we predicted the coding sequences (CDS) based on six frame translations of genomic sequences. The exon-intron boundaries were determined based on the TBLASTN alignments and the presence of GT/AG splice sites in flanking intron sequences. We also revised the predicted protein sequences if there is a discrepancy in aligned regions between BLASTP and TBLASTN results. The same gene prediction method was used to revise misannotated ORF.

### Construction of phylogenetic tree for RP genes

We inferred the phylogeny for each of the 79 RP families using the RP protein sequences collected from the 34 fungal species. Sequences were aligned through MUSCLE (Edgar 2004). The molecular phylogenetic tree was inferred by the Maximum likelihood method using RAxML with 100 bootstrap pseudo-replicates (Stamatakis 2006). The best-fit substitution model was inferred by using ProtTest (Abascal, et al. 2005). As the best substitution models are the LG model were identified for the majority of RP families, it was used in our phylogenetic reconstruction. A discrete Gamma distribution [+G] and invariable sites [+I] was used to model evolutionary rate differences among sites. We also constructed lineage-specific phylogenetic trees for duplicate RP families using representative species from each of the three fungal lineages using Neighbor-Joining method using MEGA 7 (Kumar, et al. 2016). For those RP families with almost identical amino acid sequences, we used their CDS for construction of gene trees to obtain better resolved phylogenetic trees (Supplementary Files 2-4).

### Homology microsynteny analysis

We conducted microsynteny analysis for each RP gene family in the four Schizosaccharomyces species and four Mucoromycotina species, including *R. delemar, R. microspores, P. blakesleeanus* and *B. adelaidae*. Gene order information was retrieved from genome annotations of each species obtained from NCBI. The orthologous gene groups in the fours fission yeast species and four Mucoromycotina species were respectively identified using the OrthoDB (Kriventseva, et al. 2014). For fission yeasts, we obtained a list of ten genes surrounding each RP gene (five upstream and five downstream of RP gene). For the Mucorales species, we extended our microsynteny analysis to a block of ten genes upstream and ten downstream of RP gene due to their divergent genome structures.

### Estimation of substitution rates in intron and synonymous sites

We calculated the substitution rates for every pair of duplicate RP genes for three species representing the three fungal lineages with massive RP duplications, including *S. cerevisiae, Sch. pombe, R. microspores*. The CDS and intron sequences were retrieved from NCBI, and were aligned using MUSCLE. Synonymous substitution rate was calculated using Li-Wu-Luo method with Kimura 2-parameter model (Li, et al. 1985) in MEGA 7 (Kumar, et al. 2016). Nucleotide substitution rates in intron sequences were calculated using the Kimura 2-parameter model in MEGA 7.

### Analysis of RP gene transcriptomic and physiological data

The transcriptomic data of nine budding yeasts and two fission yeasts examined were obtained from (McMillan, et al. 2019) based on CAGE. The expression abundance of an RP gene was defined as the sum of transcripts initiated from all core promoters within 500 base pairs upstream of its annotated start codon, which were normalized as TPM (tags per million mapped reads, and each tag represent one sequenced transcript). The total transcription abundance of RP genes in a species was calculated as the sum of TPM of all RP genes identified in this species. For newly predicted RP genes that were not annotated in CAGE datasets, we performed TBLASTN searches to determine their genomic locations of CDS and obtain the expression abundance data using the same criteria. The ethanol production efficiency and glucose consumption rate of 19 budding and fission yeast species were obtained from (Hagman, et al. 2013). The ethanol production efficiency was measured as grams of ethanol produced per gram of biomass per gram of glucose consumed. The glucose consumption rate was measured as grams of glucose consumed per gram of biomass per hour. If multiple biological replicates were measured for a single species, their average values were used for our analysis.

## Acknowledgements

This study was supported by the start-up fund and President’s Research Fund from Saint Louis University to ZL. We would like to thank Dr. Zhenglong Gu and Dr. Dapeng Zhang for valuable discussions and suggestions to this work.

## Supplementary Figures

**Supplementary fig. 1: The phylogenetic relationships of 34 representative fungal species.** The phylogenetic tree was inferred based on amino acid sequences of RPA polymerase II using ML method by RAxML with 100 bootstrap replicates.

**Supplementary fig. 2: The total transcription abundance of RP genes is positively correlated with ethanol production efficiency and glucose consumption rates.** (A) A scatter plot between total RP transcription abundance and ethanol production efficiency in 9 yeast species. (B) A scatter plot between the total RP transcription abundance and glucose consumption rate in 9 yeast species.

## Supplementary Tables

Table S1: List of numbers of RP genes identified in 285 fungal species

Table S2: List of numbers of RP gene identified in 34 representative fungal species based on manual curation.

Table S3: List of RP genes identified in 34 representative fungal species

Table S4: List of orthologous groups identified in four fission yeasts

Table S5: Microsynteny data of the 79 RP families in fission yeasts

Table S6: Statistics of intron number in RP genes in *S. cerevisiae* and *Sch. pombe*.

Table S7. Number of share homologous genes between microsynteny blocks near the RP paralogous genes in three Mucorales species

Table S8. Number of share homologous genes between microsynteny blocks near the RP orthologous genes in three Mucorales species

Table S9: Mutation rates of synonymous sites and intron in *S. cerevisiae, Sch. pombe* and *R. microspores*

Table S10: Transcription abundance of RP genes in 11 species based on CAGE data

## Supplementary Files

Supplementary File 1: Phylogenetic trees of RP families from the 34 representatives fungal species.

Supplementary File 2: Phylogenetic trees of RP families in the five WGD budding yeast species.

Supplementary File 3: Phylogenetic trees of RP families in the four fission yeast species.

Supplementary File 4: Phylogenetic trees of RP families in the four Mucoromycotina species.

