## Supplementary figures and images for "Parallel Concerted Evolution of Ribosomal Protein Genes in Fungi and Its Adaptive Significance"

### Supplemental Figures

#### Supplementary fig. 1

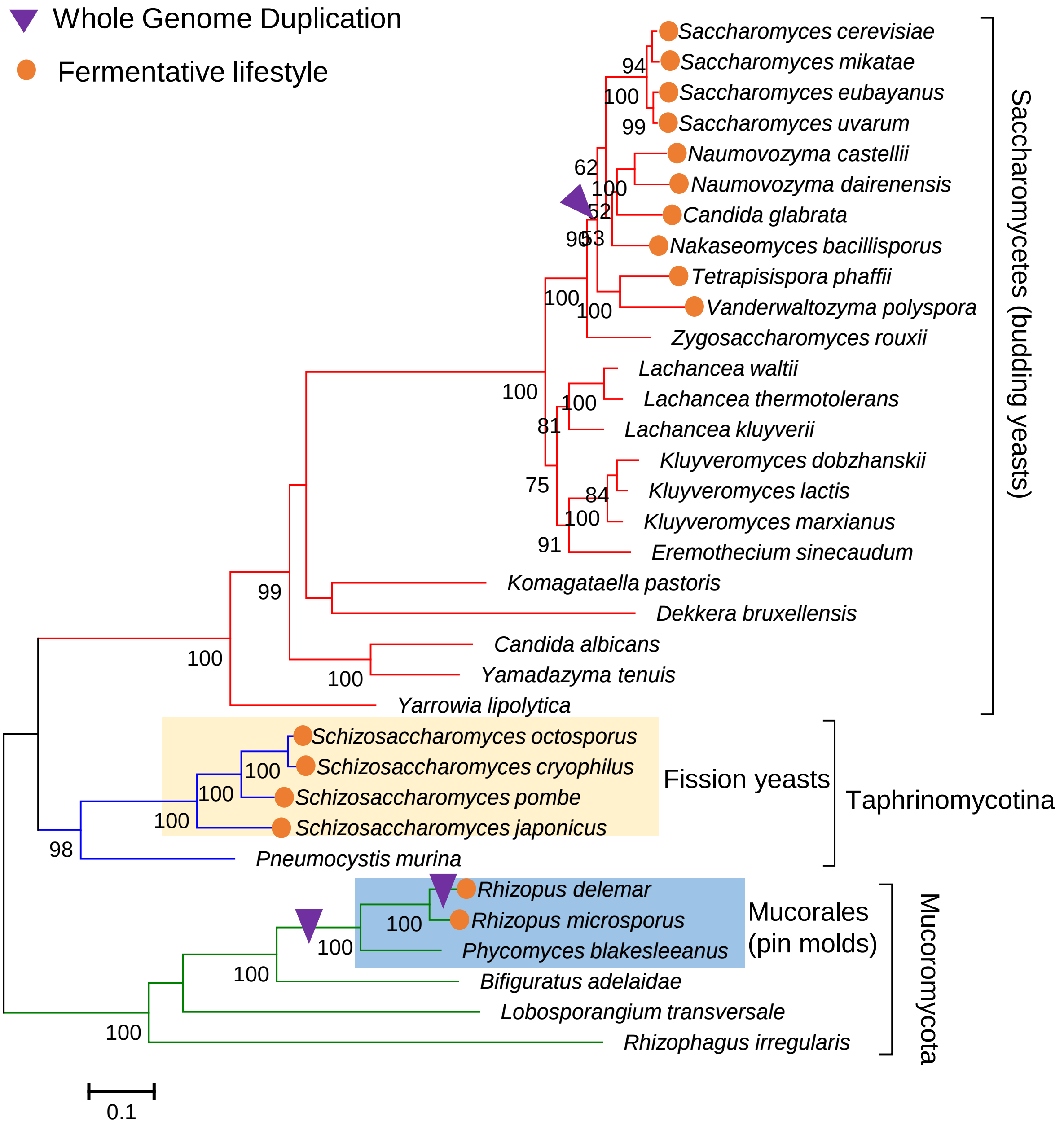

Supplementary fig. 2

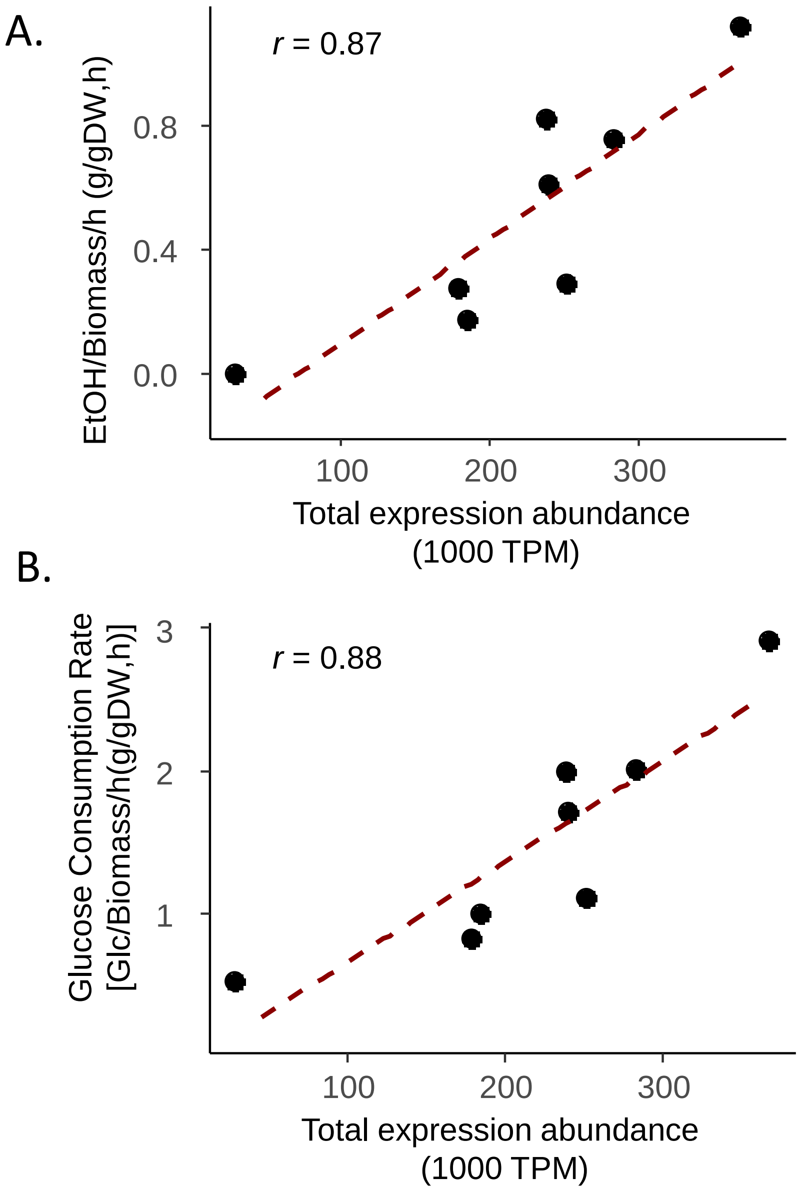
