## Supplemental File 1 for "Parallel Concerted Evolution of Ribosomal Protein Genes in Fungi and Its Adaptive Significance"

**Supplementary File 1: Phylogenetic trees of RP families from the 34 representative 34 fungal species.**


| Gene Family | Page | Gene Family | Page |
|---|---|---|---|
| RPL1 | 2 | RPP0 | 46 |
| RPL2(RPL8) | 3 | RPP1 | 47 |
| RPL3 | 4 | RPP2 | 48 |
| RPL4 | 5 | RPS0(RPSA) | 49 |
| RPL5 | 6 | RPS1(RPS3A) | 50 |
| RPL6 | 7 | RPS2 | 51 |
| RPL7 | 8 | RPS3 | 52 |
| RPL8(RPL7A) | 9 | RPS4 | 53 |
| RPL9 | 10 | RPS5 | 54 |
| RPL10 | 11 | RPS6 | 55 |
| RPL11 | 12 | RPS7 | 56 |
| RPL12 | 13 | RPS8 | 57 |
| RPL13 | 14 | RPS9 | 58 |
| RPL14 | 15 | RPS10 | 59 |
| RPL15 | 16 | RPS11 | 60 |
| RPL16 | 17 | RPS12 | 61 |
| RPL17 | 18 | RPS13 | 62 |
| RPL18 | 19 | RPS14 | 63 |
| RPL19 | 20 | RPS15 | 64 |
| RPL20 | 21 | RPS16 | 65 |
| RPL21 | 22 | RPS17 | 66 |
| RPL22 | 23 | RPS18 | 67 |
| RPL23 | 24 | RPS19 | 68 |
| RPL24 | 25 | RPS20 | 69 |
| RPL25 | 26 | RPS21 | 70 |
| RPL26 | 27 | RPS22 | 71 |
| RPL27 | 28 | RPS23 | 72 |
| RPL28(RPL27A) | 29 | RPS24 | 73 |
| RPL28e | 30 | RPS25 | 74 |
| RPL29 | 31 | RPS26 | 75 |
| RPL30 | 32 | RPS27 | 76 |
| RPL31 | 33 | RPS28 | 77 |
| RPL32 | 34 | RPS29 | 78 |
| RPL33 | 35 | RPS30 | 79 |
| RPL34 | 36 | RPS31(RPS27A) | 80 |
| RPL35 | 37 | | |
| RPL36 | 38 | | |
| RPL37 | 39 | | |
| RPL38 | 40 | | |
| RPL39 | 41 | | |
| RPL40 | 42 | | |
| RPL41 | 43 | | |
| RPL42 | 44 | | |
| RPL43 | 45 | | |


*All phylogenetic trees were inferred using amino acid sequences by maximum likelihood method with 100 bootstrap tests. Only bootstrap values above 50 are shown next to each node. The branches of budding yeast, fission yeast, and Mucoromycota species are colored in red, blue and green respectively. .

### RPL1

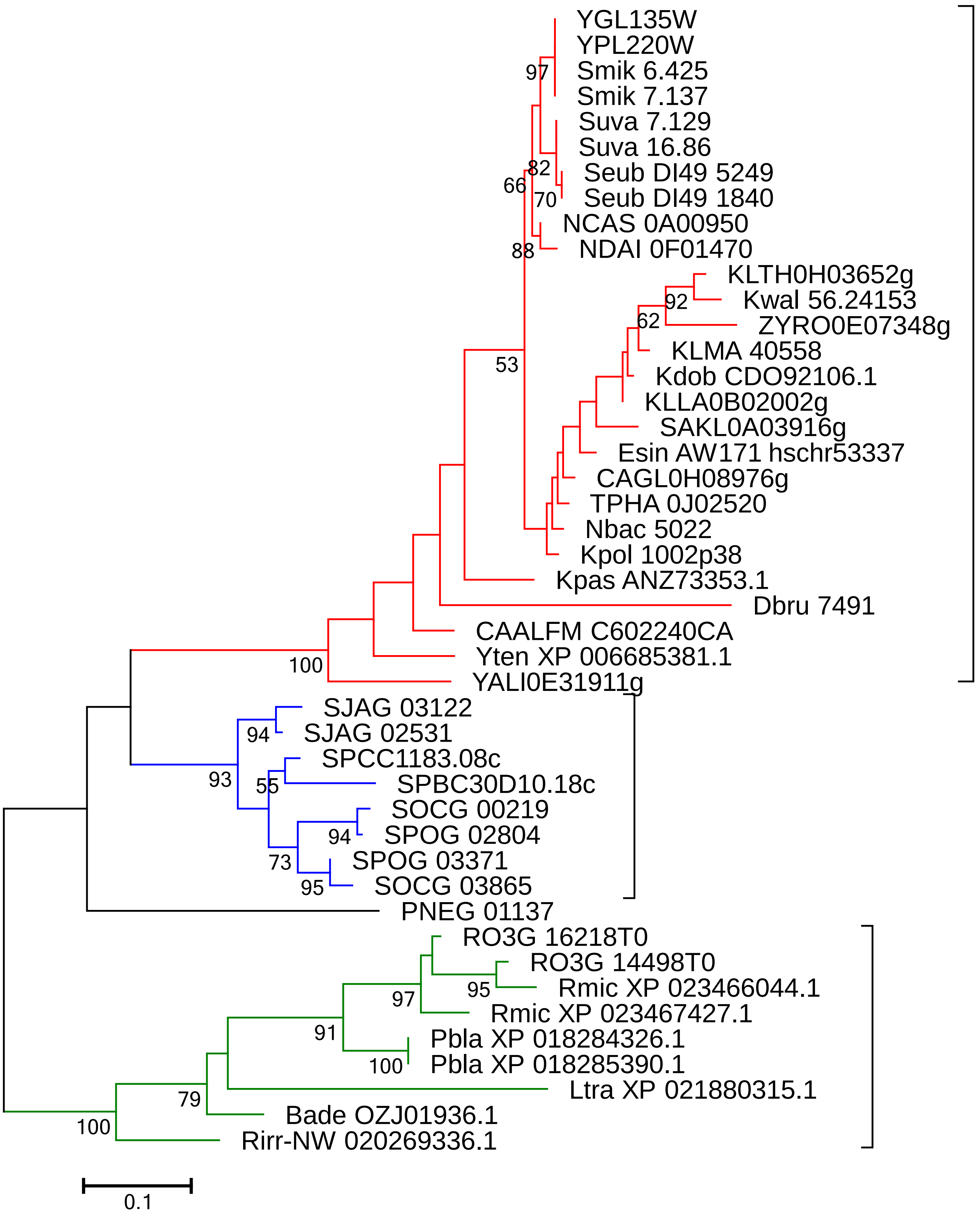

YGL135W
YPL220W
Smik 6.425
Smik 7.137
Suva 7.129
Suva 16.86
Seub DI49 5249
Seub DI49 1840
NCAS 0A00950
NDAI 0F01470
KLTH0H03652g
Kwal 56.24153
ZYRO0E07348g
KLMA 40558
Kdob CDO92106.1
KLLA0B02002g
SAKL0A03916g
Esin AW171 hschr53337
CAGL0H08976g
TPHA 0J02520
Nbac 5022
Kpol 1002p38
Kpas ANZ73353.1
Dbru 7491
CAALFM C602240CA
Yten XP 006685381.1
YALI0E31911g
SJAG 03122
SJAG 02531
SPCC1183.08c
SPBC30D10.18c
SOCG 00219
SPOG 02804
SPOG 03371
SOCG 03865
PNEG 01137
RO3G 16218T0
RO3G 14498T0
Rmic XP 023466044.1
Rmic XP 023467427.1
Pbla XP 018284326.1
Pbla XP 018285390.1
Ltra XP 021880315.1
Bade OZJ01936.1
Rirr-NW 020269336.1
97
82
66
70
88
92
62
53
100
94
93
55
94
73
95
95
97
91
100
79
100
0.1

### RPL2 (RPL8)

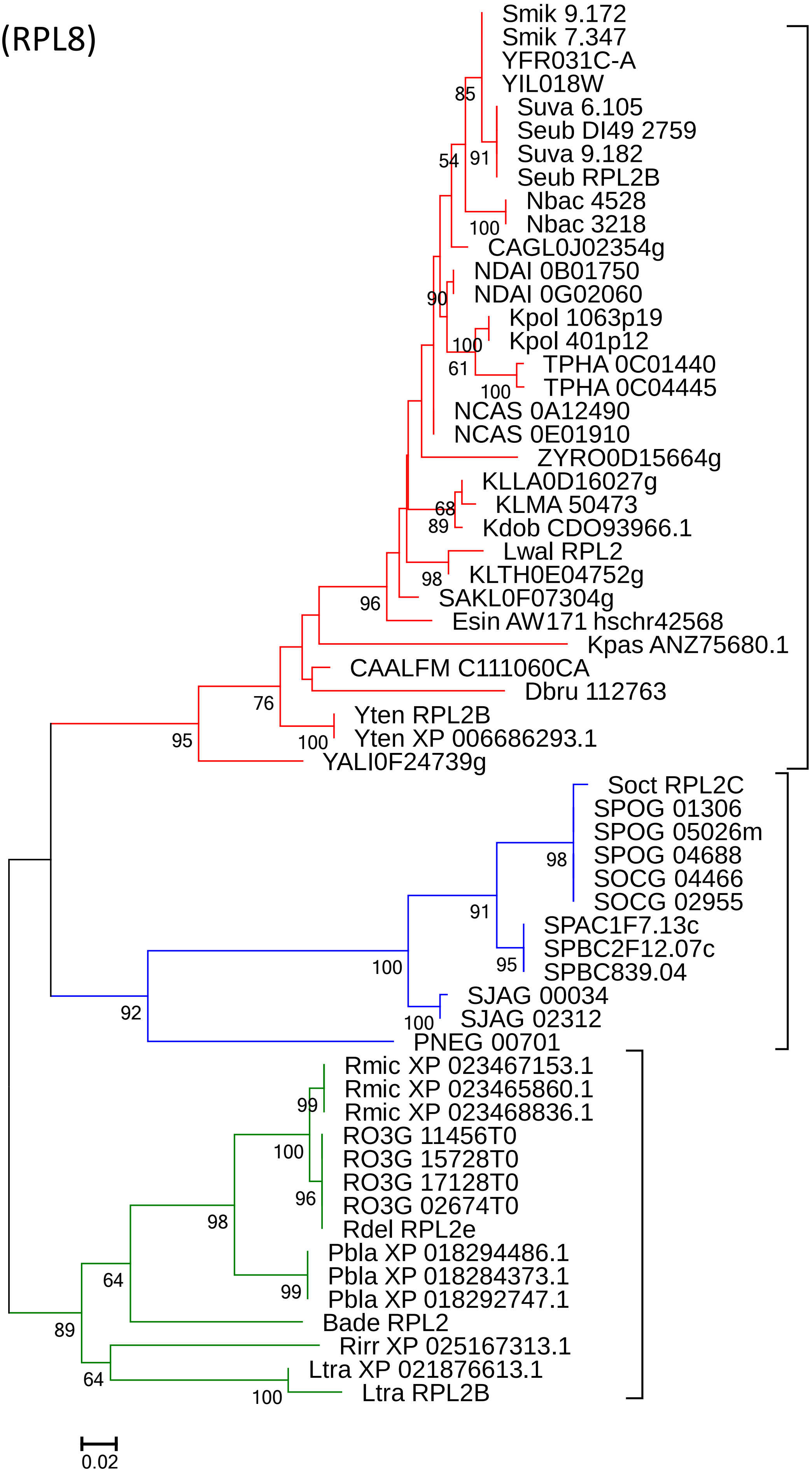

Smik 9.172
Smik 7.347
YFR031C-A
YIL018W
Suva 6.105
Seub DI49 2759
Suva 9.182
Seub RPL2B
Nbac 4528
Nbac 3218
CAGL0J02354g
NDAI 0B01750
NDAI 0G02060
Kpol 1063p19
Kpol 401p12
TPHA 0C01440
TPHA 0C04445
NCAS 0A12490
NCAS 0E01910
ZYRO0D15664g
KLLA0D16027g
KLMA 50473
Kdob CDO93966.1
Lwal RPL2
KLTH0E04752g
SAKL0F07304g
Esin AW171 hschr42568
Kpas ANZ75680.1
CAALFM C111060CA
Dbru 112763
Yten RPL2B
Yten XP 006686293.1
YALI0F24739g
Soct RPL2C
SPOG 01306
SPOG 05026m
SPOG 04688
SOCG 04466
SOCG 02955
SPAC1F7.13c
SPBC2F12.07c
SPBC839.04
SJAG 00034
SJAG 02312
PNEG 00701
Rmic XP 023467153.1
Rmic XP 023465860.1
Rmic XP 023468836.1
RO3G 11456T0
RO3G 15728T0
RO3G 17128T0
RO3G 02674T0
Rdel RPL2e
Pbla XP 018294486.1
Pbla XP 018284373.1
Pbla XP 018292747.1
Bade RPL2
Rirr XP 025167313.1
Ltra XP 021876613.1
Ltra RPL2B
85
54
91
100
90
100
61
100
68
89
98
96
76
95
100
98
91
95
100
100
92
99
100
96
98
99
64
89
64
100
0.02

### RPL3

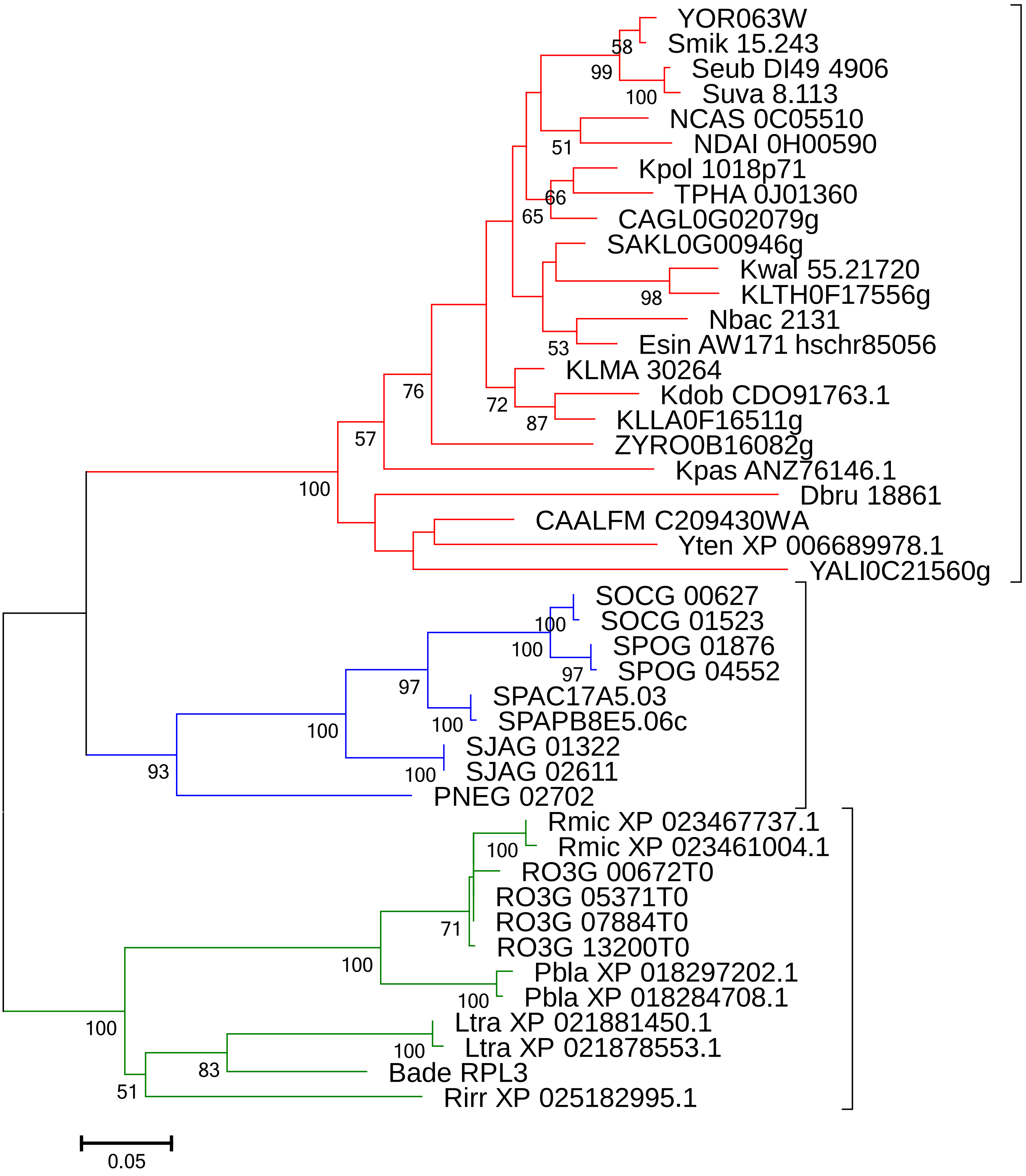
YOR063W
Smik 15.243
Seub DI49 4906
Suva 8.113
NCAS 0C05510
NDAI 0H00590
Kpol 1018p71
TPHA 0J01360
CAGL0G02079g
SAKL0G00946g
Kwal 55.21720
KLTH0F17556g
Nbac 2131
Esin AW171 hschr85056
KLMA 30264
Kdob CDO91763.1
KLLA0F16511g
ZYRO0B16082g
Kpas ANZ76146.1
Dbru 18861
CAALFM C209430WA
Yten XP 006689978.1
YALI0C21560g
SOCG 00627
SOCG 01523
SPOG 01876
SPOG 04552
SPAC17A5.03
SPAPB8E5.06c
SJAG 01322
SJAG 02611
PNEG 02702
Rmic XP 023467737.1
Rmic XP 023461004.1
RO3G 00672T0
RO3G 05371T0
RO3G 07884T0
RO3G 13200T0
Pbla XP 018297202.1
Pbla XP 018284708.1
Ltra XP 021881450.1
Ltra XP 021878553.1
Bade RPL3
Rirr XP 025182995.1
58
99
100
51
66
65
98
53
72
87
76
57
100
100
100
97
97
100
100
100
93
100
71
100
100
100
100
83
51
0.05

### RPL4

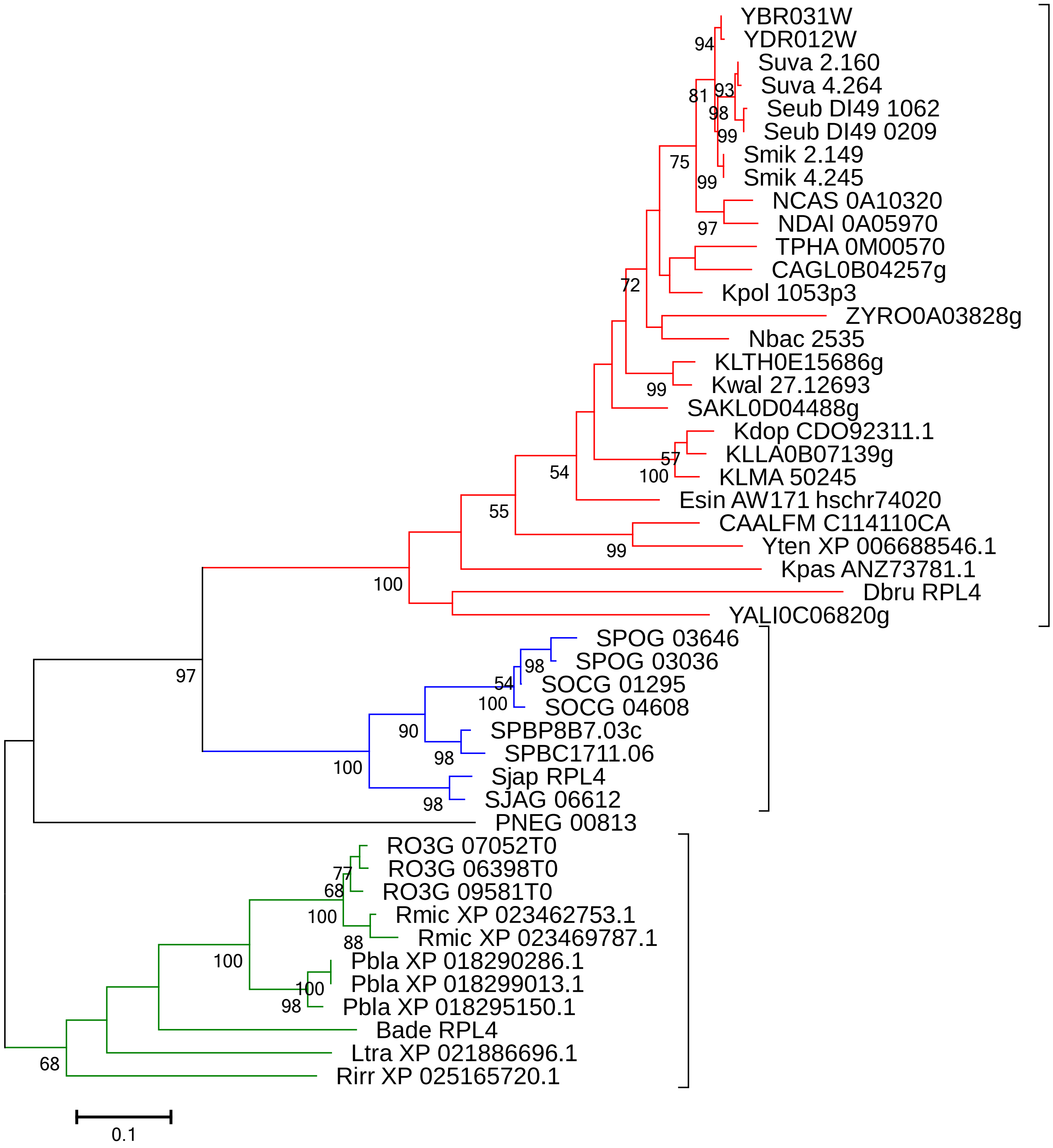
YBR031W
YDR012W
Suva 2.160
Suva 4.264
Seub DI49 1062
Seub DI49 0209
Smik 2.149
Smik 4.245
NCAS 0A10320
NDAI 0A05970
TPHA 0M00570
CAGL0B04257g
Kpol 1053p3
ZYRO0A03828g
Nbac 2535
KLTH0E15686g
Kwal 27.12693
SAKL0D04488g
Kdop CDO92311.1
KLLA0B07139g
KLMA 50245
Esin AW171 hschr74020
CAALFM C114110CA
Yten XP 006688546.1
Kpas ANZ73781.1
Dbru RPL4
YALI0C06820g
SPOG 03646
SPOG 03036
SOCG 01295
SOCG 04608
SPBP8B7.03c
SPBC1711.06
Sjap RPL4
SJAG 06612
PNEG 00813
RO3G 07052T0
RO3G 06398T0
RO3G 09581T0
Rmic XP 023462753.1
Rmic XP 023469787.1
Pbla XP 018290286.1
Pbla XP 018299013.1
Pbla XP 018295150.1
Bade RPL4
Ltra XP 021886696.1
Rirr XP 025165720.1
0.1

### RPL5

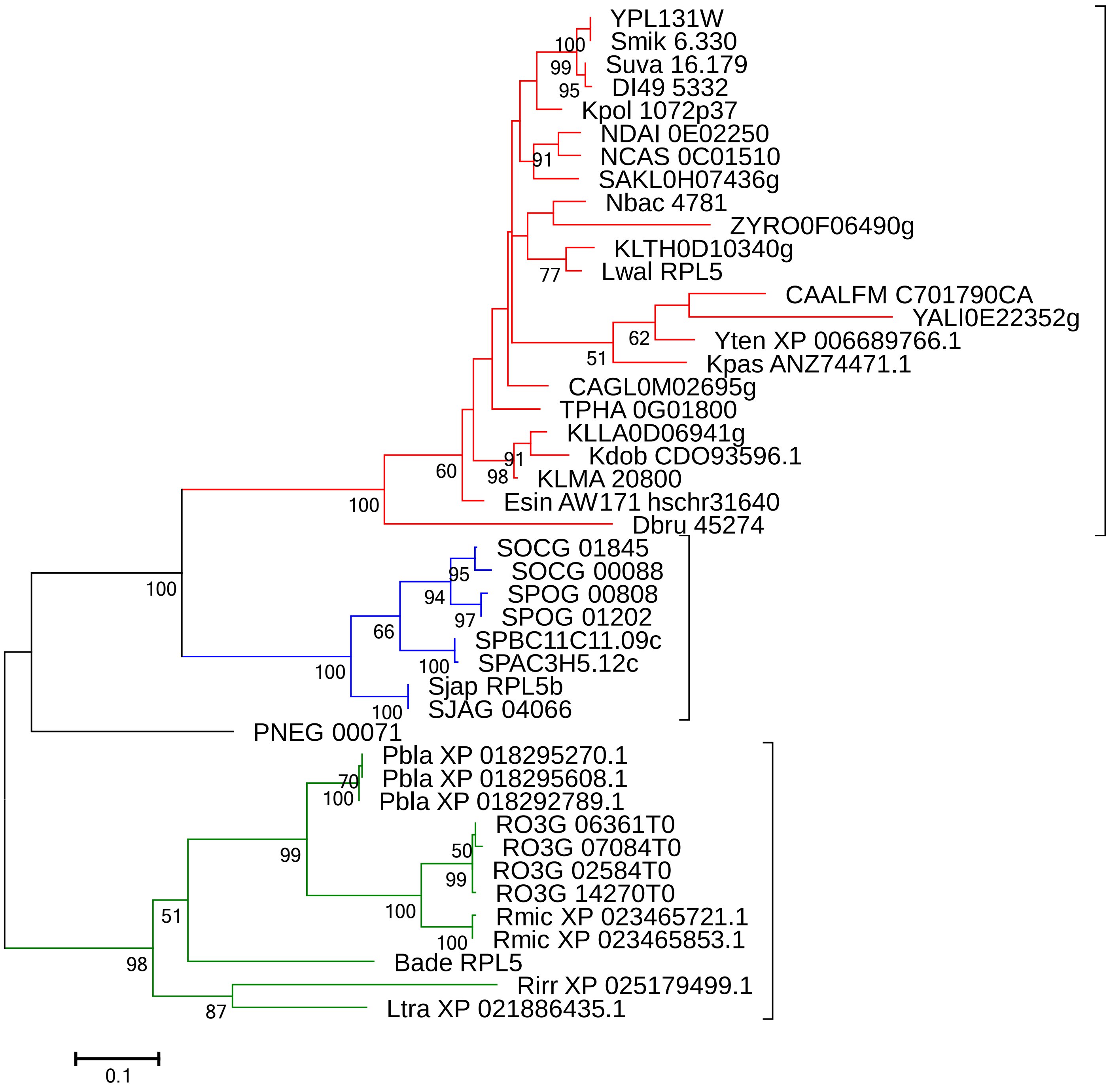
YPL131W
Smik 6.330
Suva 16.179
DI49 5332
Kpol 1072p37
NDAI 0E02250
NCAS 0C01510
SAKL0H07436g
Nbac 4781
ZYRO0F06490g
KLTH0D10340g
Lwal RPL5
CAALFM C701790CA
YALI0E22352g
Yten XP 006689766.1
Kpas ANZ74471.1
CAGL0M02695g
TPHA 0G01800
KLLA0D06941g
Kdob CDO93596.1
KLMA 20800
Esin AW171 hschr31640
Dbru 45274
SOCG 01845
SOCG 00088
SPOG 00808
SPOG 01202
SPBC11C11.09c
SPAC3H5.12c
Sjap RPL5b
SJAG 04066
PNEG 00071
Pbla XP 018295270.1
Pbla XP 018295608.1
Pbla XP 018292789.1
RO3G 06361T0
RO3G 07084T0
RO3G 02584T0
RO3G 14270T0
Rmic XP 023465721.1
Rmic XP 023465853.1
Bade RPL5
Rirr XP 025179499.1
Ltra XP 021886435.1
100
99
95
91
77
62
51
91
98
60
100
100
95
94
97
66
100
100
100
70
100
50
99
99
100
100
51
98
87
0.1

RPL6

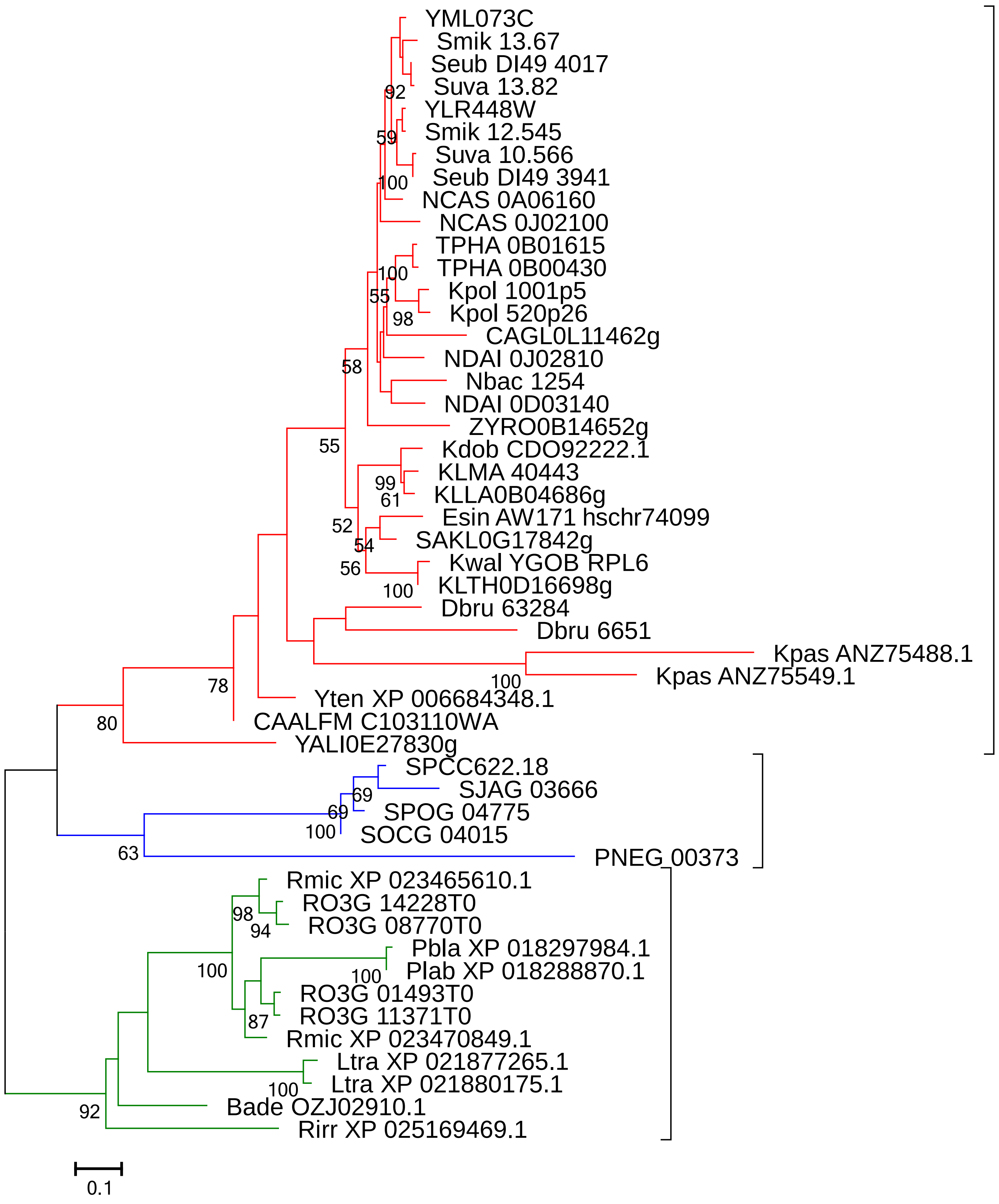
YML073C
Smik 13.67
Seub DI49 4017
Suva 13.82
92
YLR448W
Smik 12.545
59
Suva 10.566
Seub DI49 3941
100
NCAS 0A06160
NCAS 0J02100
TPHA 0B01615
TPHA 0B00430
100
Kpol 1001p5
55
Kpol 520p26
98
CAGL0L11462g
NDAI 0J02810
58
Nbac 1254
NDAI 0D03140
ZYRO0B14652g
55
Kdob CDO92222.1
KLMA 40443
99
61
KLLA0B04686g
Esin AW171 hschr74099
52
SAKL0G17842g
54
Kwal YGOB RPL6
56
KLTH0D16698g
100
Dbru 63284
Dbru 6651
Kpas ANZ75488.1
Kpas ANZ75549.1
100
78
Yten XP 006684348.1
80
CAALFM C103110WA
YALI0E27830g
SPCC622.18
SJAG 03666
69
69
SPOG 04775
100
SOCG 04015
63
PNEG 00373
Rmic XP 023465610.1
RO3G 14228T0
98
94
RO3G 08770T0
Pbla XP 018297984.1
100
Plab XP 018288870.1
100
RO3G 01493T0
RO3G 11371T0
87
Rmic XP 023470849.1
Ltra XP 021877265.1
Ltra XP 021880175.1
100
Bade OZJ02910.1
92
Rirr XP 025169469.1
0.1

RPL7

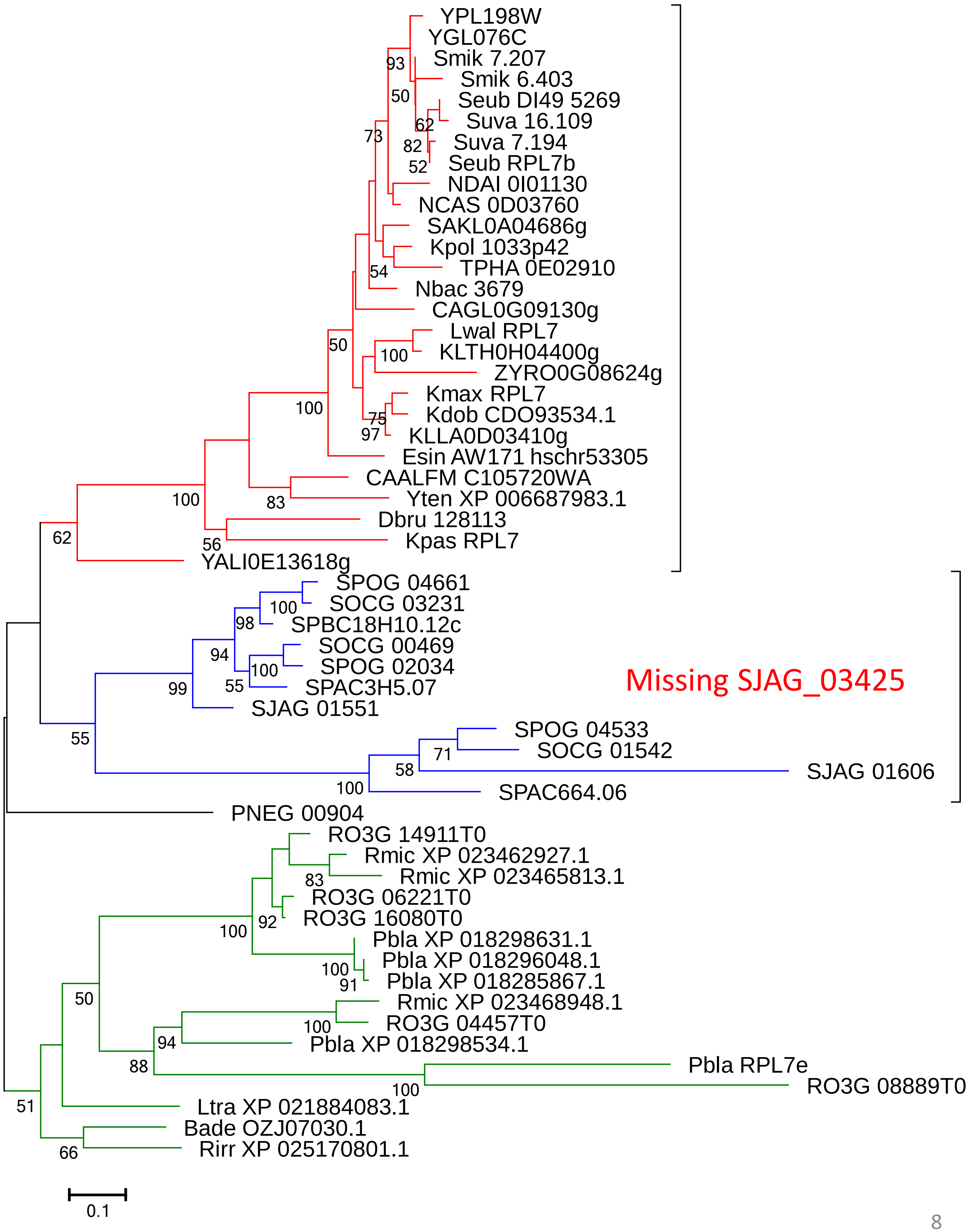
YPL198W
YGL076C
Smik 7.207
Smik 6.403
Seub DI49 5269
Suva 16.109
Suva 7.194
Seub RPL7b
NDAI 0I01130
NCAS 0D03760
SAKL0A04686g
Kpol 1033p42
TPHA 0E02910
Nbac 3679
CAGL0G09130g
Lwal RPL7
KLTH0H04400g
ZYRO0G08624g
Kmax RPL7
Kdob CDO93534.1
KLLA0D03410g
Esin AW171 hschr53305
CAALFM C105720WA
Yten XP 006687983.1
Dbru 128113
Kpas RPL7
YALI0E13618g
SPOG 04661
SOCG 03231
SPBC18H10.12c
SOCG 00469
SPOG 02034
SPAC3H5.07
SJAG 01551
Missing SJAG_03425
SPOG 04533
SOCG 01542
SJAG 01606
SPAC664.06
PNEG 00904
RO3G 14911T0
Rmic XP 023462927.1
Rmic XP 023465813.1
RO3G 06221T0
RO3G 16080T0
Pbla XP 018298631.1
Pbla XP 018296048.1
Pbla XP 018285867.1
Rmic XP 023468948.1
RO3G 04457T0
Pbla XP 018298534.1
Pbla RPL7e
RO3G 08889T0
Ltra XP 021884083.1
Bade OZJ07030.1
Rirr XP 025170801.1
0.1

### RPL8 (RPL7A)

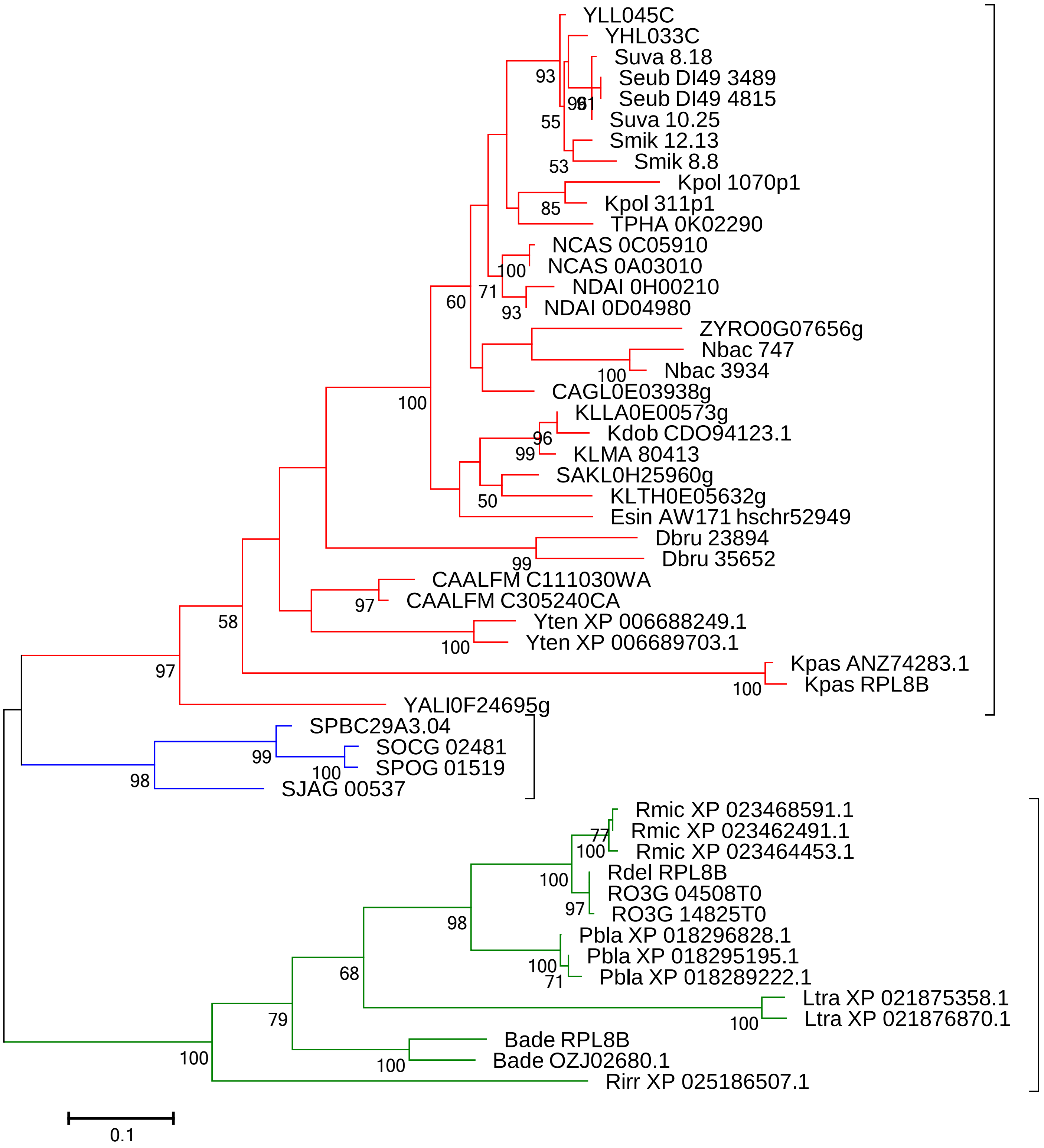
YLL045C
YHL033C
Suva 8.18
Seub DI49 3489
Seub DI49 4815
Suva 10.25
Smik 12.13
Smik 8.8
Kpol 1070p1
Kpol 311p1
TPHA 0K02290
NCAS 0C05910
NCAS 0A03010
NDAI 0H00210
NDAI 0D04980
ZYRO0G07656g
Nbac 747
Nbac 3934
CAGL0E03938g
KLLA0E00573g
Kdob CDO94123.1
KLMA 80413
SAKL0H25960g
KLTH0E05632g
Esin AW171 hschr52949
Dbru 23894
Dbru 35652
CAALFM C111030WA
CAALFM C305240CA
Yten XP 006688249.1
Yten XP 006689703.1
Kpas ANZ74283.1
Kpas RPL8B
YALI0F24695g
SPBC29A3.04
SOCG 02481
SPOG 01519
SJAG 00537
Rmic XP 023468591.1
Rmic XP 023462491.1
Rmic XP 023464453.1
Rdel RPL8B
RO3G 04508T0
RO3G 14825T0
Pbla XP 018296828.1
Pbla XP 018295195.1
Pbla XP 018289222.1
Ltra XP 021875358.1
Ltra XP 021876870.1
Bade RPL8B
Bade OZJ02680.1
Rirr XP 025186507.1
0.1

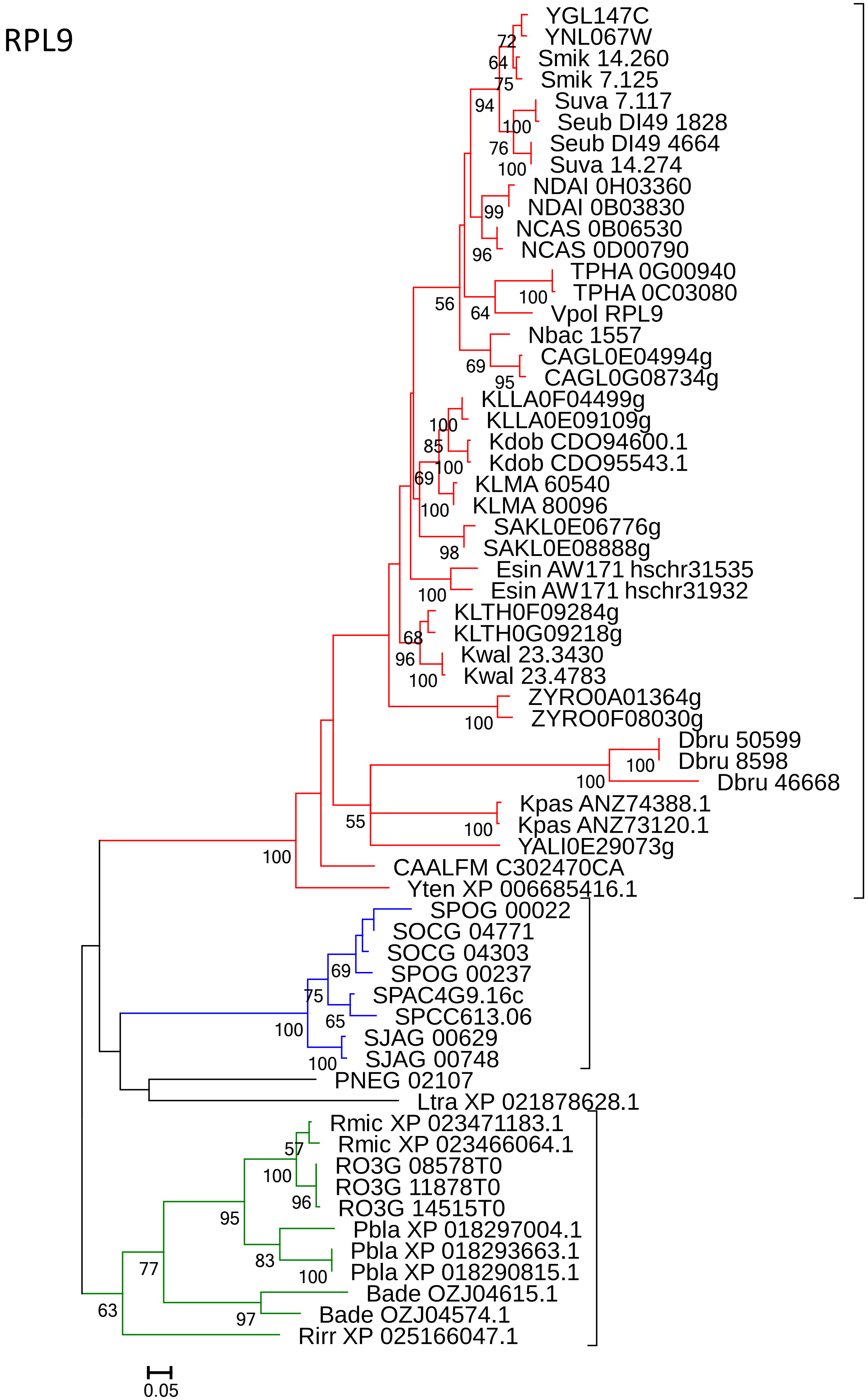
RPL9
YGL147C
YNL067W
Smik 14.260
Smik 7.125
Suva 7.117
Seub DI49 1828
Seub DI49 4664
Suva 14.274
NDAI 0H03360
NDAI 0B03830
NCAS 0B06530
NCAS 0D00790
TPHA 0G00940
TPHA 0C03080
Vpol RPL9
Nbac 1557
CAGL0E04994g
CAGL0G08734g
KLLA0F04499g
KLLA0E09109g
Kdob CDO94600.1
Kdob CDO95543.1
KLMA 60540
KLMA 80096
SAKL0E06776g
SAKL0E08888g
Esin AW171 hschr31535
Esin AW171 hschr31932
KLTH0F09284g
KLTH0G09218g
Kwal 23.3430
Kwal 23.4783
ZYRO0A01364g
ZYRO0F08030g
Dbru 50599
Dbru 8598
Dbru 46668
Kpas ANZ74388.1
Kpas ANZ73120.1
YALI0E29073g
CAALFM C302470CA
Yten XP 006685416.1
SPOG 00022
SOCG 04771
SOCG 04303
SPOG 00237
SPAC4G9.16c
SPCC613.06
SJAG 00629
SJAG 00748
PNEG 02107
Ltra XP 021878628.1
Rmic XP 023471183.1
Rmic XP 023466064.1
RO3G 08578T0
RO3G 11878T0
RO3G 14515T0
Pbla XP 018297004.1
Pbla XP 018293663.1
Pbla XP 018290815.1
Bade OZJ04615.1
Bade OZJ04574.1
Rirr XP 025166047.1
0.05

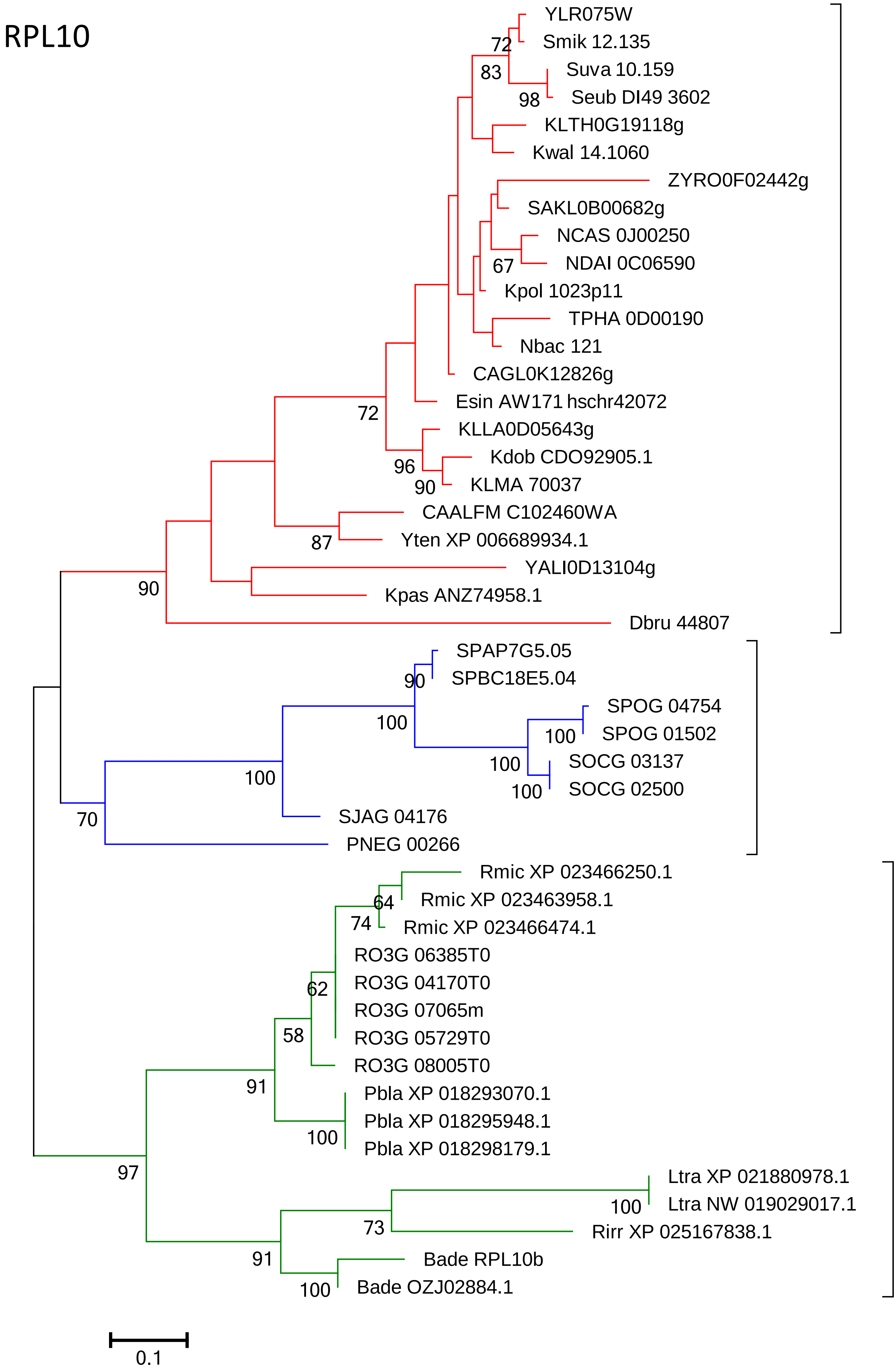
RPL10
YLR075W
Smik 12.135
Suva 10.159
Seub DI49 3602
KLTH0G19118g
Kwal 14.1060
ZYRO0F02442g
SAKL0B00682g
NCAS 0J00250
NDAI 0C06590
Kpol 1023p11
TPHA 0D00190
Nbac 121
CAGL0K12826g
Esin AW171 hschr42072
KLLA0D05643g
Kdob CDO92905.1
KLMA 70037
CAALFM C102460WA
Yten XP 006689934.1
YALI0D13104g
Kpas ANZ74958.1
Dbru 44807
SPAP7G5.05
SPBC18E5.04
SPOG 04754
SPOG 01502
SOCG 03137
SOCG 02500
SJAG 04176
PNEG 00266
Rmic XP 023466250.1
Rmic XP 023463958.1
Rmic XP 023466474.1
RO3G 06385T0
RO3G 04170T0
RO3G 07065m
RO3G 05729T0
RO3G 08005T0
Pbla XP 018293070.1
Pbla XP 018295948.1
Pbla XP 018298179.1
Ltra XP 021880978.1
Ltra NW 019029017.1
Rirr XP 025167838.1
Bade RPL10b
Bade OZJ02884.1
72
83
98
67
72
96
90
87
90
90
100
100
100
100
70
64
74
62
58
91
100
97
100
73
91
100
0.1

### RPL11

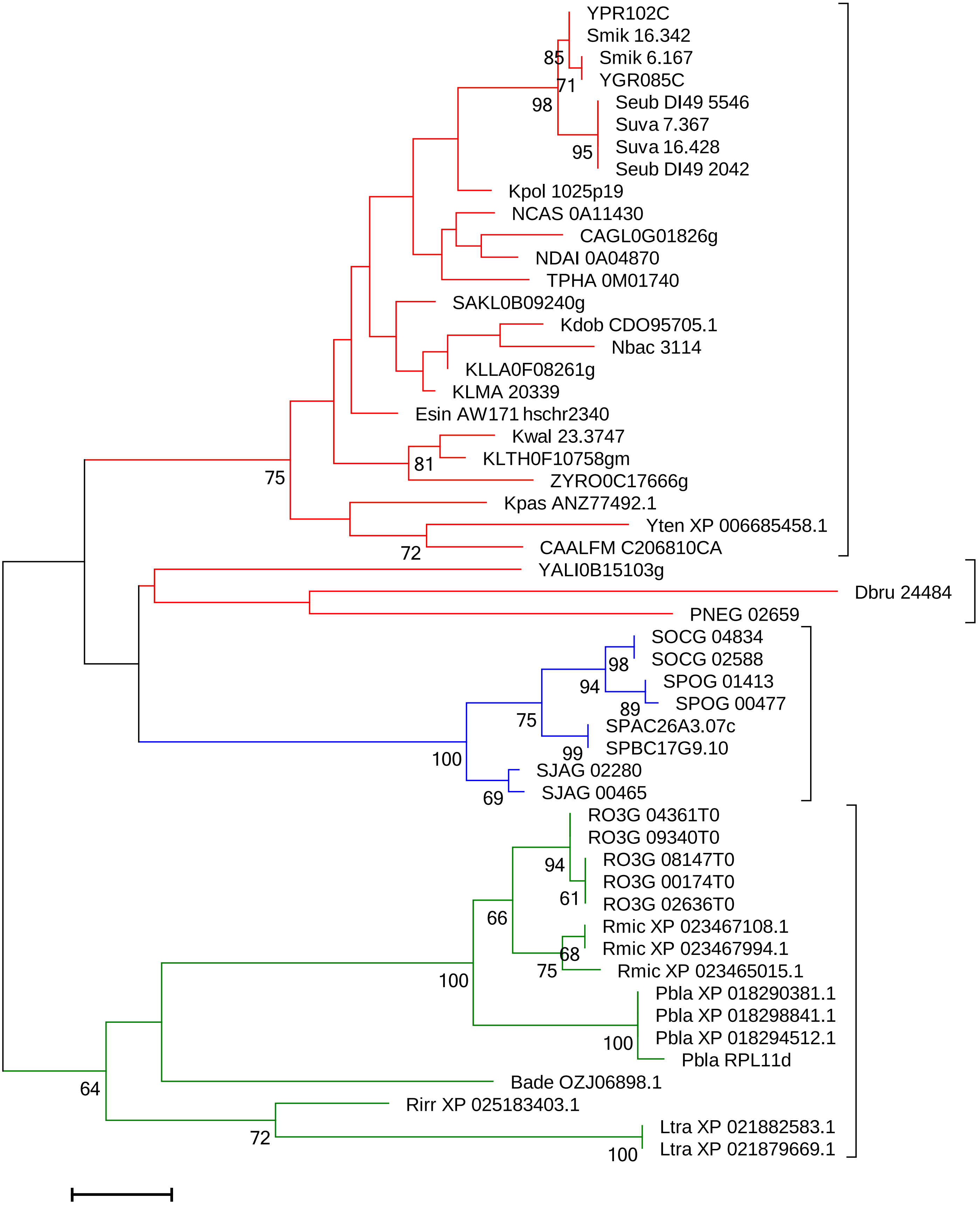

YPR102C
Smik 16.342
Smik 6.167
YGR085C
Seub DI49 5546
Suva 7.367
Suva 16.428
Seub DI49 2042
Kpol 1025p19
NCAS 0A11430
CAGL0G01826g
NDAI 0A04870
TPHA 0M01740
SAKL0B09240g
Kdob CDO95705.1
Nbac 3114
KLLA0F08261g
KLMA 20339
Esin AW171 hschr2340
Kwal 23.3747
KLTH0F10758gm
ZYRO0C17666g
Kpas ANZ77492.1
Yten XP 006685458.1
CAALFM C206810CA
YALI0B15103g
Dbru 24484
PNEG 02659
SOCG 04834
SOCG 02588
SPOG 01413
SPOG 00477
SPAC26A3.07c
SPBC17G9.10
SJAG 02280
SJAG 00465
RO3G 04361T0
RO3G 09340T0
RO3G 08147T0
RO3G 00174T0
RO3G 02636T0
Rmic XP 023467108.1
Rmic XP 023467994.1
Rmic XP 023465015.1
Pbla XP 018290381.1
Pbla XP 018298841.1
Pbla XP 018294512.1
Pbla RPL11d
Bade OZJ06898.1
Rirr XP 025183403.1
Ltra XP 021882583.1
Ltra XP 021879669.1
85
71
98
95
81
75
72
98
94
89
75
99
100
69
94
61
66
68
75
100
100
64
72
100
0.05

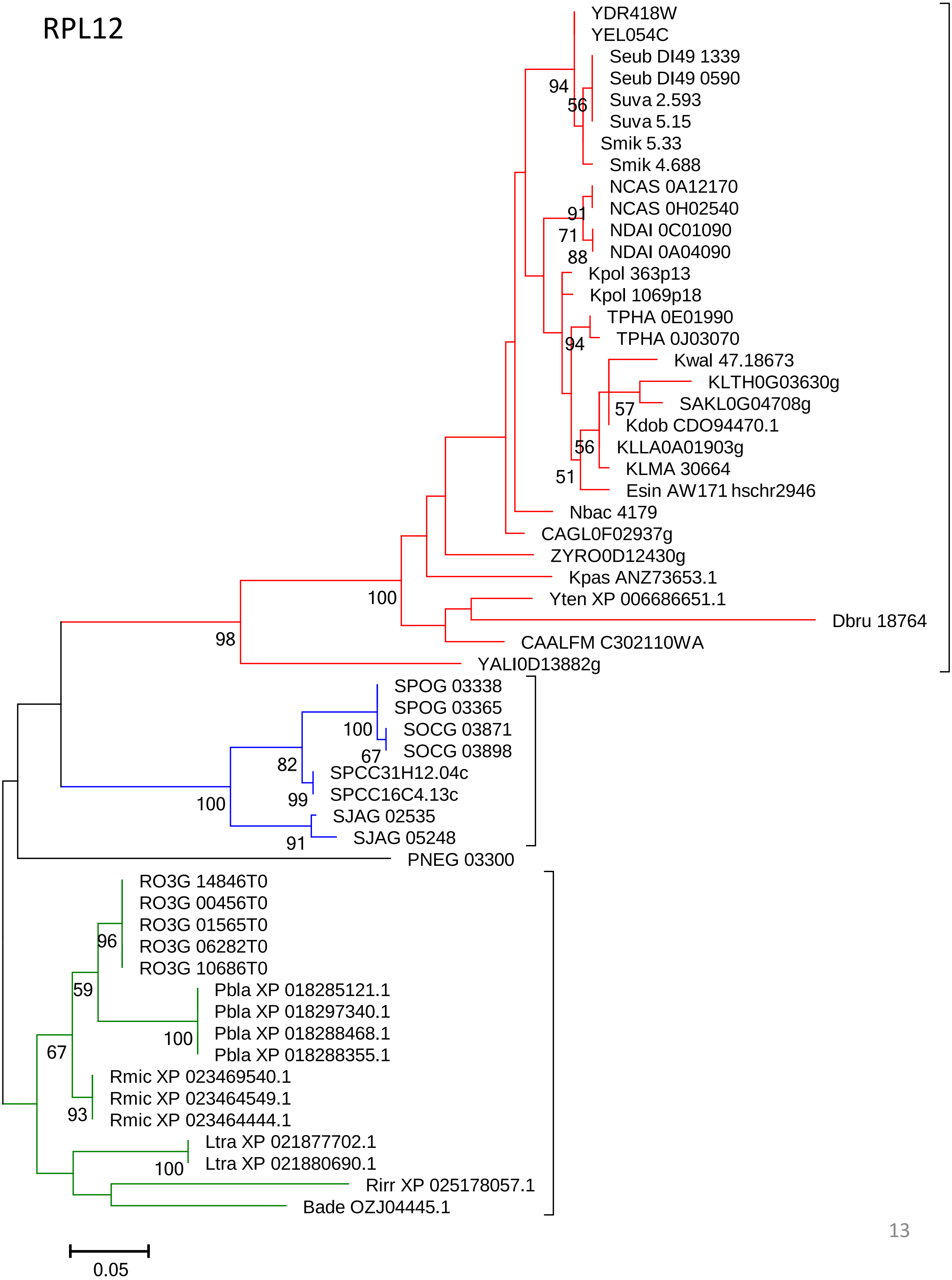
RPL12
YDR418W
YEL054C
Seub DI49 1339
Seub DI49 0590
Suva 2.593
Suva 5.15
Smik 5.33
Smik 4.688
NCAS 0A12170
NCAS 0H02540
NDAI 0C01090
NDAI 0A04090
Kpol 363p13
Kpol 1069p18
TPHA 0E01990
TPHA 0J03070
Kwal 47.18673
KLTH0G03630g
SAKL0G04708g
Kdob CDO94470.1
KLLA0A01903g
KLMA 30664
Esin AW171 hschr2946
Nbac 4179
CAGL0F02937g
ZYRO0D12430g
Kpas ANZ73653.1
Yten XP 006686651.1
Dbru 18764
CAALFM C302110WA
YALI0D13882g
SPOG 03338
SPOG 03365
SOCG 03871
SOCG 03898
SPCC31H12.04c
SPCC16C4.13c
SJAG 02535
SJAG 05248
PNEG 03300
RO3G 14846T0
RO3G 00456T0
RO3G 01565T0
RO3G 06282T0
RO3G 10686T0
Pbla XP 018285121.1
Pbla XP 018297340.1
Pbla XP 018288468.1
Pbla XP 018288355.1
Rmic XP 023469540.1
Rmic XP 023464549.1
Rmic XP 023464444.1
Ltra XP 021877702.1
Ltra XP 021880690.1
Rirr XP 025178057.1
Bade OZJ04445.1
94
56
91
71
88
94
57
56
51
100
98
100
67
82
99
100
91
96
59
100
67
93
100
0.05

### RPL13

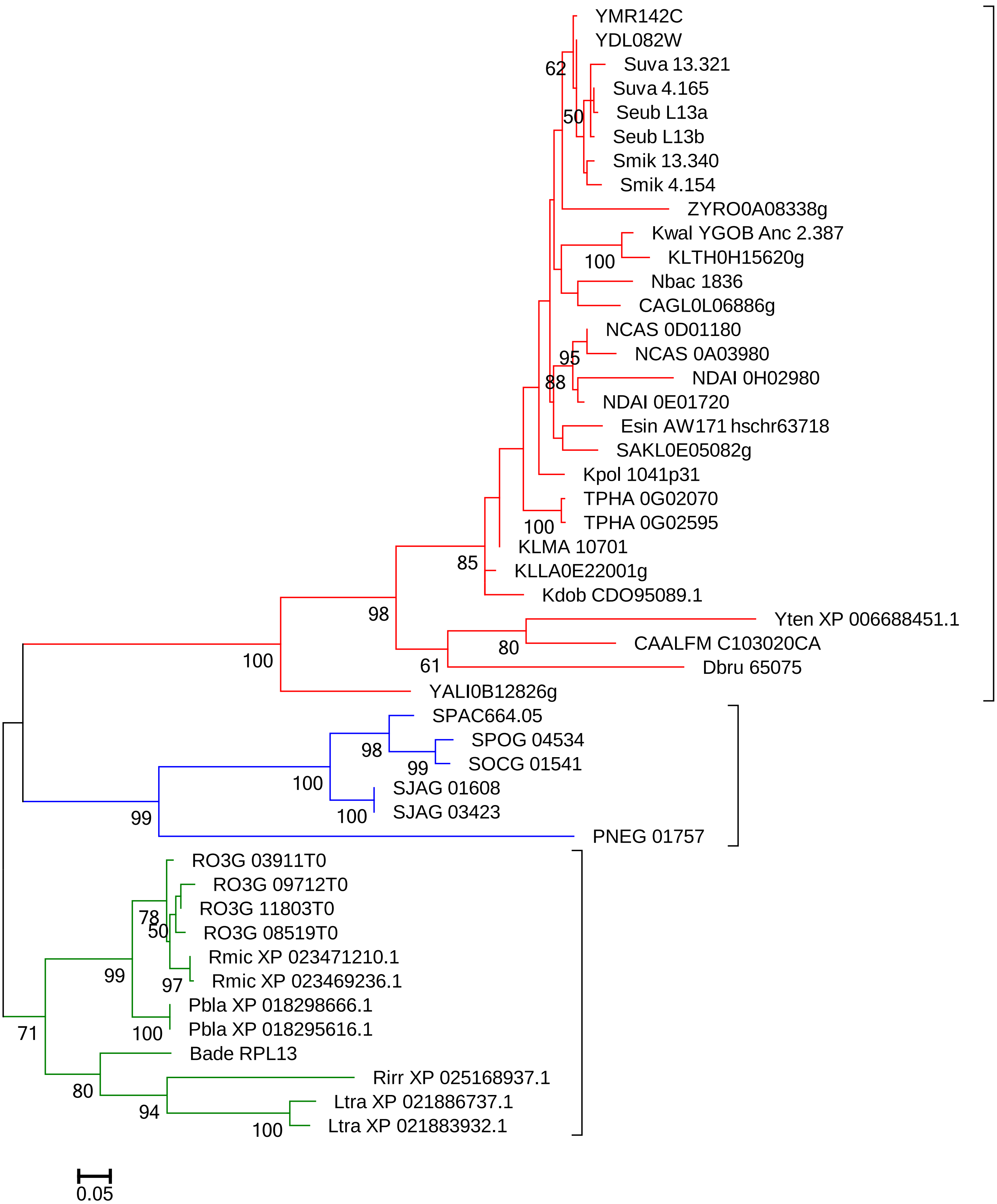
YMR142C
YDL082W
Suva 13.321
Suva 4.165
Seub L13a
Seub L13b
Smik 13.340
Smik 4.154
ZYRO0A08338g
Kwal YGOB Anc 2.387
KLTH0H15620g
Nbac 1836
CAGL0L06886g
NCAS 0D01180
NCAS 0A03980
NDAI 0H02980
NDAI 0E01720
Esin AW171 hschr63718
SAKL0E05082g
Kpol 1041p31
TPHA 0G02070
TPHA 0G02595
KLMA 10701
KLLA0E22001g
Kdob CDO95089.1
Yten XP 006688451.1
CAALFM C103020CA
Dbru 65075
YALI0B12826g
SPAC664.05
SPOG 04534
SOCG 01541
SJAG 01608
SJAG 03423
PNEG 01757
RO3G 03911T0
RO3G 09712T0
RO3G 11803T0
RO3G 08519T0
Rmic XP 023471210.1
Rmic XP 023469236.1
Pbla XP 018298666.1
Pbla XP 018295616.1
Bade RPL13
Rirr XP 025168937.1
Ltra XP 021886737.1
Ltra XP 021883932.1
62
50
100
95
88
100
85
98
80
61
100
98
99
100
100
99
78
50
99
97
100
71
80
94
100
0.05

### RPL14

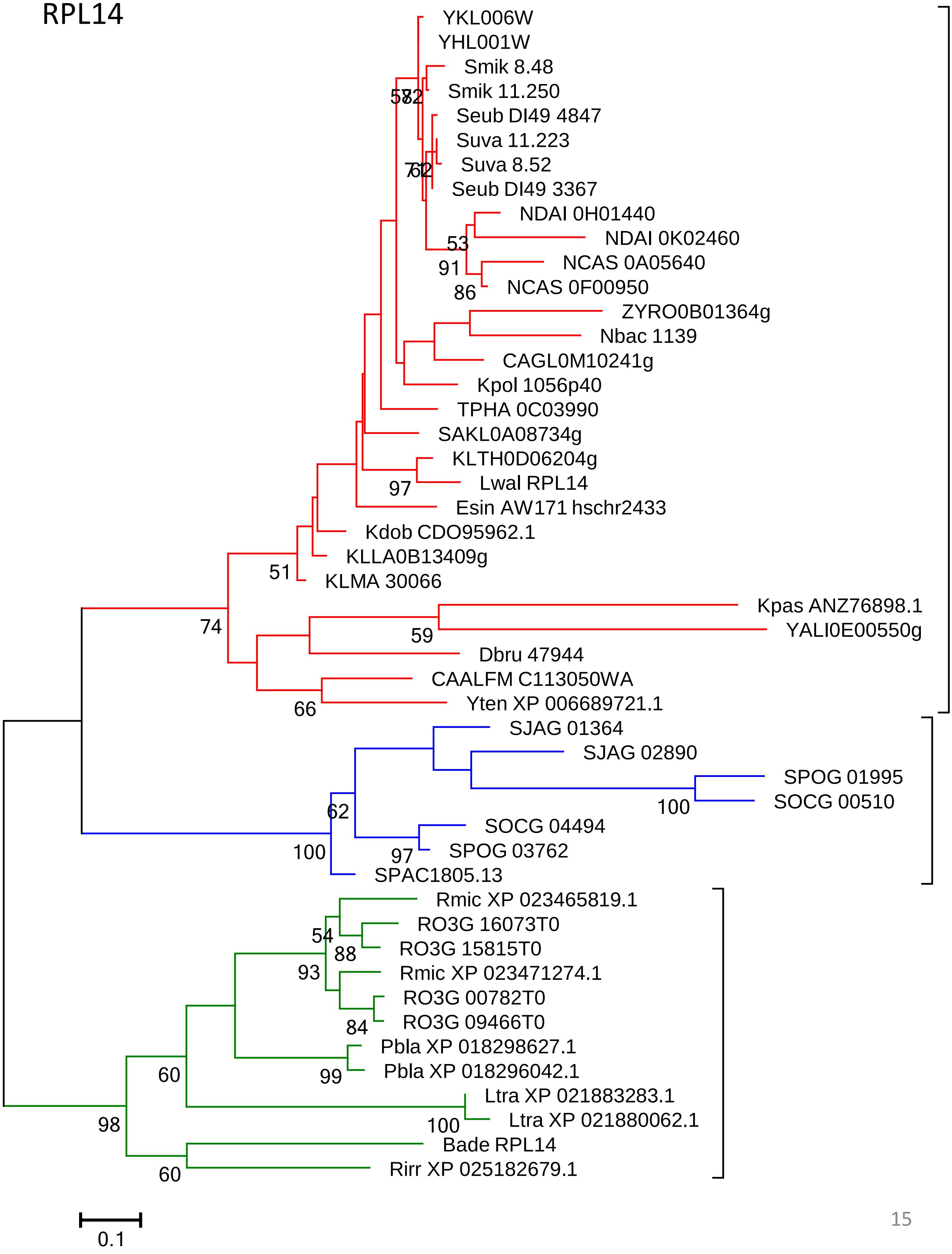

YKL006W
YHL001W
Smik 8.48
Smik 11.250
Seub DI49 4847
Suva 11.223
Suva 8.52
Seub DI49 3367
NDAI 0H01440
NDAI 0K02460
53
91
NCAS 0A05640
86
NCAS 0F00950
ZYRO0B01364g
Nbac 1139
CAGL0M10241g
Kpol 1056p40
TPHA 0C03990
SAKL0A08734g
KLTH0D06204g
97
Lwal RPL14
Esin AW171 hschr2433
Kdob CDO95962.1
KLLA0B13409g
51
KLMA 30066
Kpas ANZ76898.1
74
59
YALI0E00550g
Dbru 47944
CAALFM C113050WA
66
Yten XP 006689721.1
SJAG 01364
SJAG 02890
SPOG 01995
100
SOCG 00510
62
SOCG 04494
100
97
SPOG 03762
SPAC1805.13
Rmic XP 023465819.1
RO3G 16073T0
54
88
RO3G 15815T0
93
Rmic XP 023471274.1
RO3G 00782T0
84
RO3G 09466T0
Pbla XP 018298627.1
99
Pbla XP 018296042.1
60
Ltra XP 021883283.1
98
100
Ltra XP 021880062.1
Bade RPL14
60
Rirr XP 025182679.1
0.1

### RPL15

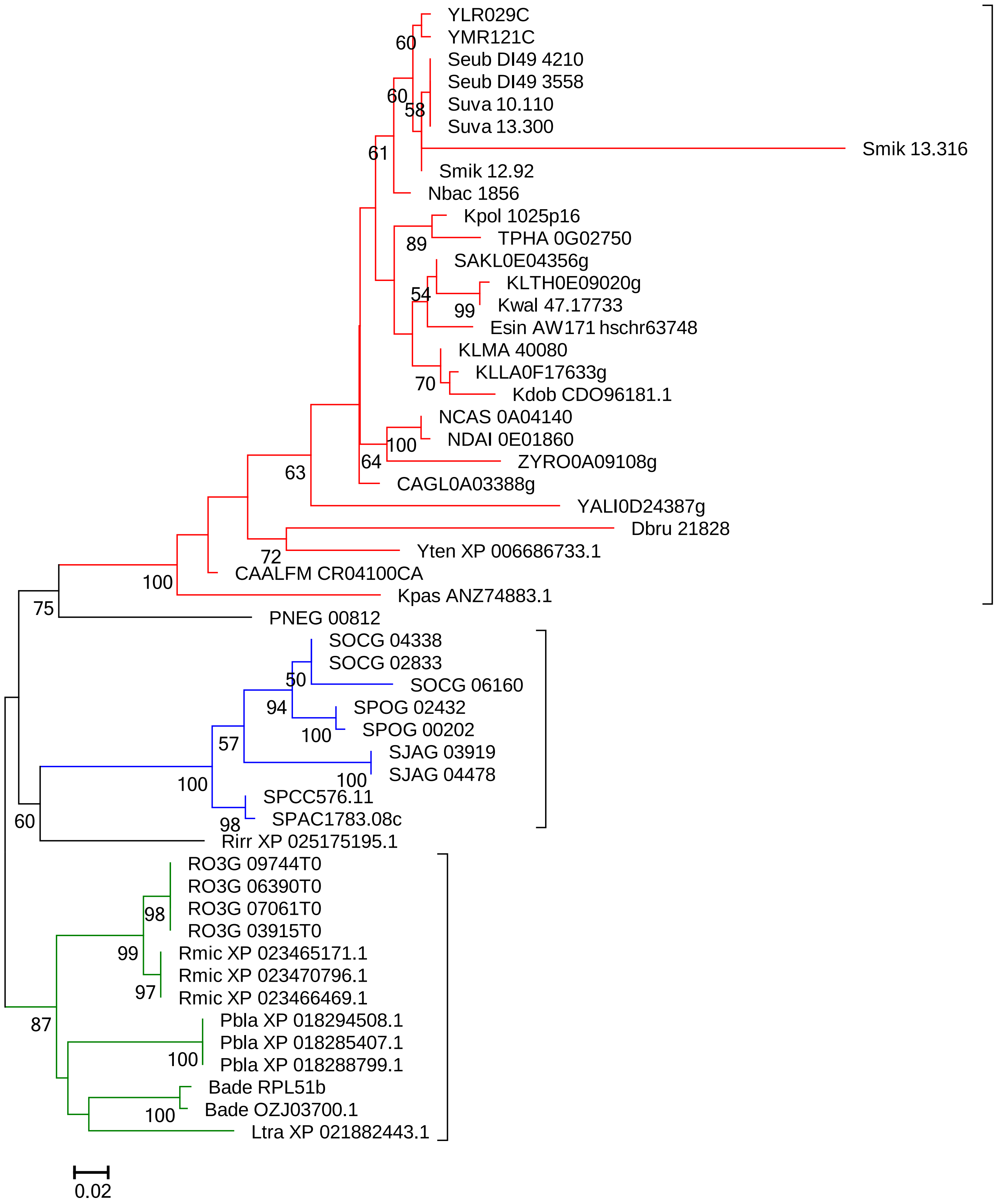
YLR029C
YMR121C
Seub DI49 4210
Seub DI49 3558
Suva 10.110
Suva 13.300
Smik 13.316
Smik 12.92
Nbac 1856
Kpol 1025p16
TPHA 0G02750
SAKL0E04356g
KLTH0E09020g
Kwal 47.17733
Esin AW171 hschr63748
KLMA 40080
KLLA0F17633g
Kdob CDO96181.1
NCAS 0A04140
NDAI 0E01860
ZYRO0A09108g
CAGL0A03388g
YALI0D24387g
Dbru 21828
Yten XP 006686733.1
CAALFM CR04100CA
Kpas ANZ74883.1
PNEG 00812
SOCG 04338
SOCG 02833
SOCG 06160
SPOG 02432
SPOG 00202
SJAG 03919
SJAG 04478
SPCC576.11
SPAC1783.08c
Rirr XP 025175195.1
RO3G 09744T0
RO3G 06390T0
RO3G 07061T0
RO3G 03915T0
Rmic XP 023465171.1
Rmic XP 023470796.1
Rmic XP 023466469.1
Pbla XP 018294508.1
Pbla XP 018285407.1
Pbla XP 018288799.1
Bade RPL51b
Bade OZJ03700.1
Ltra XP 021882443.1
0.02

### RPL16 (RPL13A)

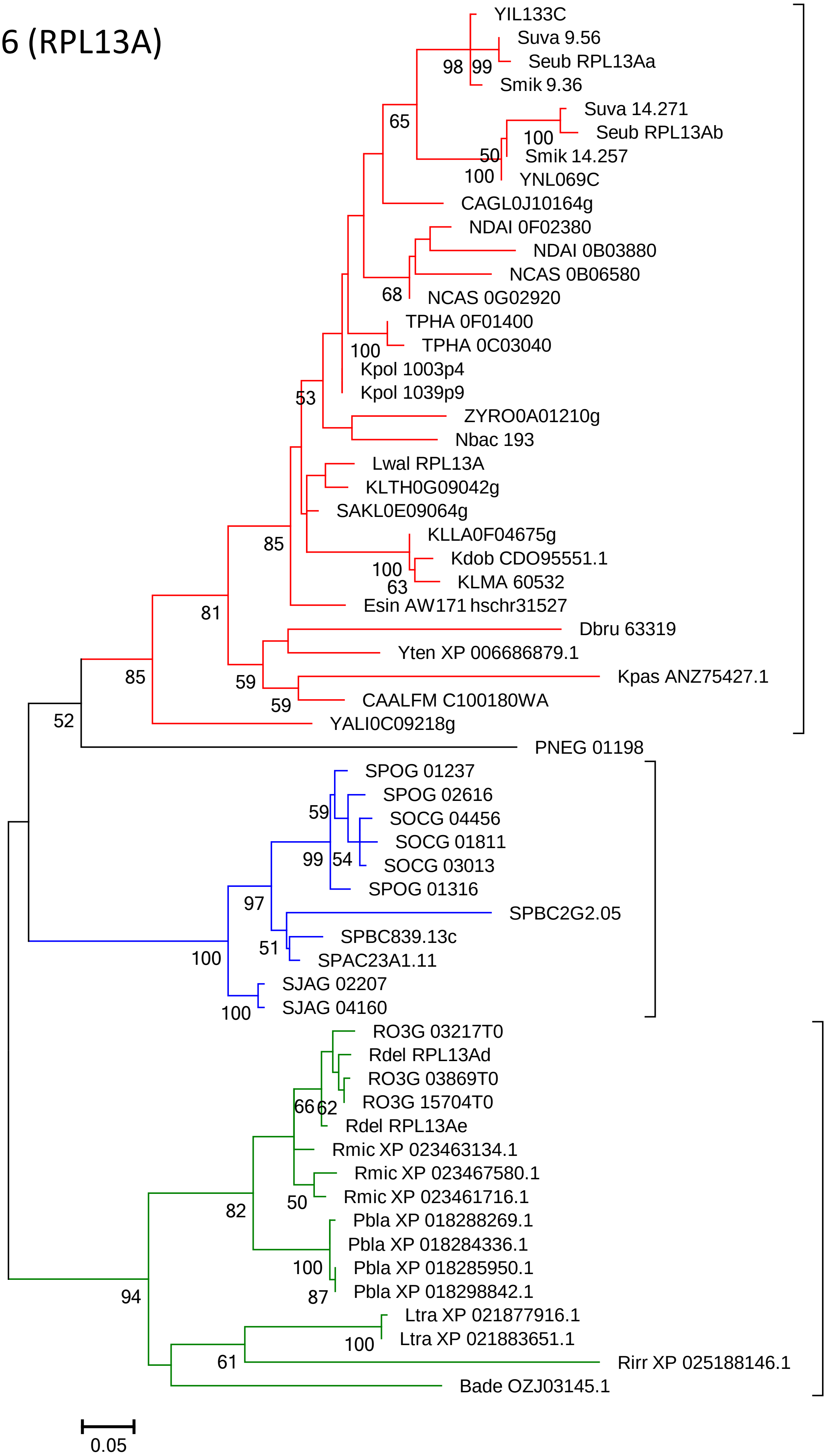

### RPL17

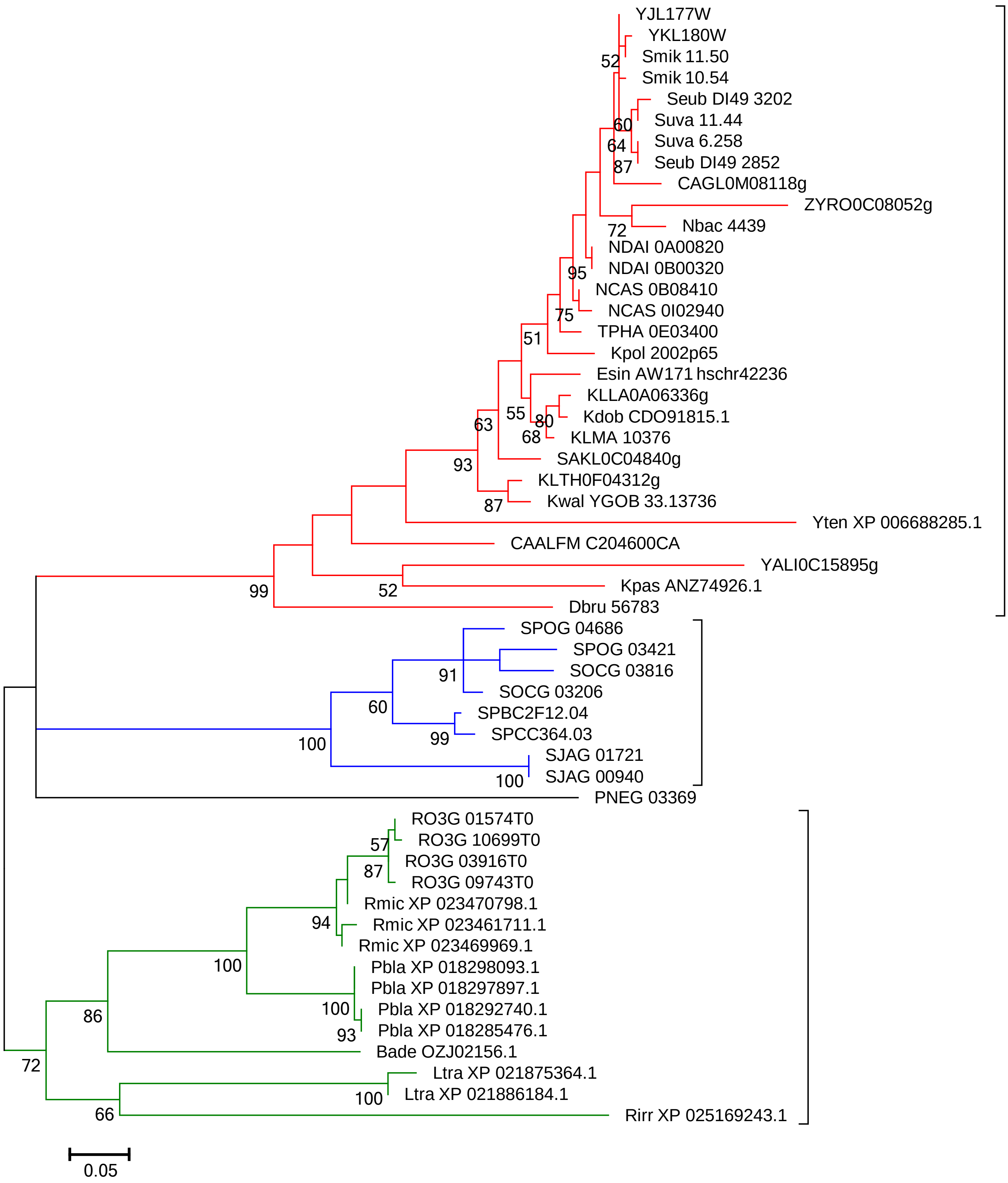
YJL177W
YKL180W
Smik 11.50
Smik 10.54
Seub DI49 3202
Suva 11.44
Suva 6.258
Seub DI49 2852
CAGL0M08118g
ZYRO0C08052g
Nbac 4439
NDAI 0A00820
NDAI 0B00320
NCAS 0B08410
NCAS 0I02940
TPHA 0E03400
Kpol 2002p65
Esin AW171 hschr42236
KLLA0A06336g
Kdob CDO91815.1
KLMA 10376
SAKL0C04840g
KLTH0F04312g
Kwal YGOB 33.13736
Yten XP 006688285.1
CAALFM C204600CA
YALI0C15895g
Kpas ANZ74926.1
Dbru 56783
SPOG 04686
SPOG 03421
SOCG 03816
SOCG 03206
SPBC2F12.04
SPCC364.03
SJAG 01721
SJAG 00940
PNEG 03369
RO3G 01574T0
RO3G 10699T0
RO3G 03916T0
RO3G 09743T0
Rmic XP 023470798.1
Rmic XP 023461711.1
Rmic XP 023469969.1
Pbla XP 018298093.1
Pbla XP 018297897.1
Pbla XP 018292740.1
Pbla XP 018285476.1
Bade OZJ02156.1
Ltra XP 021875364.1
Ltra XP 021886184.1
Rirr XP 025169243.1
52
60
64
87
72
95
75
51
55
80
68
63
93
87
99
52
91
60
99
100
100
57
87
94
100
100
93
86
72
100
66
0.05

RPL18

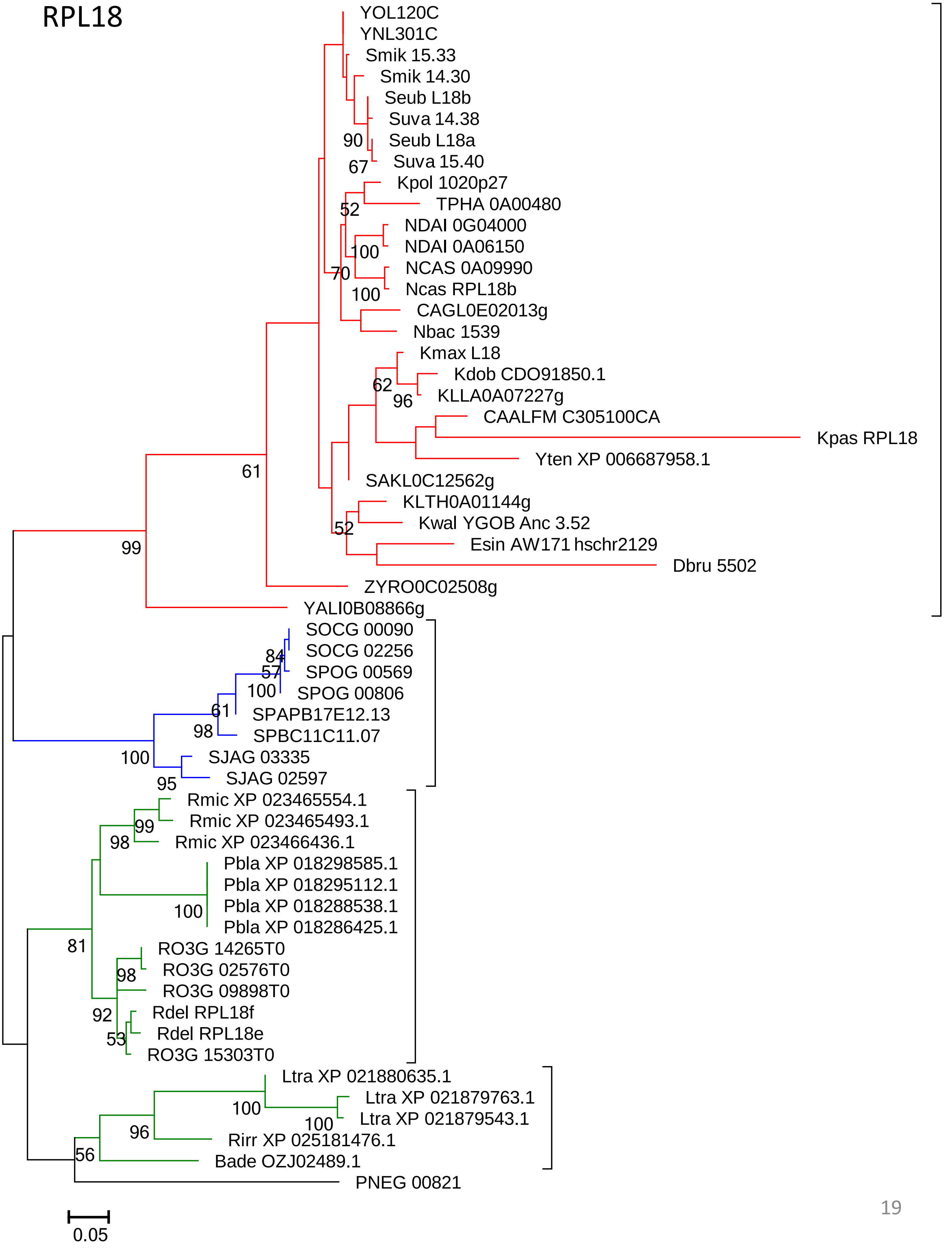
YOL120C
YNL301C
Smik 15.33
Smik 14.30
Seub L18b
Suva 14.38
Seub L18a
Suva 15.40
Kpol 1020p27
TPHA 0A00480
NDAI 0G04000
NDAI 0A06150
NCAS 0A09990
Ncas RPL18b
CAGL0E02013g
Nbac 1539
Kmax L18
Kdob CDO91850.1
KLLA0A07227g
CAALFM C305100CA
Kpas RPL18
Yten XP 006687958.1
SAKL0C12562g
KLTH0A01144g
Kwal YGOB Anc 3.52
Esin AW171 hschr2129
Dbru 5502
ZYRO0C02508g
YALI0B08866g
SOCG 00090
SOCG 02256
SPOG 00569
SPOG 00806
SPAPB17E12.13
SPBC11C11.07
SJAG 03335
SJAG 02597
Rmic XP 023465554.1
Rmic XP 023465493.1
Rmic XP 023466436.1
Pbla XP 018298585.1
Pbla XP 018295112.1
Pbla XP 018288538.1
Pbla XP 018286425.1
RO3G 14265T0
RO3G 02576T0
RO3G 09898T0
Rdel RPL18f
Rdel RPL18e
RO3G 15303T0
Ltra XP 021880635.1
Ltra XP 021879763.1
Ltra XP 021879543.1
Rirr XP 025181476.1
Bade OZJ02489.1
PNEG 00821
90
67
52
100
70
100
62
96
61
52
99
84
57
100
61
98
100
95
99
98
100
81
98
92
53
100
100
96
56
0.05

RPL19

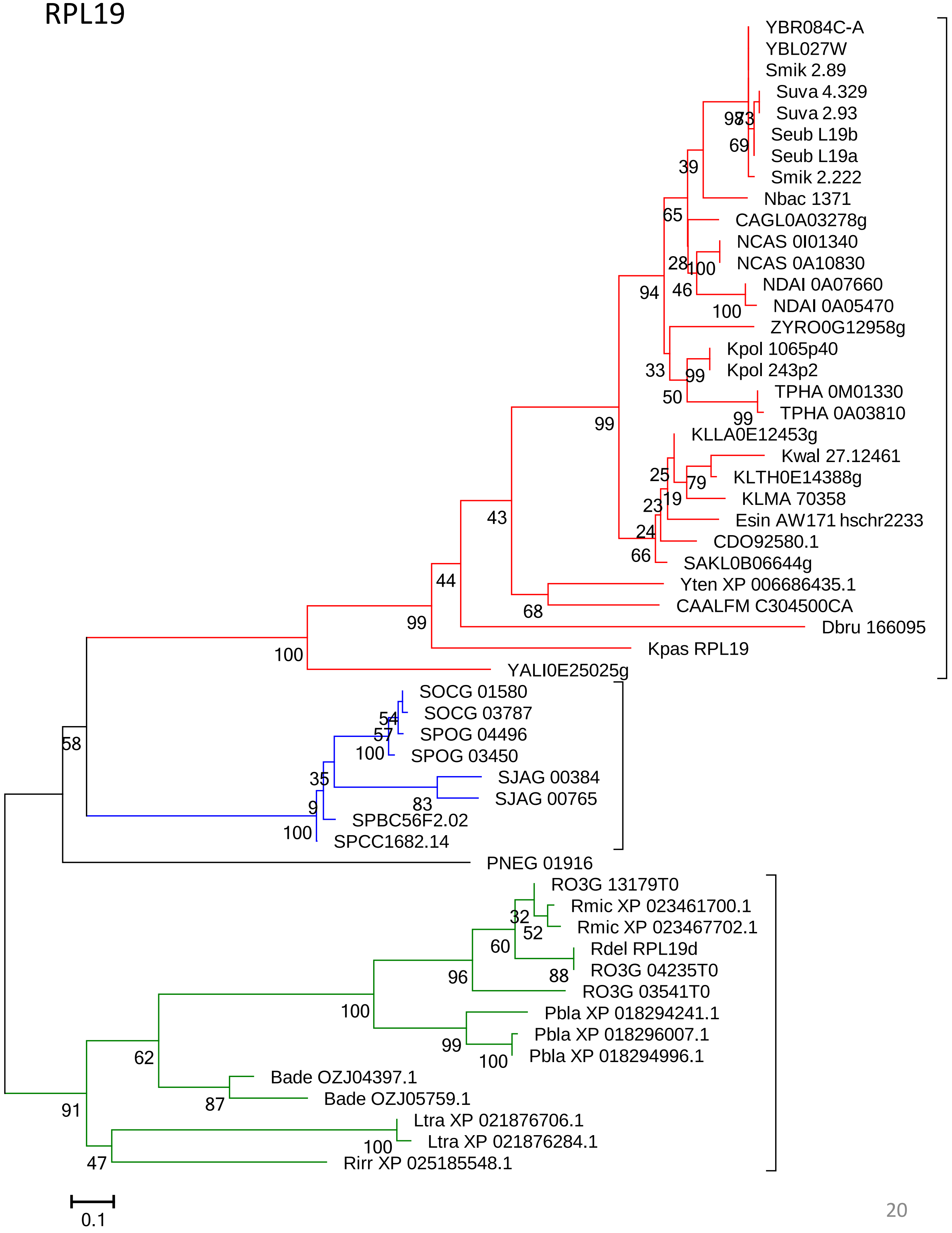
YBR084C-A
YBL027W
Smik 2.89
Suva 4.329
Suva 2.93
Seub L19b
Seub L19a
Smik 2.222
Nbac 1371
CAGL0A03278g
NCAS 0I01340
NCAS 0A10830
NDAI 0A07660
NDAI 0A05470
ZYRO0G12958g
Kpol 1065p40
Kpol 243p2
TPHA 0M01330
TPHA 0A03810
KLLA0E12453g
Kwal 27.12461
KLTH0E14388g
KLMA 70358
Esin AW171 hschr2233
CDO92580.1
SAKL0B06644g
Yten XP 006686435.1
CAALFM C304500CA
Dbru 166095
Kpas RPL19
YALI0E25025g
SOCG 01580
SOCG 03787
SPOG 04496
SPOG 03450
SJAG 00384
SJAG 00765
SPBC56F2.02
SPCC1682.14
PNEG 01916
RO3G 13179T0
Rmic XP 023461700.1
Rmic XP 023467702.1
Rdel RPL19d
RO3G 04235T0
RO3G 03541T0
Pbla XP 018294241.1
Pbla XP 018296007.1
Pbla XP 018294996.1
Bade OZJ04397.1
Bade OZJ05759.1
Ltra XP 021876706.1
Ltra XP 021876284.1
Rirr XP 025185548.1
0.1

### RPL20 (RPL18A)

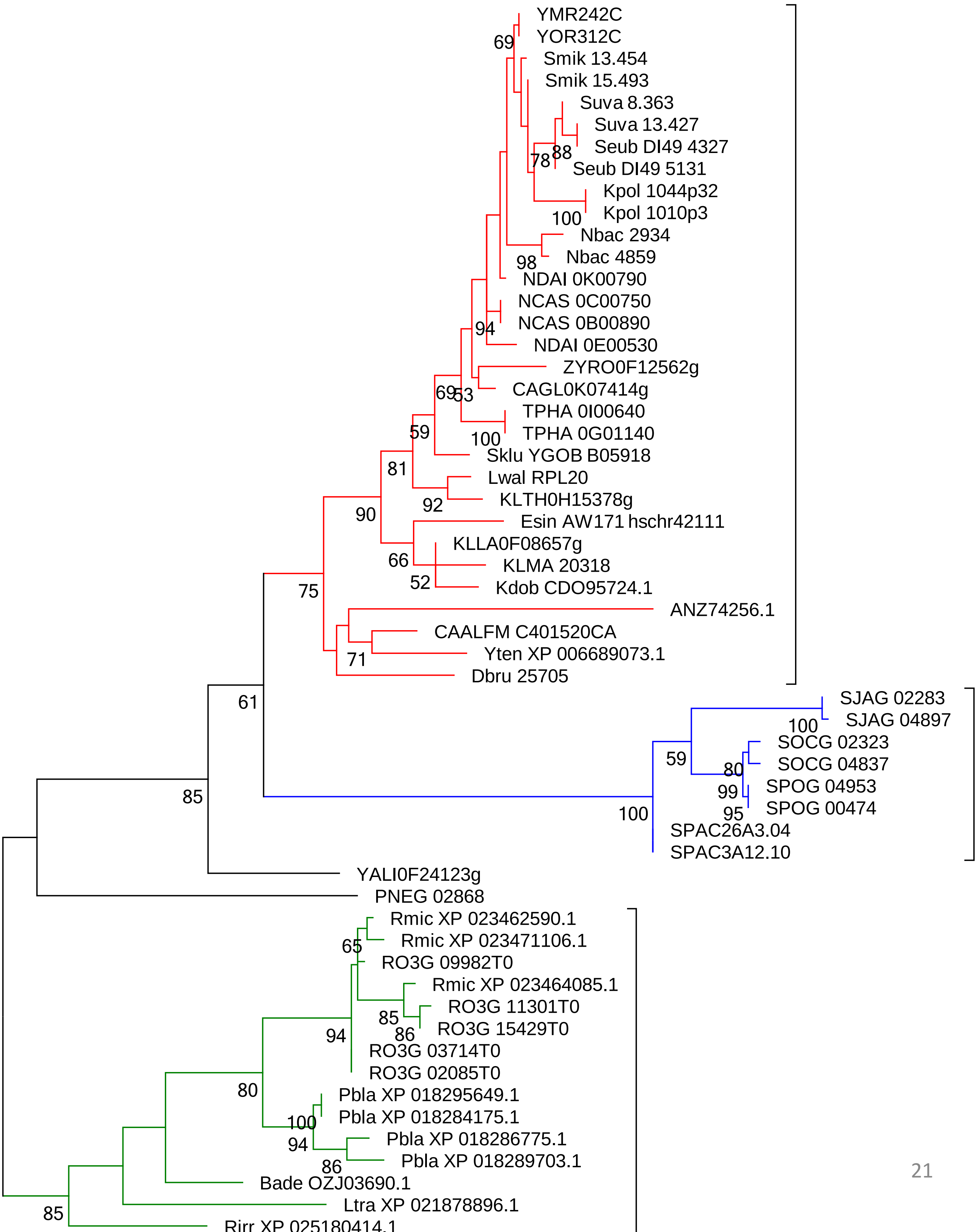

#### RPL21

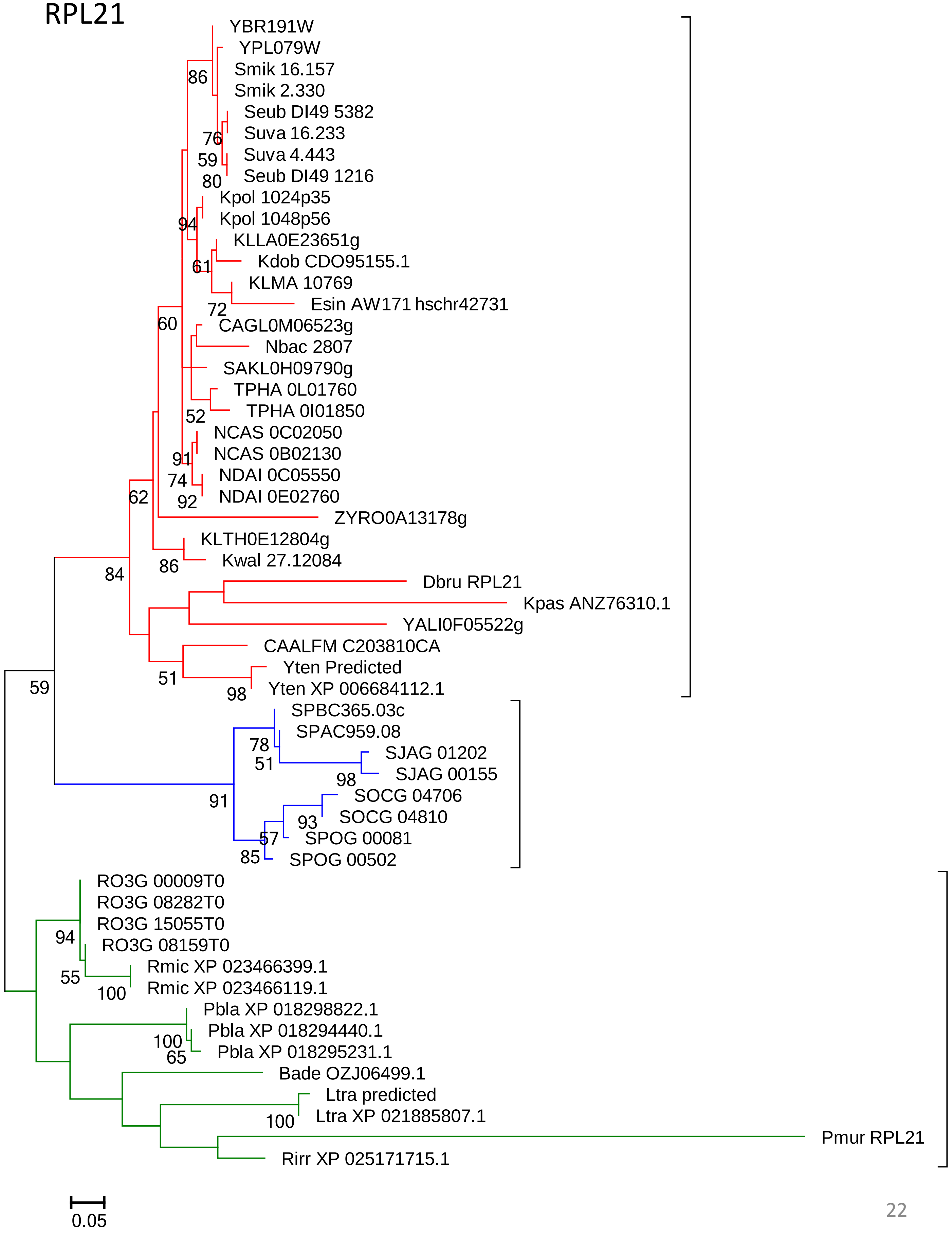

YBR191W
YPL079W
Smik 16.157
Smik 2.330
Seub DI49 5382
Suva 16.233
Suva 4.443
Seub DI49 1216
Kpol 1024p35
Kpol 1048p56
KLLA0E23651g
Kdob CDO95155.1
KLMA 10769
Esin AW171 hschr42731
CAGL0M06523g
Nbac 2807
SAKL0H09790g
TPHA 0L01760
TPHA 0I01850
NCAS 0C02050
NCAS 0B02130
NDAI 0C05550
NDAI 0E02760
ZYRO0A13178g
KLTH0E12804g
Kwal 27.12084
Dbru RPL21
Kpas ANZ76310.1
YALI0F05522g
CAALFM C203810CA
Yten Predicted
Yten XP 006684112.1
SPBC365.03c
SPAC959.08
SJAG 01202
SJAG 00155
SOCG 04706
SOCG 04810
SPOG 00081
SPOG 00502
RO3G 00009T0
RO3G 08282T0
RO3G 15055T0
RO3G 08159T0
Rmic XP 023466399.1
Rmic XP 023466119.1
Pbla XP 018298822.1
Pbla XP 018294440.1
Pbla XP 018295231.1
Bade OZJ06499.1
Ltra predicted
Ltra XP 021885807.1
Pmur RPL21
Rirr XP 025171715.1
86
76
59
80
94
61
72
60
52
91
74
92
62
86
84
51
98
59
78
51
98
91
93
57
85
94
55
100
100
65
100
0.05

### RPL22

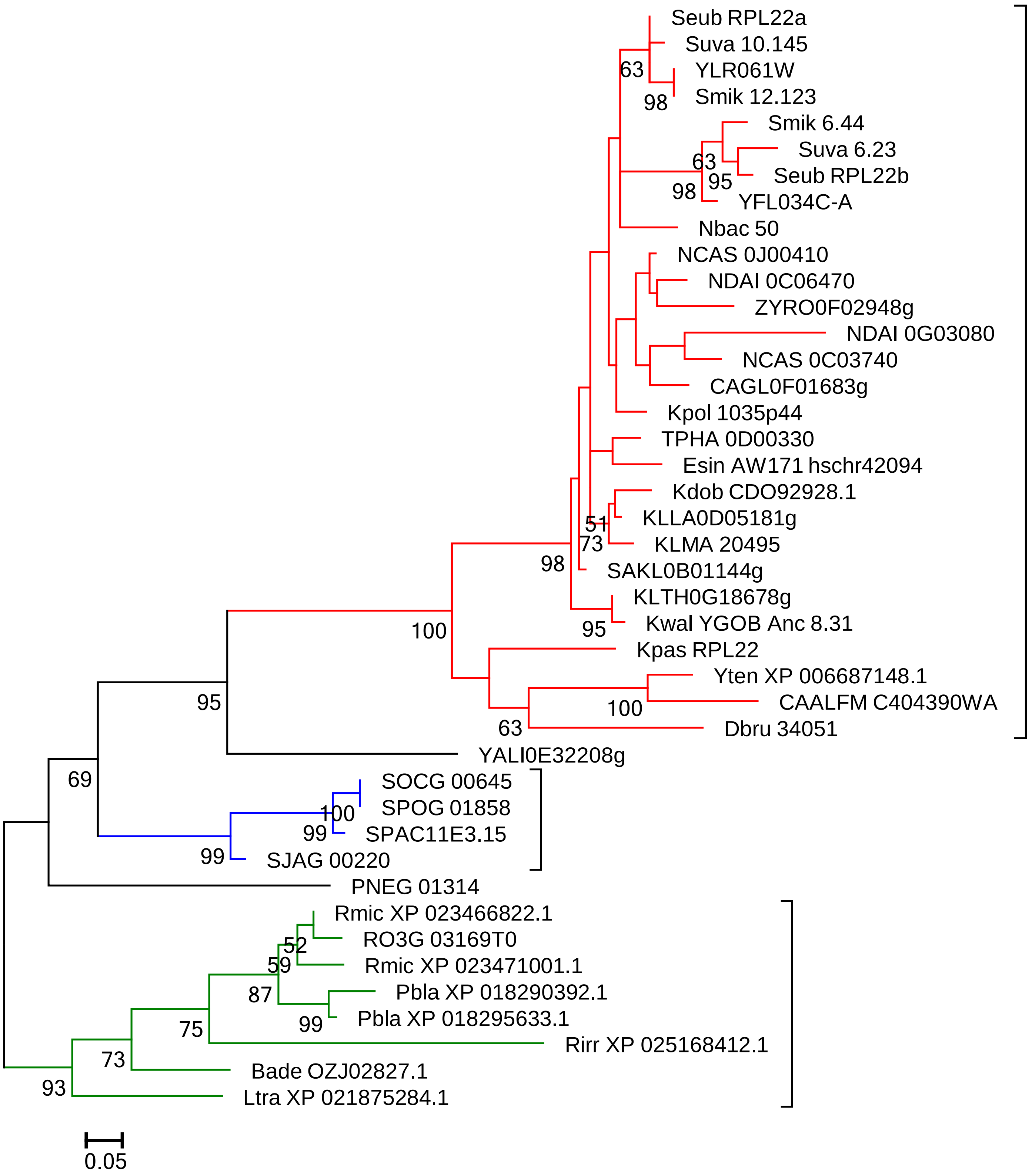
Seub RPL22a
Suva 10.145
YLR061W
Smik 12.123
Smik 6.44
Suva 6.23
Seub RPL22b
YFL034C-A
Nbac 50
NCAS 0J00410
NDAI 0C06470
ZYRO0F02948g
NDAI 0G03080
NCAS 0C03740
CAGL0F01683g
Kpol 1035p44
TPHA 0D00330
Esin AW171 hschr42094
Kdob CDO92928.1
KLLA0D05181g
KLMA 20495
SAKL0B01144g
KLTH0G18678g
Kwal YGOB Anc 8.31
Kpas RPL22
Yten XP 006687148.1
CAALFM C404390WA
Dbru 34051
YALI0E32208g
SOCG 00645
SPOG 01858
SPAC11E3.15
SJAG 00220
PNEG 01314
Rmic XP 023466822.1
RO3G 03169T0
Rmic XP 023471001.1
Pbla XP 018290392.1
Pbla XP 018295633.1
Rirr XP 025168412.1
Bade OZJ02827.1
Ltra XP 021875284.1
63
98
63
95
98
51
73
98
95
100
100
63
95
69
100
99
99
52
59
87
99
75
73
93
0.05

### RPL23

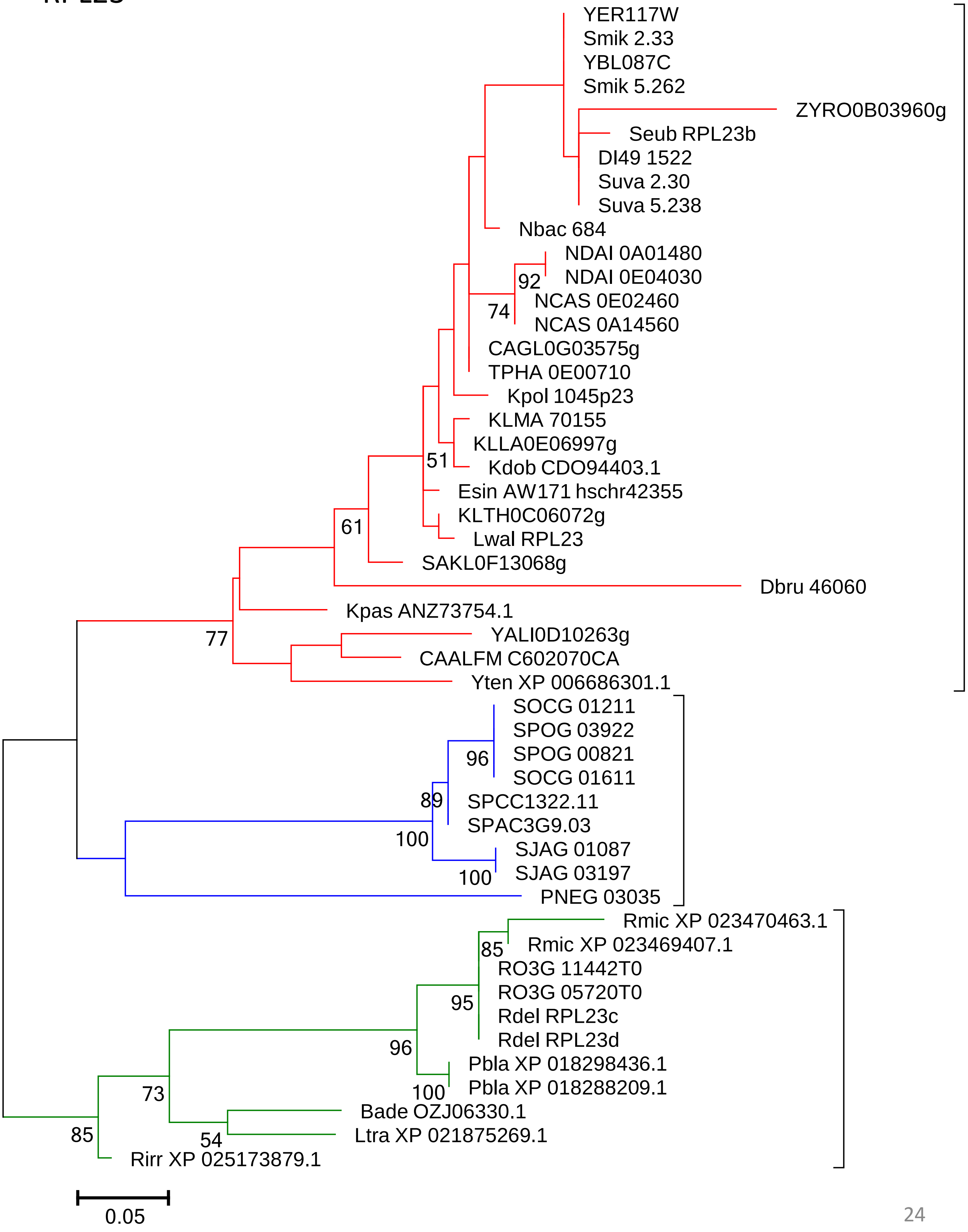

YER117W
Smik 2.33
YBL087C
Smik 5.262
ZYRO0B03960g
Seub RPL23b
DI49 1522
Suva 2.30
Suva 5.238
Nbac 684
NDAI 0A01480
92
NDAI 0E04030
NCAS 0E02460
74
NCAS 0A14560
CAGL0G03575g
TPHA 0E00710
Kpol 1045p23
KLMA 70155
KLLA0E06997g
51
Kdob CDO94403.1
Esin AW171 hschr42355
KLTH0C06072g
61
Lwal RPL23
SAKL0F13068g
Dbru 46060
Kpas ANZ73754.1
YALI0D10263g
77
CAALFM C602070CA
Yten XP 006686301.1
SOCG 01211
SPOG 03922
96
SPOG 00821
SOCG 01611
89
SPCC1322.11
SPAC3G9.03
100
SJAG 01087
100
SJAG 03197
PNEG 03035
Rmic XP 023470463.1
85
Rmic XP 023469407.1
RO3G 11442T0
95
RO3G 05720T0
Rdel RPL23c
Rdel RPL23d
96
Pbla XP 018298436.1
100
Pbla XP 018288209.1
73
Bade OZJ06330.1
54
Ltra XP 021875269.1
85
Rirr XP 025173879.1
0.05

### RPL24

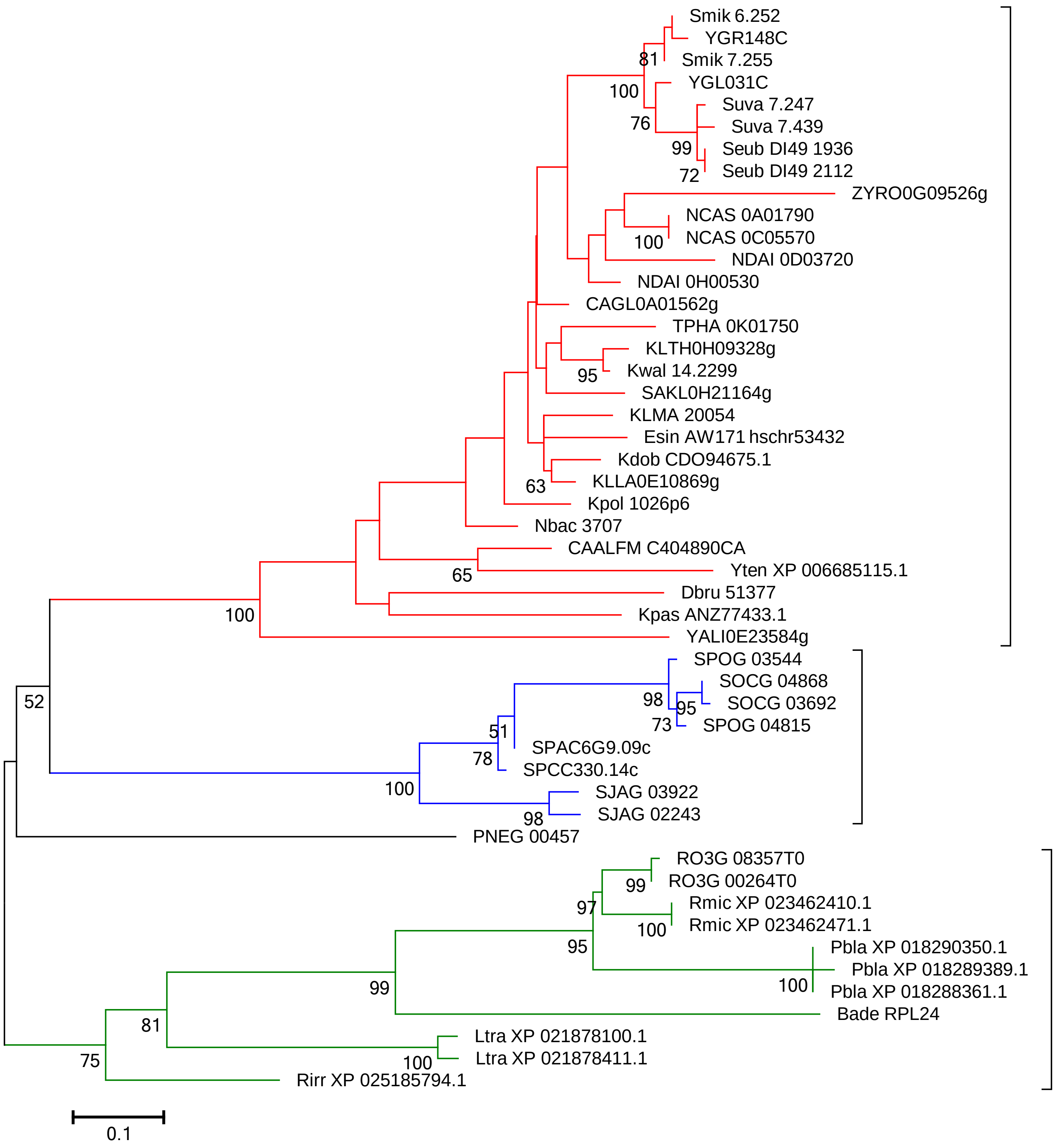

Smik 6.252
YGR148C
Smik 7.255
YGL031C
Suva 7.247
Suva 7.439
Seub DI49 1936
Seub DI49 2112
ZYRO0G09526g
NCAS 0A01790
NCAS 0C05570
NDAI 0D03720
NDAI 0H00530
CAGL0A01562g
TPHA 0K01750
KLTH0H09328g
Kwal 14.2299
SAKL0H21164g
KLMA 20054
Esin AW171 hschr53432
Kdob CDO94675.1
KLLA0E10869g
Kpol 1026p6
Nbac 3707
CAALFM C404890CA
Yten XP 006685115.1
Dbru 51377
Kpas ANZ77433.1
YALI0E23584g
SPOG 03544
SOCG 04868
SOCG 03692
SPOG 04815
SPAC6G9.09c
SPCC330.14c
SJAG 03922
SJAG 02243
PNEG 00457
RO3G 08357T0
RO3G 00264T0
Rmic XP 023462410.1
Rmic XP 023462471.1
Pbla XP 018290350.1
Pbla XP 018289389.1
Pbla XP 018288361.1
Bade RPL24
Ltra XP 021878100.1
Ltra XP 021878411.1
Rirr XP 025185794.1
81
100
76
99
72
100
95
63
65
100
98
95
73
51
78
100
98
52
99
97
100
95
100
99
81
100
75
0.1

### RPL25 (RPL23A)

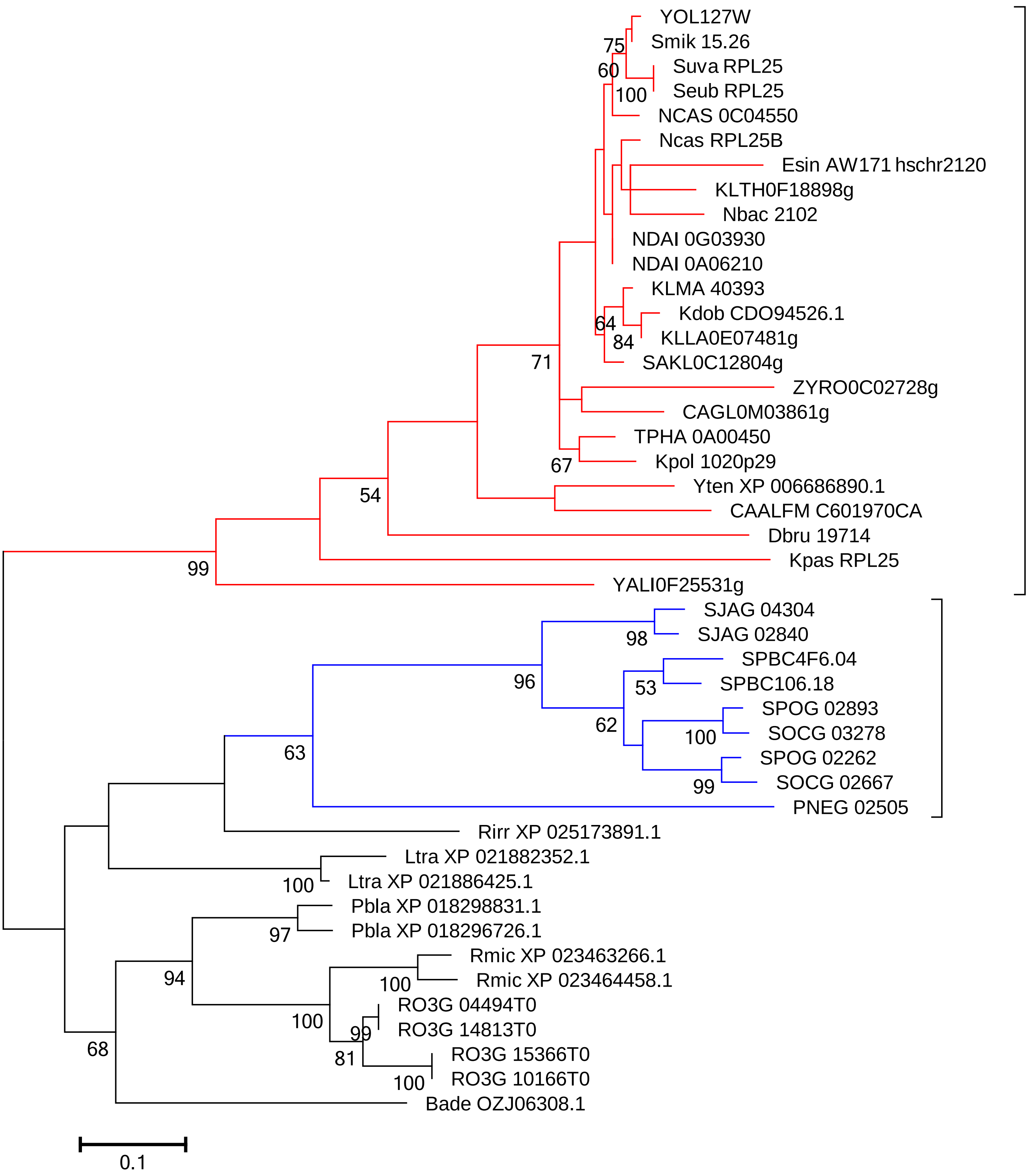

### RPL26

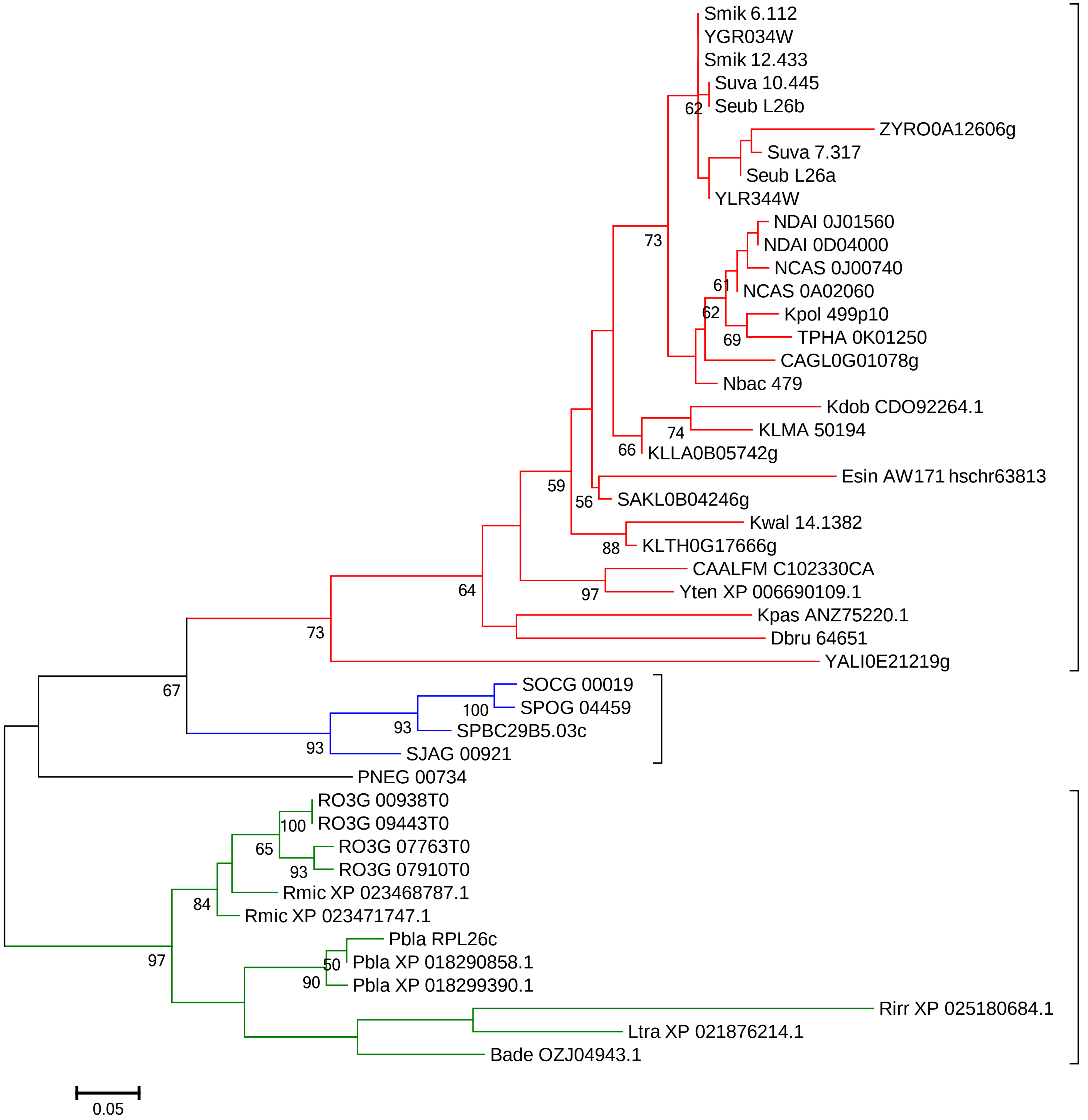

Smik 6.112
YGR034W
Smik 12.433
Suva 10.445
Seub L26b
62
ZYRO0A12606g
Suva 7.317
Seub L26a
YLR344W
NDAI 0J01560
NDAI 0D04000
NCAS 0J00740
61
NCAS 0A02060
62
Kpol 499p10
69
TPHA 0K01250
CAGL0G01078g
Nbac 479
73
Kdob CDO92264.1
74
KLMA 50194
66
KLLA0B05742g
Esin AW171 hschr63813
59
56
SAKL0B04246g
Kwal 14.1382
88
KLTH0G17666g
CAALFM C102330CA
64
97
Yten XP 006690109.1
Kpas ANZ75220.1
Dbru 64651
73
YALI0E21219g
SOCG 00019
100
SPOG 04459
67
93
SPBC29B5.03c
93
SJAG 00921
PNEG 00734
RO3G 00938T0
100
RO3G 09443T0
65
RO3G 07763T0
93
RO3G 07910T0
Rmic XP 023468787.1
84
Rmic XP 023471747.1
Pbla RPL26c
50
Pbla XP 018290858.1
97
90
Pbla XP 018299390.1
Rirr XP 025180684.1
Ltra XP 021876214.1
Bade OZJ04943.1
0.05

### RPL27

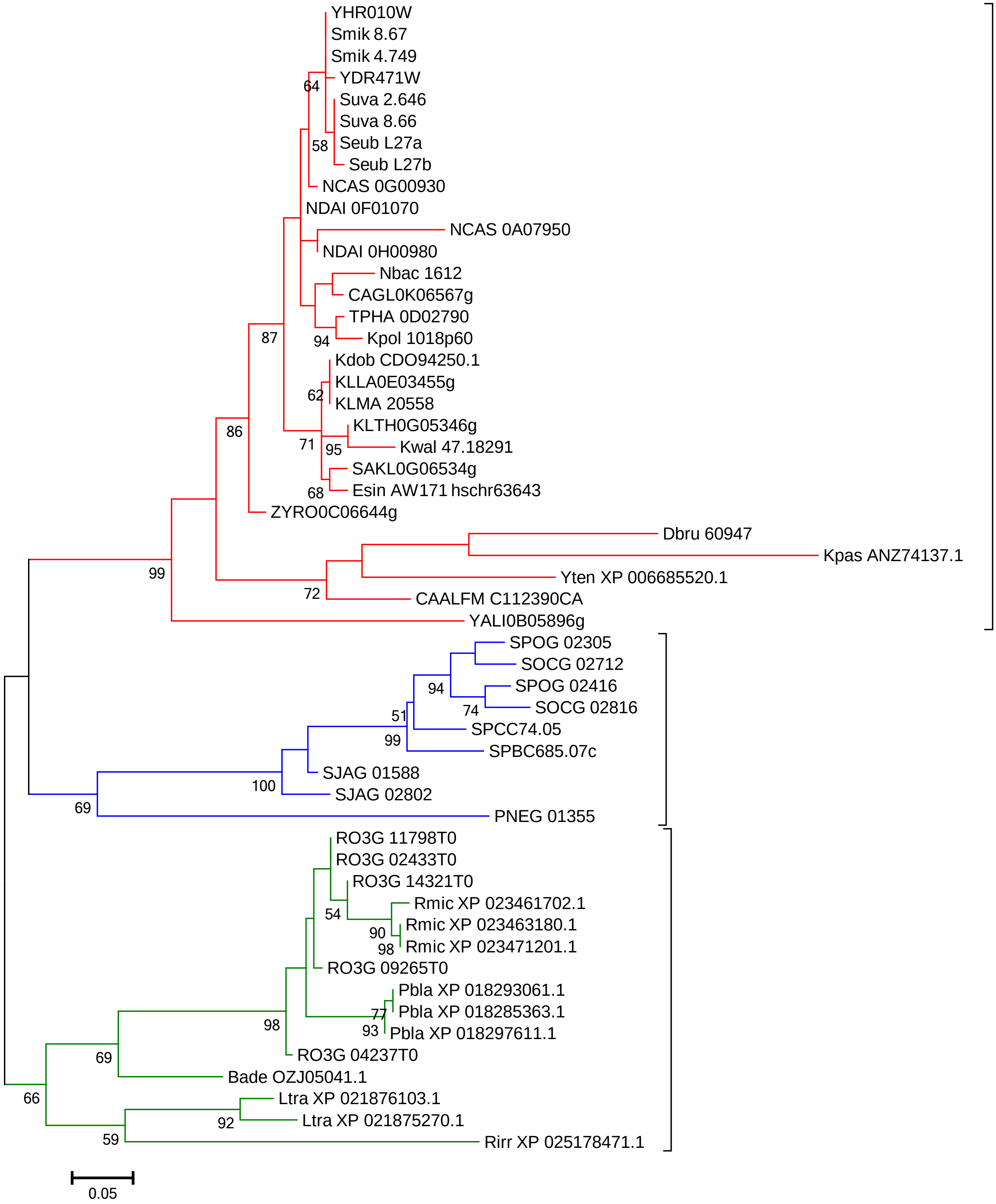

YHR010W
Smik 8.67
Smik 4.749
YDR471W
Suva 2.646
Suva 8.66
Seub L27a
Seub L27b
NCAS 0G00930
NDAI 0F01070
NCAS 0A07950
NDAI 0H00980
Nbac 1612
CAGL0K06567g
TPHA 0D02790
Kpol 1018p60
Kdob CDO94250.1
KLLA0E03455g
KLMA 20558
KLTH0G05346g
Kwal 47.18291
SAKL0G06534g
Esin AW171 hschr63643
ZYRO0C06644g
Dbru 60947
Kpas ANZ74137.1
Yten XP 006685520.1
CAALFM C112390CA
YALI0B05896g
SPOG 02305
SOCG 02712
SPOG 02416
SOCG 02816
SPCC74.05
SPBC685.07c
SJAG 01588
SJAG 02802
PNEG 01355
RO3G 11798T0
RO3G 02433T0
RO3G 14321T0
Rmic XP 023461702.1
Rmic XP 023463180.1
Rmic XP 023471201.1
RO3G 09265T0
Pbla XP 018293061.1
Pbla XP 018285363.1
Pbla XP 018297611.1
RO3G 04237T0
Bade OZJ05041.1
Ltra XP 021876103.1
Ltra XP 021875270.1
Rirr XP 025178471.1
64
58
87
94
62
86
71
95
68
99
72
94
74
51
99
100
69
54
90
98
77
93
98
69
66
92
59
0.05

### RPL28 (RPL27A)

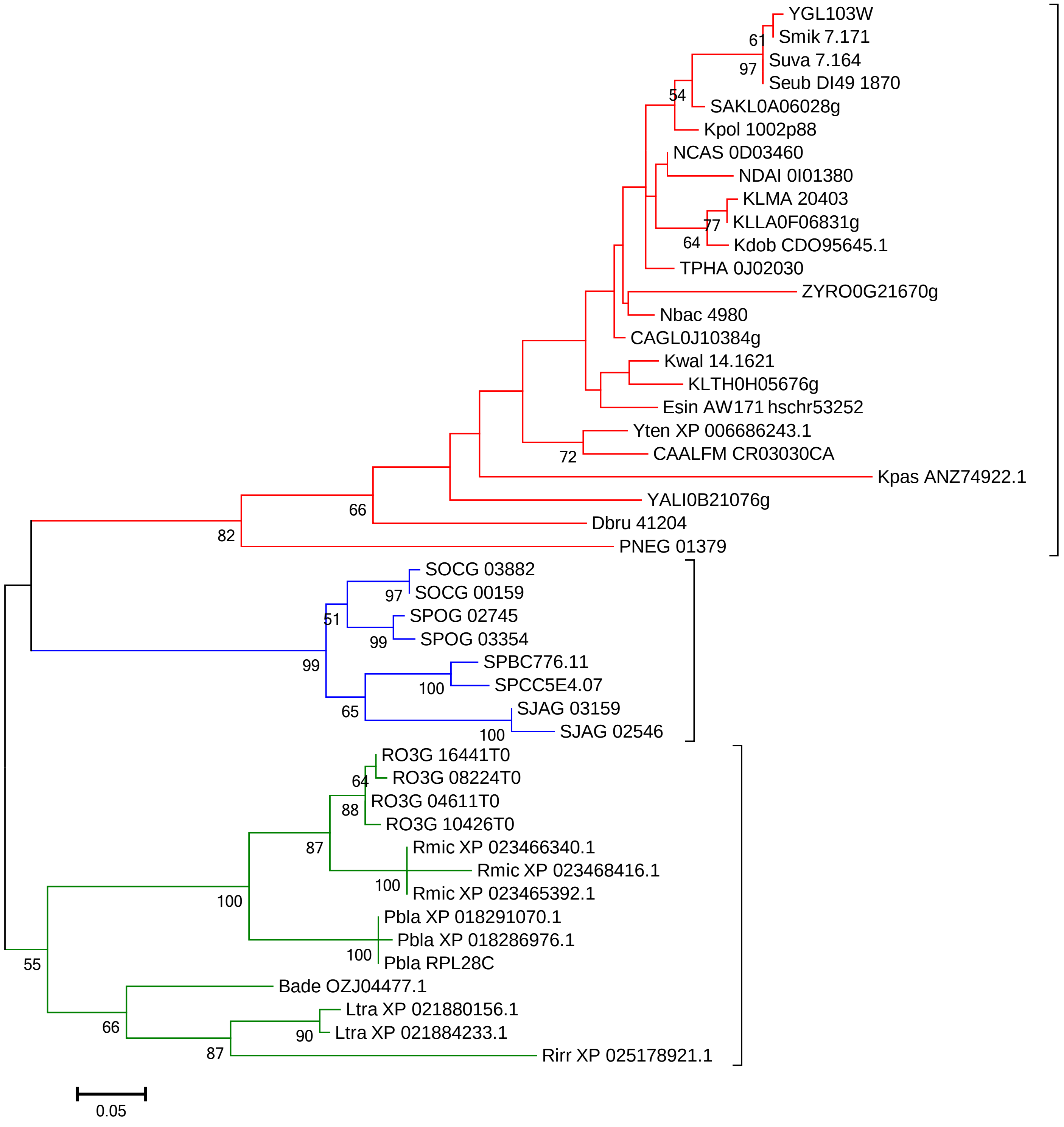

### RPL28e

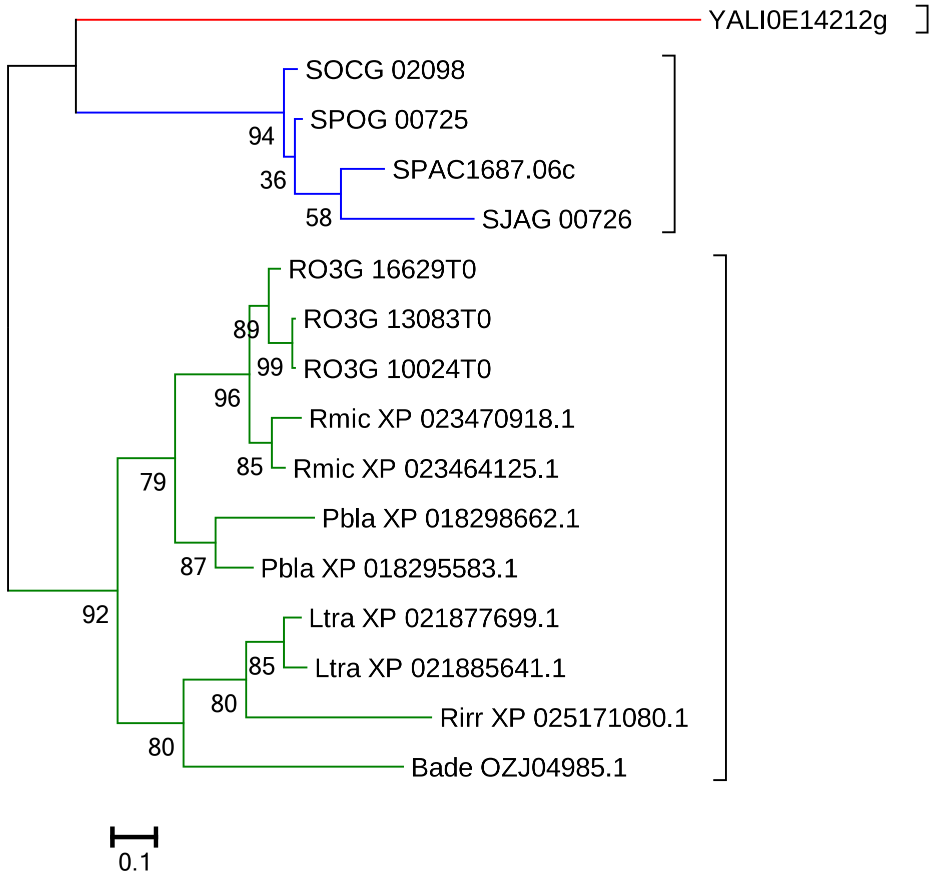
YALI0E14212g
SOCG 02098
SPOG 00725
SPAC1687.06c
SJAG 00726
RO3G 16629T0
RO3G 13083T0
RO3G 10024T0
Rmic XP 023470918.1
Rmic XP 023464125.1
Pbla XP 018298662.1
Pbla XP 018295583.1
Ltra XP 021877699.1
Ltra XP 021885641.1
Rirr XP 025171080.1
Bade OZJ04985.1
94
36
58
89
99
96
85
79
87
92
85
80
80
0.1

### RPL29

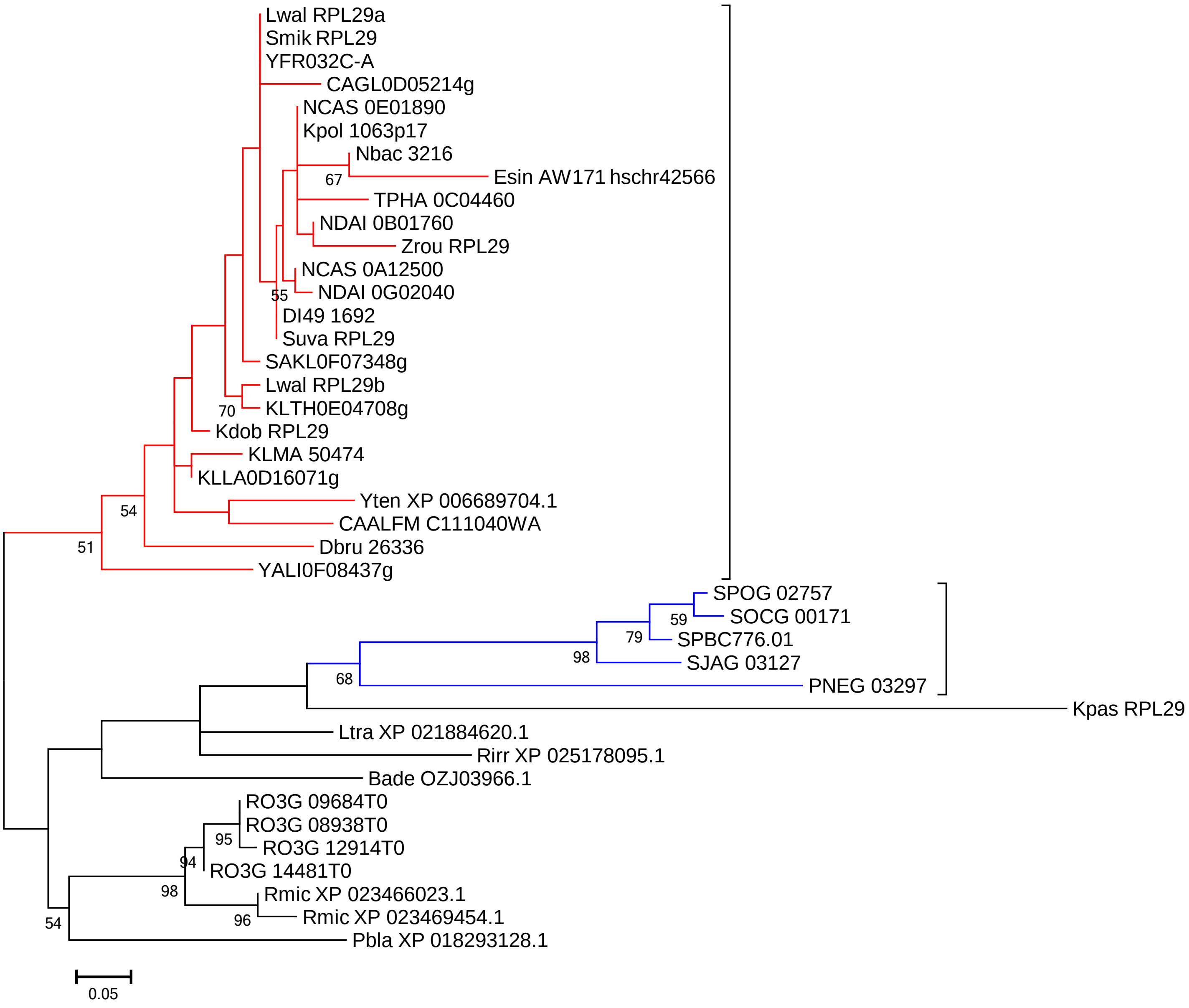

Lwal RPL29a
Smik RPL29
YFR032C-A
CAGL0D05214g
NCAS 0E01890
Kpol 1063p17
Nbac 3216
67
Esin AW171 hschr42566
TPHA 0C04460
NDAI 0B01760
Zrou RPL29
NCAS 0A12500
55
NDAI 0G02040
DI49 1692
Suva RPL29
SAKL0F07348g
Lwal RPL29b
70
KLTH0E04708g
Kdob RPL29
KLMA 50474
KLLA0D16071g
Yten XP 006689704.1
54
CAALFM C111040WA
Dbru 26336
51
YALI0F08437g
SPOG 02757
59
SOCG 00171
79
SPBC776.01
98
SJAG 03127
68
PNEG 03297
Kpas RPL29
Ltra XP 021884620.1
Rirr XP 025178095.1
Bade OZJ03966.1
RO3G 09684T0
RO3G 08938T0
95
RO3G 12914T0
94
RO3G 14481T0
98
Rmic XP 023466023.1
96
Rmic XP 023469454.1
54
Pbla XP 018293128.1
0.05

### RPL30

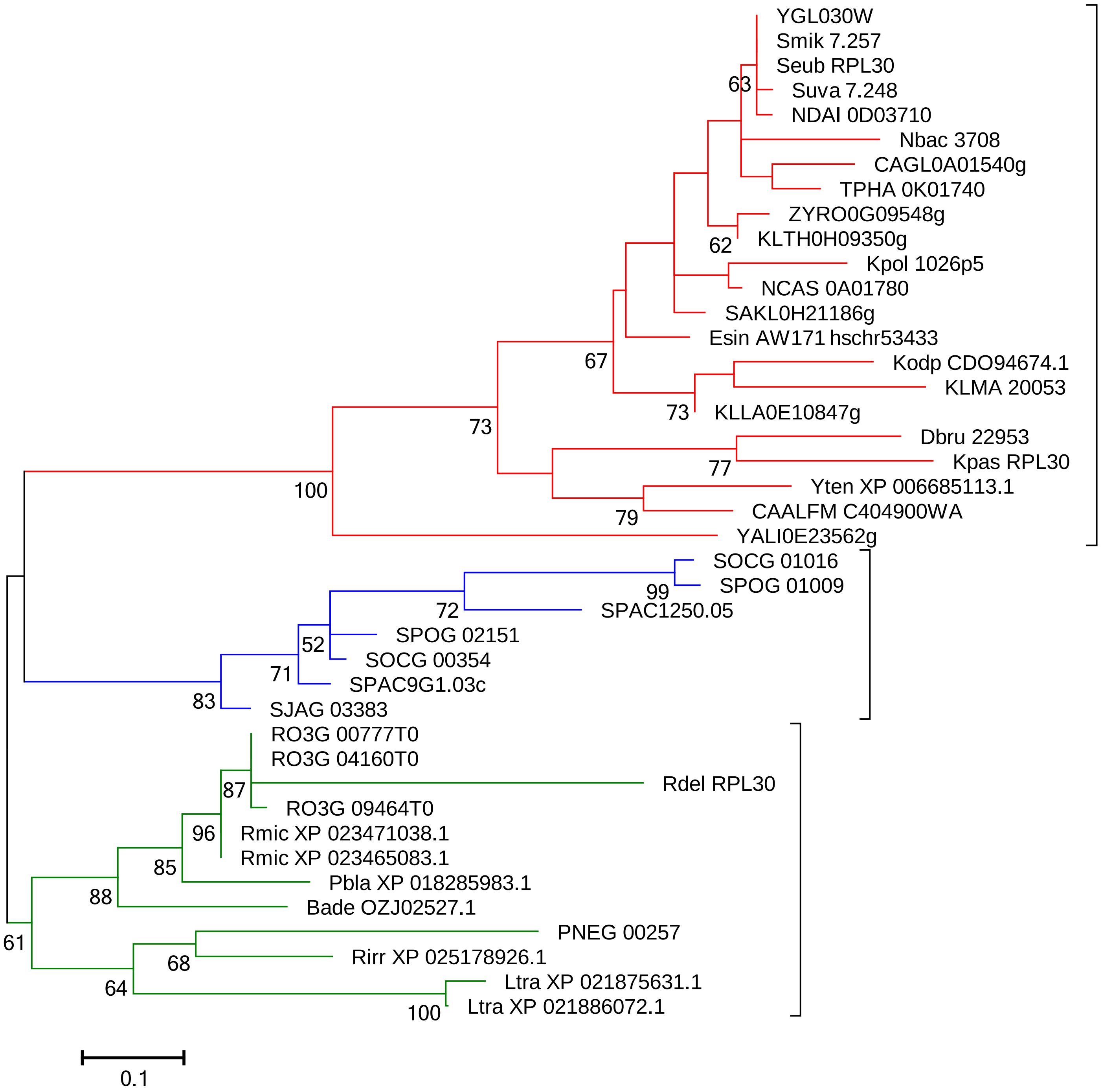

YGL030W
Smik 7.257
Seub RPL30
63
Suva 7.248
NDAI 0D03710
Nbac 3708
CAGL0A01540g
TPHA 0K01740
ZYRO0G09548g
62
KLTH0H09350g
Kpol 1026p5
NCAS 0A01780
SAKL0H21186g
Esin AW171 hschr53433
67
Kodp CDO94674.1
KLMA 20053
73
KLLA0E10847g
73
Dbru 22953
77
Kpas RPL30
Yten XP 006685113.1
79
CAALFM C404900WA
100
YALI0E23562g
SOCG 01016
99
SPOG 01009
72
SPAC1250.05
SPOG 02151
52
SOCG 00354
71
SPAC9G1.03c
83
SJAG 03383
RO3G 00777T0
RO3G 04160T0
Rdel RPL30
87
RO3G 09464T0
96
Rmic XP 023471038.1
Rmic XP 023465083.1
85
Pbla XP 018285983.1
88
Bade OZJ02527.1
61
PNEG 00257
68
Rirr XP 025178926.1
64
Ltra XP 021875631.1
100
Ltra XP 021886072.1
0.1

### RPL31

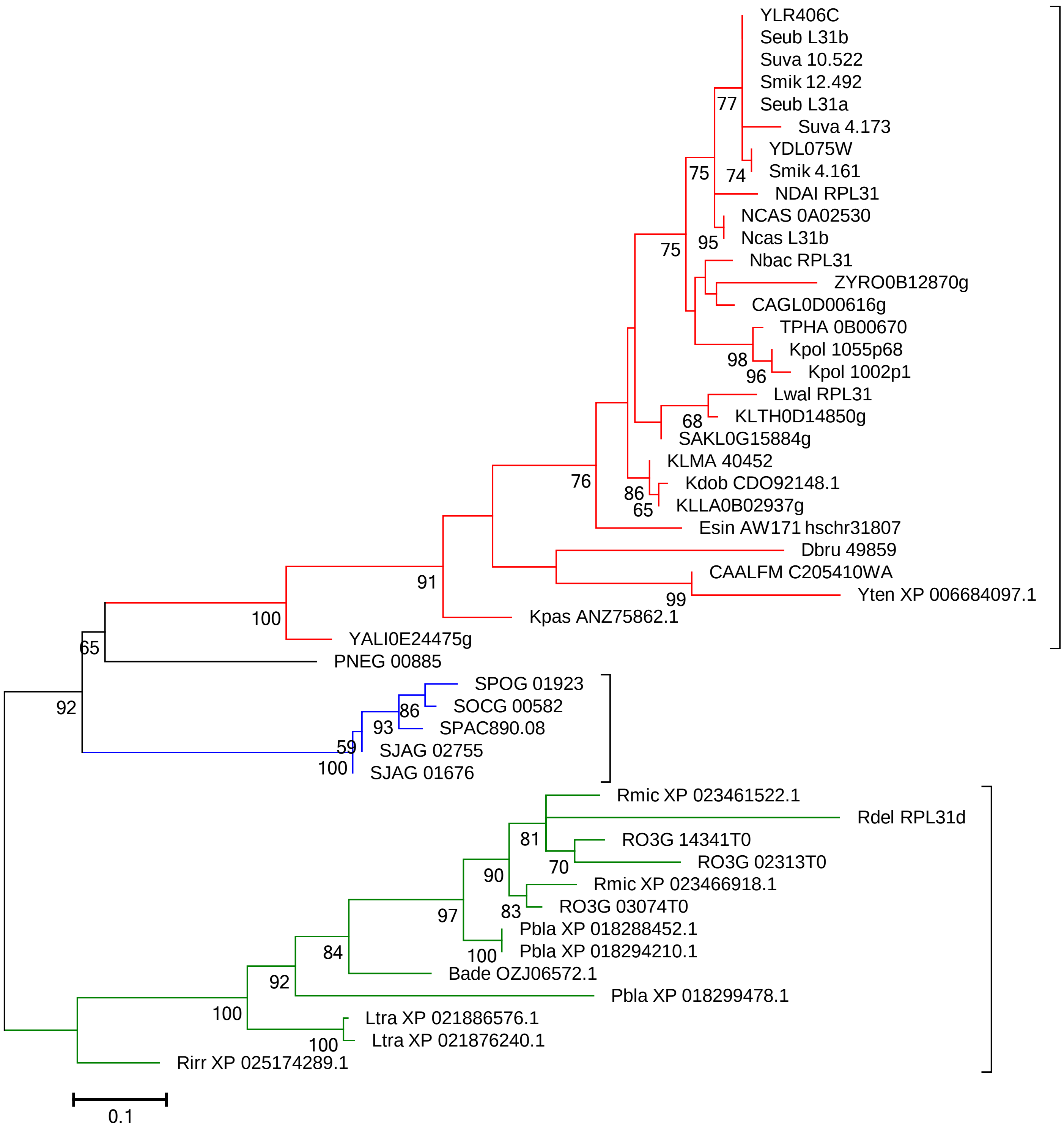
YLR406C
Seub L31b
Suva 10.522
Smik 12.492
Seub L31a
Suva 4.173
YDL075W
Smik 4.161
NDAI RPL31
NCAS 0A02530
Ncas L31b
Nbac RPL31
ZYRO0B12870g
CAGL0D00616g
TPHA 0B00670
Kpol 1055p68
Kpol 1002p1
Lwal RPL31
KLTH0D14850g
SAKL0G15884g
KLMA 40452
Kdob CDO92148.1
KLLA0B02937g
Esin AW171 hschr31807
Dbru 49859
CAALFM C205410WA
Yten XP 006684097.1
Kpas ANZ75862.1
YALI0E24475g
PNEG 00885
SPOG 01923
SOCG 00582
SPAC890.08
SJAG 02755
SJAG 01676
Rmic XP 023461522.1
Rdel RPL31d
RO3G 14341T0
RO3G 02313T0
Rmic XP 023466918.1
RO3G 03074T0
Pbla XP 018288452.1
Pbla XP 018294210.1
Bade OZJ06572.1
Pbla XP 018299478.1
Ltra XP 021886576.1
Ltra XP 021876240.1
Rirr XP 025174289.1
77
75
74
75
95
98
96
68
76
86
65
91
99
100
65
92
86
93
59
100
81
70
90
83
97
100
84
92
100
100
0.1

### RPL32

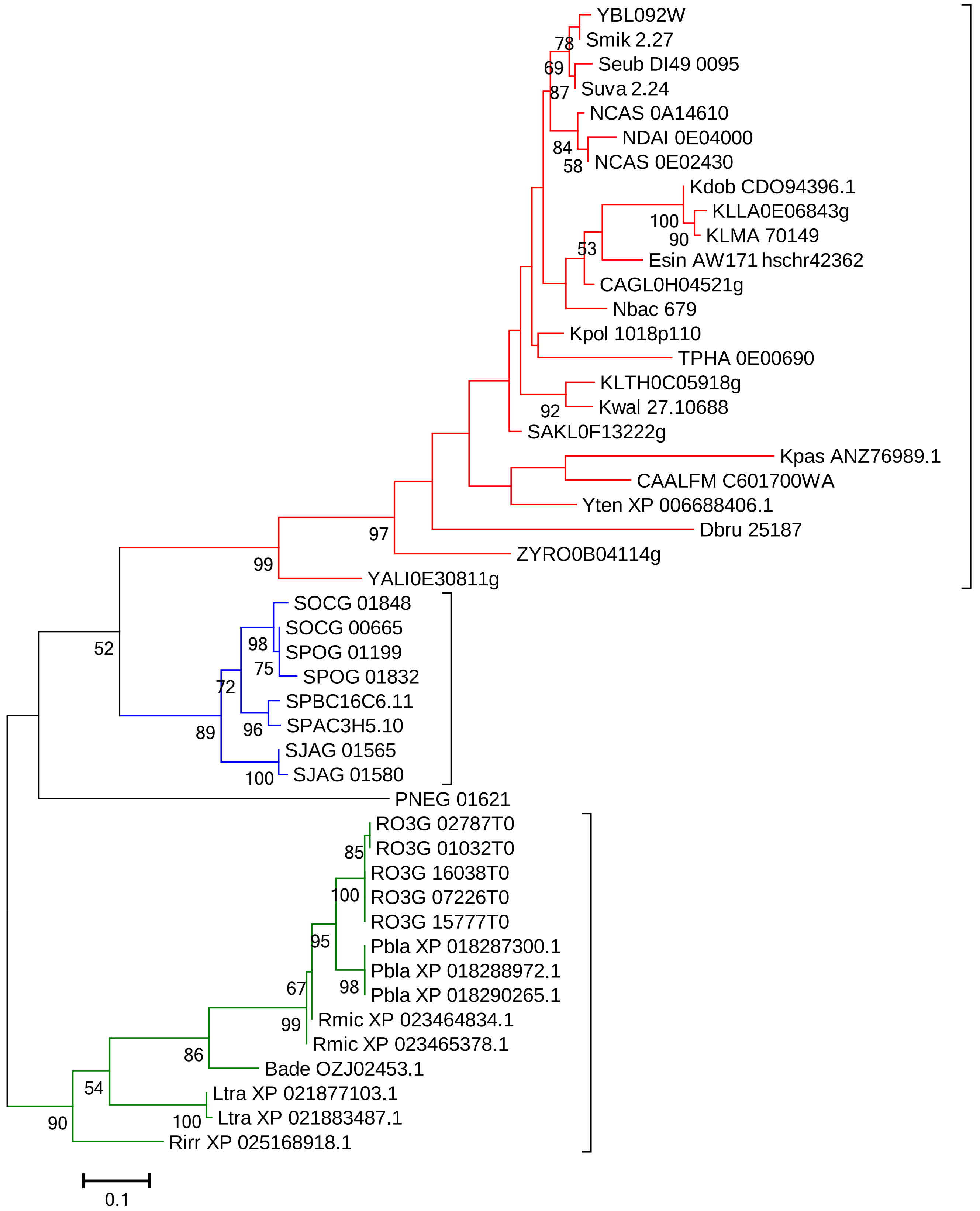

YBL092W
Smik 2.27
Seub DI49 0095
Suva 2.24
NCAS 0A14610
NDAI 0E04000
NCAS 0E02430
Kdob CDO94396.1
KLLA0E06843g
KLMA 70149
Esin AW171 hschr42362
CAGL0H04521g
Nbac 679
Kpol 1018p110
TPHA 0E00690
KLTH0C05918g
Kwal 27.10688
SAKL0F13222g
Kpas ANZ76989.1
CAALFM C601700WA
Yten XP 006688406.1
Dbru 25187
ZYRO0B04114g
YALI0E30811g
SOCG 01848
SOCG 00665
SPOG 01199
SPOG 01832
SPBC16C6.11
SPAC3H5.10
SJAG 01565
SJAG 01580
PNEG 01621
RO3G 02787T0
RO3G 01032T0
RO3G 16038T0
RO3G 07226T0
RO3G 15777T0
Pbla XP 018287300.1
Pbla XP 018288972.1
Pbla XP 018290265.1
Rmic XP 023464834.1
Rmic XP 023465378.1
Bade OZJ02453.1
Ltra XP 021877103.1
Ltra XP 021883487.1
Rirr XP 025168918.1
78
69
87
84
58
100
90
53
92
97
99
52
98
75
72
89
96
100
85
100
95
67
98
99
86
54
100
90
0.1

### RPL33 (RPL35A)

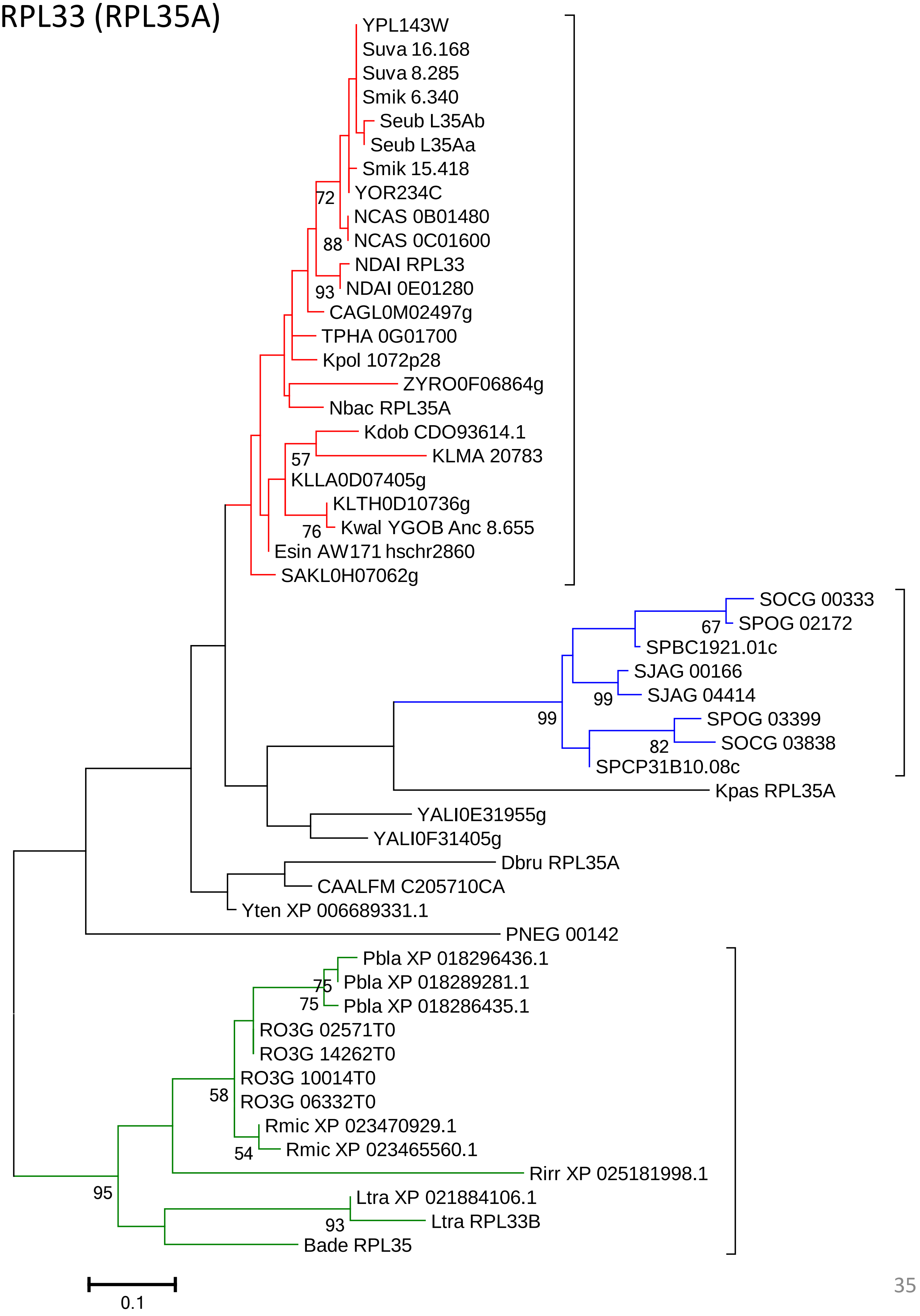

### RPL34

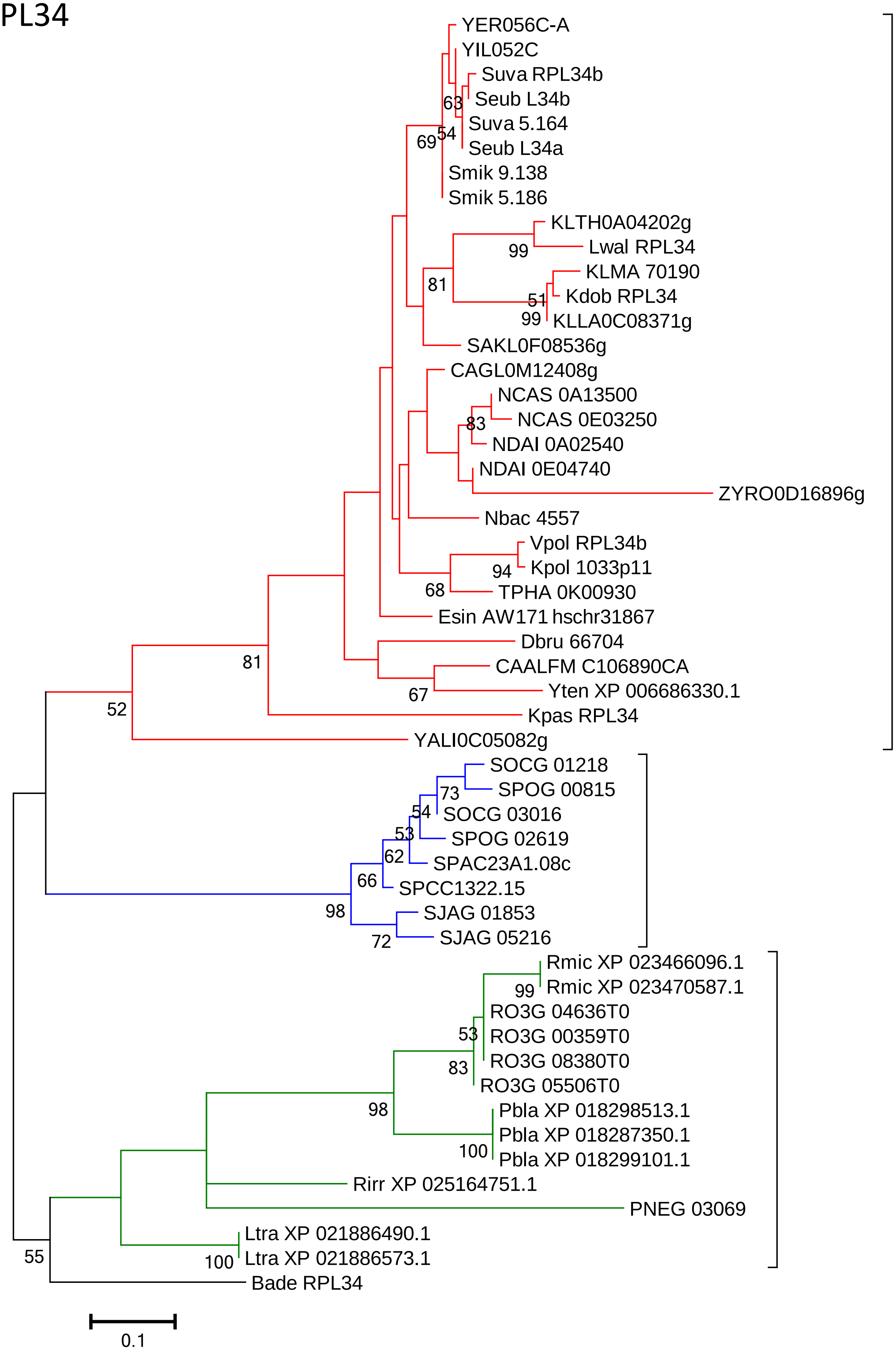
YER056C-A
YIL052C
Suva RPL34b
Seub L34b
Suva 5.164
Seub L34a
Smik 9.138
Smik 5.186
KLTH0A04202g
Lwal RPL34
KLMA 70190
Kdob RPL34
KLLA0C08371g
SAKL0F08536g
CAGL0M12408g
NCAS 0A13500
NCAS 0E03250
NDAI 0A02540
NDAI 0E04740
ZYRO0D16896g
Nbac 4557
Vpol RPL34b
Kpol 1033p11
TPHA 0K00930
Esin AW171 hschr31867
Dbru 66704
CAALFM C106890CA
Yten XP 006686330.1
Kpas RPL34
YALI0C05082g
SOCG 01218
SPOG 00815
SOCG 03016
SPOG 02619
SPAC23A1.08c
SPCC1322.15
SJAG 01853
SJAG 05216
Rmic XP 023466096.1
Rmic XP 023470587.1
RO3G 04636T0
RO3G 00359T0
RO3G 08380T0
RO3G 05506T0
Pbla XP 018298513.1
Pbla XP 018287350.1
Pbla XP 018299101.1
Rirr XP 025164751.1
PNEG 03069
Ltra XP 021886490.1
Ltra XP 021886573.1
Bade RPL34
63
54
69
99
81
51
99
83
94
68
81
67
52
73
54
53
62
66
98
72
99
53
83
98
100
55
100
0.1

### RPL35

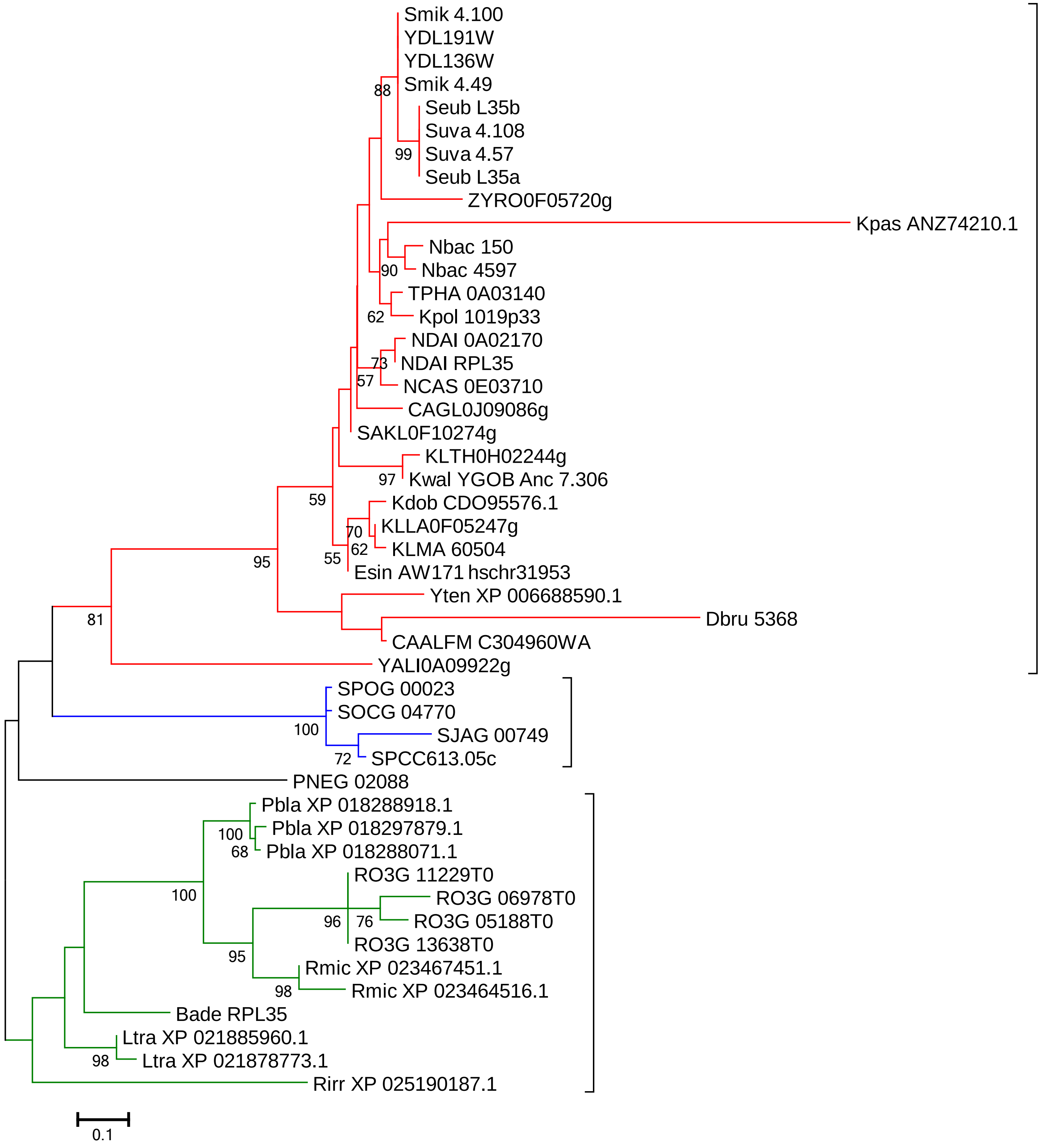

Smik 4.100
YDL191W
YDL136W
Smik 4.49
Seub L35b
Suva 4.108
Suva 4.57
Seub L35a
ZYRO0F05720g
Kpas ANZ74210.1
Nbac 150
Nbac 4597
TPHA 0A03140
Kpol 1019p33
NDAI 0A02170
NDAI RPL35
NCAS 0E03710
CAGL0J09086g
SAKL0F10274g
KLTH0H02244g
Kwal YGOB Anc 7.306
Kdob CDO95576.1
KLLA0F05247g
KLMA 60504
Esin AW171 hschr31953
Yten XP 006688590.1
Dbru 5368
CAALFM C304960WA
YALI0A09922g
SPOG 00023
SOCG 04770
SJAG 00749
SPCC613.05c
PNEG 02088
Pbla XP 018288918.1
Pbla XP 018297879.1
Pbla XP 018288071.1
RO3G 11229T0
RO3G 06978T0
RO3G 05188T0
RO3G 13638T0
Rmic XP 023467451.1
Rmic XP 023464516.1
Bade RPL35
Ltra XP 021885960.1
Ltra XP 021878773.1
Rirr XP 025190187.1
88
99
90
62
73
57
97
59
70
62
55
95
81
100
72
100
68
100
96
76
95
98
98
0.1

### RPL36

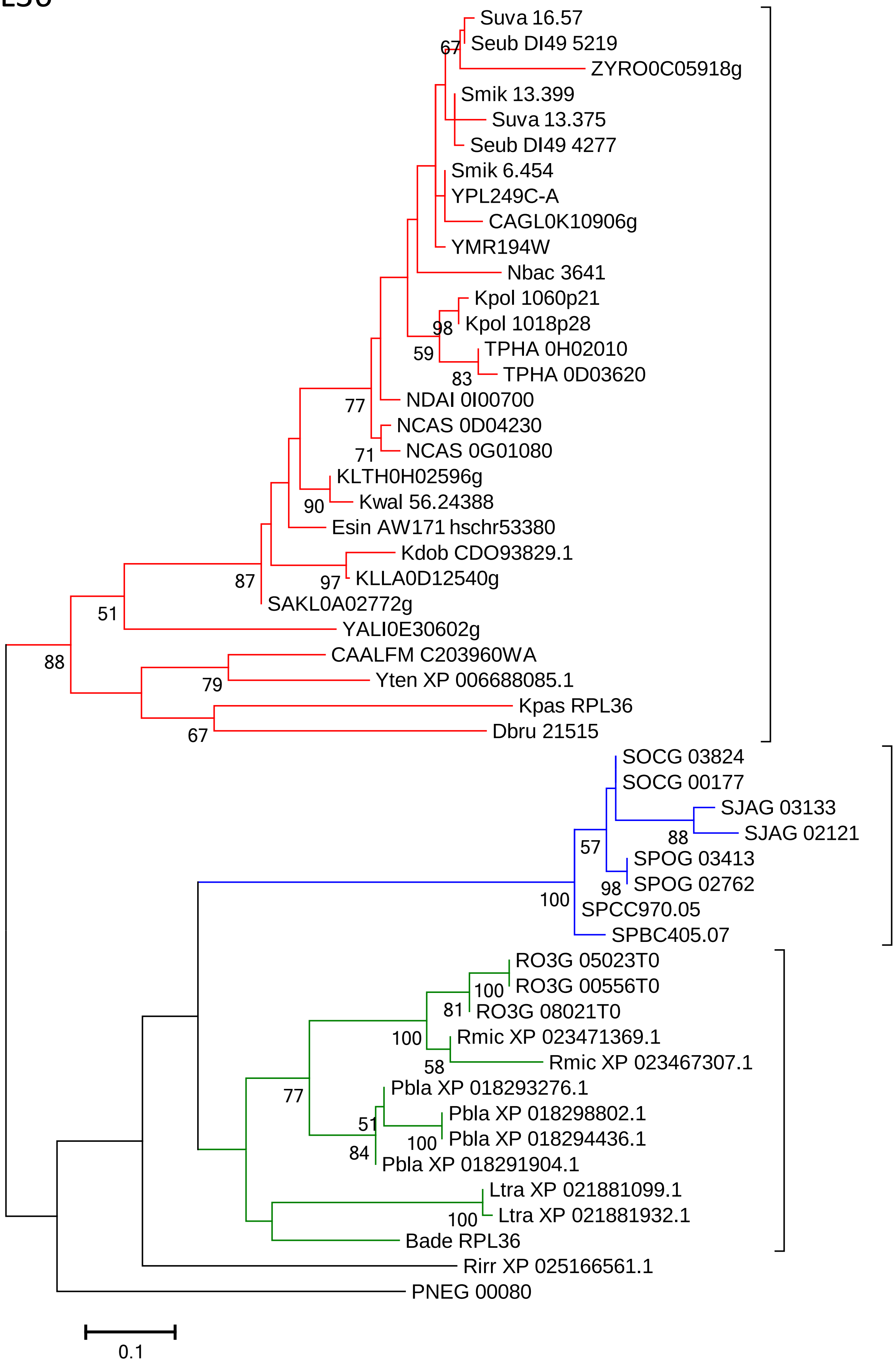

Suva 16.57
Seub DI49 5219
67
ZYRO0C05918g
Smik 13.399
Suva 13.375
Seub DI49 4277
Smik 6.454
YPL249C-A
CAGL0K10906g
YMR194W
Nbac 3641
Kpol 1060p21
98
Kpol 1018p28
59
TPHA 0H02010
83
TPHA 0D03620
77
NDAI 0I00700
NCAS 0D04230
71
NCAS 0G01080
KLTH0H02596g
90
Kwal 56.24388
Esin AW171 hschr53380
Kdob CDO93829.1
87
97
KLLA0D12540g
SAKL0A02772g
51
YALI0E30602g
CAALFM C203960WA
88
79
Yten XP 006688085.1
Kpas RPL36
67
Dbru 21515
SOCG 03824
SOCG 00177
SJAG 03133
88
SJAG 02121
57
SPOG 03413
98
SPOG 02762
100
SPCC970.05
SPBC405.07
RO3G 05023T0
100
RO3G 00556T0
81
RO3G 08021T0
100
Rmic XP 023471369.1
58
Rmic XP 023467307.1
Pbla XP 018293276.1
77
51
Pbla XP 018298802.1
100
Pbla XP 018294436.1
84
Pbla XP 018291904.1
Ltra XP 021881099.1
100
Ltra XP 021881932.1
Bade RPL36
Rirr XP 025166561.1
PNEG 00080
0.1

### RPL37

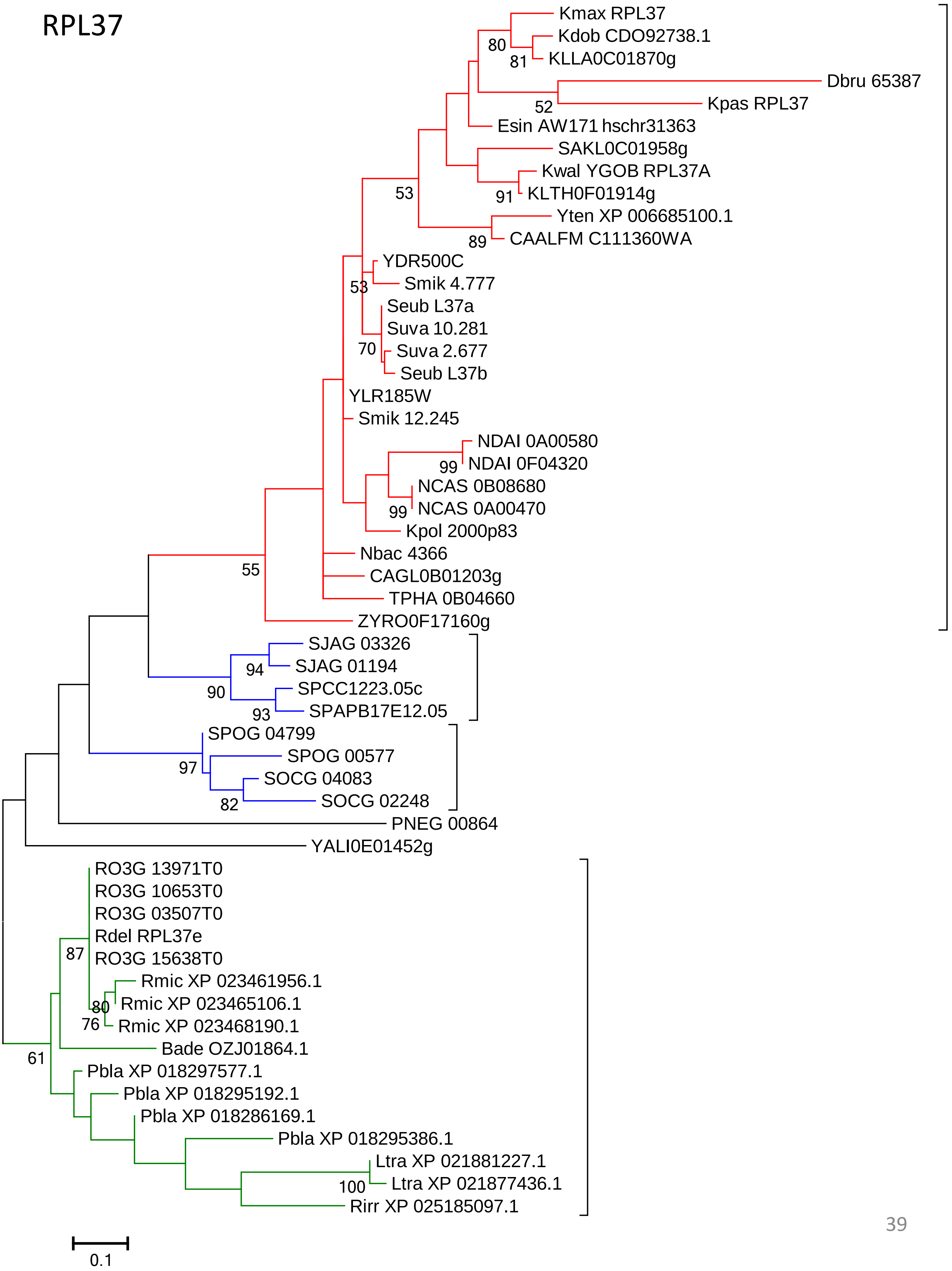

Kmax RPL37
Kdob CDO92738.1
KLLA0C01870g
Dbru 65387
Kpas RPL37
Esin AW171 hschr31363
SAKL0C01958g
Kwal YGOB RPL37A
KLTH0F01914g
Yten XP 006685100.1
CAALFM C111360WA
YDR500C
Smik 4.777
Seub L37a
Suva 10.281
Suva 2.677
Seub L37b
YLR185W
Smik 12.245
NDAI 0A00580
NDAI 0F04320
NCAS 0B08680
NCAS 0A00470
Kpol 2000p83
Nbac 4366
CAGL0B01203g
TPHA 0B04660
ZYRO0F17160g
SJAG 03326
SJAG 01194
SPCC1223.05c
SPAPB17E12.05
SPOG 04799
SPOG 00577
SOCG 04083
SOCG 02248
PNEG 00864
YALI0E01452g
RO3G 13971T0
RO3G 10653T0
RO3G 03507T0
Rdel RPL37e
RO3G 15638T0
Rmic XP 023461956.1
Rmic XP 023465106.1
Rmic XP 023468190.1
Bade OZJ01864.1
Pbla XP 018297577.1
Pbla XP 018295192.1
Pbla XP 018286169.1
Pbla XP 018295386.1
Ltra XP 021881227.1
Ltra XP 021877436.1
Rirr XP 025185097.1
80
81
52
53
91
89
53
70
99
99
55
94
90
93
97
82
87
80
76
61
100
0.1

### RPL38

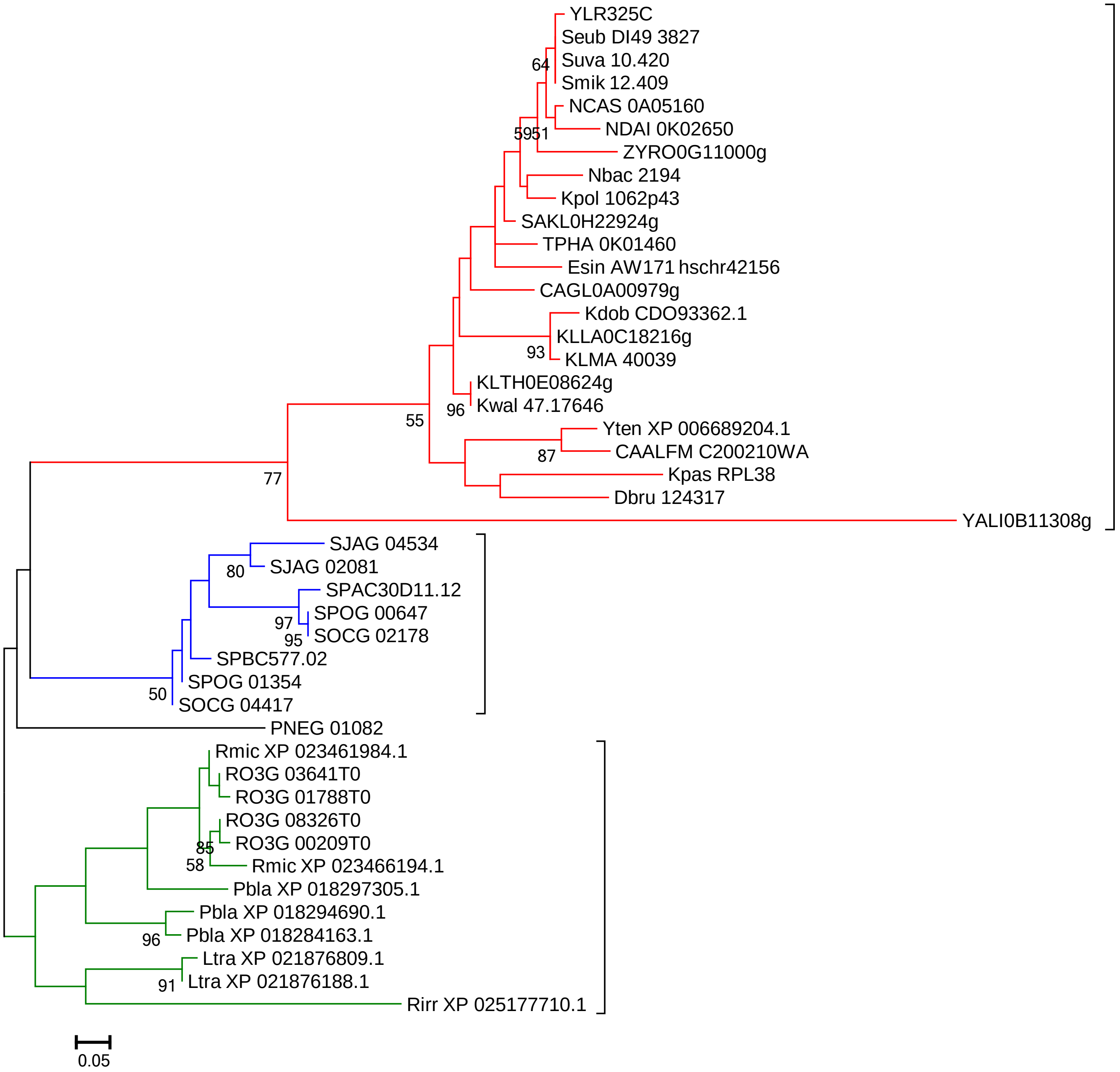
YLR325C
Seub DI49 3827
Suva 10.420
Smik 12.409
NCAS 0A05160
NDAI 0K02650
ZYRO0G11000g
Nbac 2194
Kpol 1062p43
SAKL0H22924g
TPHA 0K01460
Esin AW171 hschr42156
CAGL0A00979g
Kdob CDO93362.1
KLLA0C18216g
KLMA 40039
KLTH0E08624g
Kwal 47.17646
Yten XP 006689204.1
CAALFM C200210WA
Kpas RPL38
Dbru 124317
YALI0B11308g
SJAG 04534
SJAG 02081
SPAC30D11.12
SPOG 00647
SOCG 02178
SPBC577.02
SPOG 01354
SOCG 04417
PNEG 01082
Rmic XP 023461984.1
RO3G 03641T0
RO3G 01788T0
RO3G 08326T0
RO3G 00209T0
Rmic XP 023466194.1
Pbla XP 018297305.1
Pbla XP 018294690.1
Pbla XP 018284163.1
Ltra XP 021876809.1
Ltra XP 021876188.1
Rirr XP 025177710.1
64
5951
93
96
55
87
77
80
97
95
50
85
58
96
91
0.05

RPL39

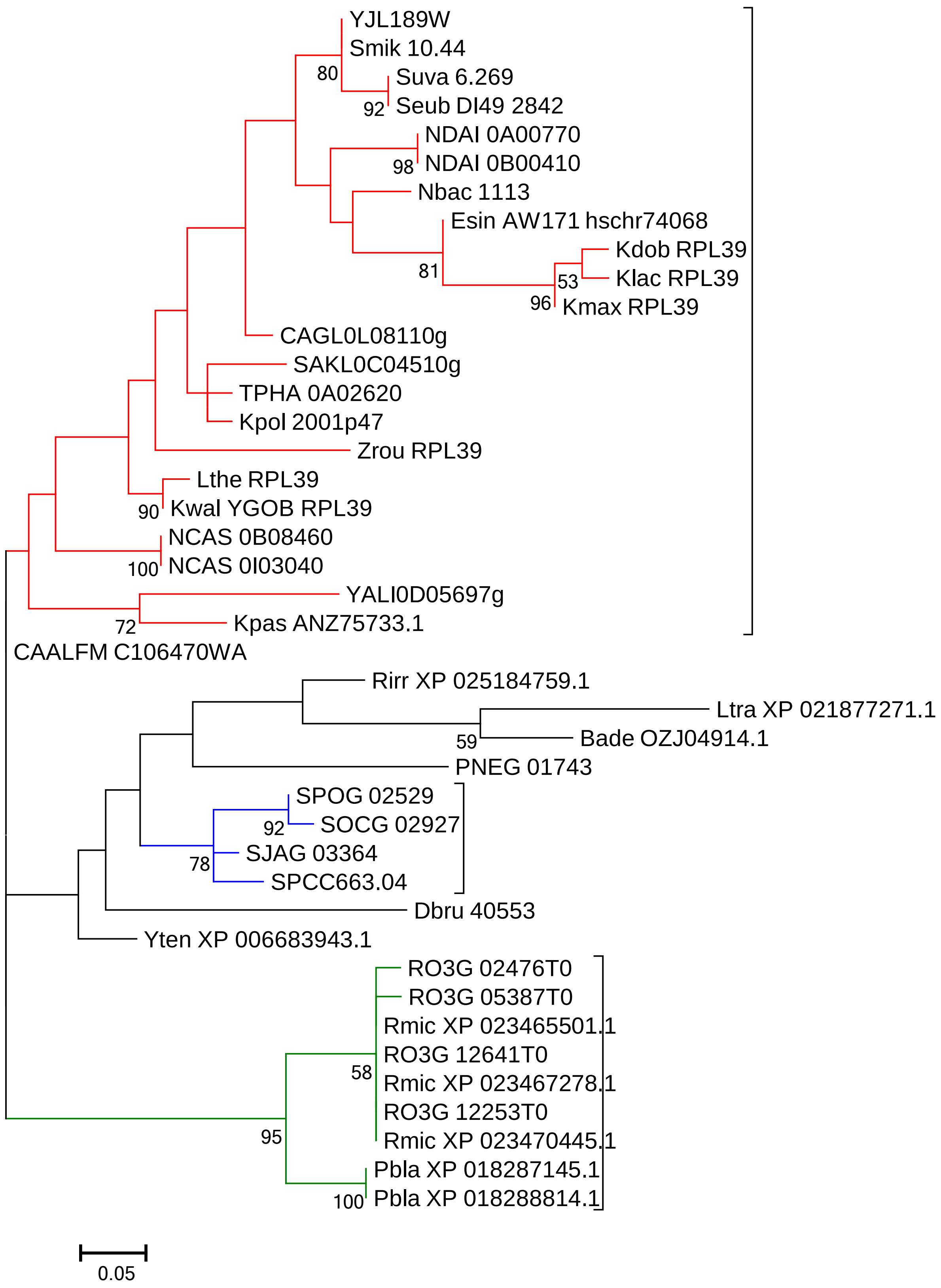
YJL189W
Smik 10.44
80
Suva 6.269
92
Seub DI49 2842
NDAI 0A00770
98
NDAI 0B00410
Nbac 1113
Esin AW171 hschr74068
Kdob RPL39
53
Klac RPL39
81
96
Kmax RPL39
CAGL0L08110g
SAKL0C04510g
TPHA 0A02620
Kpol 2001p47
Zrou RPL39
Lthe RPL39
90
Kwal YGOB RPL39
NCAS 0B08460
100
NCAS 0I03040
YALI0D05697g
72
Kpas ANZ75733.1
CAALFM C106470WA
Rirr XP 025184759.1
Ltra XP 021877271.1
59
Bade OZJ04914.1
PNEG 01743
SPOG 02529
92
SOCG 02927
SJAG 03364
78
SPCC663.04
Dbru 40553
Yten XP 006683943.1
RO3G 02476T0
RO3G 05387T0
Rmic XP 023465501.1
RO3G 12641T0
58
Rmic XP 023467278.1
RO3G 12253T0
95
Rmic XP 023470445.1
Pbla XP 018287145.1
100
Pbla XP 018288814.1
0.05

### RPL40

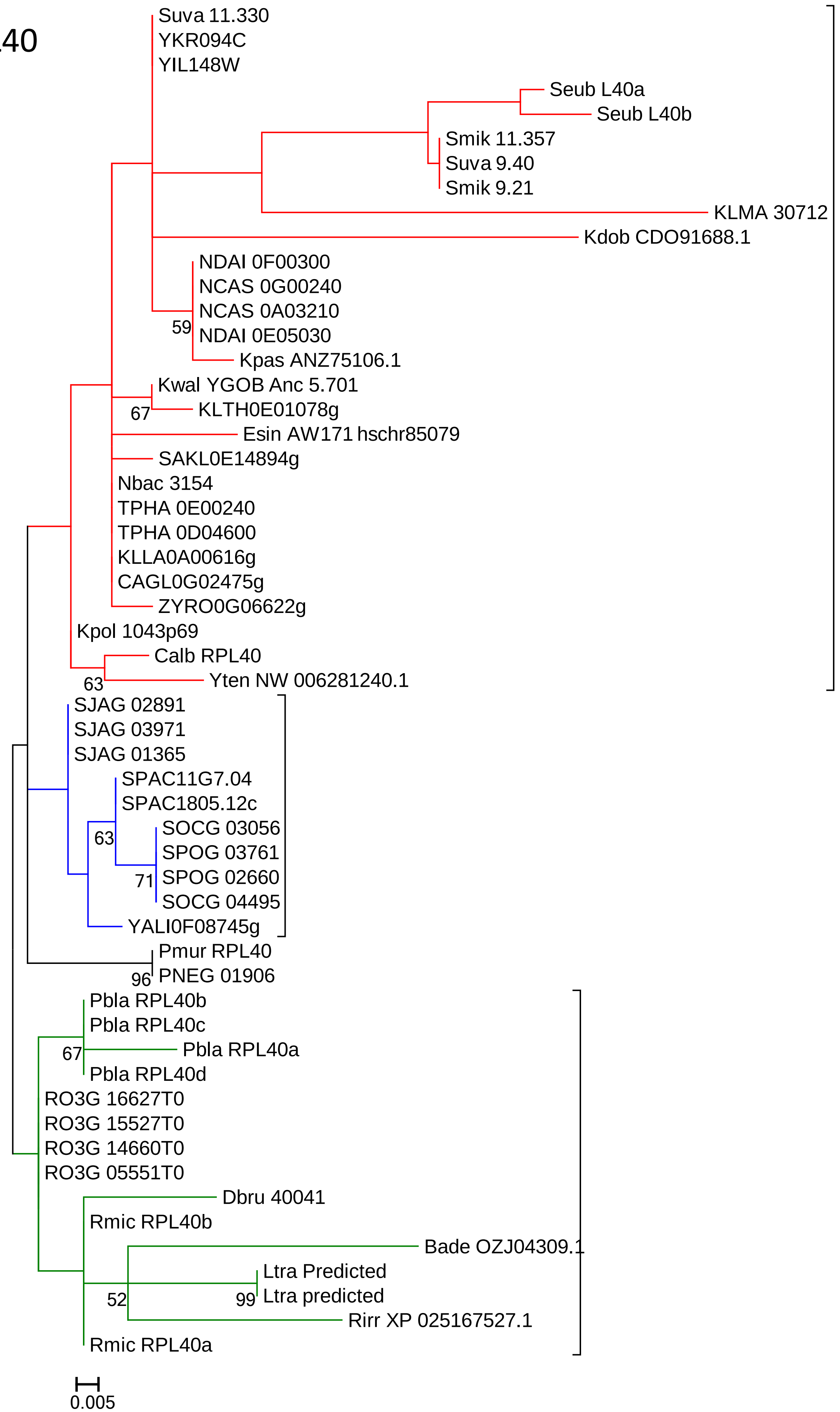
Suva 11.330
YKR094C
YIL148W
Seub L40a
Seub L40b
Smik 11.357
Suva 9.40
Smik 9.21
KLMA 30712
Kdob CDO91688.1
NDAI 0F00300
NCAS 0G00240
NCAS 0A03210
59
NDAI 0E05030
Kpas ANZ75106.1
Kwal YGOB Anc 5.701
67
KLTH0E01078g
Esin AW171 hschr85079
SAKL0E14894g
Nbac 3154
TPHA 0E00240
TPHA 0D04600
KLLA0A00616g
CAGL0G02475g
ZYRO0G06622g
Kpol 1043p69
Calb RPL40
63
Yten NW 006281240.1
SJAG 02891
SJAG 03971
SJAG 01365
SPAC11G7.04
SPAC1805.12c
63
SOCG 03056
SPOG 03761
71
SPOG 02660
SOCG 04495
YALI0F08745g
Pmur RPL40
96
PNEG 01906
Pbla RPL40b
Pbla RPL40c
67
Pbla RPL40a
Pbla RPL40d
RO3G 16627T0
RO3G 15527T0
RO3G 14660T0
RO3G 05551T0
Dbru 40041
Rmic RPL40b
Bade OZJ04309.1
Ltra Predicted
52
99
Ltra predicted
Rirr XP 025167527.1
Rmic RPL40a
0.005

### RPL41

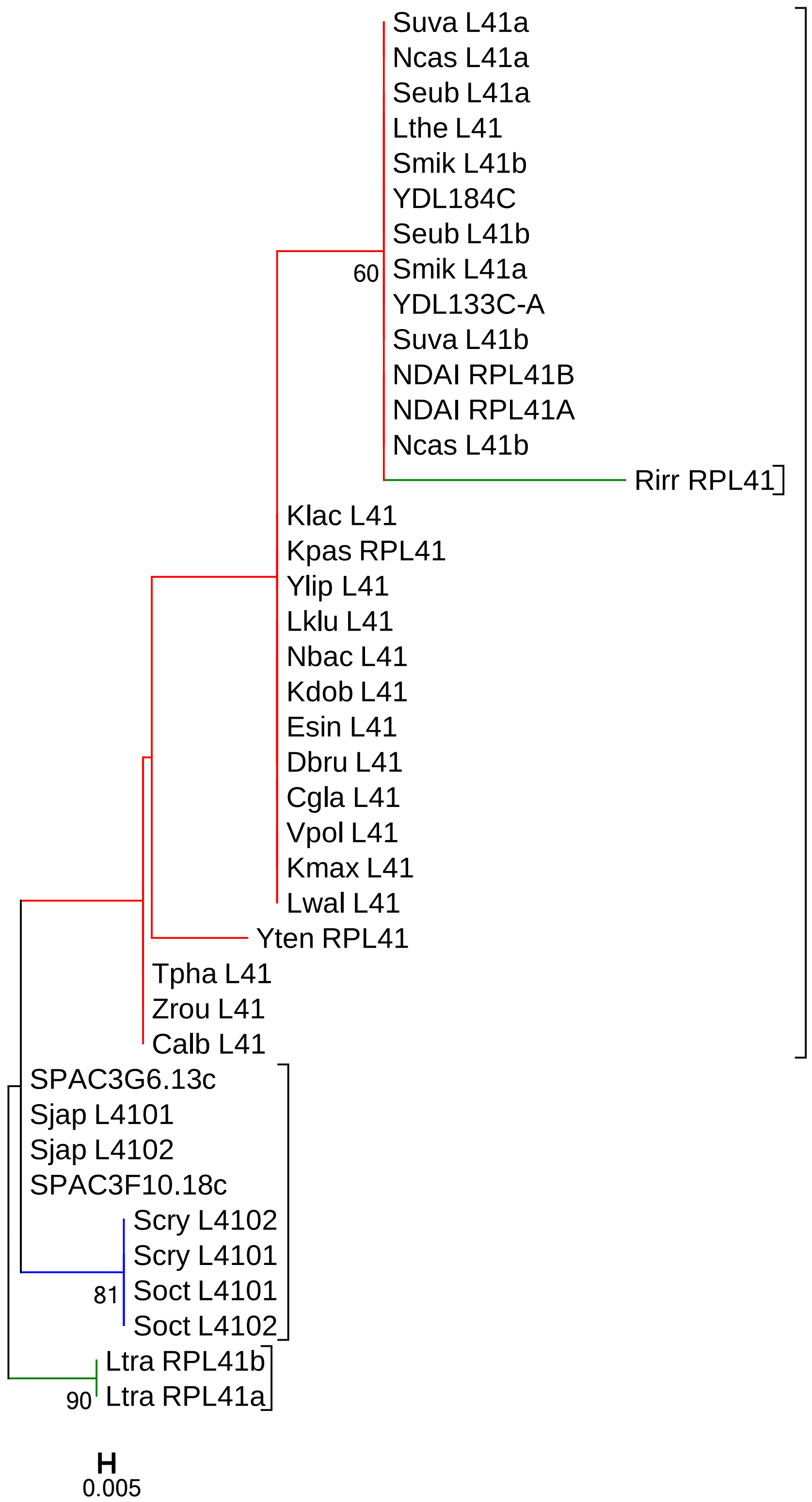
Suva L41a
Ncas L41a
Seub L41a
Lthe L41
Smik L41b
YDL184C
Seub L41b
60
Smik L41a
YDL133C-A
Suva L41b
NDAI RPL41B
NDAI RPL41A
Ncas L41b
Rirr RPL41
Klac L41
Kpas RPL41
Ylip L41
Lklu L41
Nbac L41
Kdob L41
Esin L41
Dbru L41
Cgla L41
Vpol L41
Kmax L41
Lwal L41
Yten RPL41
Tpha L41
Zrou L41
Calb L41
SPAC3G6.13c
Sjap L4101
Sjap L4102
SPAC3F10.18c
Scry L4102
Scry L4101
81
Soct L4101
Soct L4102
Ltra RPL41b
90
Ltra RPL41a
0.005

### RPL42 (RPL36A)

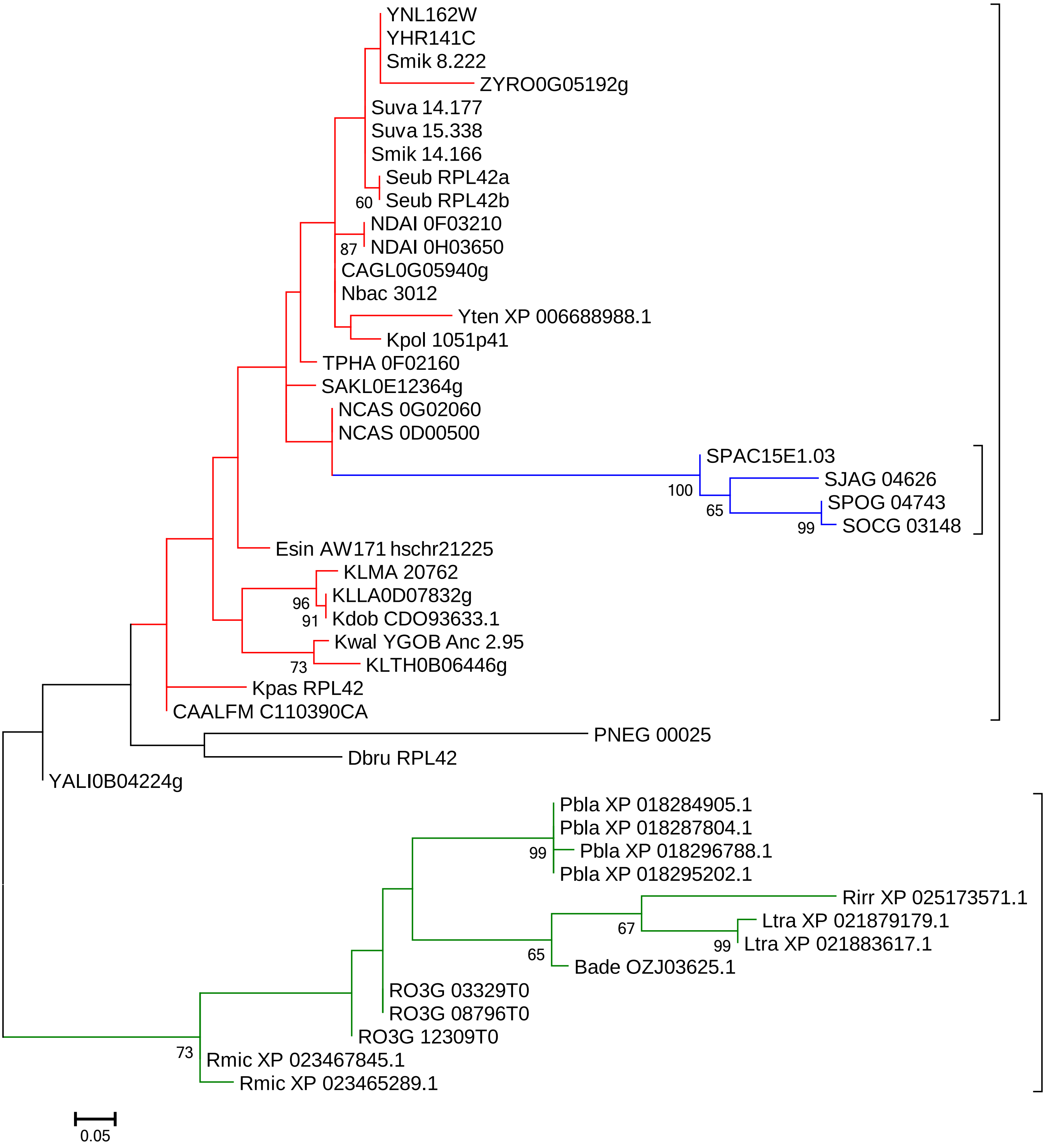

### RPL43 (RPL37A)

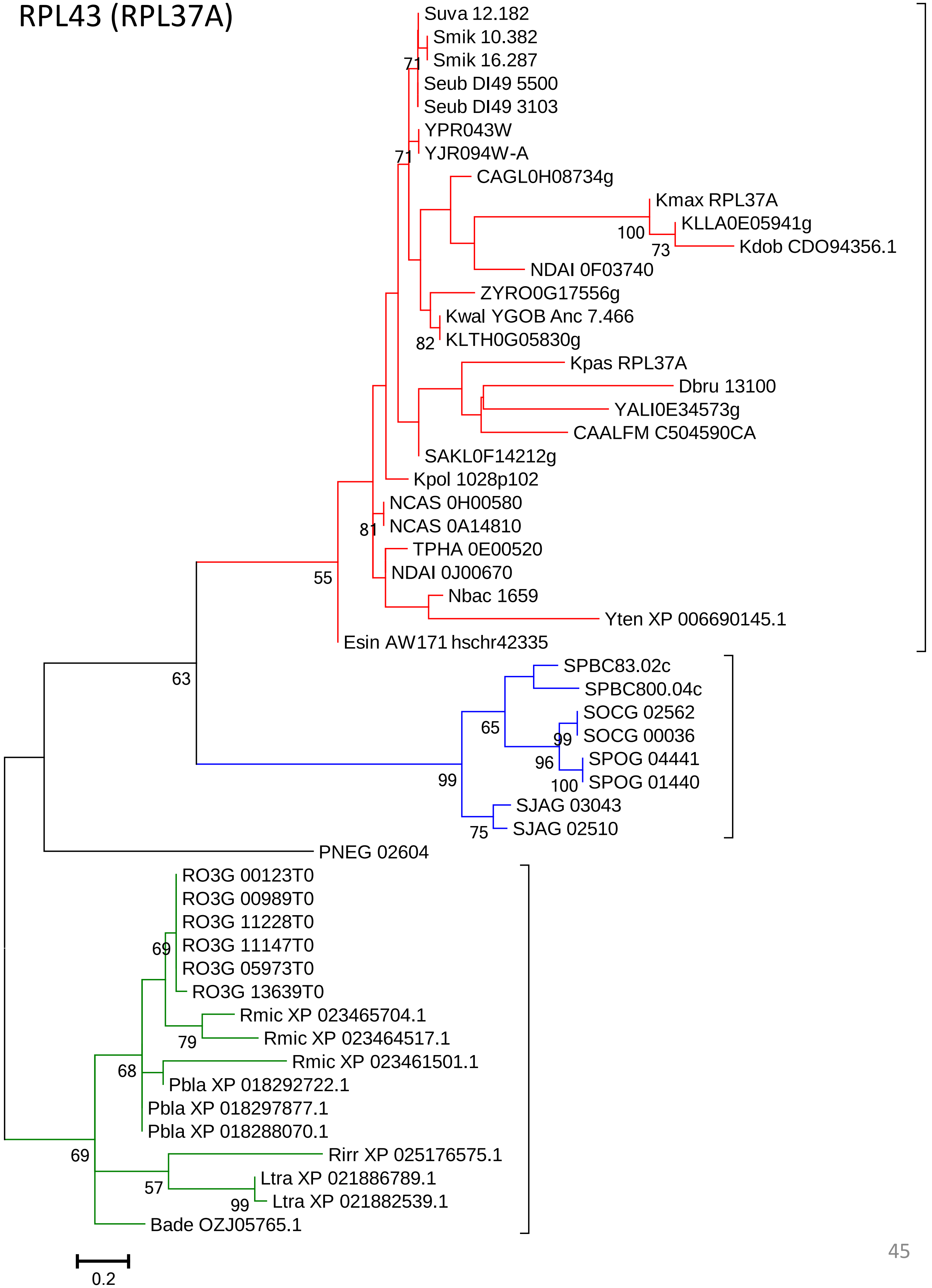

### RPP0 (RLP0)

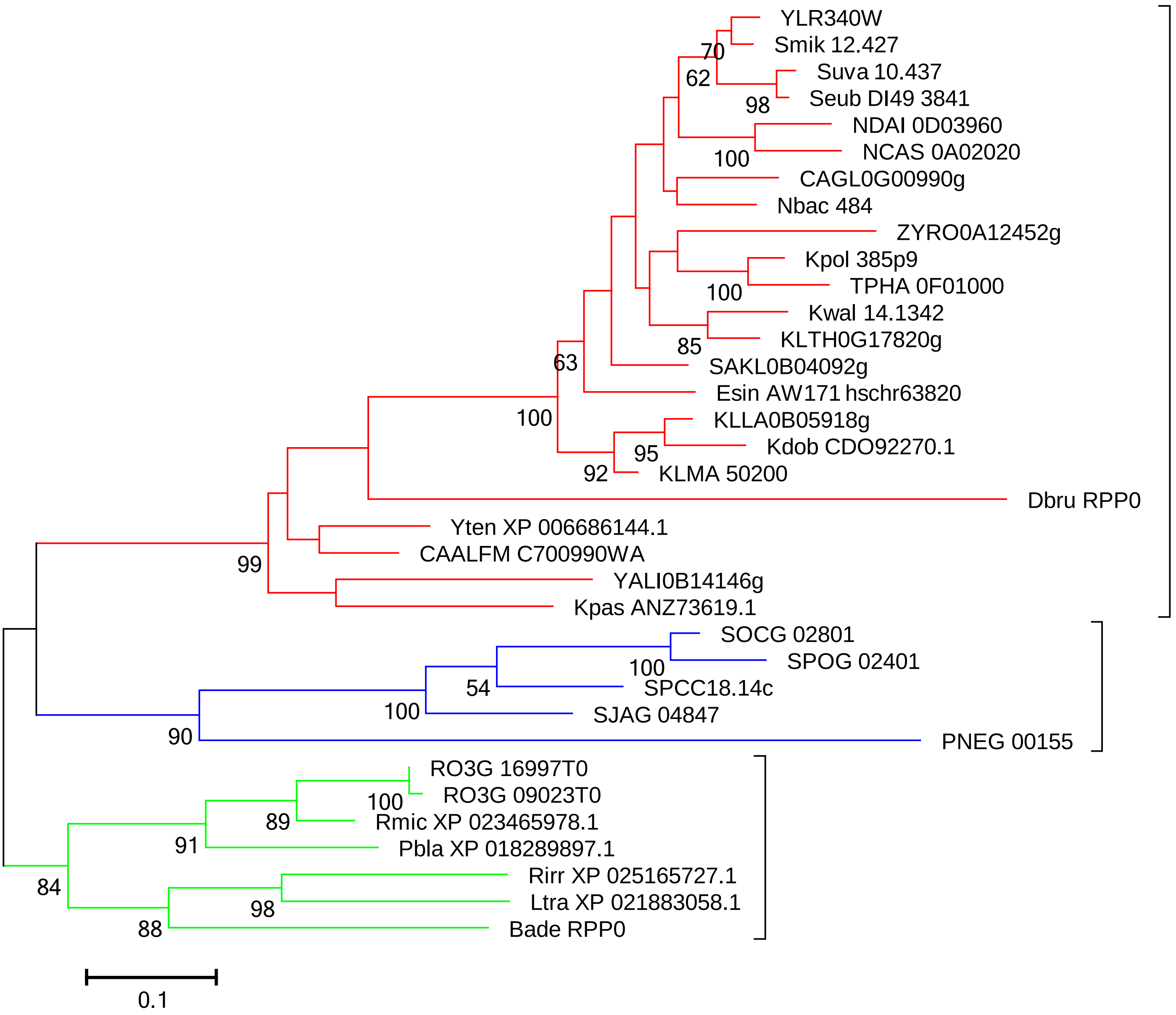
YLR340W
Smik 12.427
Suva 10.437
Seub DI49 3841
NDAI 0D03960
NCAS 0A02020
CAGL0G00990g
Nbac 484
ZYRO0A12452g
Kpol 385p9
TPHA 0F01000
Kwal 14.1342
KLTH0G17820g
SAKL0B04092g
Esin AW171 hschr63820
KLLA0B05918g
Kdob CDO92270.1
KLMA 50200
Dbru RPP0
Yten XP 006686144.1
CAALFM C700990WA
YALI0B14146g
Kpas ANZ73619.1
SOCG 02801
SPOG 02401
SPCC18.14c
SJAG 04847
PNEG 00155
RO3G 16997T0
RO3G 09023T0
Rmic XP 023465978.1
Pbla XP 018289897.1
Rirr XP 025165727.1
Ltra XP 021883058.1
Bade RPP0
70
62
98
100
100
85
63
100
95
92
99
100
54
100
90
100
89
91
84
98
88
0.1

### RPP1 (RLP1)

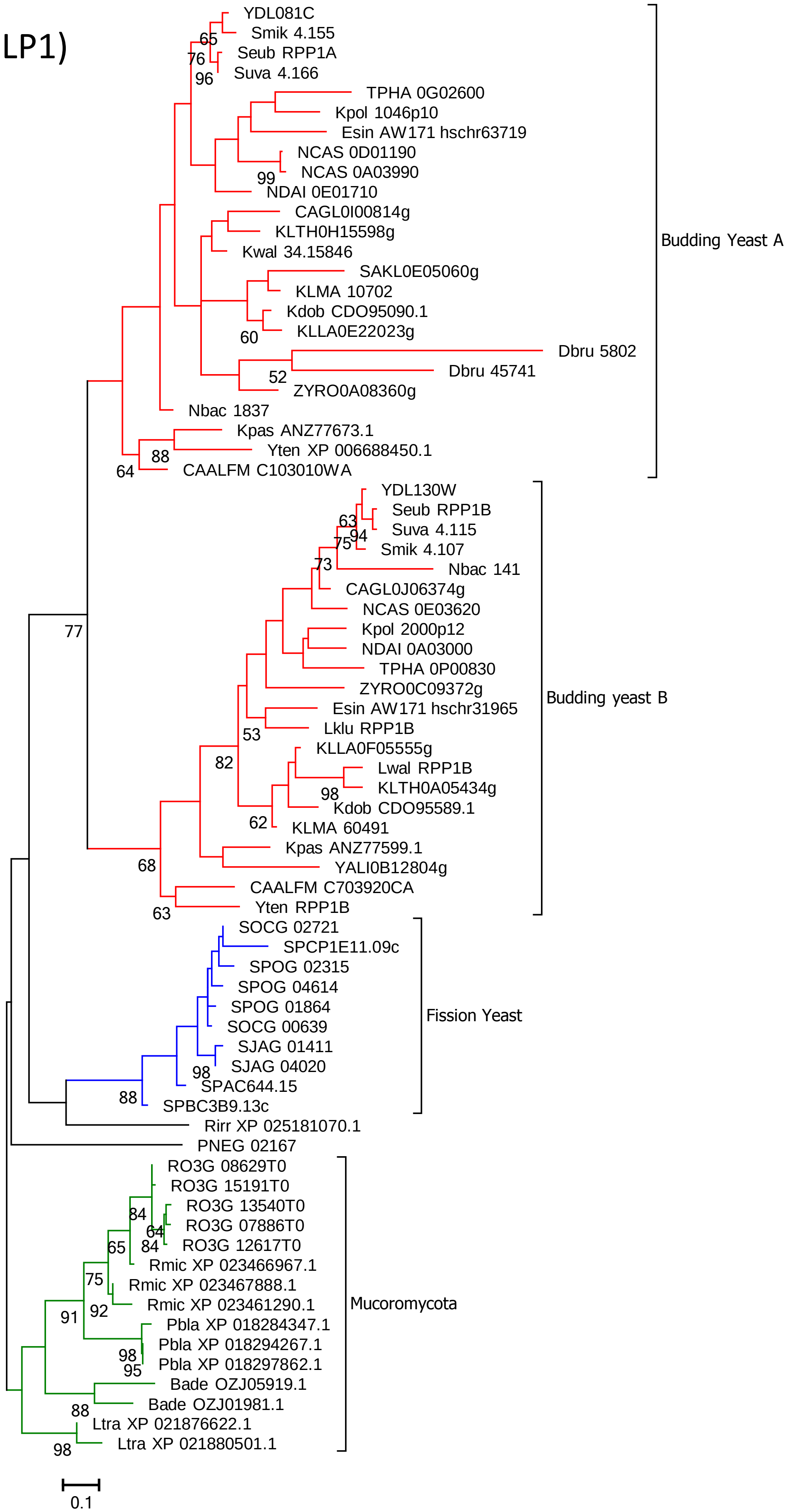
YDL081C
Smik 4.155
Seub RPP1A
Suva 4.166
TPHA 0G02600
Kpol 1046p10
Esin AW171 hschr63719
NCAS 0D01190
NCAS 0A03990
NDAI 0E01710
CAGL0I00814g
KLTH0H15598g
Kwal 34.15846
SAKL0E05060g
KLMA 10702
Kdob CDO95090.1
KLLA0E22023g
Dbru 5802
Dbru 45741
ZYRO0A08360g
Nbac 1837
Kpas ANZ77673.1
Yten XP 006688450.1
CAALFM C103010WA
Budding Yeast A
YDL130W
Seub RPP1B
Suva 4.115
Smik 4.107
Nbac 141
CAGL0J06374g
NCAS 0E03620
Kpol 2000p12
NDAI 0A03000
TPHA 0P00830
ZYRO0C09372g
Esin AW171 hschr31965
Lklu RPP1B
KLLA0F05555g
Lwal RPP1B
KLTH0A05434g
Kdob CDO95589.1
KLMA 60491
Kpas ANZ77599.1
YALI0B12804g
CAALFM C703920CA
Yten RPP1B
Budding yeast B
SOCG 02721
SPCP1E11.09c
SPOG 02315
SPOG 04614
SPOG 01864
SOCG 00639
SJAG 01411
SJAG 04020
SPAC644.15
SPBC3B9.13c
Fission Yeast
Rirr XP 025181070.1
PNEG 02167
RO3G 08629T0
RO3G 15191T0
RO3G 13540T0
RO3G 07886T0
RO3G 12617T0
Rmic XP 023466967.1
Rmic XP 023467888.1
Rmic XP 023461290.1
Pbla XP 018284347.1
Pbla XP 018294267.1
Pbla XP 018297862.1
Bade OZJ05919.1
Bade OZJ01981.1
Ltra XP 021876622.1
Ltra XP 021880501.1
Mucoromycota
65
76
96
99
60
52
88
64
63
94
75
73
77
53
82
98
62
68
63
98
88
84
64
84
65
75
91
92
98
95
88
98
0.1

### RPP2 (RLP2)

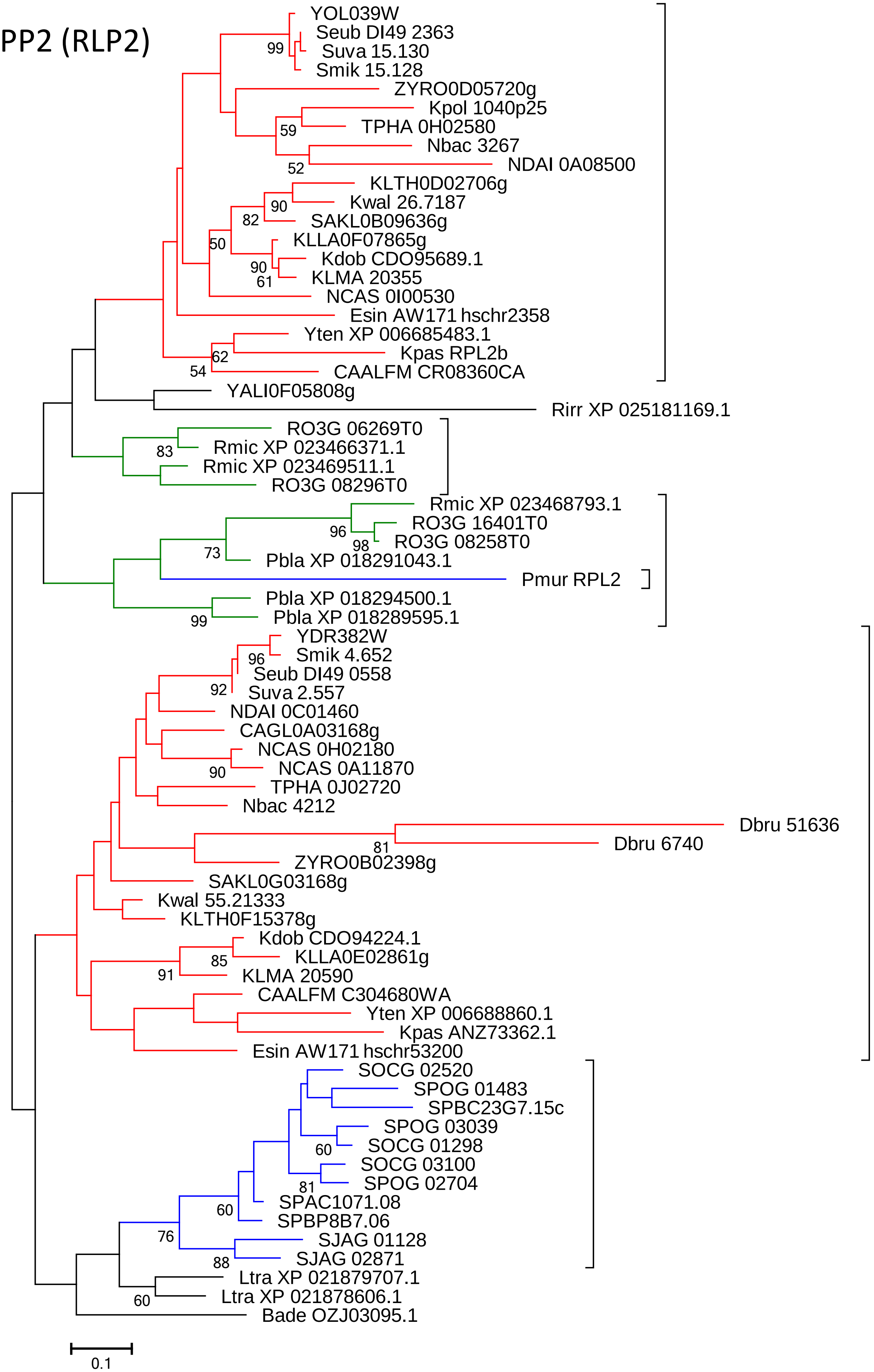

YOL039W
Seub DI49 2363
Suva 15.130
Smik 15.128
ZYRO0D05720g
Kpol 1040p25
TPHA 0H02580
Nbac 3267
NDAI 0A08500
KLTH0D02706g
Kwal 26.7187
SAKL0B09636g
KLLA0F07865g
Kdob CDO95689.1
KLMA 20355
NCAS 0I00530
Esin AW171 hschr2358
Yten XP 006685483.1
Kpas RPL2b
CAALFM CR08360CA
YALI0F05808g
Rirr XP 025181169.1
RO3G 06269T0
Rmic XP 023466371.1
Rmic XP 023469511.1
RO3G 08296T0
Rmic XP 023468793.1
RO3G 16401T0
RO3G 08258T0
Pbla XP 018291043.1
Pmur RPL2
Pbla XP 018294500.1
Pbla XP 018289595.1
YDR382W
Smik 4.652
Seub DI49 0558
Suva 2.557
NDAI 0C01460
CAGL0A03168g
NCAS 0H02180
NCAS 0A11870
TPHA 0J02720
Nbac 4212
Dbru 51636
Dbru 6740
ZYRO0B02398g
SAKL0G03168g
Kwal 55.21333
KLTH0F15378g
Kdob CDO94224.1
KLLA0E02861g
KLMA 20590
CAALFM C304680WA
Yten XP 006688860.1
Kpas ANZ73362.1
Esin AW171 hschr53200
SOCG 02520
SPOG 01483
SPBC23G7.15c
SPOG 03039
SOCG 01298
SOCG 03100
SPOG 02704
SPAC1071.08
SPBP8B7.06
SJAG 01128
SJAG 02871
Ltra XP 021879707.1
Ltra XP 021878606.1
Bade OZJ03095.1
99
59
52
90
82
50
90
61
62
54
83
96
98
73
99
96
92
90
81
85
91
60
81
60
76
88
60
0.1

### RPS0 (RPSA)

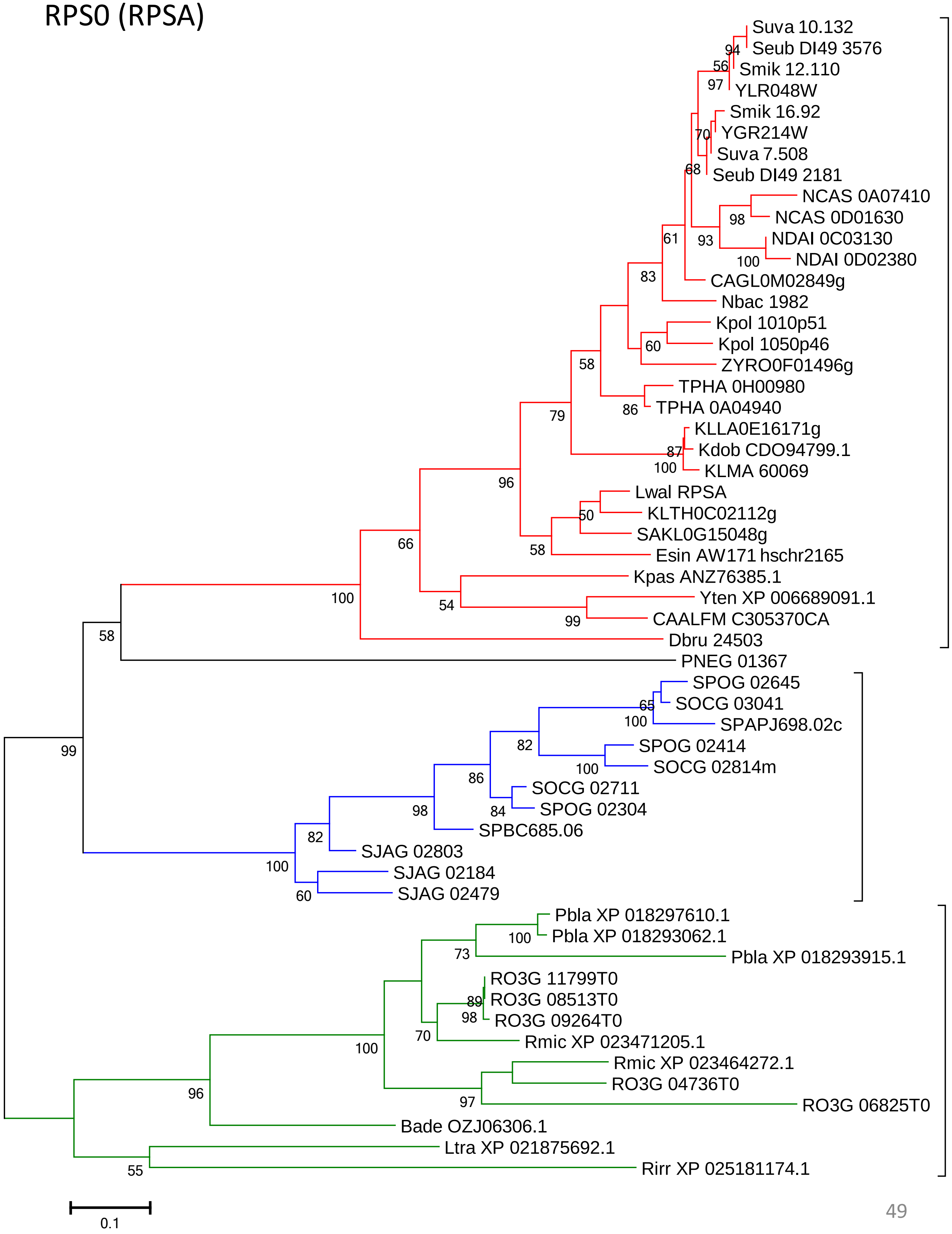

Suva 10.132
94
Seub DI49 3576
56
Smik 12.110
97
YLR048W
Smik 16.92
70
YGR214W
Suva 7.508
68
Seub DI49 2181
NCAS 0A07410
98
NCAS 0D01630
61
93
NDAI 0C03130
100
NDAI 0D02380
83
CAGL0M02849g
Nbac 1982
Kpol 1010p51
60
Kpol 1050p46
58
ZYRO0F01496g
TPHA 0H00980
86
TPHA 0A04940
79
KLLA0E16171g
87
Kdob CDO94799.1
100
KLMA 60069
96
Lwal RPSA
50
KLTH0C02112g
66
SAKL0G15048g
58
Esin AW171 hschr2165
Kpas ANZ76385.1
100
54
Yten XP 006689091.1
99
CAALFM C305370CA
58
Dbru 24503
PNEG 01367
SPOG 02645
65
SOCG 03041
100
SPAPJ698.02c
82
SPOG 02414
99
100
SOCG 02814m
86
SOCG 02711
84
SPOG 02304
98
SPBC685.06
82
SJAG 02803
100
SJAG 02184
60
SJAG 02479
Pbla XP 018297610.1
100
Pbla XP 018293062.1
73
Pbla XP 018293915.1
RO3G 11799T0
89
RO3G 08513T0
98
RO3G 09264T0
70
Rmic XP 023471205.1
100
Rmic XP 023464272.1
RO3G 04736T0
97
RO3G 06825T0
96
Bade OZJ06306.1
Ltra XP 021875692.1
55
Rirr XP 025181174.1
0.1

### RPS1 (RPS3A)

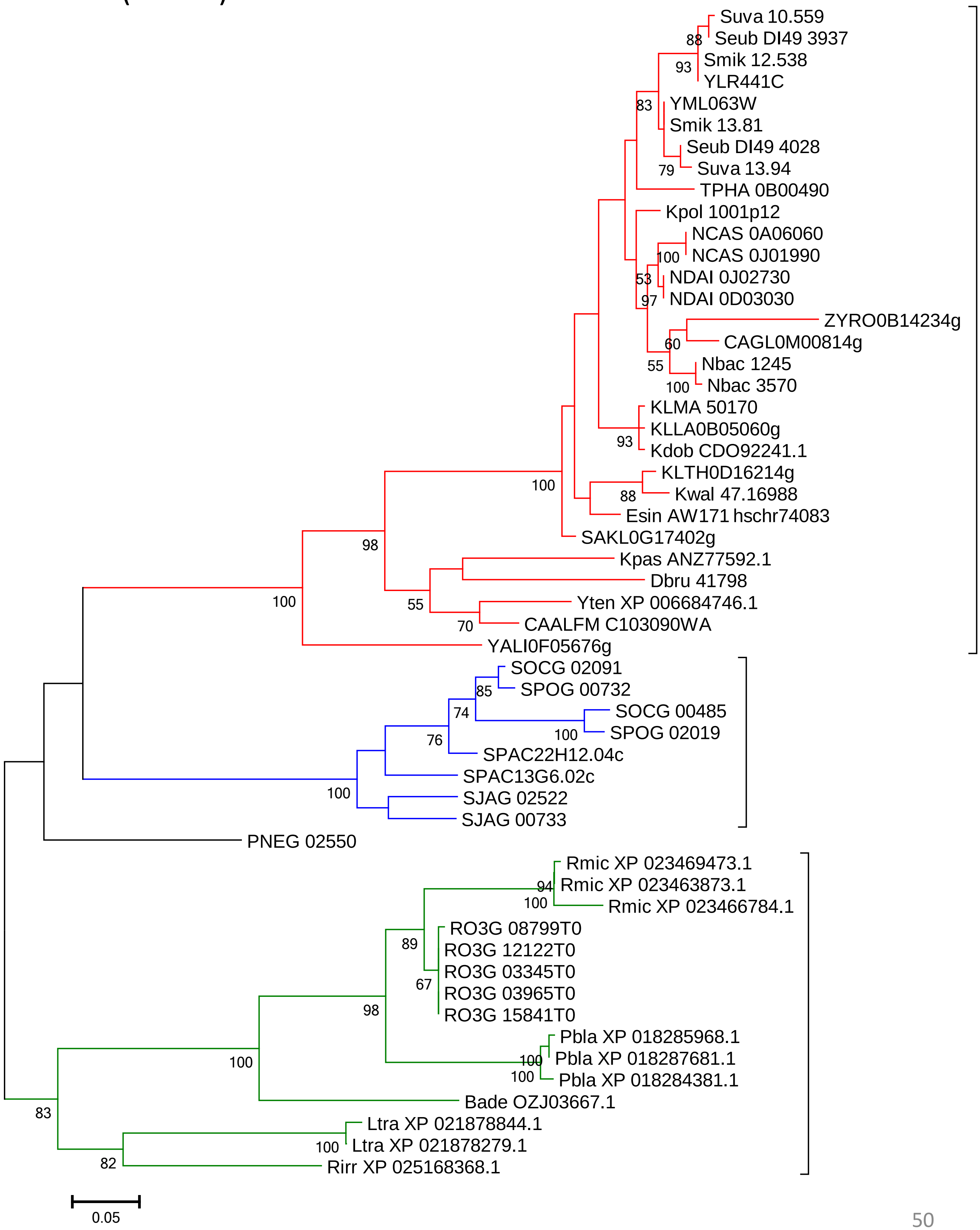
Suva 10.559
Seub DI49 3937
Smik 12.538
YLR441C
YML063W
Smik 13.81
Seub DI49 4028
Suva 13.94
TPHA 0B00490
Kpol 1001p12
NCAS 0A06060
NCAS 0J01990
NDAI 0J02730
NDAI 0D03030
ZYRO0B14234g
CAGL0M00814g
Nbac 1245
Nbac 3570
KLMA 50170
KLLA0B05060g
Kdob CDO92241.1
KLTH0D16214g
Kwal 47.16988
Esin AW171 hschr74083
SAKL0G17402g
Kpas ANZ77592.1
Dbru 41798
Yten XP 006684746.1
CAALFM C103090WA
YALI0F05676g
SOCG 02091
SPOG 00732
SOCG 00485
SPOG 02019
SPAC22H12.04c
SPAC13G6.02c
SJAG 02522
SJAG 00733
PNEG 02550
Rmic XP 023469473.1
Rmic XP 023463873.1
Rmic XP 023466784.1
RO3G 08799T0
RO3G 12122T0
RO3G 03345T0
RO3G 03965T0
RO3G 15841T0
Pbla XP 018285968.1
Pbla XP 018287681.1
Pbla XP 018284381.1
Bade OZJ03667.1
Ltra XP 021878844.1
Ltra XP 021878279.1
Rirr XP 025168368.1
0.05

### RPS2

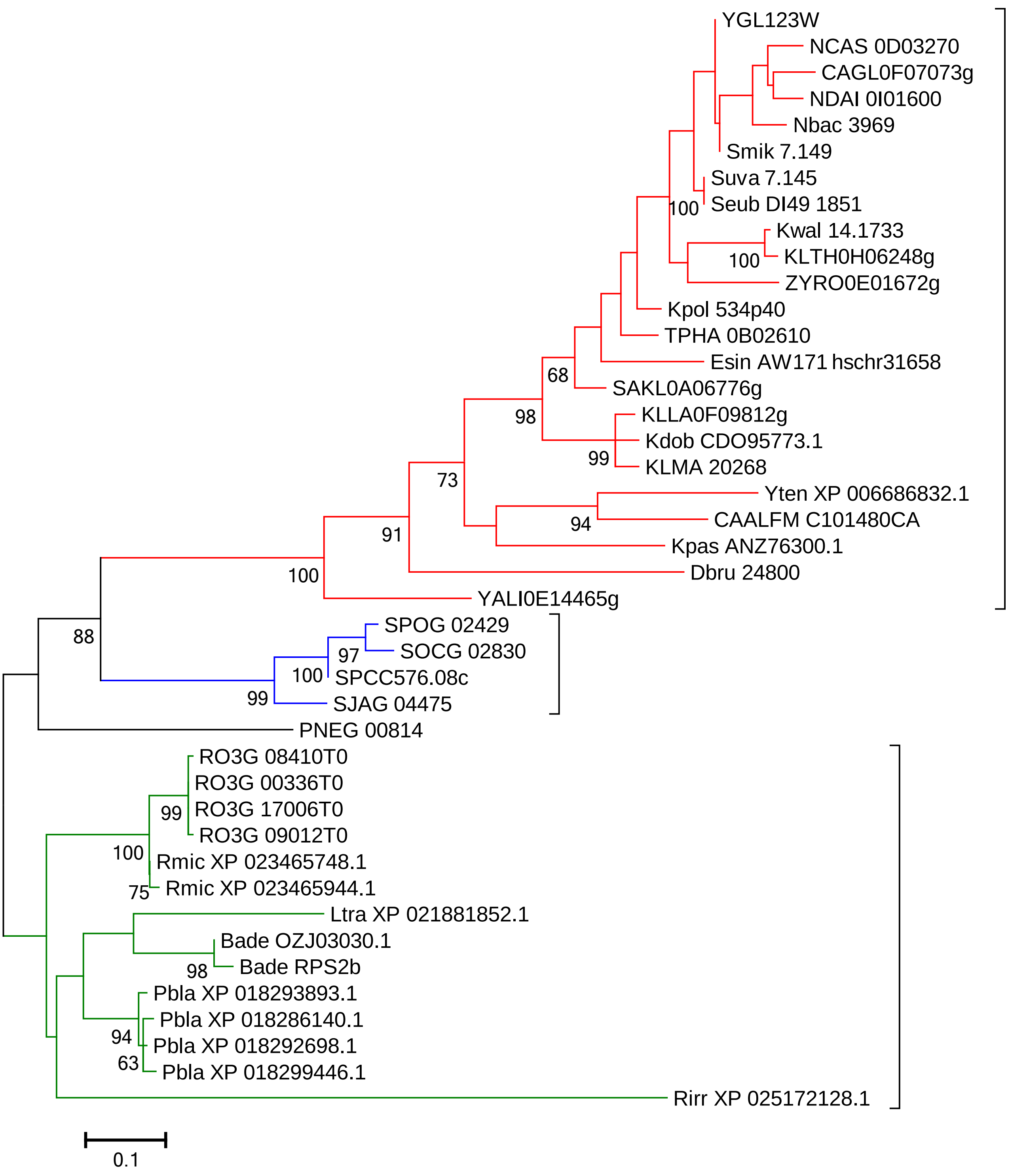
YGL123W
NCAS 0D03270
CAGL0F07073g
NDAI 0I01600
Nbac 3969
Smik 7.149
Suva 7.145
Seub DI49 1851
100
Kwal 14.1733
100
KLTH0H06248g
ZYRO0E01672g
Kpol 534p40
TPHA 0B02610
Esin AW171 hschr31658
68
SAKL0A06776g
98
KLLA0F09812g
Kdob CDO95773.1
99
KLMA 20268
73
Yten XP 006686832.1
94
CAALFM C101480CA
91
Kpas ANZ76300.1
100
Dbru 24800
YALI0E14465g
88
SPOG 02429
97
SOCG 02830
100
SPCC576.08c
99
SJAG 04475
PNEG 00814
RO3G 08410T0
RO3G 00336T0
99
RO3G 17006T0
RO3G 09012T0
100
Rmic XP 023465748.1
75
Rmic XP 023465944.1
Ltra XP 021881852.1
Bade OZJ03030.1
98
Bade RPS2b
Pbla XP 018293893.1
Pbla XP 018286140.1
94
Pbla XP 018292698.1
63
Pbla XP 018299446.1
Rirr XP 025172128.1
0.1

### RPS3

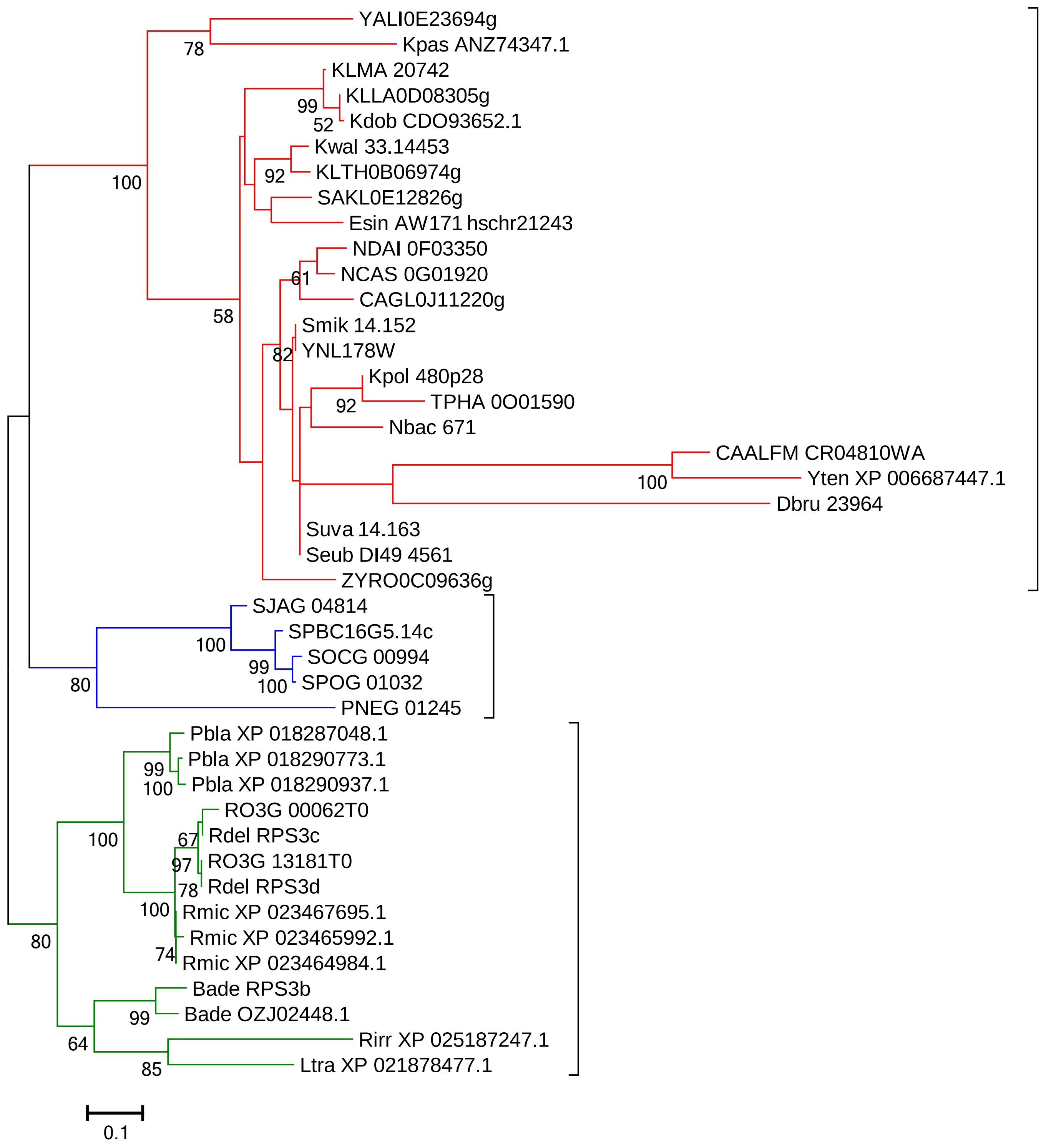
YALI0E23694g
Kpas ANZ74347.1
KLMA 20742
KLLA0D08305g
Kdob CDO93652.1
Kwal 33.14453
KLTH0B06974g
SAKL0E12826g
Esin AW171 hschr21243
NDAI 0F03350
NCAS 0G01920
CAGL0J11220g
Smik 14.152
YNL178W
Kpol 480p28
TPHA 0O01590
Nbac 671
CAALFM CR04810WA
Yten XP 006687447.1
Dbru 23964
Suva 14.163
Seub DI49 4561
ZYRO0C09636g
SJAG 04814
SPBC16G5.14c
SOCG 00994
SPOG 01032
PNEG 01245
Pbla XP 018287048.1
Pbla XP 018290773.1
Pbla XP 018290937.1
RO3G 00062T0
Rdel RPS3c
RO3G 13181T0
Rdel RPS3d
Rmic XP 023467695.1
Rmic XP 023465992.1
Rmic XP 023464984.1
Bade RPS3b
Bade OZJ02448.1
Rirr XP 025187247.1
Ltra XP 021878477.1
78
99
52
100
92
61
58
82
92
100
100
99
100
80
99
100
67
100
97
78
100
74
80
99
64
85
0.1

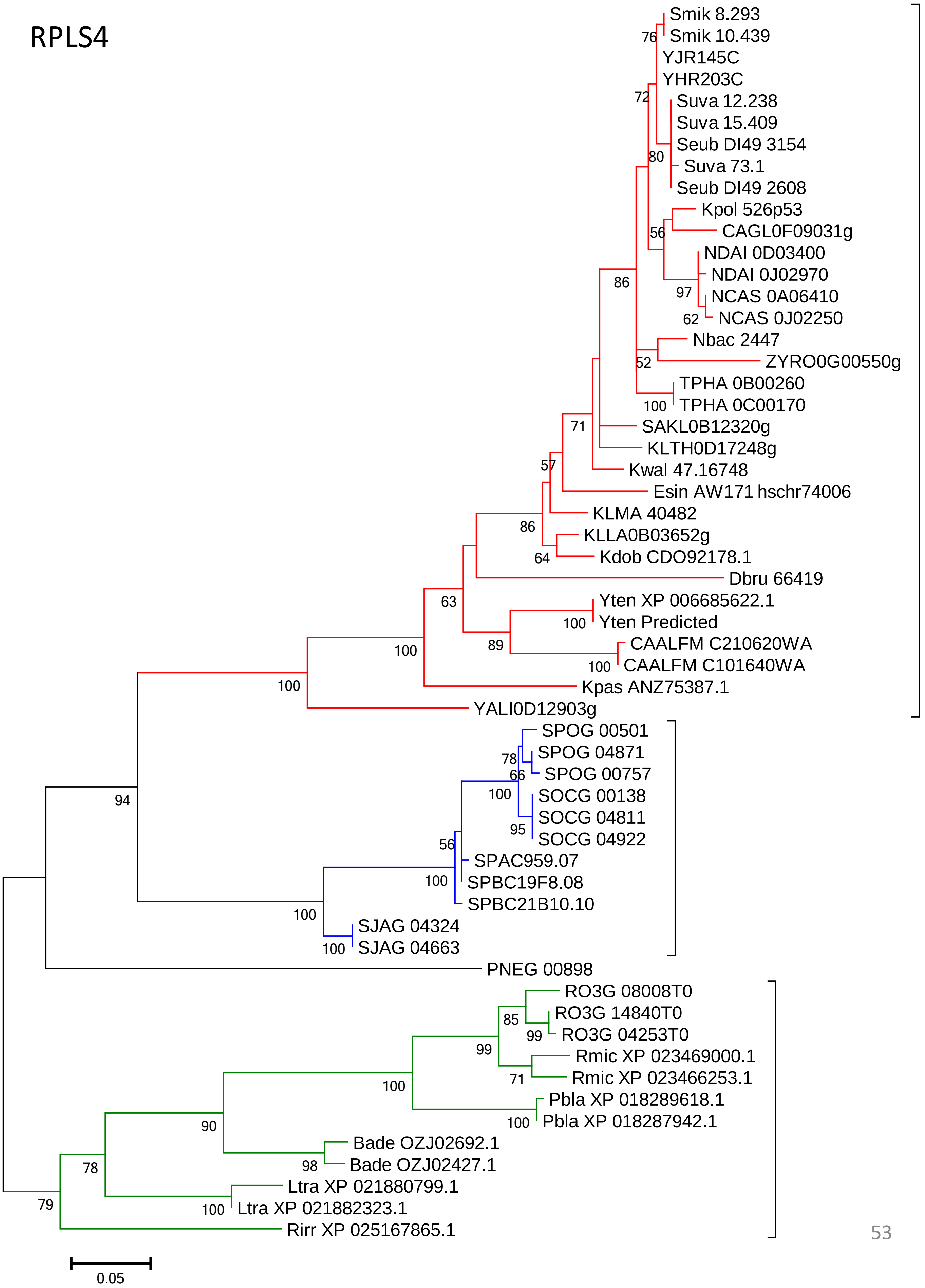

RPLS4
Smik 8.293
Smik 10.439
YJR145C
YHR203C
Suva 12.238
Suva 15.409
Seub DI49 3154
Suva 73.1
Seub DI49 2608
Kpol 526p53
CAGL0F09031g
NDAI 0D03400
NDAI 0J02970
NCAS 0A06410
NCAS 0J02250
Nbac 2447
ZYRO0G00550g
TPHA 0B00260
TPHA 0C00170
SAKL0B12320g
KLTH0D17248g
Kwal 47.16748
Esin AW171 hschr74006
KLMA 40482
KLLA0B03652g
Kdob CDO92178.1
Dbru 66419
Yten XP 006685622.1
Yten Predicted
CAALFM C210620WA
CAALFM C101640WA
Kpas ANZ75387.1
YALI0D12903g
SPOG 00501
SPOG 04871
SPOG 00757
SOCG 00138
SOCG 04811
SOCG 04922
SPAC959.07
SPBC19F8.08
SPBC21B10.10
SJAG 04324
SJAG 04663
PNEG 00898
RO3G 08008T0
RO3G 14840T0
RO3G 04253T0
Rmic XP 023469000.1
Rmic XP 023466253.1
Pbla XP 018289618.1
Pbla XP 018287942.1
Bade OZJ02692.1
Bade OZJ02427.1
Ltra XP 021880799.1
Ltra XP 021882323.1
Rirr XP 025167865.1
0.05

### RPS5

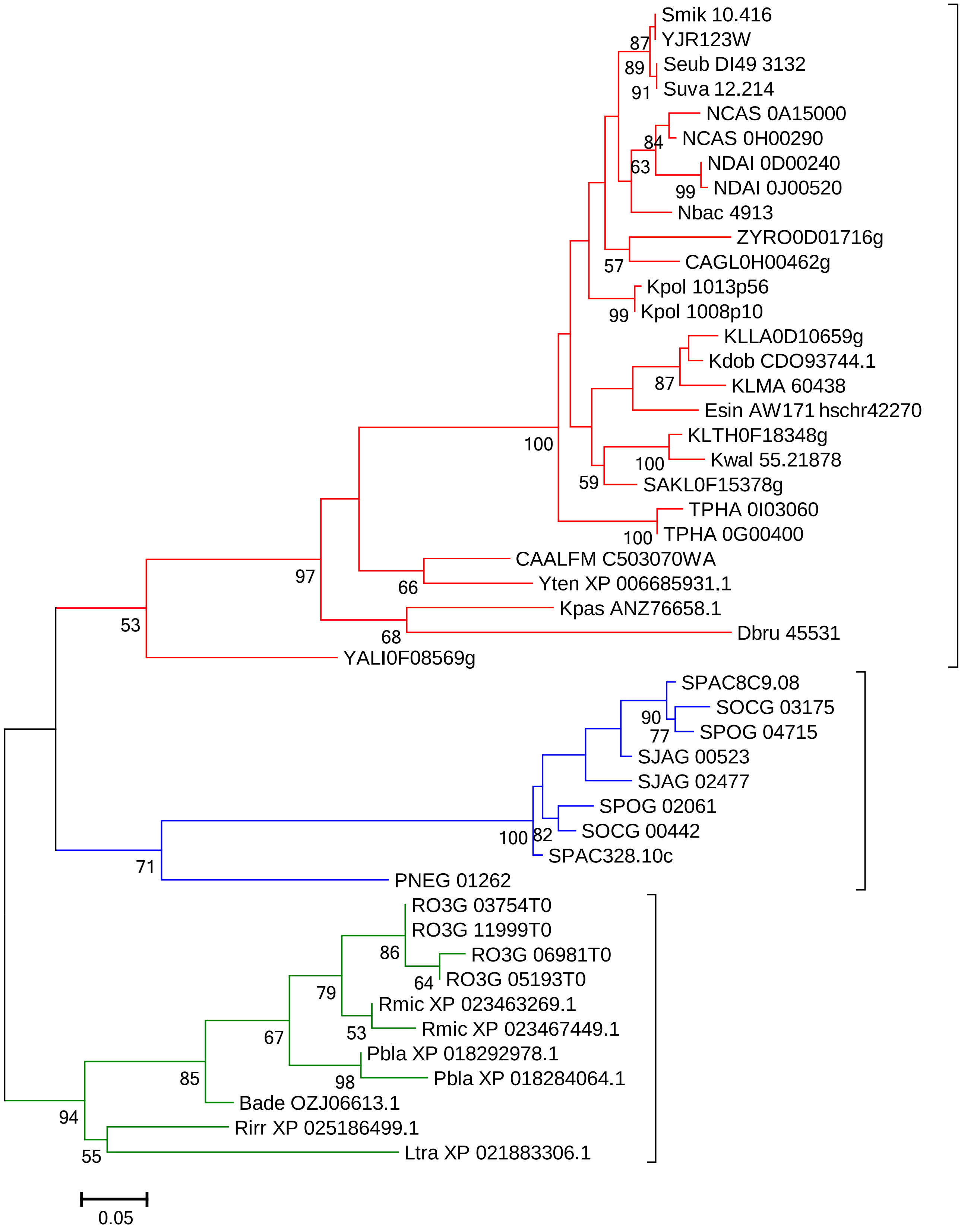
Smik 10.416
YJR123W
Seub DI49 3132
Suva 12.214
NCAS 0A15000
NCAS 0H00290
NDAI 0D00240
NDAI 0J00520
Nbac 4913
ZYRO0D01716g
CAGL0H00462g
Kpol 1013p56
Kpol 1008p10
KLLA0D10659g
Kdob CDO93744.1
KLMA 60438
Esin AW171 hschr42270
KLTH0F18348g
Kwal 55.21878
SAKL0F15378g
TPHA 0I03060
TPHA 0G00400
CAALFM C503070WA
Yten XP 006685931.1
Kpas ANZ76658.1
Dbru 45531
YALI0F08569g
SPAC8C9.08
SOCG 03175
SPOG 04715
SJAG 00523
SJAG 02477
SPOG 02061
SOCG 00442
SPAC328.10c
PNEG 01262
RO3G 03754T0
RO3G 11999T0
RO3G 06981T0
RO3G 05193T0
Rmic XP 023463269.1
Rmic XP 023467449.1
Pbla XP 018292978.1
Pbla XP 018284064.1
Bade OZJ06613.1
Rirr XP 025186499.1
Ltra XP 021883306.1
87
89
91
84
63
99
57
99
87
100
100
59
100
97
66
68
53
90
77
100
82
71
86
64
79
53
67
98
85
94
55
0.05

### RPS6

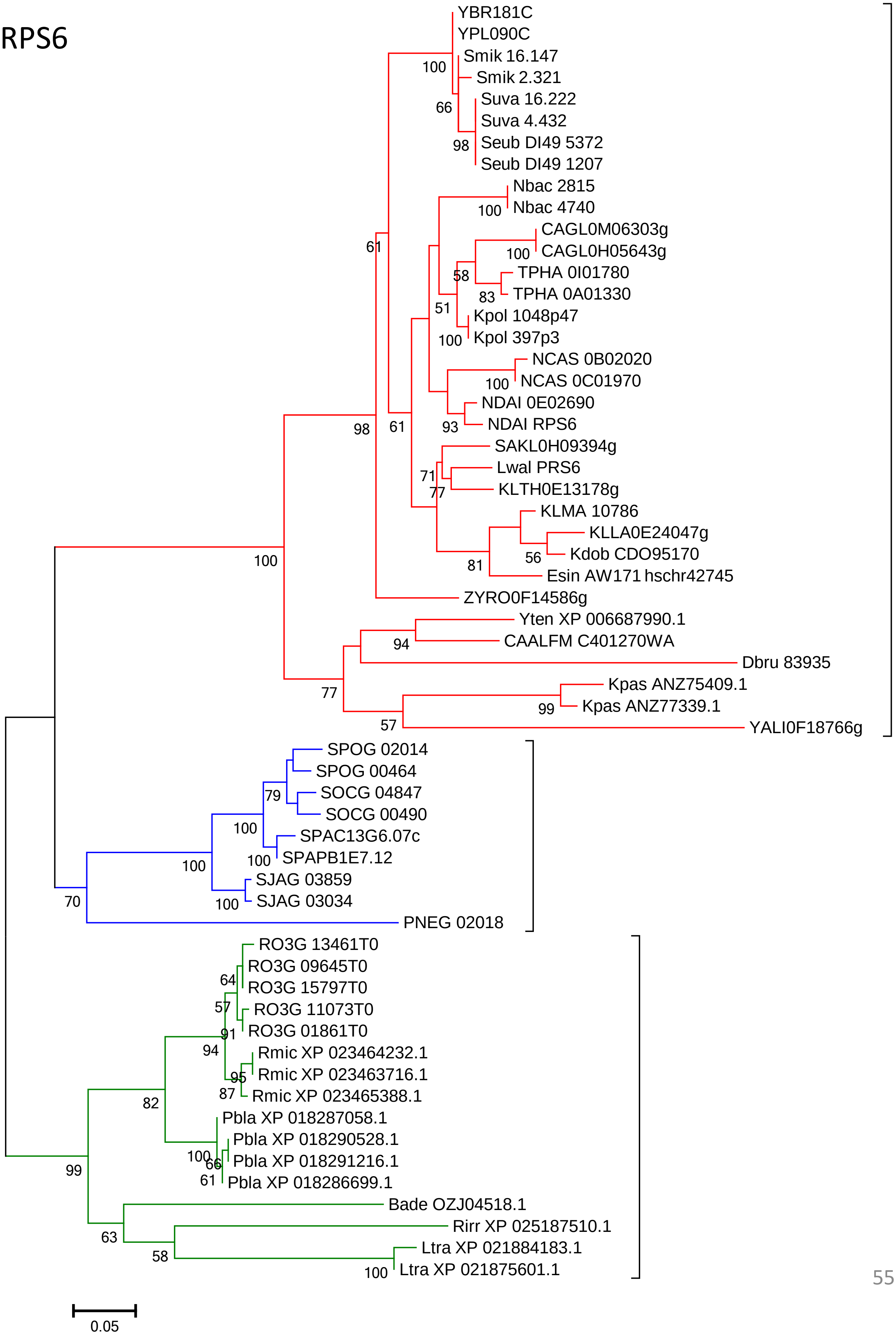
YBR181C
YPL090C
Smik 16.147
Smik 2.321
Suva 16.222
Suva 4.432
Seub DI49 5372
Seub DI49 1207
Nbac 2815
Nbac 4740
CAGL0M06303g
CAGL0H05643g
TPHA 0I01780
TPHA 0A01330
Kpol 1048p47
Kpol 397p3
NCAS 0B02020
NCAS 0C01970
NDAI 0E02690
NDAI RPS6
SAKL0H09394g
Lwal PRS6
KLTH0E13178g
KLMA 10786
KLLA0E24047g
Kdob CDO95170
Esin AW171 hschr42745
ZYRO0F14586g
Yten XP 006687990.1
CAALFM C401270WA
Dbru 83935
Kpas ANZ75409.1
Kpas ANZ77339.1
YALI0F18766g
SPOG 02014
SPOG 00464
SOCG 04847
SOCG 00490
SPAC13G6.07c
SPAPB1E7.12
SJAG 03859
SJAG 03034
PNEG 02018
RO3G 13461T0
RO3G 09645T0
RO3G 15797T0
RO3G 11073T0
RO3G 01861T0
Rmic XP 023464232.1
Rmic XP 023463716.1
Rmic XP 023465388.1
Pbla XP 018287058.1
Pbla XP 018290528.1
Pbla XP 018291216.1
Pbla XP 018286699.1
Bade OZJ04518.1
Rirr XP 025187510.1
Ltra XP 021884183.1
Ltra XP 021875601.1
0.05

### RPS7

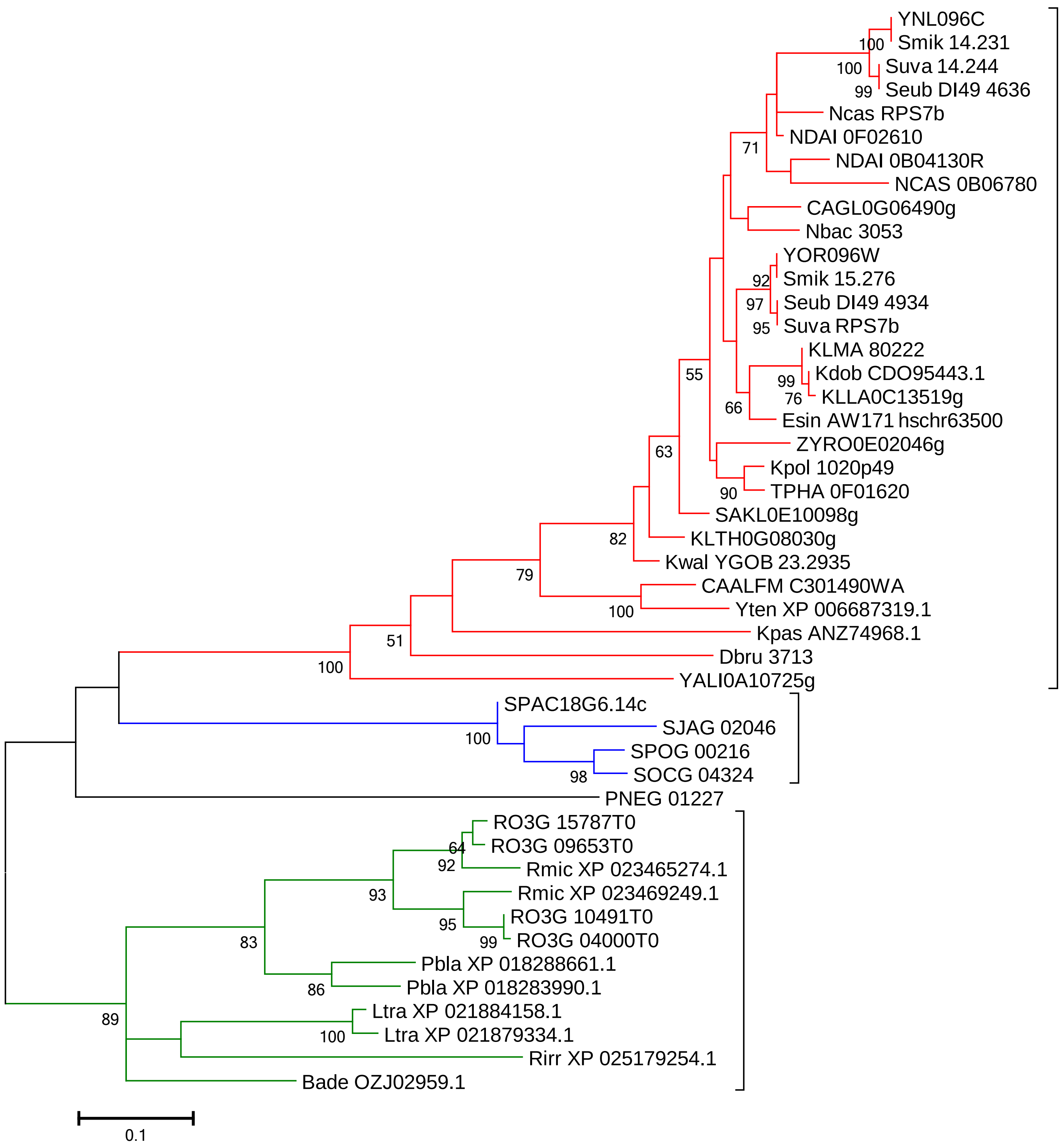
YNL096C
Smik 14.231
Suva 14.244
Seub DI49 4636
Ncas RPS7b
NDAI 0F02610
NDAI 0B04130R
NCAS 0B06780
CAGL0G06490g
Nbac 3053
YOR096W
Smik 15.276
Seub DI49 4934
Suva RPS7b
KLMA 80222
Kdob CDO95443.1
KLLA0C13519g
Esin AW171 hschr63500
ZYRO0E02046g
Kpol 1020p49
TPHA 0F01620
SAKL0E10098g
KLTH0G08030g
Kwal YGOB 23.2935
CAALFM C301490WA
Yten XP 006687319.1
Kpas ANZ74968.1
Dbru 3713
YALI0A10725g
SPAC18G6.14c
SJAG 02046
SPOG 00216
SOCG 04324
PNEG 01227
RO3G 15787T0
RO3G 09653T0
Rmic XP 023465274.1
Rmic XP 023469249.1
RO3G 10491T0
RO3G 04000T0
Pbla XP 018288661.1
Pbla XP 018283990.1
Ltra XP 021884158.1
Ltra XP 021879334.1
Rirr XP 025179254.1
Bade OZJ02959.1
100
100
99
71
92
97
95
99
76
66
55
90
63
82
79
100
51
100
100
98
64
92
93
95
99
83
86
100
89
0.1

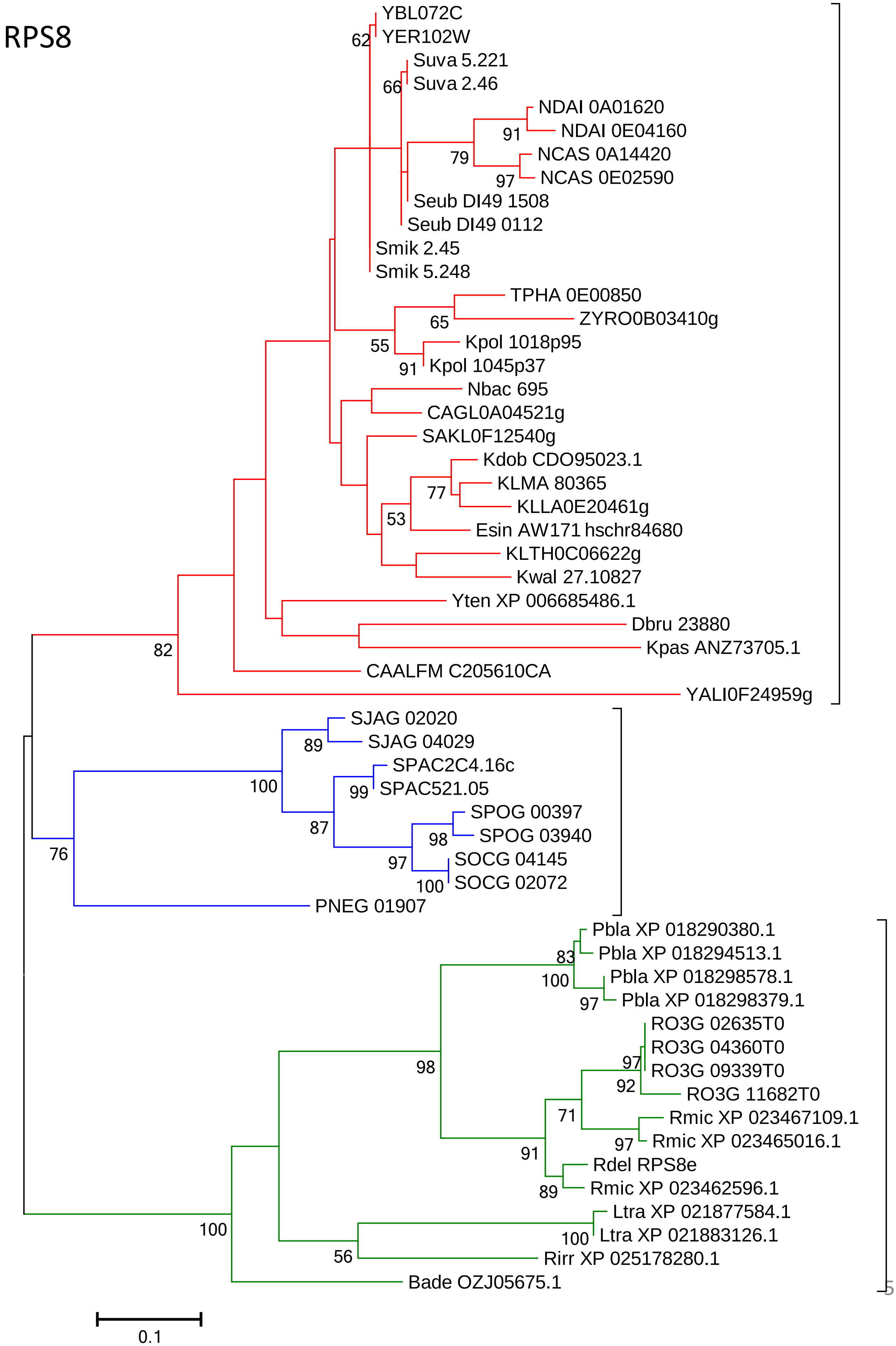
RPS8
YBL072C
YER102W
62
Suva 5.221
Suva 2.46
66
NDAI 0A01620
NDAI 0E04160
91
NCAS 0A14420
79
NCAS 0E02590
97
Seub DI49 1508
Seub DI49 0112
Smik 2.45
Smik 5.248
TPHA 0E00850
ZYRO0B03410g
65
Kpol 1018p95
55
Kpol 1045p37
91
Nbac 695
CAGL0A04521g
SAKL0F12540g
Kdob CDO95023.1
KLMA 80365
77
KLLA0E20461g
53
Esin AW171 hschr84680
KLTH0C06622g
Kwal 27.10827
Yten XP 006685486.1
Dbru 23880
Kpas ANZ73705.1
82
CAALFM C205610CA
YALI0F24959g
SJAG 02020
89
SJAG 04029
SPAC2C4.16c
100
99
SPAC521.05
SPOG 00397
87
SPOG 03940
98
SOCG 04145
97
76
100
SOCG 02072
PNEG 01907
Pbla XP 018290380.1
Pbla XP 018294513.1
83
Pbla XP 018298578.1
100
Pbla XP 018298379.1
97
RO3G 02635T0
RO3G 04360T0
97
RO3G 09339T0
98
92
RO3G 11682T0
71
Rmic XP 023467109.1
97
Rmic XP 023465016.1
91
Rdel RPS8e
89
Rmic XP 023462596.1
Ltra XP 021877584.1
100
100
Ltra XP 021883126.1
56
Rirr XP 025178280.1
Bade OZJ05675.1
0.1

### RPS9

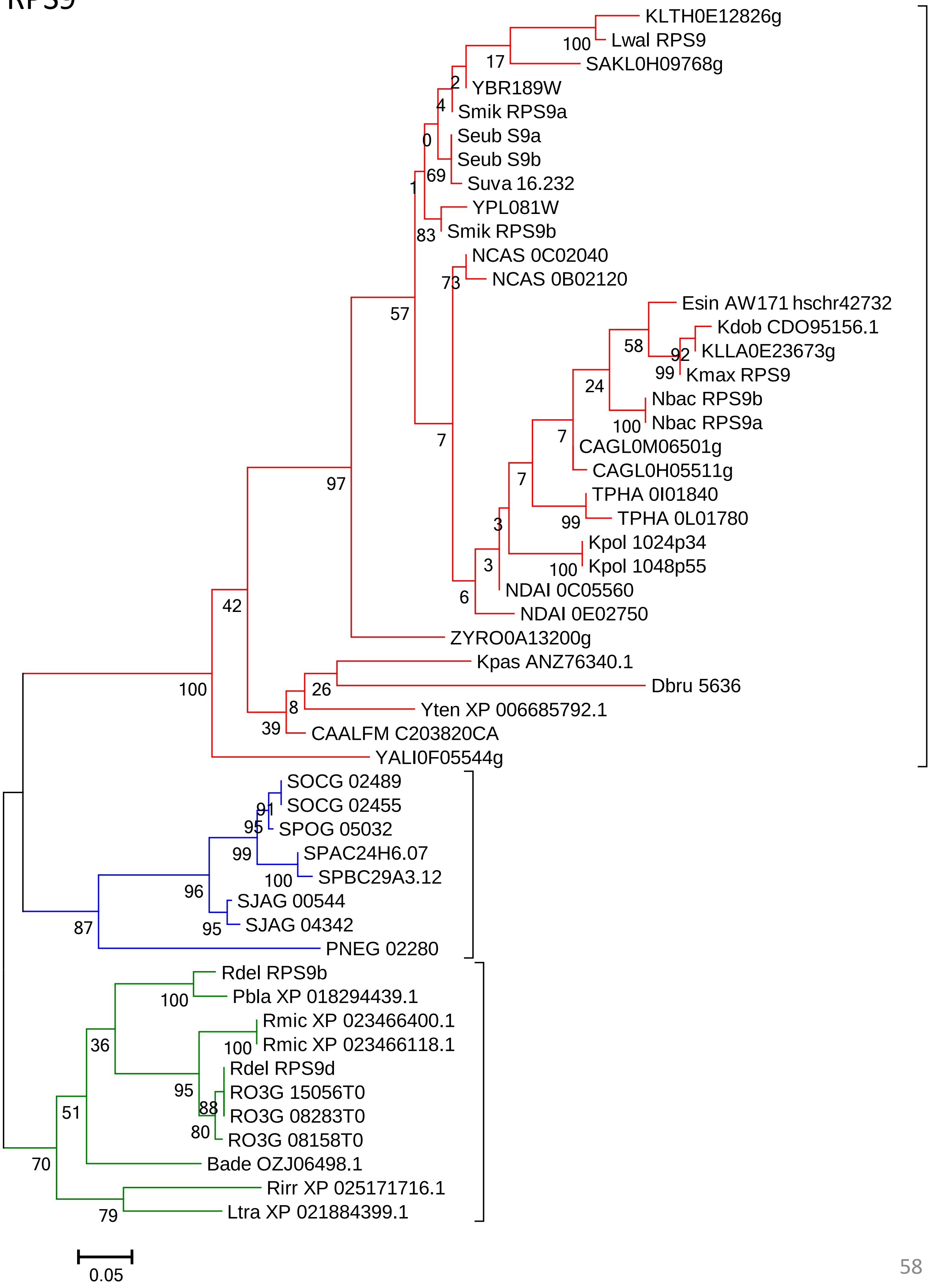

KLTH0E12826g
Lwal RPS9
SAKL0H09768g
YBR189W
Smik RPS9a
Seub S9a
Seub S9b
Suva 16.232
YPL081W
Smik RPS9b
NCAS 0C02040
NCAS 0B02120
Esin AW171 hschr42732
Kdob CDO95156.1
KLLA0E23673g
Kmax RPS9
Nbac RPS9b
Nbac RPS9a
CAGL0M06501g
CAGL0H05511g
TPHA 0I01840
TPHA 0L01780
Kpol 1024p34
Kpol 1048p55
NDAI 0C05560
NDAI 0E02750
ZYRO0A13200g
Kpas ANZ76340.1
Dbru 5636
Yten XP 006685792.1
CAALFM C203820CA
YALI0F05544g
SOCG 02489
SOCG 02455
SPOG 05032
SPAC24H6.07
SPBC29A3.12
SJAG 00544
SJAG 04342
PNEG 02280
Rdel RPS9b
Pbla XP 018294439.1
Rmic XP 023466400.1
Rmic XP 023466118.1
Rdel RPS9d
RO3G 15056T0
RO3G 08283T0
RO3G 08158T0
Bade OZJ06498.1
Rirr XP 025171716.1
Ltra XP 021884399.1
100
17
2
4
0
69
1
83
73
57
58
92
99
24
100
7
7
99
3
3
100
6
7
97
42
100
26
8
39
91
95
99
100
96
95
87
100
36
100
95
88
80
51
70
79
0.05

### RPS10

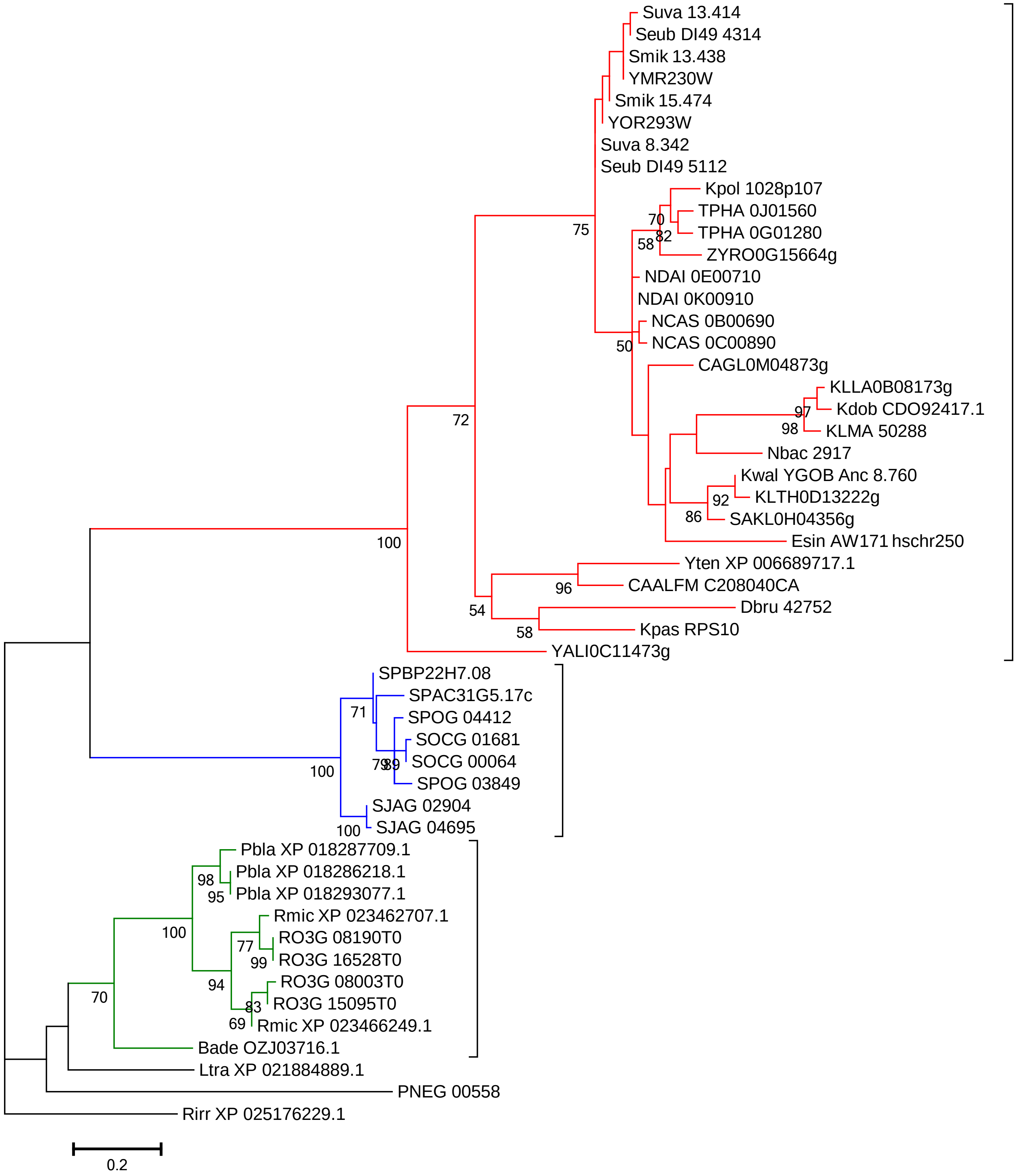

Suva 13.414
Seub DI49 4314
Smik 13.438
YMR230W
Smik 15.474
YOR293W
Suva 8.342
Seub DI49 5112
Kpol 1028p107
TPHA 0J01560
TPHA 0G01280
ZYRO0G15664g
NDAI 0E00710
NDAI 0K00910
NCAS 0B00690
NCAS 0C00890
CAGL0M04873g
KLLA0B08173g
Kdob CDO92417.1
KLMA 50288
Nbac 2917
Kwal YGOB Anc 8.760
KLTH0D13222g
SAKL0H04356g
Esin AW171 hschr250
Yten XP 006689717.1
CAALFM C208040CA
Dbru 42752
Kpas RPS10
YALI0C11473g
SPBP22H7.08
SPAC31G5.17c
SPOG 04412
SOCG 01681
SOCG 00064
SPOG 03849
SJAG 02904
SJAG 04695
Pbla XP 018287709.1
Pbla XP 018286218.1
Pbla XP 018293077.1
Rmic XP 023462707.1
RO3G 08190T0
RO3G 16528T0
RO3G 08003T0
RO3G 15095T0
Rmic XP 023466249.1
Bade OZJ03716.1
Ltra XP 021884889.1
PNEG 00558
Rirr XP 025176229.1
75
70
82
58
50
97
98
72
92
86
100
96
54
58
71
79
89
100
100
98
95
100
77
99
94
83
69
70
0.2

### RPS11

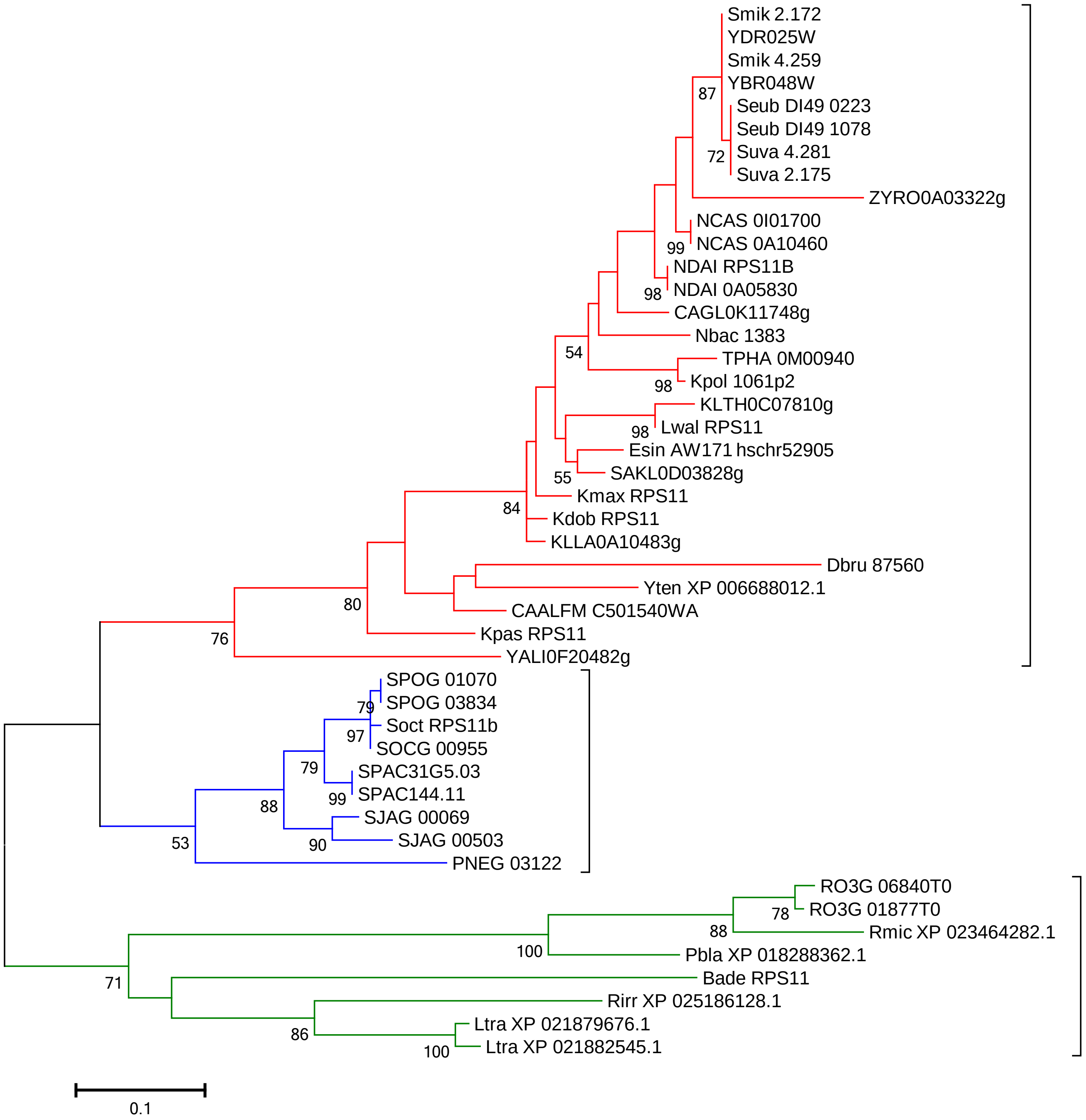

Smik 2.172
YDR025W
Smik 4.259
YBR048W
Seub DI49 0223
Seub DI49 1078
Suva 4.281
Suva 2.175
87
72
ZYRO0A03322g
NCAS 0I01700
NCAS 0A10460
99
NDAI RPS11B
NDAI 0A05830
98
CAGL0K11748g
Nbac 1383
54
TPHA 0M00940
Kpol 1061p2
98
KLTH0C07810g
Lwal RPS11
98
Esin AW171 hschr52905
SAKL0D03828g
55
Kmax RPS11
84
Kdob RPS11
KLLA0A10483g
Dbru 87560
Yten XP 006688012.1
80
CAALFM C501540WA
Kpas RPS11
76
YALI0F20482g
SPOG 01070
SPOG 03834
79
Soct RPS11b
97
SOCG 00955
79
SPAC31G5.03
SPAC144.11
99
88
SJAG 00069
SJAG 00503
90
53
PNEG 03122
RO3G 06840T0
RO3G 01877T0
78
88
Rmic XP 023464282.1
100
Pbla XP 018288362.1
71
Bade RPS11
Rirr XP 025186128.1
86
Ltra XP 021879676.1
Ltra XP 021882545.1
100
0.1

RPS12

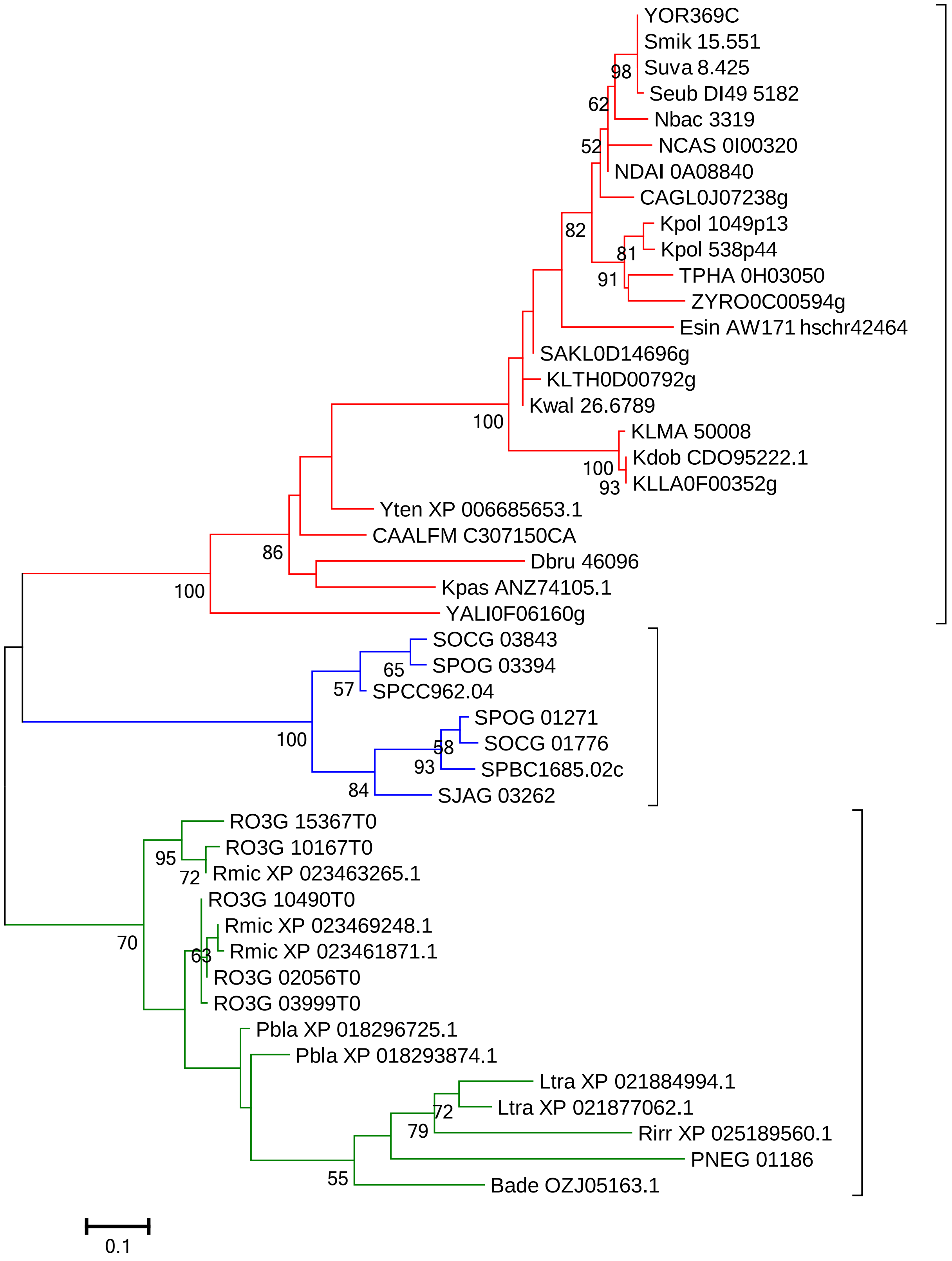
YOR369C
Smik 15.551
Suva 8.425
Seub DI49 5182
Nbac 3319
NCAS 0I00320
NDAI 0A08840
CAGL0J07238g
Kpol 1049p13
Kpol 538p44
TPHA 0H03050
ZYRO0C00594g
Esin AW171 hschr42464
SAKL0D14696g
KLTH0D00792g
Kwal 26.6789
KLMA 50008
Kdob CDO95222.1
KLLA0F00352g
Yten XP 006685653.1
CAALFM C307150CA
Dbru 46096
Kpas ANZ74105.1
YALI0F06160g
SOCG 03843
SPOG 03394
SPCC962.04
SPOG 01271
SOCG 01776
SPBC1685.02c
SJAG 03262
RO3G 15367T0
RO3G 10167T0
Rmic XP 023463265.1
RO3G 10490T0
Rmic XP 023469248.1
Rmic XP 023461871.1
RO3G 02056T0
RO3G 03999T0
Pbla XP 018296725.1
Pbla XP 018293874.1
Ltra XP 021884994.1
Ltra XP 021877062.1
Rirr XP 025189560.1
PNEG 01186
Bade OZJ05163.1
98
62
52
82
81
91
100
100
93
86
100
65
57
100
58
93
84
95
72
70
63
72
79
55
0.1

#### RPS13

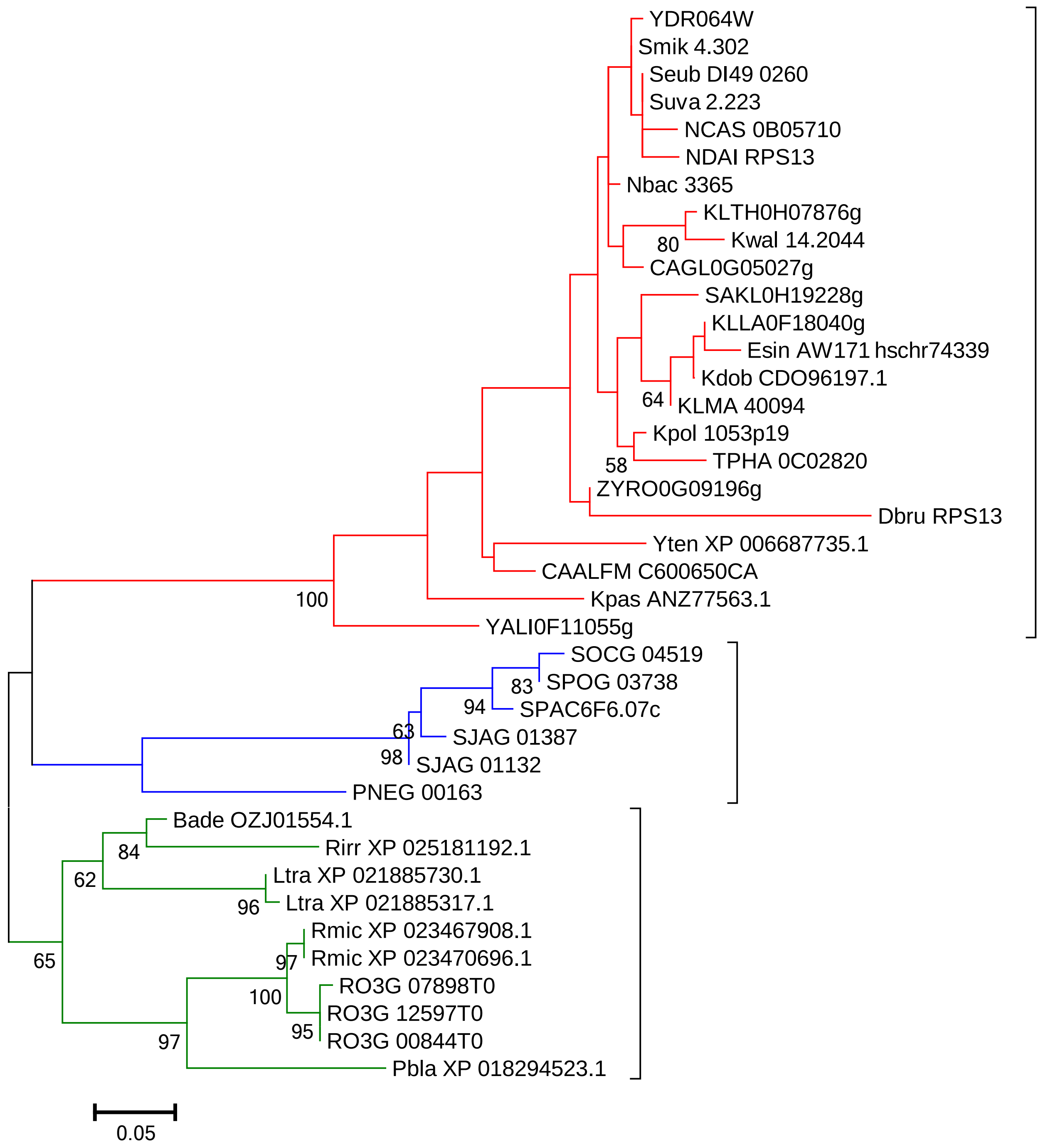
YDR064W
Smik 4.302
Seub DI49 0260
Suva 2.223
NCAS 0B05710
NDAI RPS13
Nbac 3365
KLTH0H07876g
Kwal 14.2044
80
CAGL0G05027g
SAKL0H19228g
KLLA0F18040g
Esin AW171 hschr74339
Kdob CDO96197.1
64
KLMA 40094
Kpol 1053p19
58
TPHA 0C02820
ZYRO0G09196g
Dbru RPS13
Yten XP 006687735.1
CAALFM C600650CA
Kpas ANZ77563.1
100
YALI0F11055g
SOCG 04519
83
SPOG 03738
94
SPAC6F6.07c
63
SJAG 01387
98
SJAG 01132
PNEG 00163
Bade OZJ01554.1
84
Rirr XP 025181192.1
62
Ltra XP 021885730.1
96
Ltra XP 021885317.1
Rmic XP 023467908.1
97
Rmic XP 023470696.1
65
100
RO3G 07898T0
RO3G 12597T0
95
RO3G 00844T0
97
Pbla XP 018294523.1
0.05

### RPS14

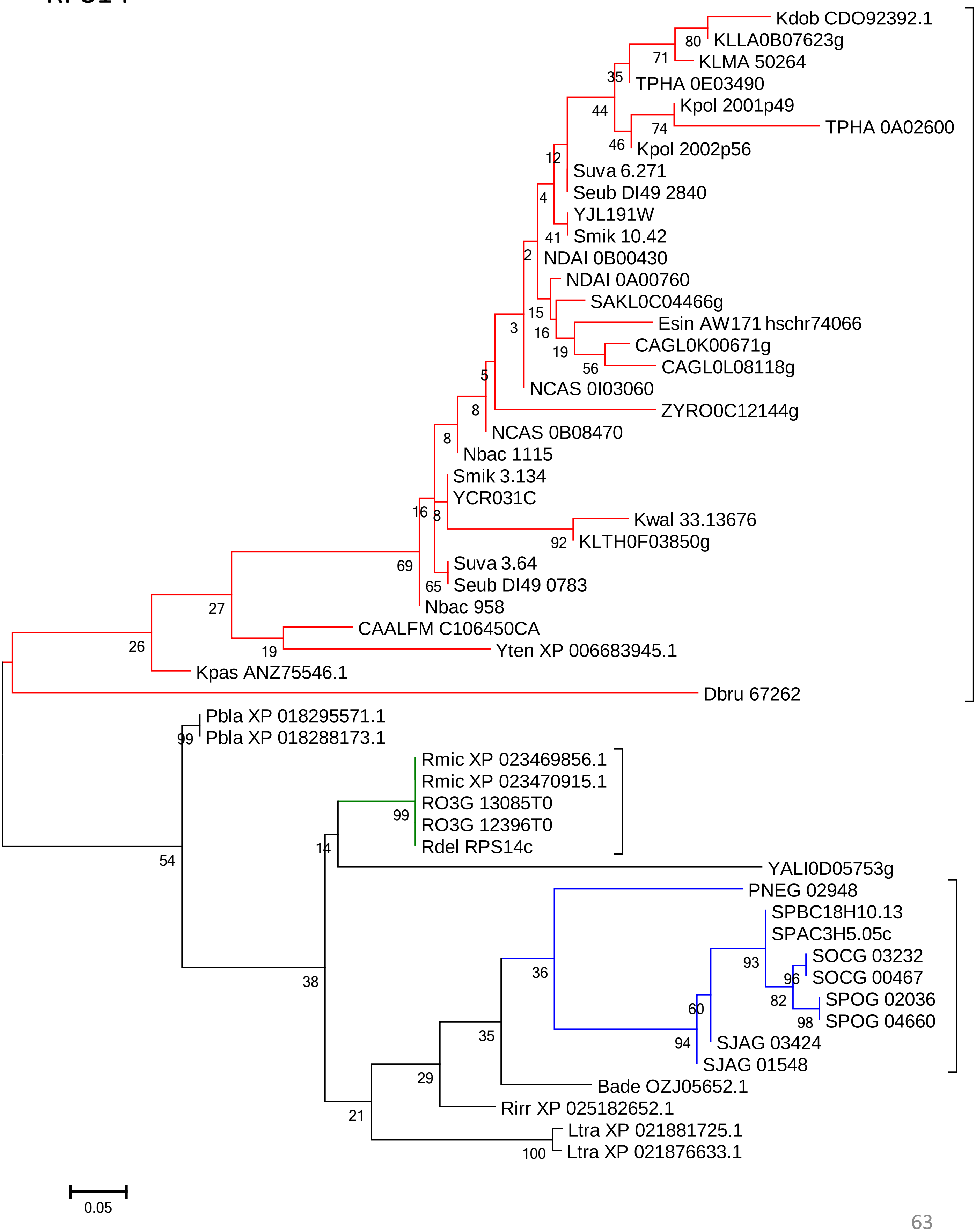
Kdob CDO92392.1
KLLA0B07623g
KLMA 50264
TPHA 0E03490
Kpol 2001p49
TPHA 0A02600
Kpol 2002p56
Suva 6.271
Seub DI49 2840
YJL191W
Smik 10.42
NDAI 0B00430
NDAI 0A00760
SAKL0C04466g
Esin AW171 hschr74066
CAGL0K00671g
CAGL0L08118g
NCAS 0I03060
ZYRO0C12144g
NCAS 0B08470
Nbac 1115
Smik 3.134
YCR031C
Kwal 33.13676
KLTH0F03850g
Suva 3.64
Seub DI49 0783
Nbac 958
CAALFM C106450CA
Yten XP 006683945.1
Kpas ANZ75546.1
Dbru 67262
Pbla XP 018295571.1
Pbla XP 018288173.1
Rmic XP 023469856.1
Rmic XP 023470915.1
RO3G 13085T0
RO3G 12396T0
Rdel RPS14c
YALI0D05753g
PNEG 02948
SPBC18H10.13
SPAC3H5.05c
SOCG 03232
SOCG 00467
SPOG 02036
SPOG 04660
SJAG 03424
SJAG 01548
Bade OZJ05652.1
Rirr XP 025182652.1
Ltra XP 021881725.1
Ltra XP 021876633.1
0.05

### RPS15

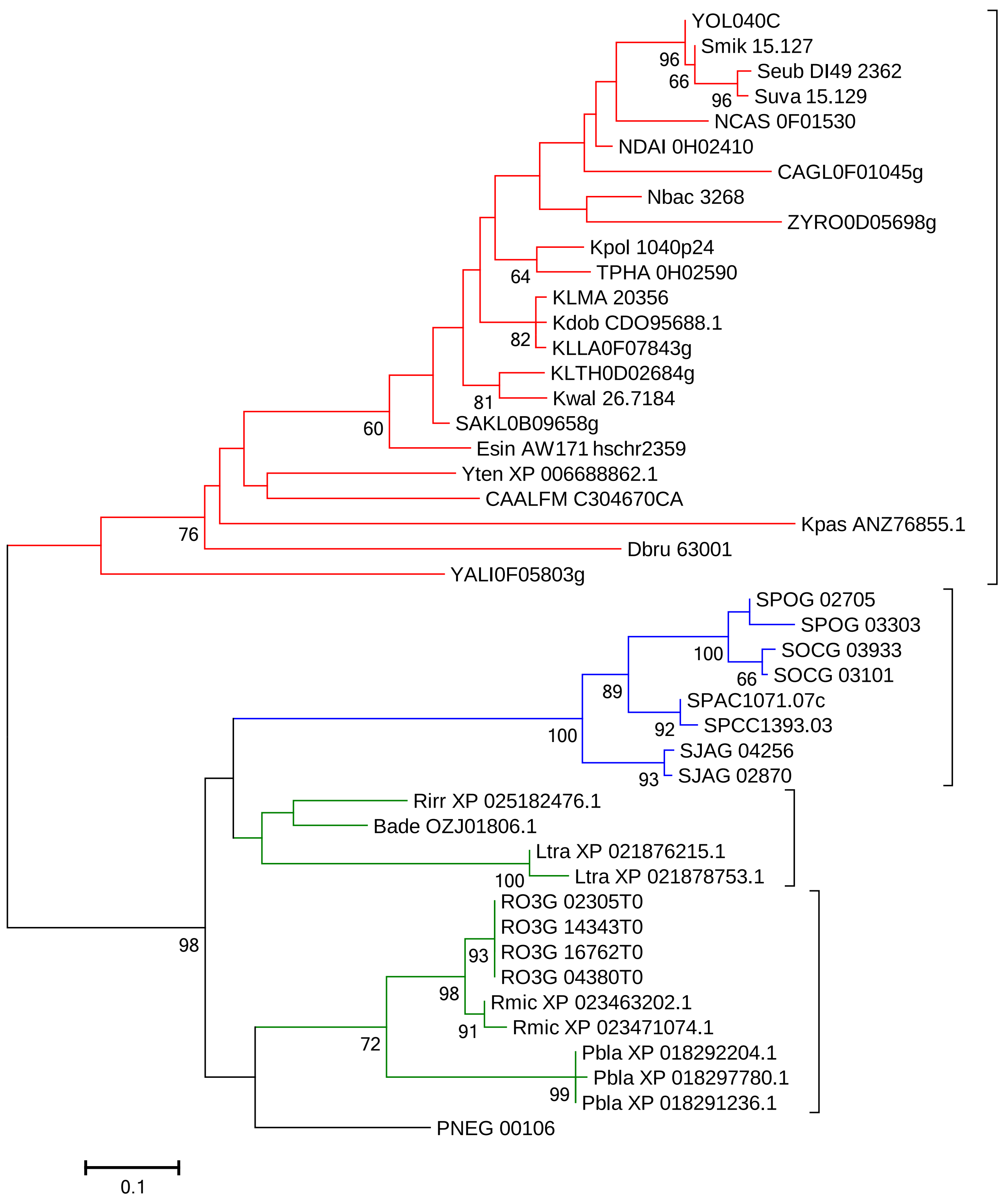
YOL040C
Smik 15.127
Seub DI49 2362
Suva 15.129
NCAS 0F01530
NDAI 0H02410
CAGL0F01045g
Nbac 3268
ZYRO0D05698g
Kpol 1040p24
TPHA 0H02590
KLMA 20356
Kdob CDO95688.1
KLLA0F07843g
KLTH0D02684g
Kwal 26.7184
SAKL0B09658g
Esin AW171 hschr2359
Yten XP 006688862.1
CAALFM C304670CA
Kpas ANZ76855.1
Dbru 63001
YALI0F05803g
SPOG 02705
SPOG 03303
SOCG 03933
SOCG 03101
SPAC1071.07c
SPCC1393.03
SJAG 04256
SJAG 02870
Rirr XP 025182476.1
Bade OZJ01806.1
Ltra XP 021876215.1
Ltra XP 021878753.1
RO3G 02305T0
RO3G 14343T0
RO3G 16762T0
RO3G 04380T0
Rmic XP 023463202.1
Rmic XP 023471074.1
Pbla XP 018292204.1
Pbla XP 018297780.1
Pbla XP 018291236.1
PNEG 00106
96
66
96
64
82
81
60
76
100
66
89
92
100
93
100
93
98
91
72
99
98
0.1

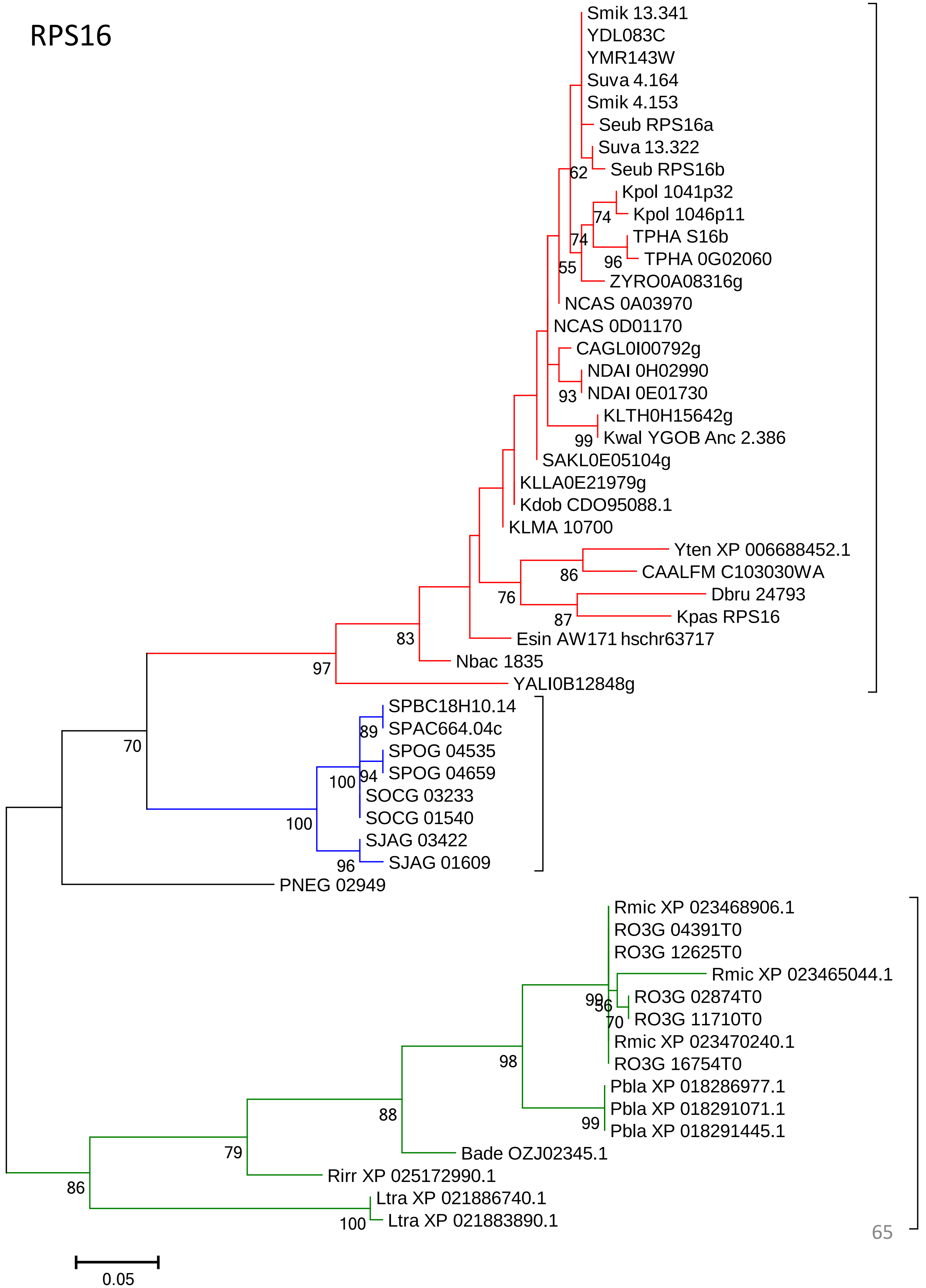
RPS16
Smik 13.341
YDL083C
YMR143W
Suva 4.164
Smik 4.153
Seub RPS16a
Suva 13.322
Seub RPS16b
Kpol 1041p32
Kpol 1046p11
TPHA S16b
TPHA 0G02060
ZYRO0A08316g
NCAS 0A03970
NCAS 0D01170
CAGL0I00792g
NDAI 0H02990
NDAI 0E01730
KLTH0H15642g
Kwal YGOB Anc 2.386
SAKL0E05104g
KLLA0E21979g
Kdob CDO95088.1
KLMA 10700
Yten XP 006688452.1
CAALFM C103030WA
Dbru 24793
Kpas RPS16
Esin AW171 hschr63717
Nbac 1835
YALI0B12848g
SPBC18H10.14
SPAC664.04c
SPOG 04535
SPOG 04659
SOCG 03233
SOCG 01540
SJAG 03422
SJAG 01609
PNEG 02949
Rmic XP 023468906.1
RO3G 04391T0
RO3G 12625T0
Rmic XP 023465044.1
RO3G 02874T0
RO3G 11710T0
Rmic XP 023470240.1
RO3G 16754T0
Pbla XP 018286977.1
Pbla XP 018291071.1
Pbla XP 018291445.1
Bade OZJ02345.1
Rirr XP 025172990.1
Ltra XP 021886740.1
Ltra XP 021883890.1
0.05

RPS17

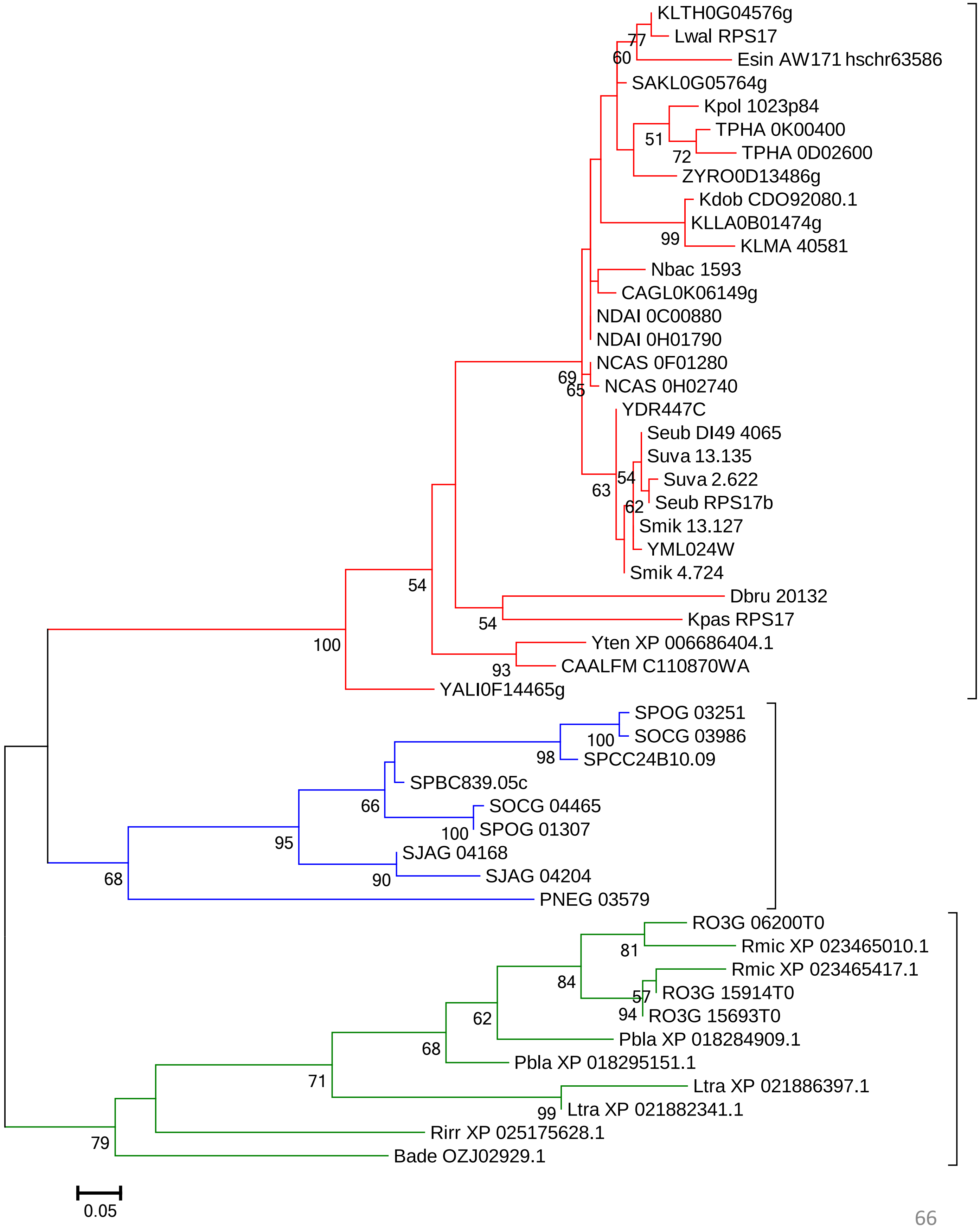
KLTH0G04576g
Lwal RPS17
Esin AW171 hschr63586
SAKL0G05764g
Kpol 1023p84
TPHA 0K00400
TPHA 0D02600
ZYRO0D13486g
Kdob CDO92080.1
KLLA0B01474g
KLMA 40581
Nbac 1593
CAGL0K06149g
NDAI 0C00880
NDAI 0H01790
NCAS 0F01280
NCAS 0H02740
YDR447C
Seub DI49 4065
Suva 13.135
Suva 2.622
Seub RPS17b
Smik 13.127
YML024W
Smik 4.724
Dbru 20132
Kpas RPS17
Yten XP 006686404.1
CAALFM C110870WA
YALI0F14465g
SPOG 03251
SOCG 03986
SPCC24B10.09
SPBC839.05c
SOCG 04465
SPOG 01307
SJAG 04168
SJAG 04204
PNEG 03579
RO3G 06200T0
Rmic XP 023465010.1
Rmic XP 023465417.1
RO3G 15914T0
RO3G 15693T0
Pbla XP 018284909.1
Pbla XP 018295151.1
Ltra XP 021886397.1
Ltra XP 021882341.1
Rirr XP 025175628.1
Bade OZJ02929.1
0.05

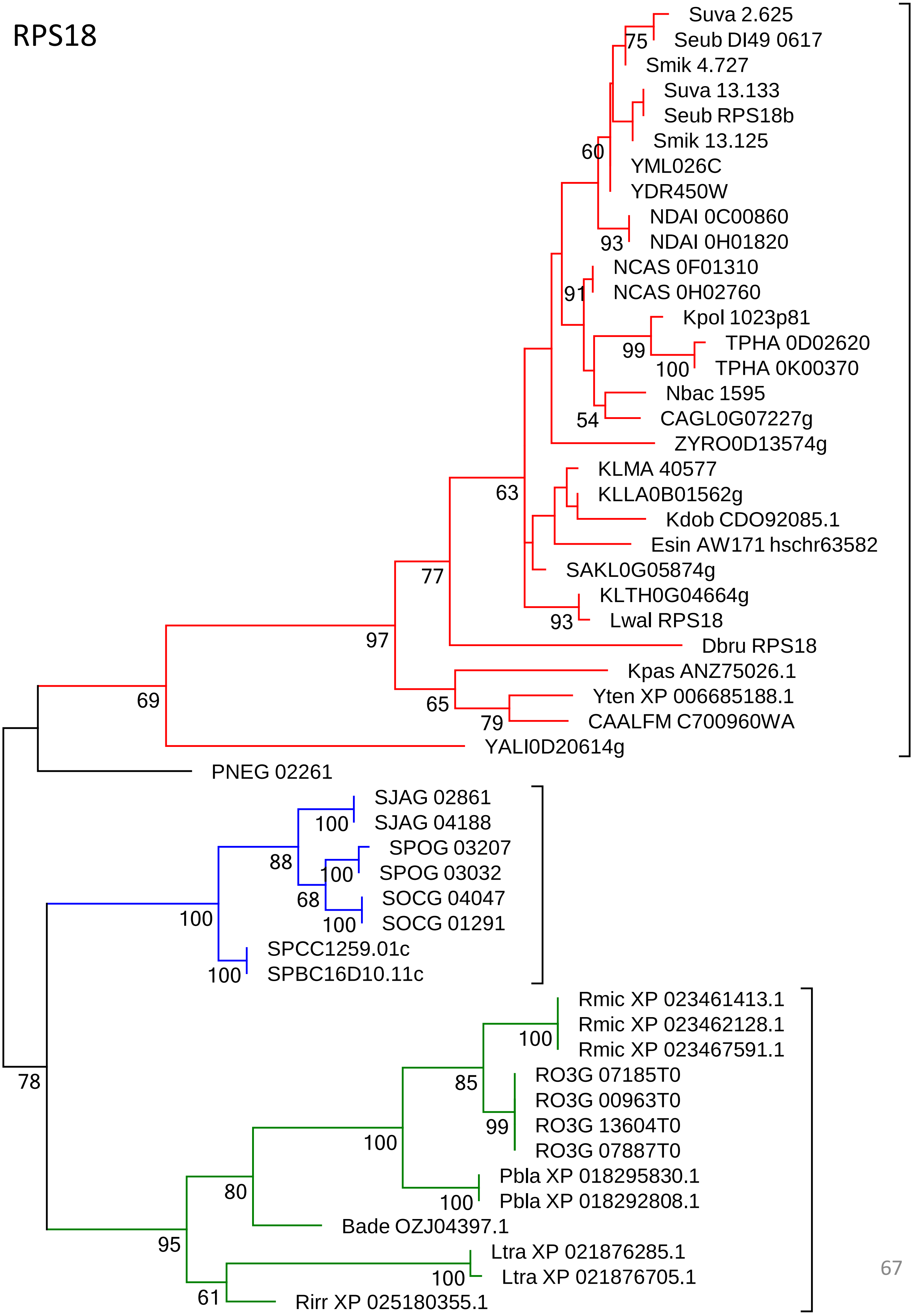
RPS18
Suva 2.625
Seub DI49 0617
75
Smik 4.727
Suva 13.133
Seub RPS18b
Smik 13.125
60
YML026C
YDR450W
NDAI 0C00860
93
NDAI 0H01820
NCAS 0F01310
91
NCAS 0H02760
Kpol 1023p81
99
TPHA 0D02620
100
TPHA 0K00370
Nbac 1595
54
CAGL0G07227g
ZYRO0D13574g
KLMA 40577
63
KLLA0B01562g
Kdob CDO92085.1
Esin AW171 hschr63582
77
SAKL0G05874g
KLTH0G04664g
93
Lwal RPS18
97
Dbru RPS18
Kpas ANZ75026.1
65
Yten XP 006685188.1
69
79
CAALFM C700960WA
YALI0D20614g
PNEG 02261
SJAG 02861
100
SJAG 04188
SPOG 03207
88
100
SPOG 03032
68
SOCG 04047
100
100
SOCG 01291
SPCC1259.01c
100
SPBC16D10.11c
Rmic XP 023461413.1
Rmic XP 023462128.1
100
Rmic XP 023467591.1
RO3G 07185T0
85
RO3G 00963T0
78
99
RO3G 13604T0
100
RO3G 07887T0
Pbla XP 018295830.1
80
100
Pbla XP 018292808.1
Bade OZJ04397.1
95
Ltra XP 021876285.1
100
Ltra XP 021876705.1
61
Rirr XP 025180355.1

### RPS19

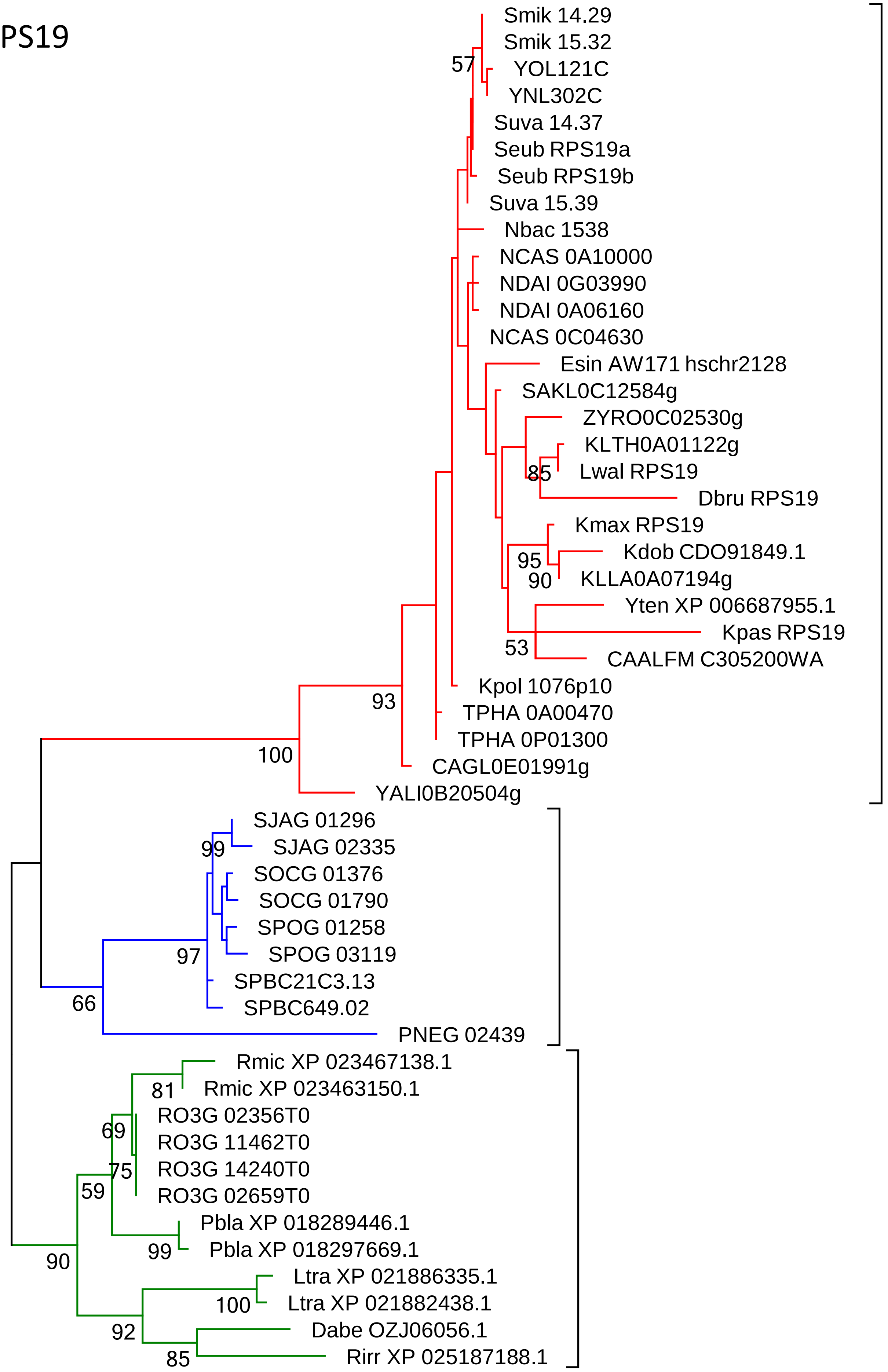
Smik 14.29
Smik 15.32
57
YOL121C
YNL302C
Suva 14.37
Seub RPS19a
Seub RPS19b
Suva 15.39
Nbac 1538
NCAS 0A10000
NDAI 0G03990
NDAI 0A06160
NCAS 0C04630
Esin AW171 hschr2128
SAKL0C12584g
ZYRO0C02530g
KLTH0A01122g
85
Lwal RPS19
Dbru RPS19
Kmax RPS19
Kdob CDO91849.1
95
90
KLLA0A07194g
Yten XP 006687955.1
Kpas RPS19
53
CAALFM C305200WA
Kpol 1076p10
93
TPHA 0A00470
TPHA 0P01300
100
CAGL0E01991g
YALI0B20504g
SJAG 01296
SJAG 02335
99
SOCG 01376
SOCG 01790
SPOG 01258
97
SPOG 03119
SPBC21C3.13
66
SPBC649.02
PNEG 02439
Rmic XP 023467138.1
81
Rmic XP 023463150.1
RO3G 02356T0
69
RO3G 11462T0
75
RO3G 14240T0
59
RO3G 02659T0
Pbla XP 018289446.1
90
99
Pbla XP 018297669.1
Ltra XP 021886335.1
100
Ltra XP 021882438.1
92
Dabe OZJ06056.1
85
Rirr XP 025187188.1

### RPS20

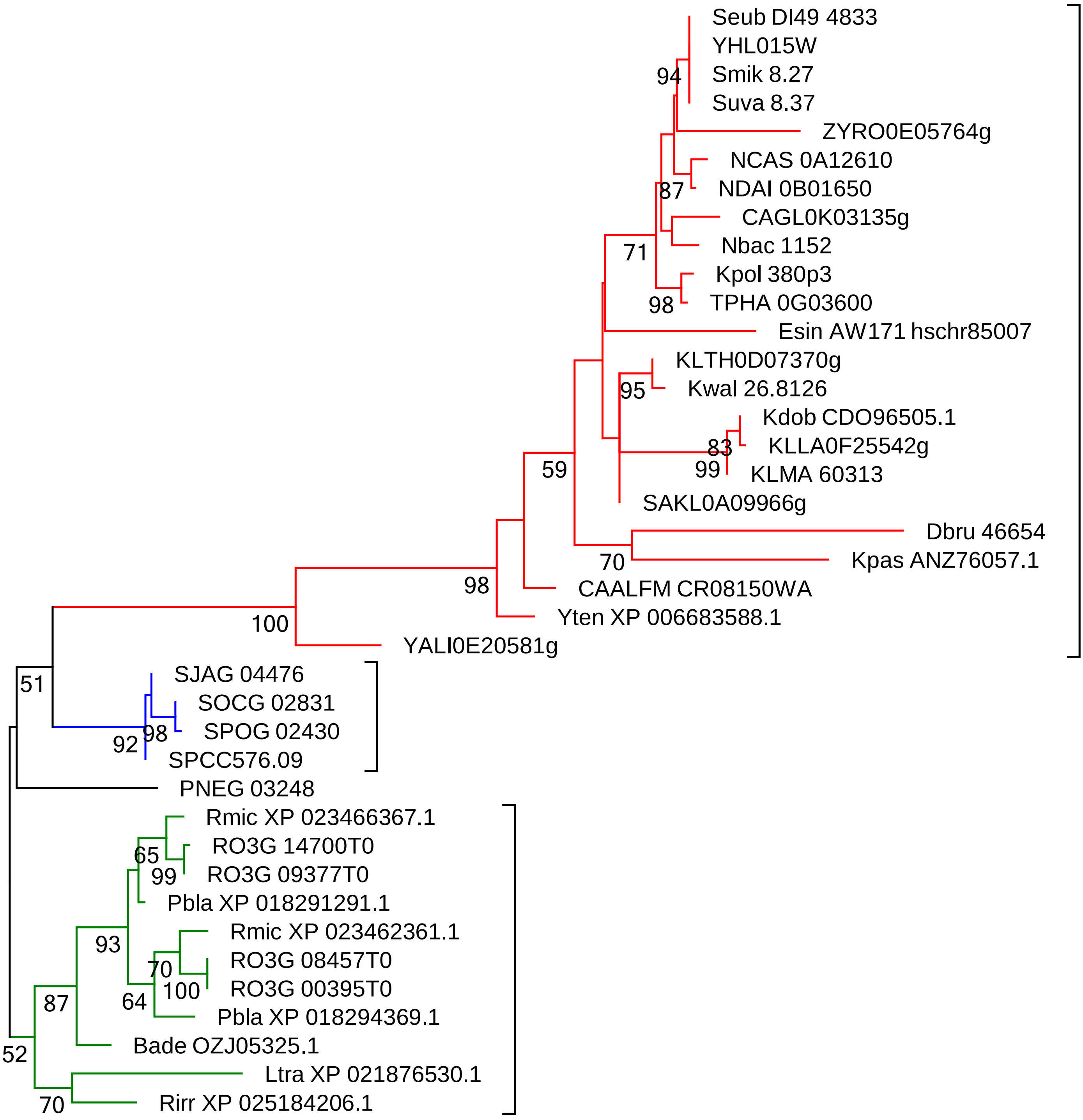

Seub DI49 4833
YHL015W
Smik 8.27
Suva 8.37
94
ZYRO0E05764g
NCAS 0A12610
NDAI 0B01650
87
CAGL0K03135g
Nbac 1152
71
Kpol 380p3
TPHA 0G03600
98
Esin AW171 hschr85007
KLTH0D07370g
Kwal 26.8126
95
Kdob CDO96505.1
KLLA0F25542g
83
99
KLMA 60313
SAKL0A09966g
59
Dbru 46654
Kpas ANZ76057.1
70
CAALFM CR08150WA
98
Yten XP 006683588.1
100
YALI0E20581g
SJAG 04476
SOCG 02831
SPOG 02430
98
92
SPCC576.09
51
PNEG 03248
Rmic XP 023466367.1
RO3G 14700T0
65
99
RO3G 09377T0
Pbla XP 018291291.1
Rmic XP 023462361.1
93
RO3G 08457T0
70
RO3G 00395T0
100
64
Pbla XP 018294369.1
87
Bade OZJ05325.1
52
Ltra XP 021876530.1
70
Rirr XP 025184206.1

### RPS21

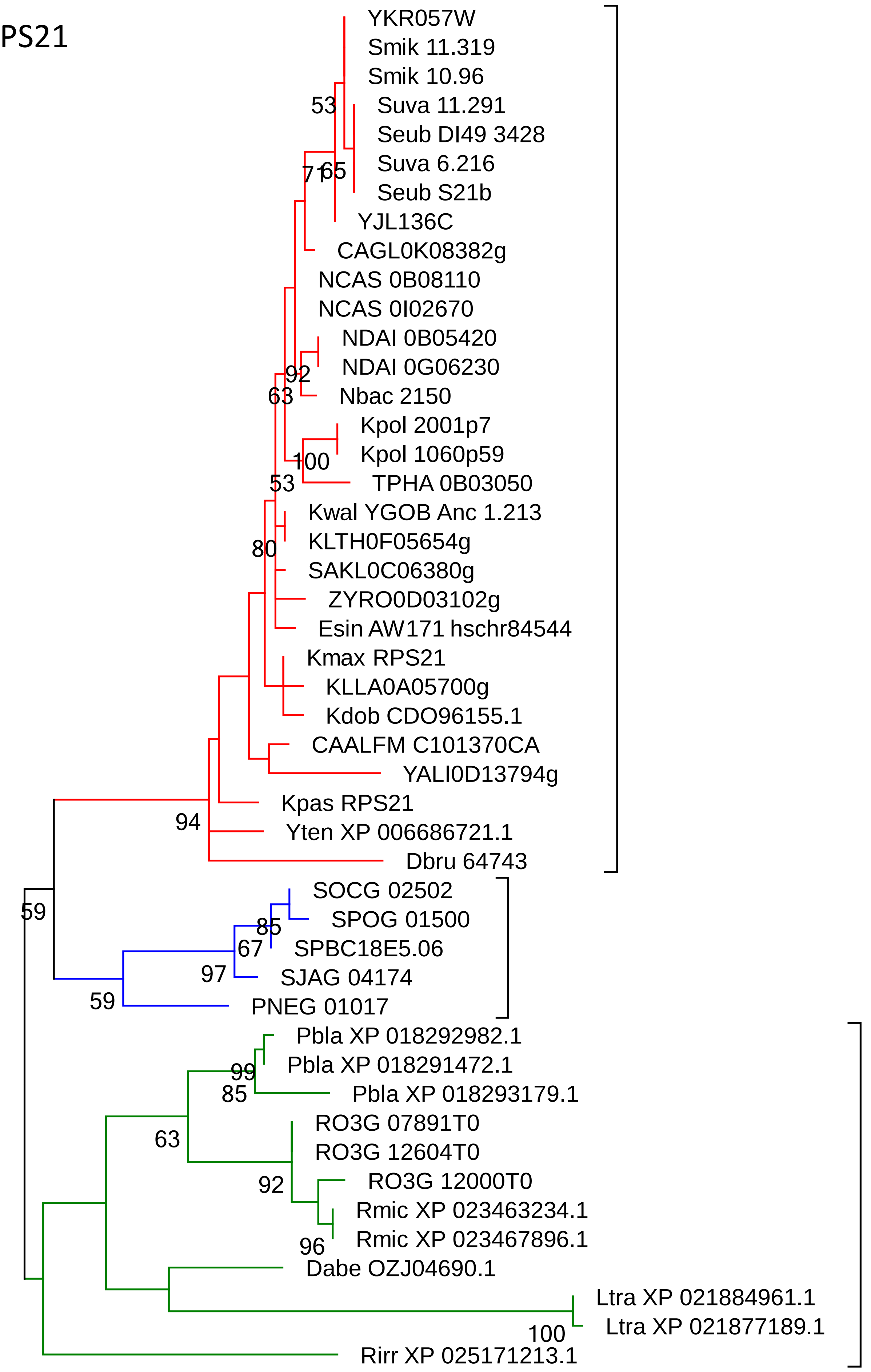
YKR057W
Smik 11.319
Smik 10.96
Suva 11.291
Seub DI49 3428
Suva 6.216
Seub S21b
YJL136C
CAGL0K08382g
NCAS 0B08110
NCAS 0I02670
NDAI 0B05420
NDAI 0G06230
Nbac 2150
Kpol 2001p7
Kpol 1060p59
TPHA 0B03050
Kwal YGOB Anc 1.213
KLTH0F05654g
SAKL0C06380g
ZYRO0D03102g
Esin AW171 hschr84544
Kmax RPS21
KLLA0A05700g
Kdob CDO96155.1
CAALFM C101370CA
YALI0D13794g
Kpas RPS21
Yten XP 006686721.1
Dbru 64743
SOCG 02502
SPOG 01500
SPBC18E5.06
SJAG 04174
PNEG 01017
Pbla XP 018292982.1
Pbla XP 018291472.1
Pbla XP 018293179.1
RO3G 07891T0
RO3G 12604T0
RO3G 12000T0
Rmic XP 023463234.1
Rmic XP 023467896.1
Dabe OZJ04690.1
Ltra XP 021884961.1
Ltra XP 021877189.1
Rirr XP 025171213.1
53
71
65
92
63
100
53
80
94
59
85
67
97
59
99
85
63
92
96
100

### RPS22 (RPS15A)

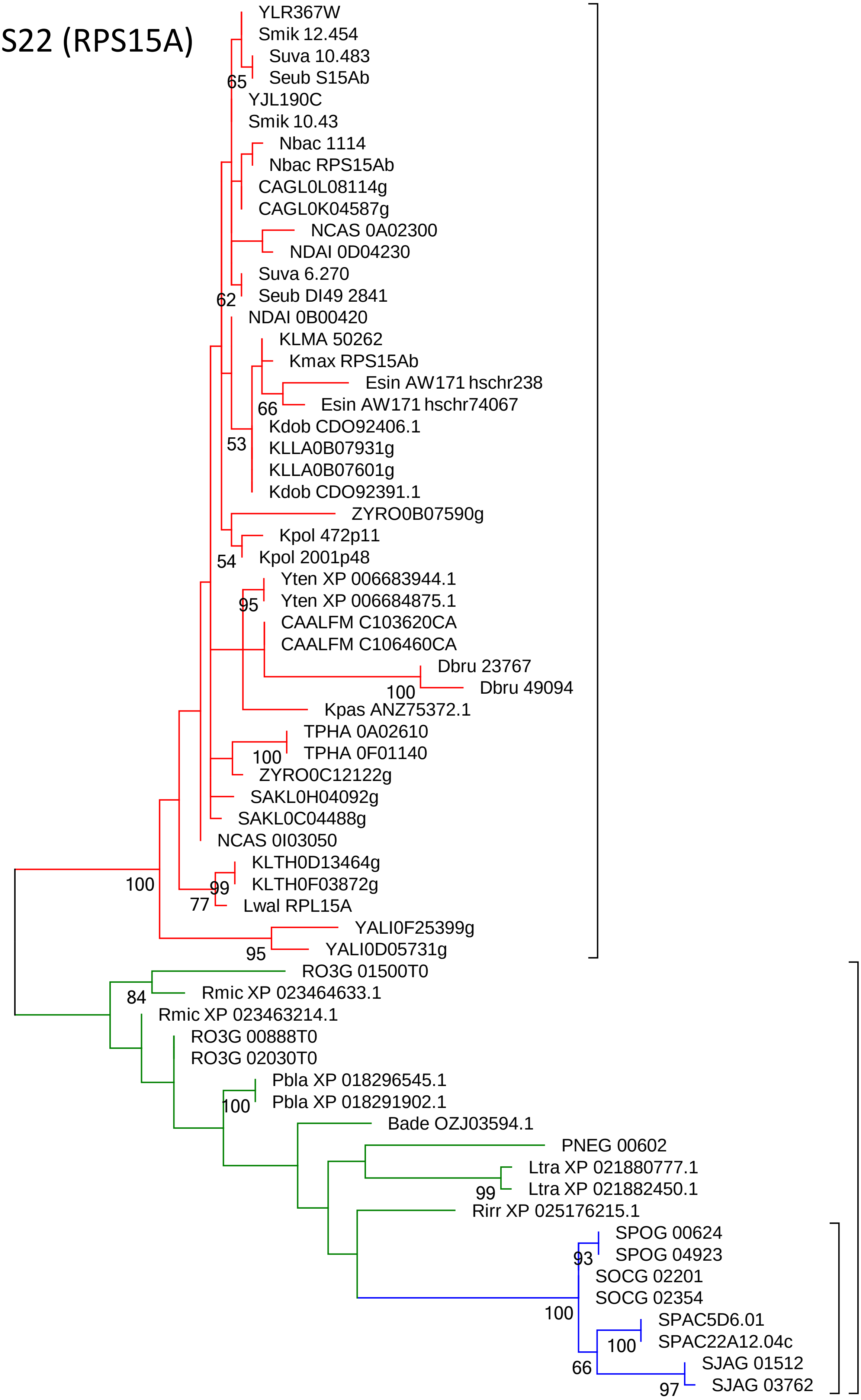

### RPS23

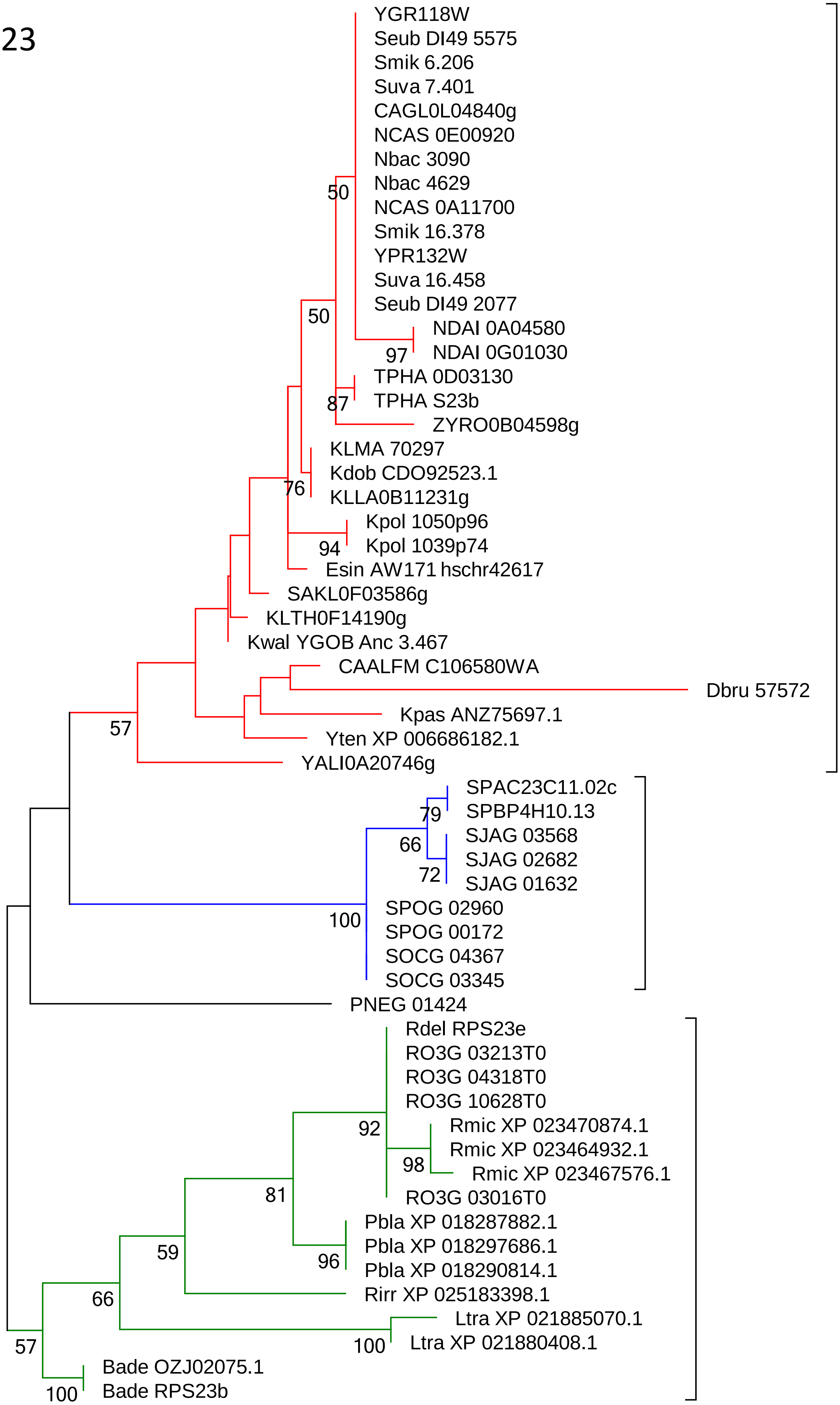
YGR118W
Seub DI49 5575
Smik 6.206
Suva 7.401
CAGL0L04840g
NCAS 0E00920
Nbac 3090
Nbac 4629
NCAS 0A11700
Smik 16.378
YPR132W
Suva 16.458
Seub DI49 2077
NDAI 0A04580
NDAI 0G01030
TPHA 0D03130
TPHA S23b
ZYRO0B04598g
KLMA 70297
Kdob CDO92523.1
KLLA0B11231g
Kpol 1050p96
Kpol 1039p74
Esin AW171 hschr42617
SAKL0F03586g
KLTH0F14190g
Kwal YGOB Anc 3.467
CAALFM C106580WA
Dbru 57572
Kpas ANZ75697.1
Yten XP 006686182.1
YALI0A20746g
SPAC23C11.02c
SPBP4H10.13
SJAG 03568
SJAG 02682
SJAG 01632
SPOG 02960
SPOG 00172
SOCG 04367
SOCG 03345
PNEG 01424
Rdel RPS23e
RO3G 03213T0
RO3G 04318T0
RO3G 10628T0
Rmic XP 023470874.1
Rmic XP 023464932.1
Rmic XP 023467576.1
RO3G 03016T0
Pbla XP 018287882.1
Pbla XP 018297686.1
Pbla XP 018290814.1
Rirr XP 025183398.1
Ltra XP 021885070.1
Ltra XP 021880408.1
Bade OZJ02075.1
Bade RPS23b
50
50
97
87
76
94
57
79
66
72
100
92
98
81
96
59
66
100
57
100

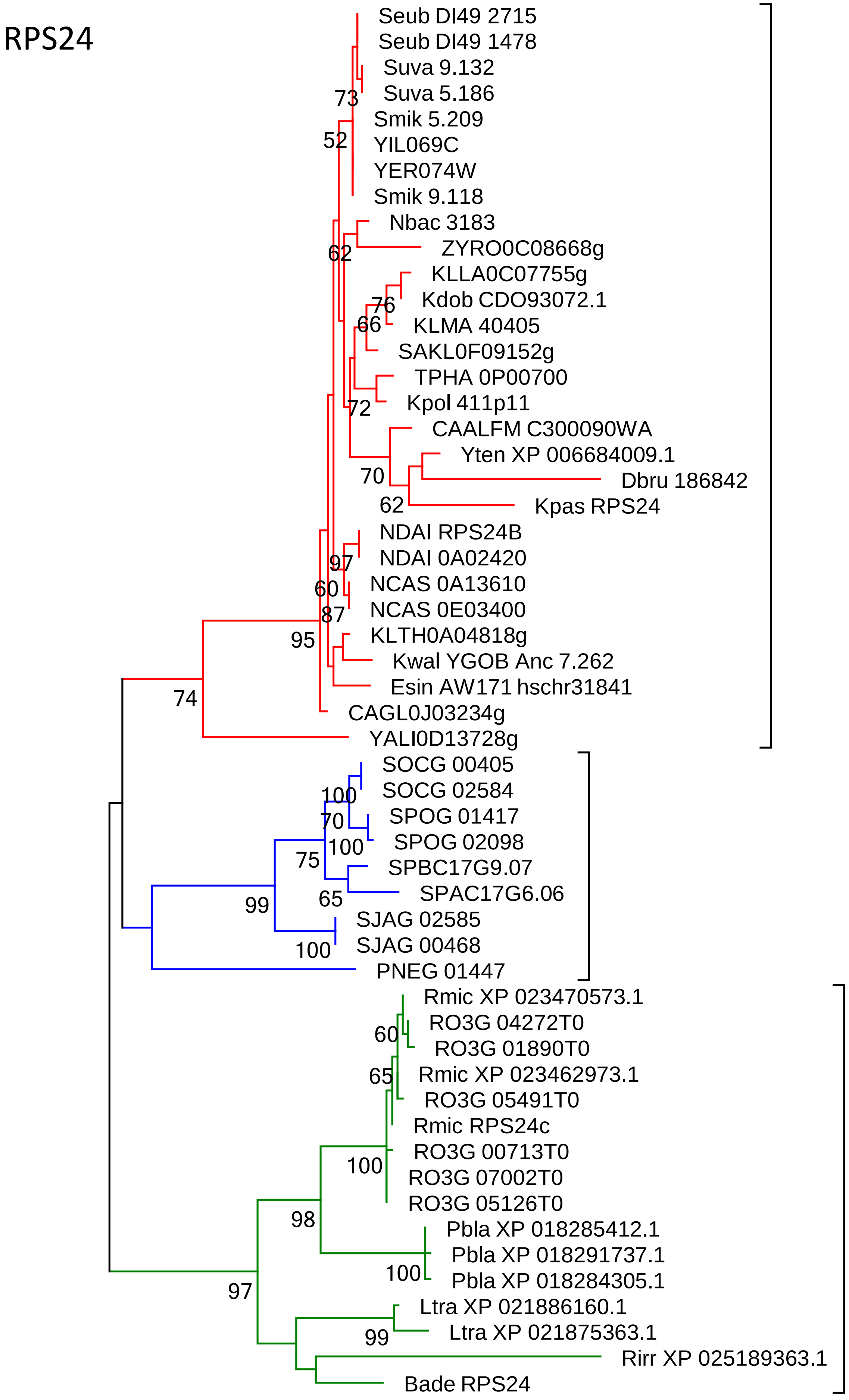
RPS24
Seub DI49 2715
Seub DI49 1478
Suva 9.132
Suva 5.186
73
Smik 5.209
52
YIL069C
YER074W
Smik 9.118
Nbac 3183
ZYRO0C08668g
62
KLLA0C07755g
Kdob CDO93072.1
76
66
KLMA 40405
SAKL0F09152g
TPHA 0P00700
Kpol 411p11
72
CAALFM C300090WA
Yten XP 006684009.1
70
Dbru 186842
62
Kpas RPS24
NDAI RPS24B
97
NDAI 0A02420
60
NCAS 0A13610
NCAS 0E03400
87
KLTH0A04818g
95
Kwal YGOB Anc 7.262
Esin AW171 hschr31841
74
CAGL0J03234g
YALI0D13728g
SOCG 00405
SOCG 02584
100
SPOG 01417
70
SPOG 02098
100
75
SPBC17G9.07
SPAC17G6.06
65
99
SJAG 02585
SJAG 00468
100
PNEG 01447
Rmic XP 023470573.1
RO3G 04272T0
60
RO3G 01890T0
Rmic XP 023462973.1
65
RO3G 05491T0
Rmic RPS24c
RO3G 00713T0
100
RO3G 07002T0
RO3G 05126T0
98
Pbla XP 018285412.1
Pbla XP 018291737.1
100
Pbla XP 018284305.1
97
Ltra XP 021886160.1
Ltra XP 021875363.1
99
Rirr XP 025189363.1
Bade RPS24

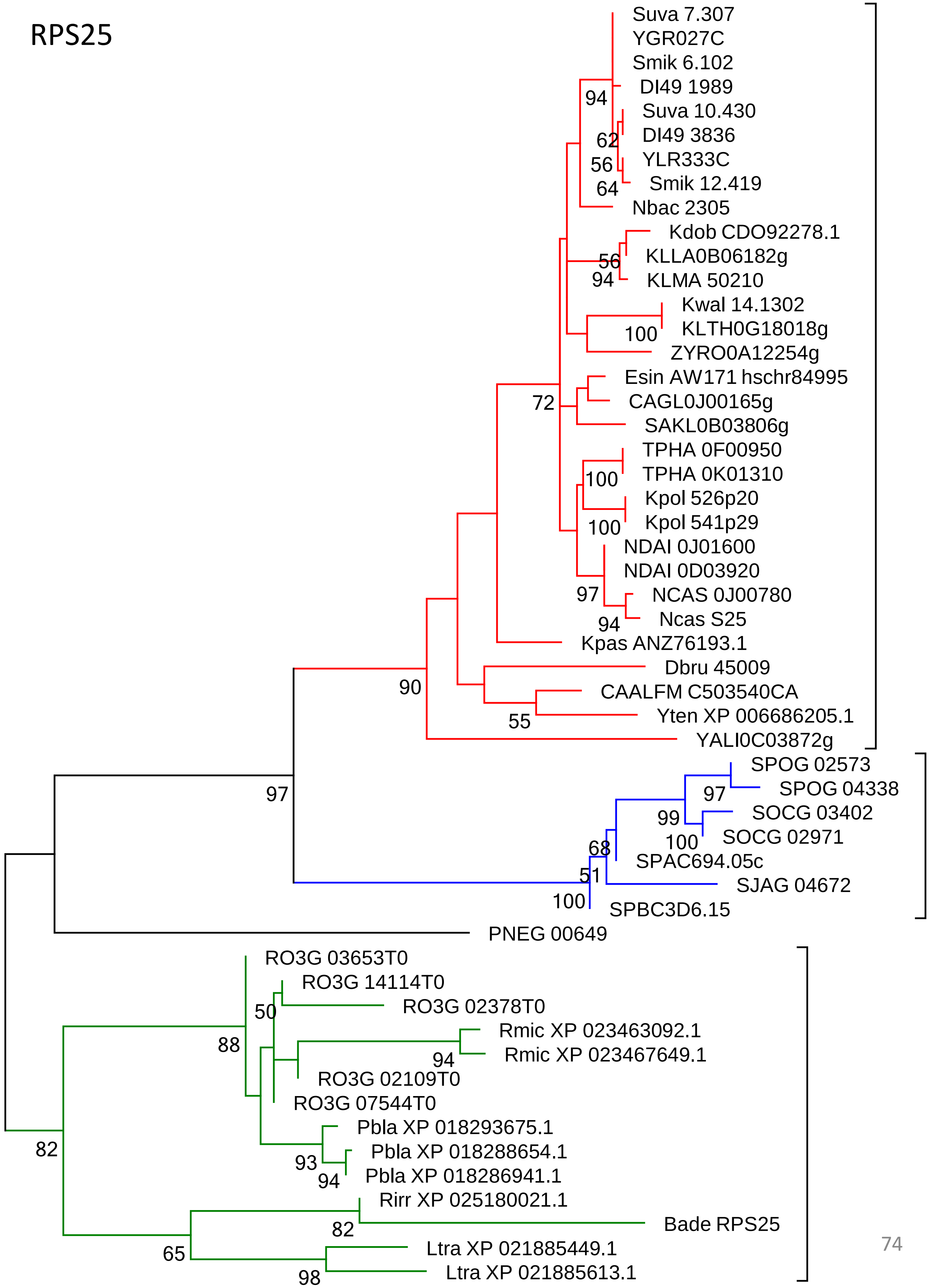
RPS25
Suva 7.307
YGR027C
Smik 6.102
DI49 1989
Suva 10.430
DI49 3836
YLR333C
Smik 12.419
Nbac 2305
Kdob CDO92278.1
KLLA0B06182g
KLMA 50210
Kwal 14.1302
KLTH0G18018g
ZYRO0A12254g
Esin AW171 hschr84995
CAGL0J00165g
SAKL0B03806g
TPHA 0F00950
TPHA 0K01310
Kpol 526p20
Kpol 541p29
NDAI 0J01600
NDAI 0D03920
NCAS 0J00780
Ncas S25
Kpas ANZ76193.1
Dbru 45009
CAALFM C503540CA
Yten XP 006686205.1
YALI0C03872g
SPOG 02573
SPOG 04338
SOCG 03402
SOCG 02971
SPAC694.05c
SJAG 04672
SPBC3D6.15
PNEG 00649
RO3G 03653T0
RO3G 14114T0
RO3G 02378T0
Rmic XP 023463092.1
Rmic XP 023467649.1
RO3G 02109T0
RO3G 07544T0
Pbla XP 018293675.1
Pbla XP 018288654.1
Pbla XP 018286941.1
Rirr XP 025180021.1
Bade RPS25
Ltra XP 021885449.1
Ltra XP 021885613.1
94
62
56
64
56
94
100
72
100
100
97
94
90
55
97
97
99
100
68
51
100
50
88
94
93
94
82
82
65
98

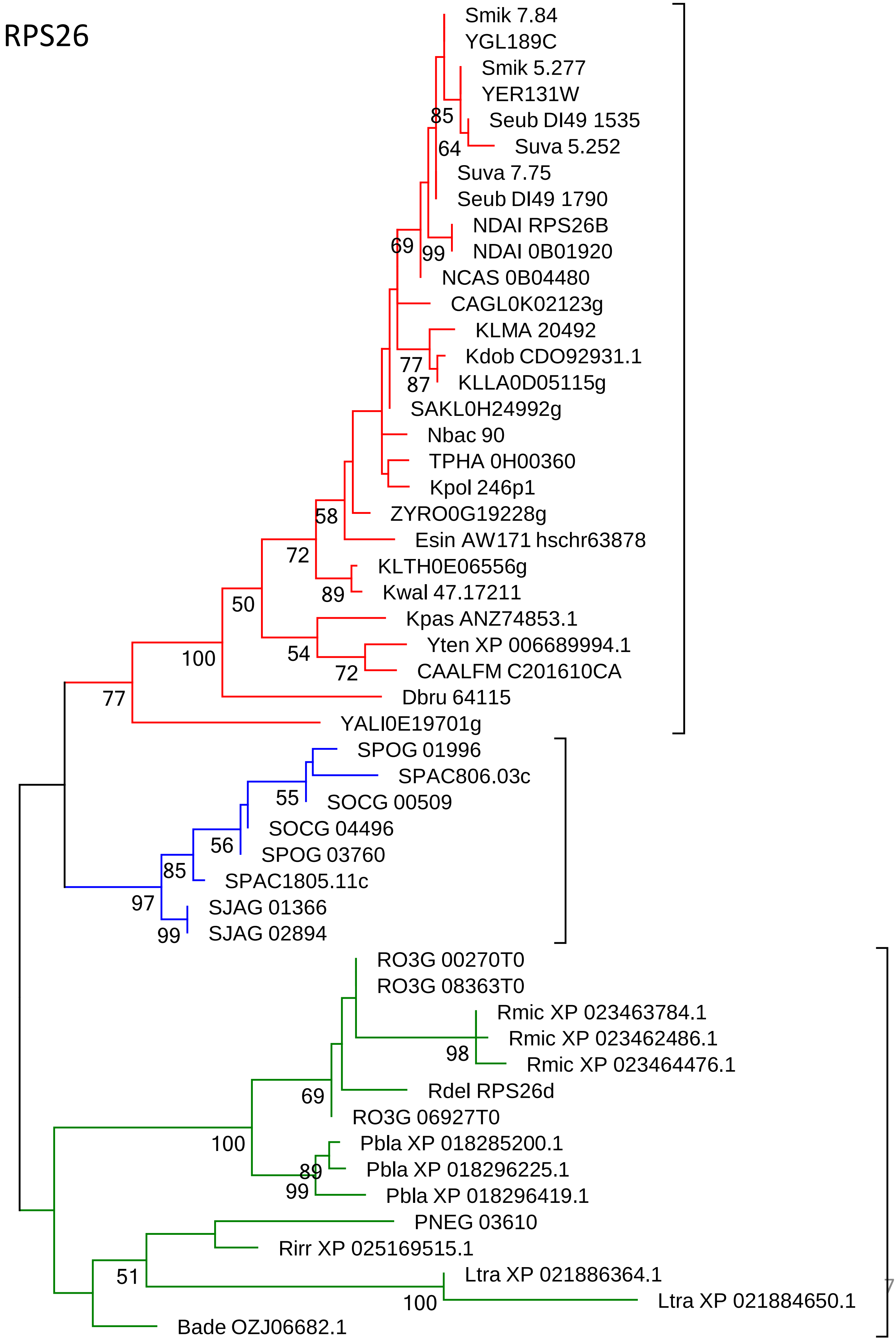
RPS26
Smik 7.84
YGL189C
Smik 5.277
YER131W
85
Seub DI49 1535
64
Suva 5.252
Suva 7.75
Seub DI49 1790
NDAI RPS26B
69
99
NDAI 0B01920
NCAS 0B04480
CAGL0K02123g
KLMA 20492
Kdob CDO92931.1
77
87
KLLA0D05115g
SAKL0H24992g
Nbac 90
TPHA 0H00360
Kpol 246p1
58
ZYRO0G19228g
Esin AW171 hschr63878
72
KLTH0E06556g
89
Kwal 47.17211
50
Kpas ANZ74853.1
Yten XP 006689994.1
100
54
72
CAALFM C201610CA
77
Dbru 64115
YALI0E19701g
SPOG 01996
SPAC806.03c
55
SOCG 00509
SOCG 04496
56
SPOG 03760
85
SPAC1805.11c
97
SJAG 01366
99
SJAG 02894
RO3G 00270T0
RO3G 08363T0
Rmic XP 023463784.1
Rmic XP 023462486.1
98
Rmic XP 023464476.1
Rdel RPS26d
69
RO3G 06927T0
100
Pbla XP 018285200.1
89
Pbla XP 018296225.1
99
Pbla XP 018296419.1
PNEG 03610
Rirr XP 025169515.1
51
Ltra XP 021886364.1
100
Ltra XP 021884650.1
Bade OZJ06682.1

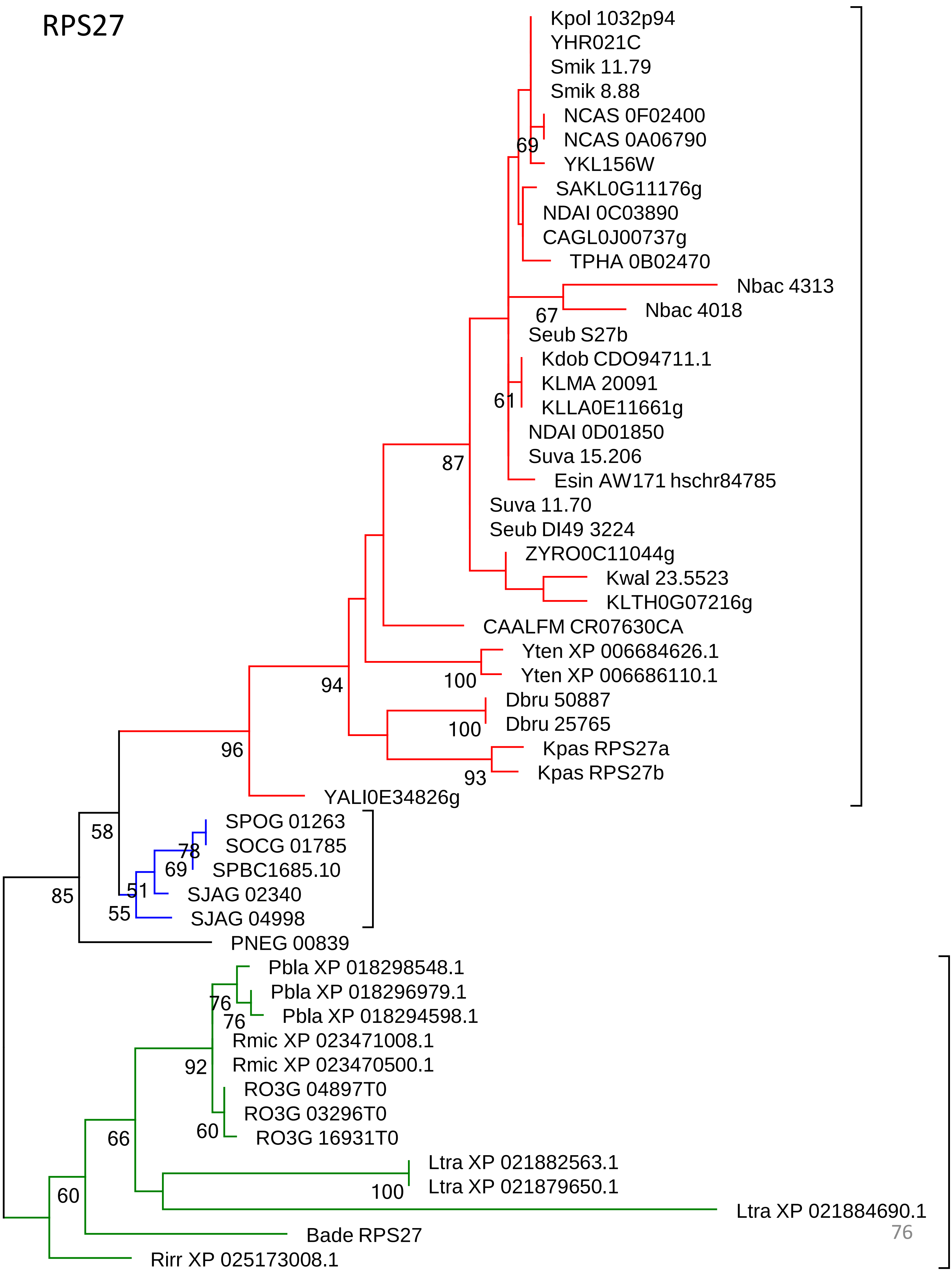

RPS27
Kpol 1032p94
YHR021C
Smik 11.79
Smik 8.88
NCAS 0F02400
NCAS 0A06790
69
YKL156W
SAKL0G11176g
NDAI 0C03890
CAGL0J00737g
TPHA 0B02470
Nbac 4313
Nbac 4018
67
Seub S27b
Kdob CDO94711.1
KLMA 20091
61
KLLA0E11661g
NDAI 0D01850
Suva 15.206
87
Esin AW171 hschr84785
Suva 11.70
Seub DI49 3224
ZYRO0C11044g
Kwal 23.5523
KLTH0G07216g
CAALFM CR07630CA
Yten XP 006684626.1
100
Yten XP 006686110.1
94
Dbru 50887
100
Dbru 25765
96
Kpas RPS27a
93
Kpas RPS27b
YALI0E34826g
58
SPOG 01263
78
SOCG 01785
69
SPBC1685.10
85
51
SJAG 02340
55
SJAG 04998
PNEG 00839
Pbla XP 018298548.1
Pbla XP 018296979.1
76
76
Pbla XP 018294598.1
Rmic XP 023471008.1
92
Rmic XP 023470500.1
RO3G 04897T0
RO3G 03296T0
66
60
RO3G 16931T0
Ltra XP 021882563.1
100
Ltra XP 021879650.1
60
Ltra XP 021884690.1
Bade RPS27
Rirr XP 025173008.1

### RPS28

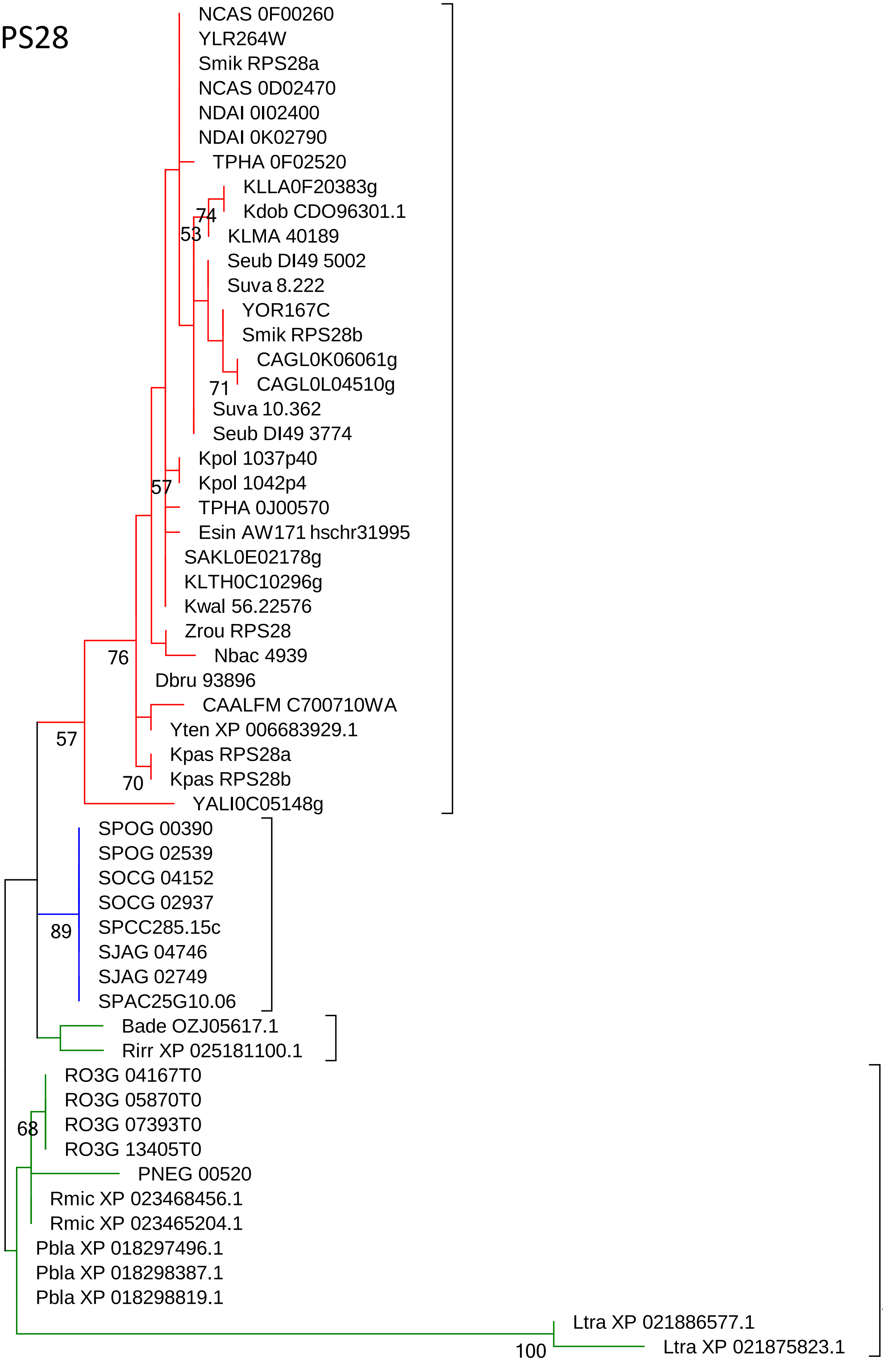
NCAS 0F00260
YLR264W
Smik RPS28a
NCAS 0D02470
NDAI 0I02400
NDAI 0K02790
TPHA 0F02520
KLLA0F20383g
Kdob CDO96301.1
74
53
KLMA 40189
Seub DI49 5002
Suva 8.222
YOR167C
Smik RPS28b
CAGL0K06061g
71
CAGL0L04510g
Suva 10.362
Seub DI49 3774
Kpol 1037p40
Kpol 1042p4
57
TPHA 0J00570
Esin AW171 hschr31995
SAKL0E02178g
KLTH0C10296g
Kwal 56.22576
Zrou RPS28
Nbac 4939
76
Dbru 93896
CAALFM C700710WA
57
Yten XP 006683929.1
Kpas RPS28a
70
Kpas RPS28b
YALI0C05148g
SPOG 00390
SPOG 02539
SOCG 04152
SOCG 02937
89
SPCC285.15c
SJAG 04746
SJAG 02749
SPAC25G10.06
Bade OZJ05617.1
Rirr XP 025181100.1
RO3G 04167T0
RO3G 05870T0
68
RO3G 07393T0
RO3G 13405T0
PNEG 00520
Rmic XP 023468456.1
Rmic XP 023465204.1
Pbla XP 018297496.1
Pbla XP 018298387.1
Pbla XP 018298819.1
Ltra XP 021886577.1
100
Ltra XP 021875823.1

### RPS29

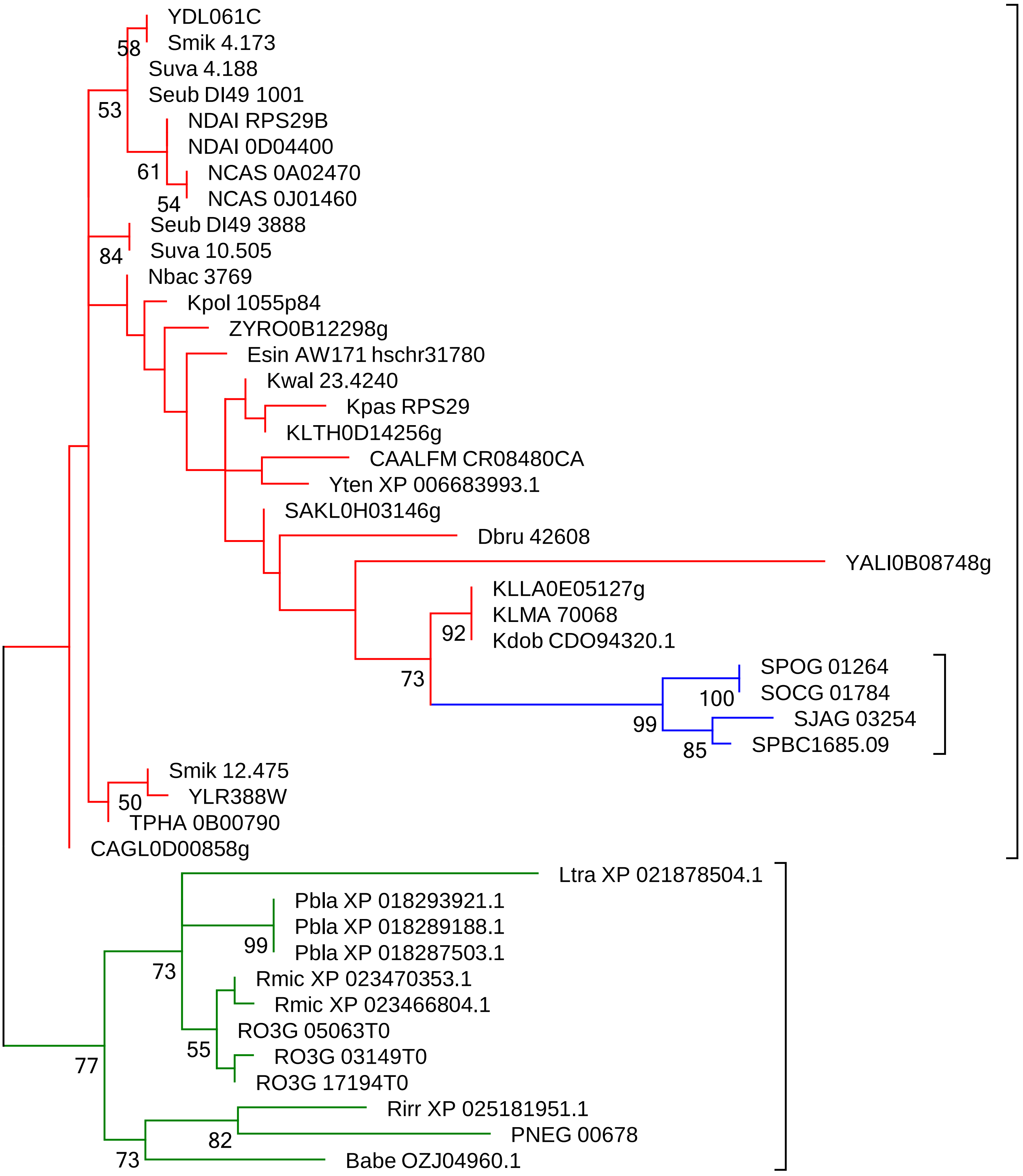
YDL061C
Smik 4.173
58
Suva 4.188
Seub DI49 1001
53
NDAI RPS29B
NDAI 0D04400
61
NCAS 0A02470
54
NCAS 0J01460
Seub DI49 3888
84
Suva 10.505
Nbac 3769
Kpol 1055p84
ZYRO0B12298g
Esin AW171 hschr31780
Kwal 23.4240
Kpas RPS29
KLTH0D14256g
CAALFM CR08480CA
Yten XP 006683993.1
SAKL0H03146g
Dbru 42608
YALI0B08748g
KLLA0E05127g
KLMA 70068
92
Kdob CDO94320.1
SPOG 01264
73
100
SOCG 01784
99
SJAG 03254
85
SPBC1685.09
Smik 12.475
50
YLR388W
TPHA 0B00790
CAGL0D00858g
Ltra XP 021878504.1
Pbla XP 018293921.1
Pbla XP 018289188.1
99
Pbla XP 018287503.1
73
Rmic XP 023470353.1
Rmic XP 023466804.1
RO3G 05063T0
55
RO3G 03149T0
77
RO3G 17194T0
Rirr XP 025181951.1
82
PNEG 00678
73
Babe OZJ04960.1

### RPS30

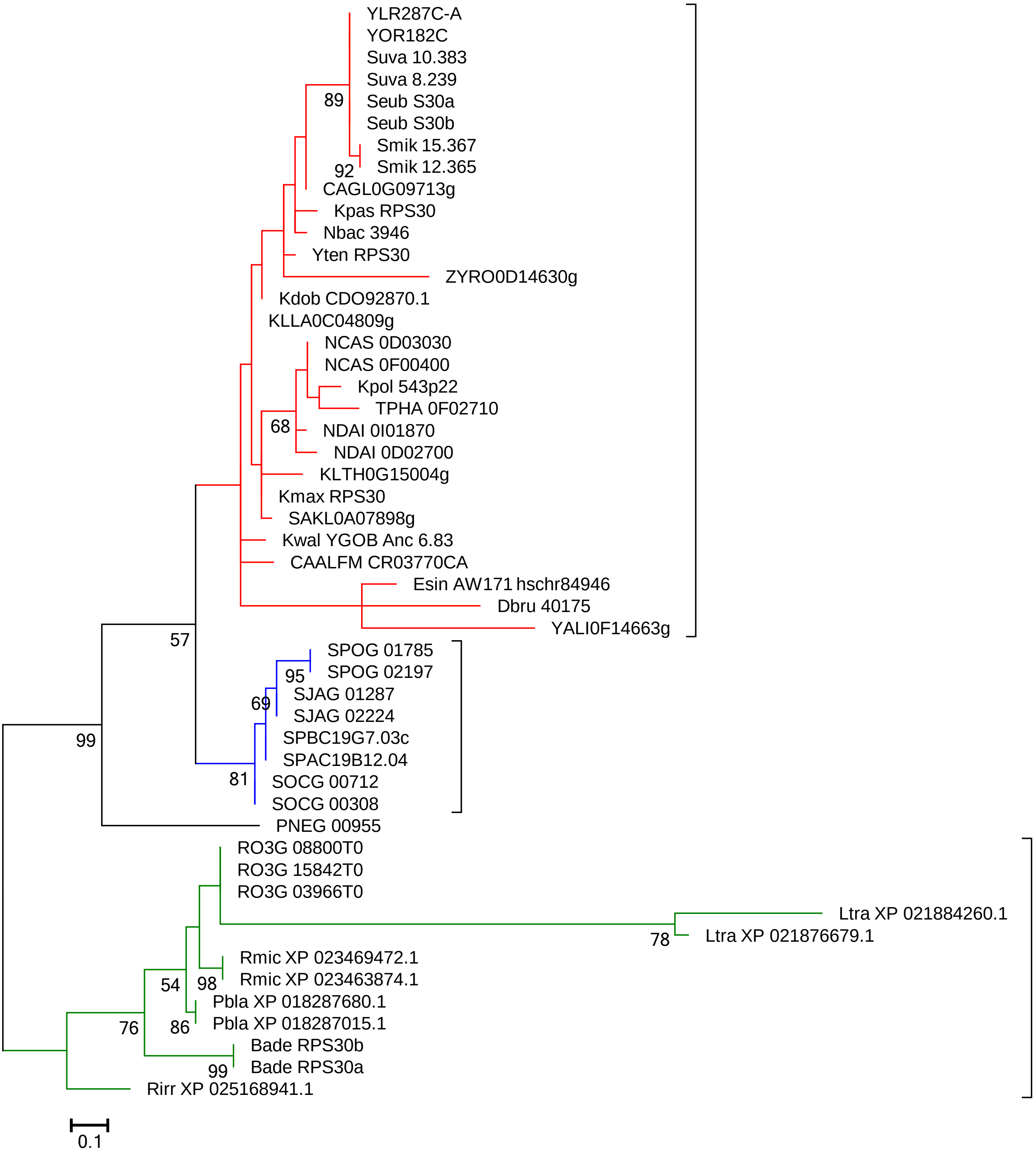
YLR287C-A
YOR182C
Suva 10.383
Suva 8.239
Seub S30a
Seub S30b
Smik 15.367
Smik 12.365
89
92
CAGL0G09713g
Kpas RPS30
Nbac 3946
Yten RPS30
ZYRO0D14630g
Kdob CDO92870.1
KLLA0C04809g
NCAS 0D03030
NCAS 0F00400
Kpol 543p22
TPHA 0F02710
68
NDAI 0I01870
NDAI 0D02700
KLTH0G15004g
Kmax RPS30
SAKL0A07898g
Kwal YGOB Anc 6.83
CAALFM CR03770CA
Esin AW171 hschr84946
Dbru 40175
YALI0F14663g
57
SPOG 01785
SPOG 02197
95
SJAG 01287
69
SJAG 02224
SPBC19G7.03c
SPAC19B12.04
81
SOCG 00712
SOCG 00308
99
PNEG 00955
RO3G 08800T0
RO3G 15842T0
RO3G 03966T0
Ltra XP 021884260.1
78
Ltra XP 021876679.1
Rmic XP 023469472.1
54
98
Rmic XP 023463874.1
Pbla XP 018287680.1
76
86
Pbla XP 018287015.1
Bade RPS30b
99
Bade RPS30a
Rirr XP 025168941.1
0.1

### RPS31 (RPS27A)

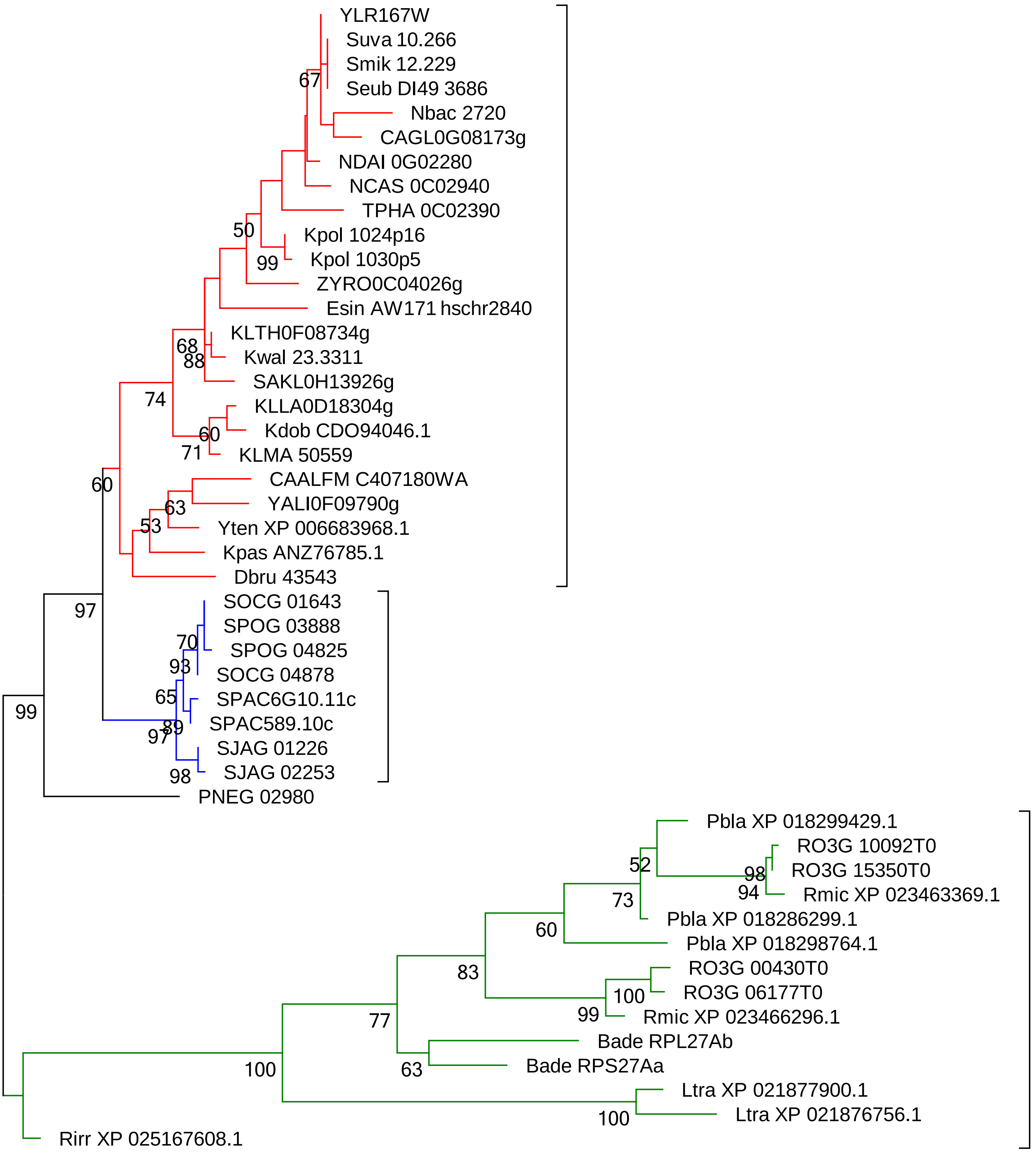
