## Supplemental File 2 for "Parallel Concerted Evolution of Ribosomal Protein Genes in Fungi and Its Adaptive Significance"

**Supplementary File 2: Phylogenetic trees of RP families in the five WGD budding yeast species.**

| Gene family | Category | Page | Gene family | Category | Page | Gene family | Category | Page |
|---|---|---|---|---|---|---|---|---|
| RPL1 | A | 2 | RPL6 | C | 12 | RPL22 | D | 20 |
| RPL4 | A | 2 | RPL16 | C | 12 | RPL37 | D | 20 |
| RPL7 | A | 2 | RPL21 | C | 12 | RPP1 | D | 20 |
| RPL8 | A | 3 | RPL23 | C | 13 | RPP2 | D | 21 |
| RPL9 | A | 3 | RPL26 | C | 13 | RPS7 | D | 21 |
| RPL12 | A | 3 | RPL27 | C | 13 | RPS27 | D | 21 |
| RPL13 | A | 4 | RPL31 | C | 14 | RPS29 | D | 22 |
| RPL14 | A | 4 | RPL33 | C | 14 | | | |
| RPL15 | A | 4 | RPL34 | C | 14 | | | |
| RPL18 | A | 5 | RPL36 | C | 15 | | | |
| RPL20 | A | 5 | RPL40 | C | 15 | | | |
| RPL35 | A | 5 | RPL41 | C | 15 | | | |
| RPL42 | A | 6 | RPS0 | C | 16 | | | |
| RPL43 | A | 6 | RPS1 | C | 16 | | | |
| RPS4 | A | 6 | RPS9 | C | 16 | | | |
| RPS6 | A | 7 | RPS10 | C | 17 | | | |
| RPS8 | A | 7 | RPS17 | C | 17 | | | |
| RPS11 | A | 7 | RPS21 | C | 17 | | | |
| RPS18 | A | 8 | RPS22 | C | 18 | | | |
| RPS19 | A | 8 | RPS23 | C | 18 | | | |
| RPS24 | A | 8 | RPS25 | C | 18 | | | |
| RPL2 | B | 9 | RPS26 | C | 19 | | | |
| RPL11 | B | 9 | RPS28 | C | 19 | | | |
| RPL17 | B | 9 | RPS30 | C | 19 | | | |
| RPL19 | B | 10 | | | | | | |
| RPL24 | B | 10 | | | | | | |
| RPS14 | B | 10 | | | | | | |
| RPS16 | B | 11 | | | | | | |

*The phylogenetic trees were built based on amino acid sequences of four WGD budding yeast species, unless specially specified. Each tree was reconstructed by Neighbor-Joining method using Poisson correction with 1000 Bootstrap replicates. For some trees, if the phylogenetic tree topology is not resolved due to high degree of sequence similarity, we used their CDS sequences to build their phylogenetic tree.

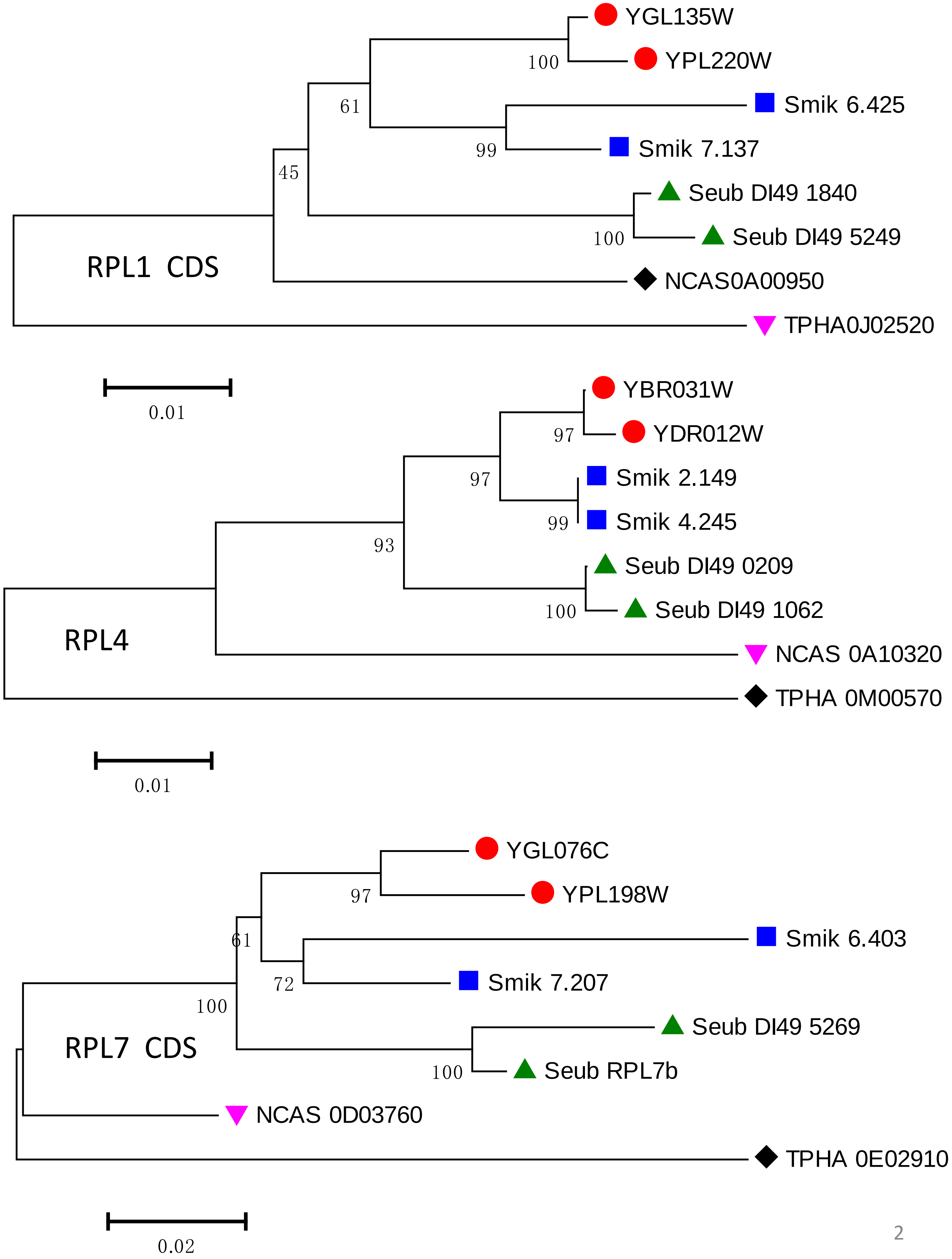
YGL135W
YPL220W
Smik 6.425
Smik 7.137
Seub DI49 1840
Seub DI49 5249
NCAS0A00950
TPHA0J02520
100
61
99
45
100
RPL1 CDS
0.01
YBR031W
YDR012W
Smik 2.149
Smik 4.245
Seub DI49 0209
Seub DI49 1062
NCAS 0A10320
TPHA 0M00570
97
97
99
93
100
RPL4
0.01
YGL076C
YPL198W
Smik 6.403
Smik 7.207
Seub DI49 5269
Seub RPL7b
NCAS 0D03760
TPHA 0E02910
97
61
72
100
100
RPL7 CDS
0.02

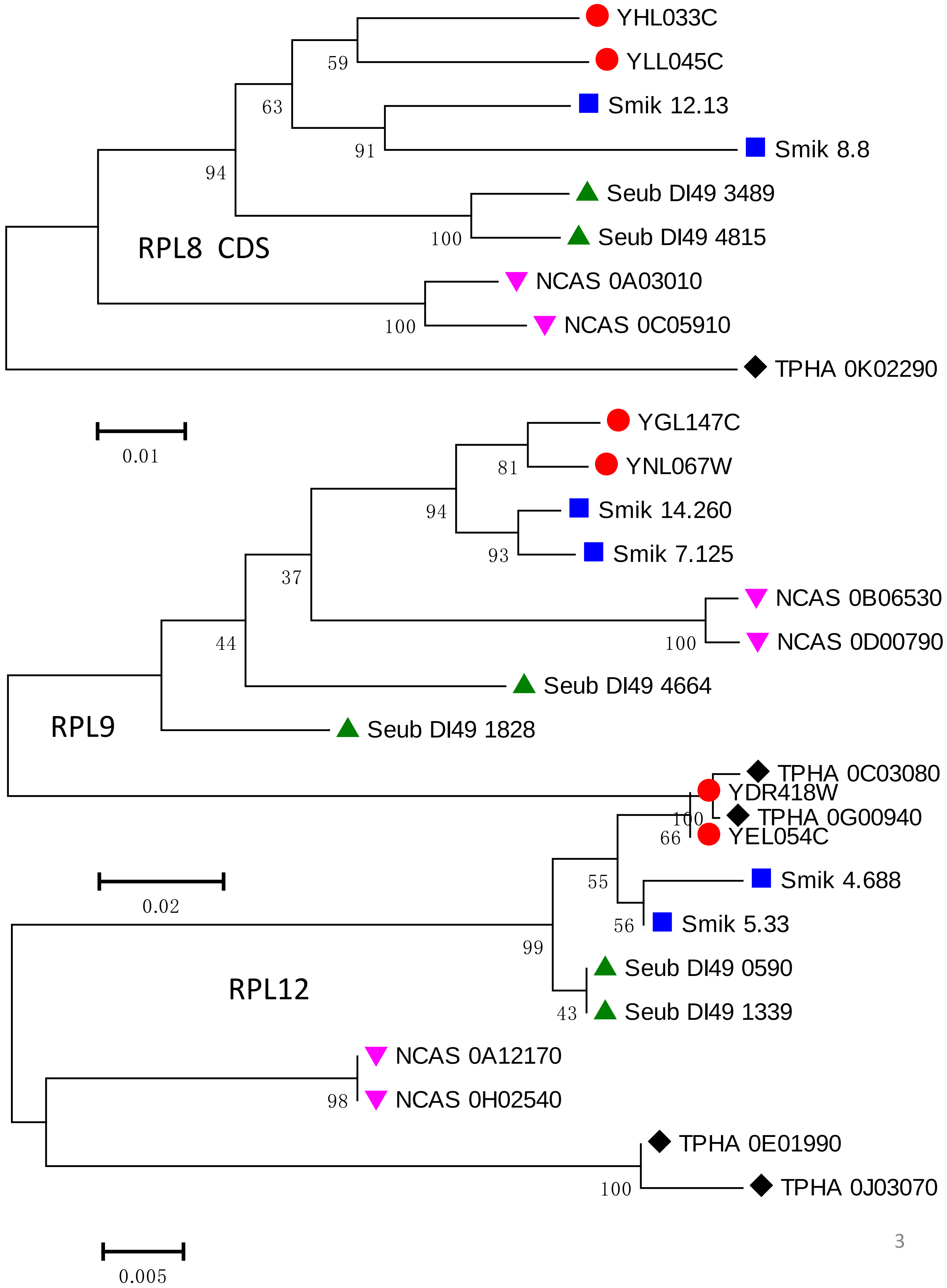
RPL8 CDS
YHL033C
YLL045C
Smik 12.13
Smik 8.8
Seub DI49 3489
Seub DI49 4815
NCAS 0A03010
NCAS 0C05910
TPHA 0K02290
59
63
91
94
100
100
0.01
RPL9
YGL147C
YNL067W
Smik 14.260
Smik 7.125
NCAS 0B06530
NCAS 0D00790
Seub DI49 4664
Seub DI49 1828
81
94
93
37
100
44
0.02
RPL12
TPHA 0C03080
YDR418W
TPHA 0G00940
YEL054C
Smik 4.688
Smik 5.33
Seub DI49 0590
Seub DI49 1339
NCAS 0A12170
NCAS 0H02540
TPHA 0E01990
TPHA 0J03070
100
66
55
56
99
43
98
100
0.005

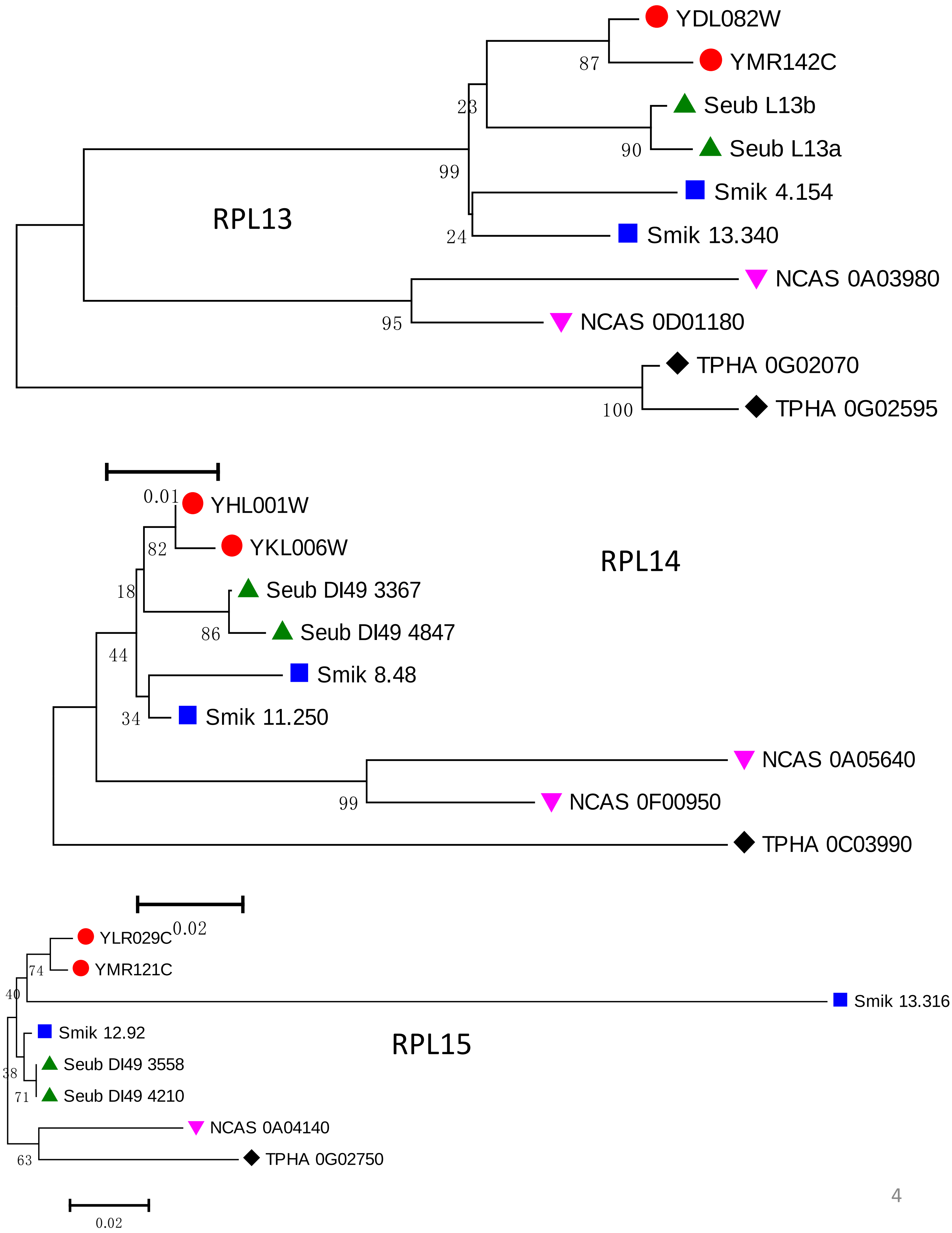
YDL082W
YMR142C
87
23
Seub L13b
Seub L13a
90
99
Smik 4.154
RPL13
24
Smik 13.340
NCAS 0A03980
95
NCAS 0D01180
TPHA 0G02070
100
TPHA 0G02595
0.01
YHL001W
82
YKL006W
RPL14
18
Seub DI49 3367
86
Seub DI49 4847
44
Smik 8.48
34
Smik 11.250
NCAS 0A05640
99
NCAS 0F00950
TPHA 0C03990
0.02
YLR029C
74
YMR121C
40
Smik 13.316
Smik 12.92
RPL15
38
Seub DI49 3558
71
Seub DI49 4210
NCAS 0A04140
63
TPHA 0G02750
0.02

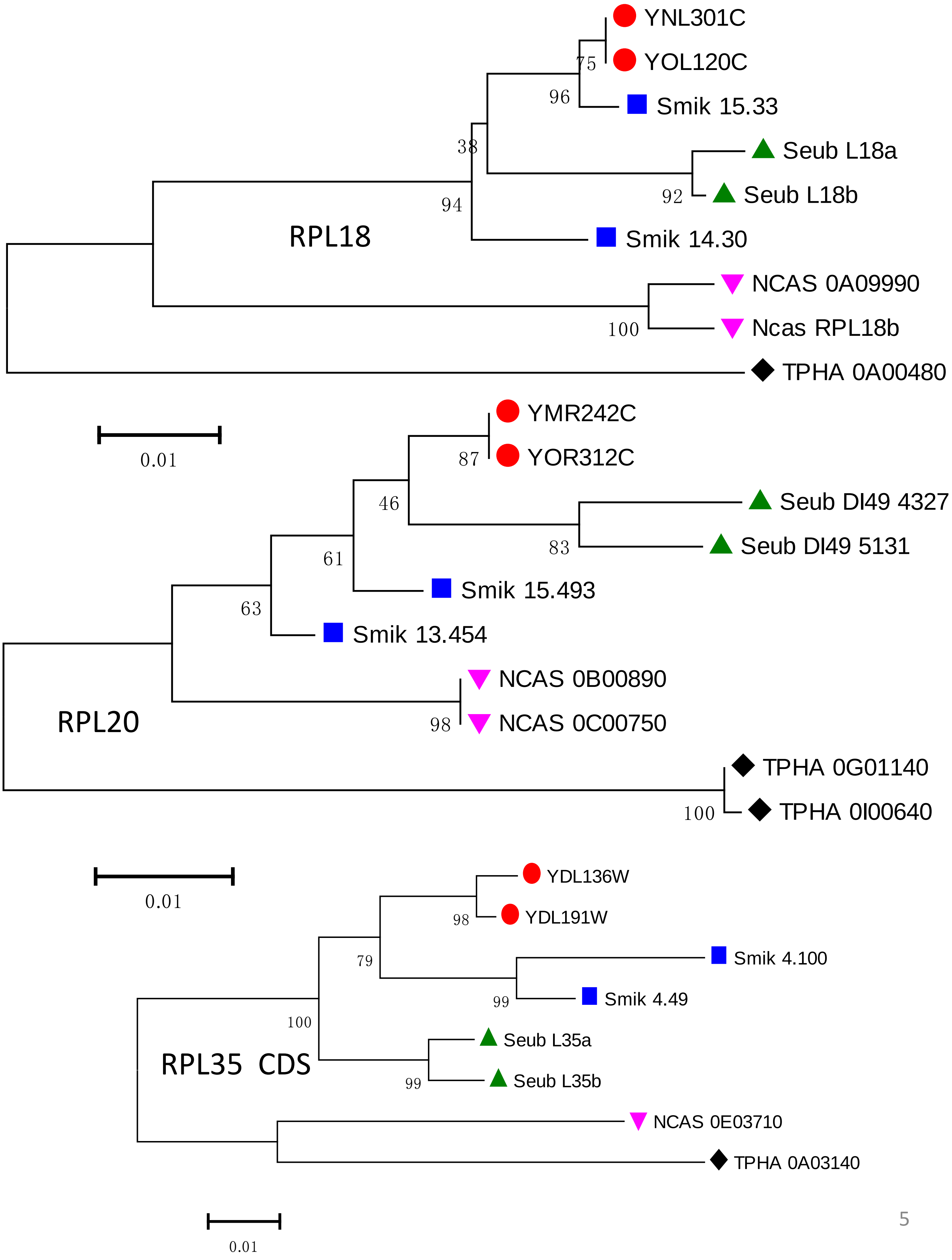
YNL301C
YOL120C
75
96
Smik 15.33
38
Seub L18a
92
Seub L18b
94
RPL18
Smik 14.30
NCAS 0A09990
100
Ncas RPL18b
TPHA 0A00480
0.01
YMR242C
87
YOR312C
46
Seub DI49 4327
83
Seub DI49 5131
61
Smik 15.493
63
Smik 13.454
NCAS 0B00890
98
NCAS 0C00750
RPL20
TPHA 0G01140
100
TPHA 0I00640
0.01
YDL136W
98
YDL191W
79
Smik 4.100
99
Smik 4.49
100
Seub L35a
RPL35 CDS
99
Seub L35b
NCAS 0E03710
TPHA 0A03140
0.01

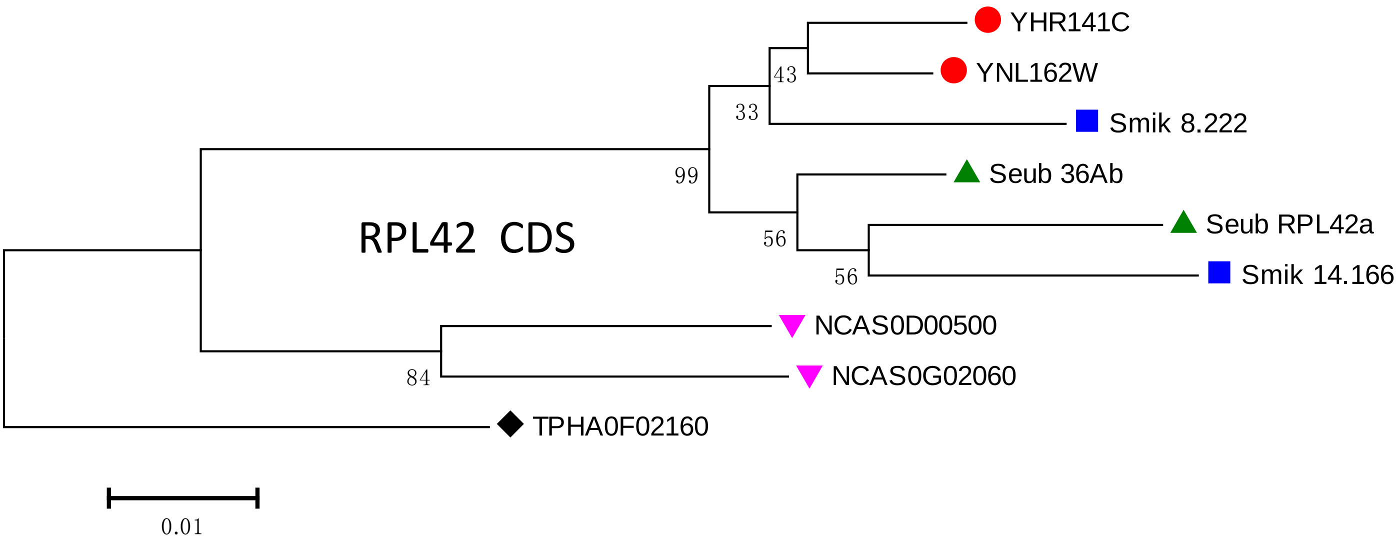
RPL42 CDS
YHR141C
YNL162W
Smik 8.222
Seub 36Ab
Seub RPL42a
Smik 14.166
NCAS0D00500
NCAS0G02060
TPHA0F02160
43
33
99
56
56
84
0.01

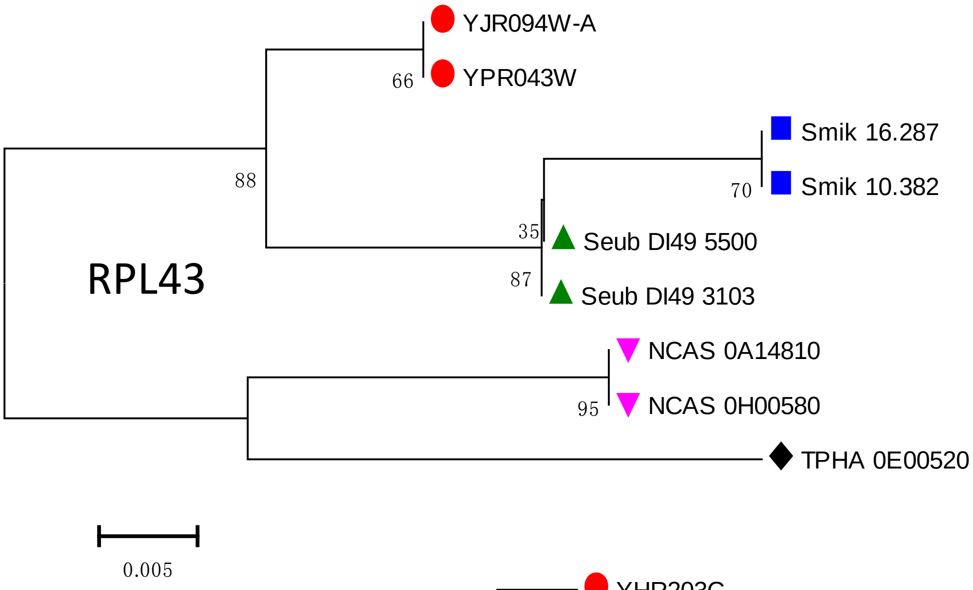
RPL43
YJR094W-A
YPR043W
Smik 16.287
Smik 10.382
Seub DI49 5500
Seub DI49 3103
NCAS 0A14810
NCAS 0H00580
TPHA 0E00520
66
88
70
35
87
95
0.005

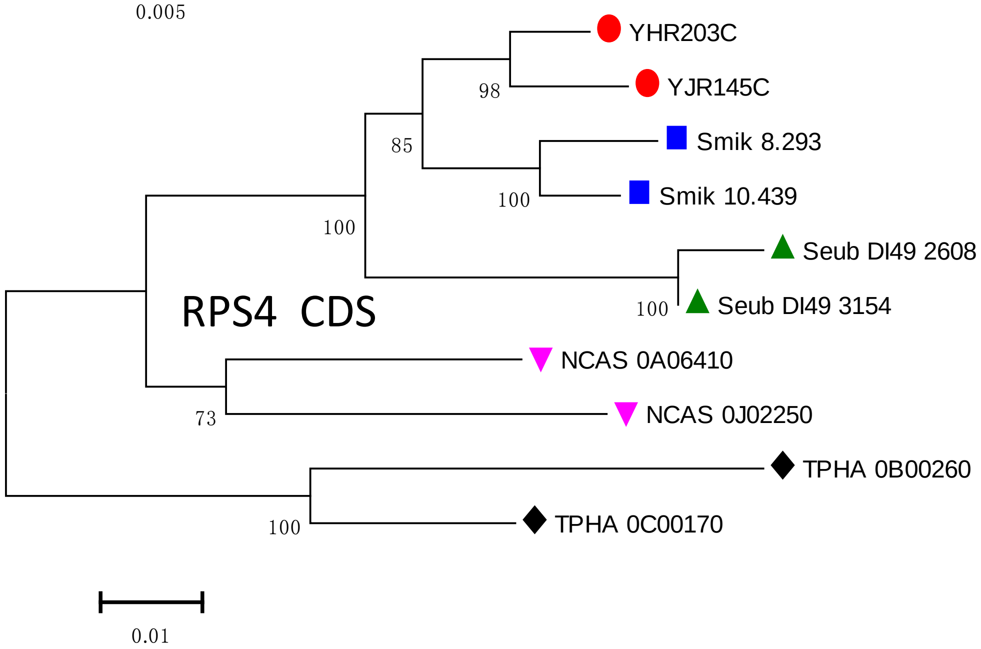
RPS4 CDS
YHR203C
YJR145C
Smik 8.293
Smik 10.439
Seub DI49 2608
Seub DI49 3154
NCAS 0A06410
NCAS 0J02250
TPHA 0B00260
TPHA 0C00170
98
85
100
100
100
73
100
0.01

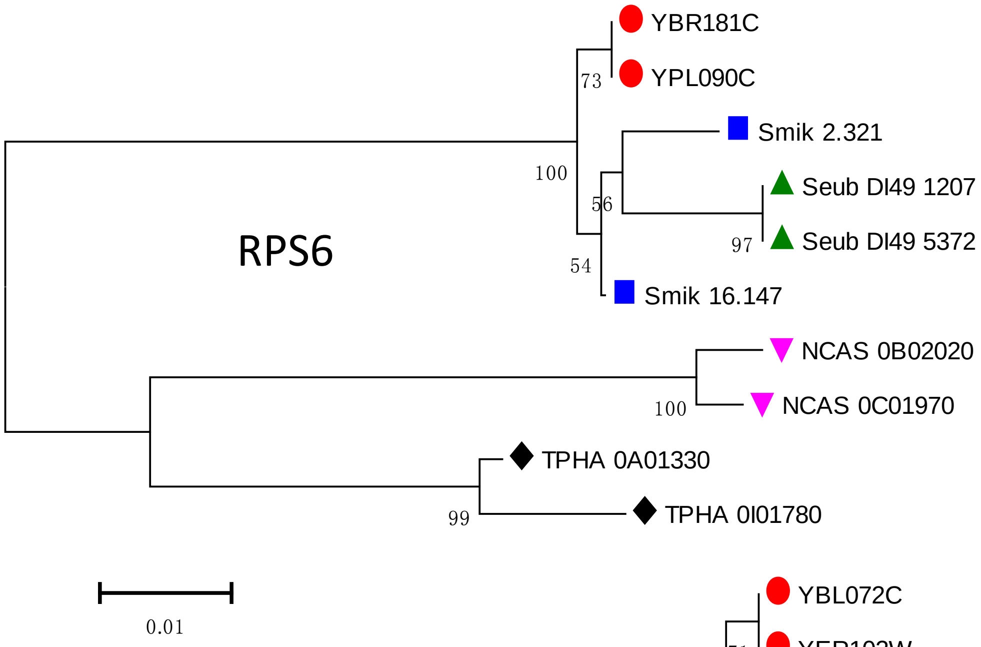
YBR181C
YPL090C
73
Smik 2.321
100
56
Seub DI49 1207
RPS6
97
Seub DI49 5372
54
Smik 16.147
NCAS 0B02020
100
NCAS 0C01970
TPHA 0A01330
99
TPHA 0I01780
0.01

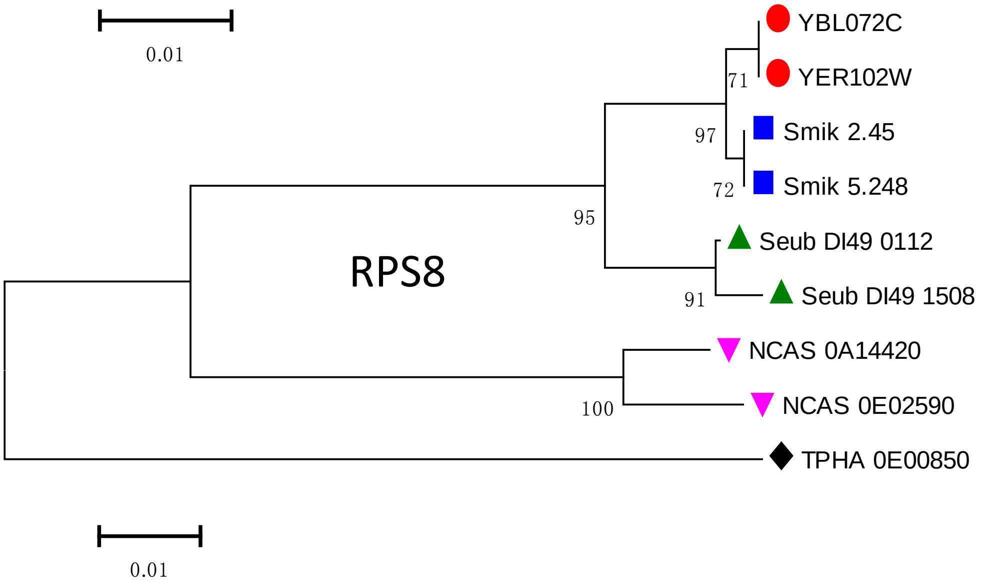
YBL072C
71
YER102W
97
Smik 2.45
72
Smik 5.248
95
Seub DI49 0112
RPS8
91
Seub DI49 1508
NCAS 0A14420
100
NCAS 0E02590
TPHA 0E00850
0.01

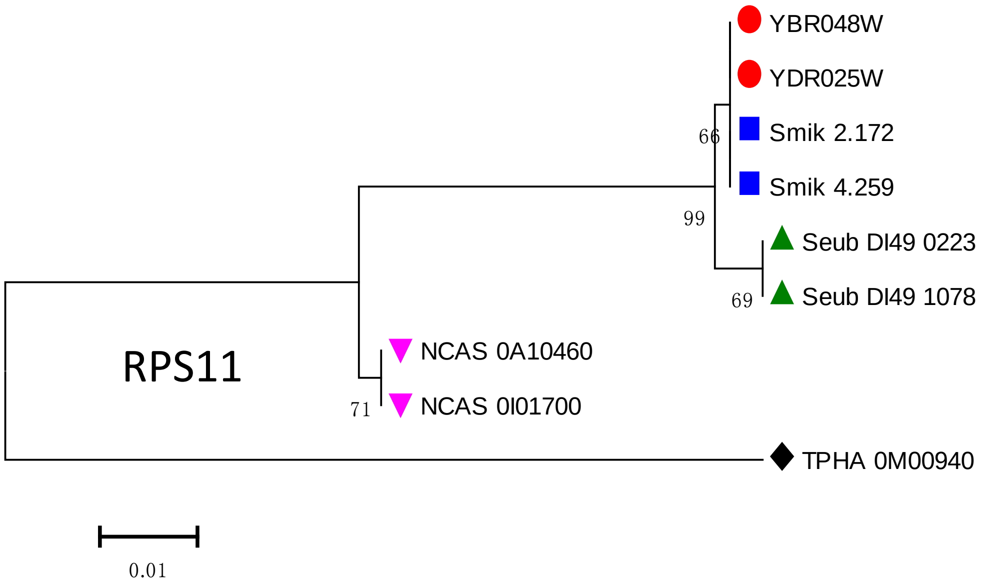
YBR048W
YDR025W
66
Smik 2.172
Smik 4.259
99
Seub DI49 0223
69
Seub DI49 1078
NCAS 0A10460
RPS11
71
NCAS 0I01700
TPHA 0M00940
0.01

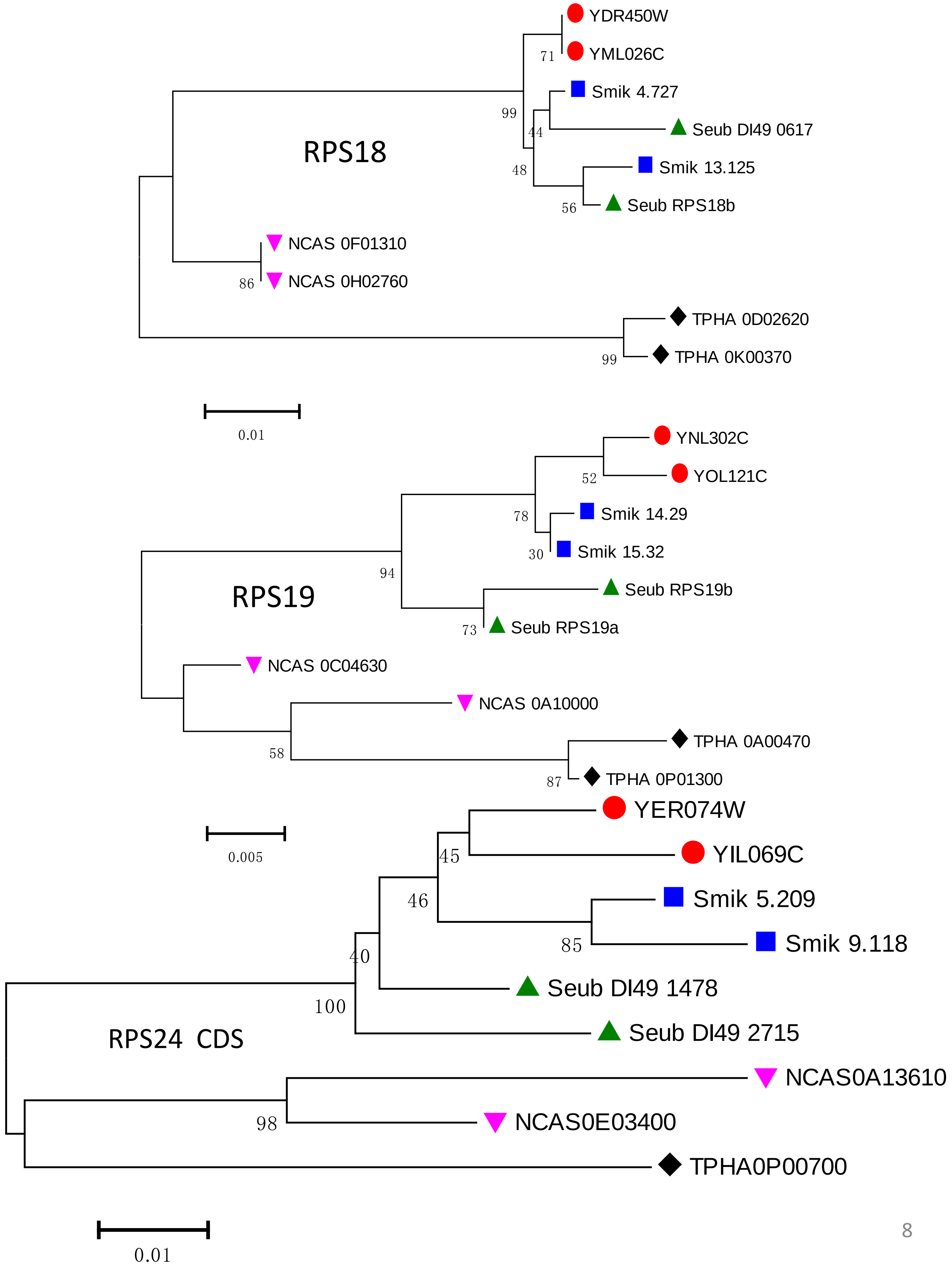
RPS18
YDR450W
YML026C
71
Smik 4.727
Seub DI49 0617
99
44
Smik 13.125
48
Seub RPS18b
56
NCAS 0F01310
NCAS 0H02760
86
TPHA 0D02620
TPHA 0K00370
99
0.01
RPS19
YNL302C
YOL121C
52
Smik 14.29
78
Smik 15.32
30
Seub RPS19b
94
Seub RPS19a
73
NCAS 0C04630
NCAS 0A10000
TPHA 0A00470
58
TPHA 0P01300
87
0.005
RPS24 CDS
YER074W
YIL069C
45
Smik 5.209
46
Smik 9.118
85
40
Seub DI49 1478
100
Seub DI49 2715
NCAS0A13610
98
NCAS0E03400
TPHA0P00700
0.01

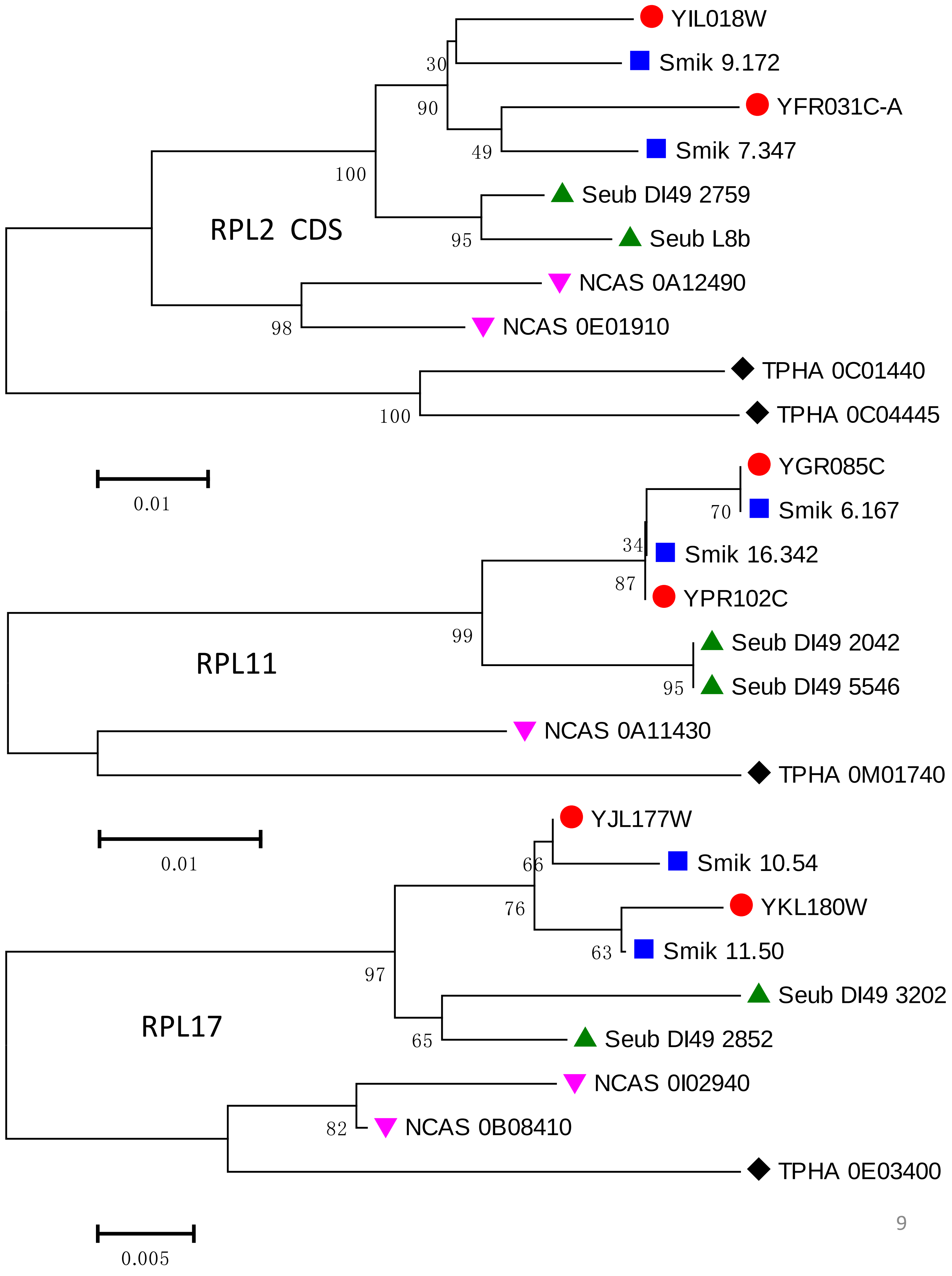

RPL2 CDS
YIL018W
Smik 9.172
YFR031C-A
Smik 7.347
Seub DI49 2759
Seub L8b
NCAS 0A12490
NCAS 0E01910
TPHA 0C01440
TPHA 0C04445
30
90
49
100
95
98
100
0.01
RPL11
YGR085C
Smik 6.167
Smik 16.342
YPR102C
Seub DI49 2042
Seub DI49 5546
NCAS 0A11430
TPHA 0M01740
70
34
87
99
95
0.01
RPL17
YJL177W
Smik 10.54
YKL180W
Smik 11.50
Seub DI49 3202
Seub DI49 2852
NCAS 0I02940
NCAS 0B08410
TPHA 0E03400
66
76
63
97
65
82
0.005

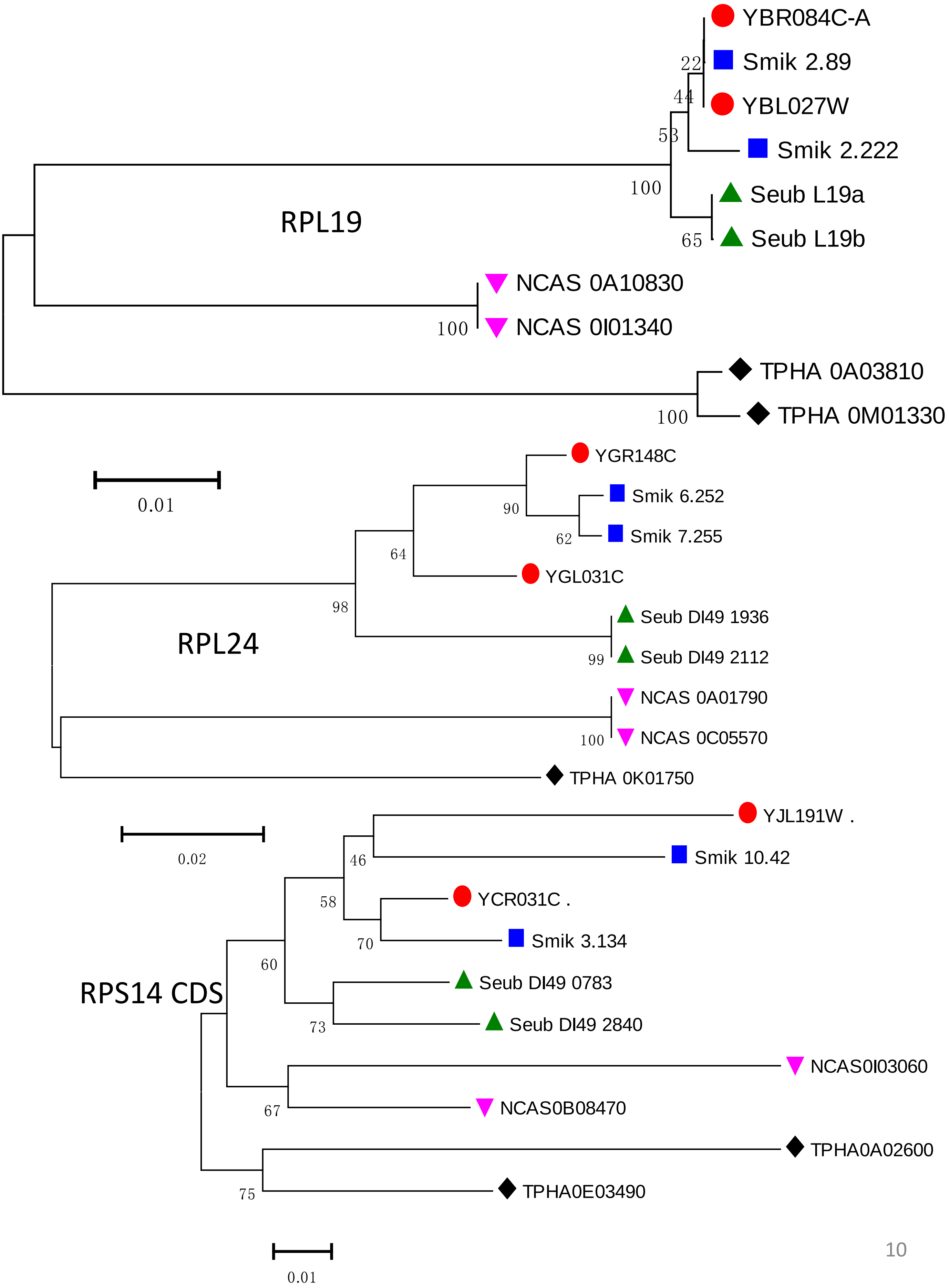
RPL19
YBR084C-A
Smik 2.89
YBL027W
Smik 2.222
Seub L19a
Seub L19b
NCAS 0A10830
NCAS 0I01340
TPHA 0A03810
TPHA 0M01330
22
44
53
100
65
100
100
0.01
RPL24
YGR148C
Smik 6.252
Smik 7.255
YGL031C
Seub DI49 1936
Seub DI49 2112
NCAS 0A01790
NCAS 0C05570
TPHA 0K01750
90
62
64
98
99
100
0.02
RPS14 CDS
YJL191W .
Smik 10.42
YCR031C .
Smik 3.134
Seub DI49 0783
Seub DI49 2840
NCAS0I03060
NCAS0B08470
TPHA0A02600
TPHA0E03490
46
58
70
60
73
67
75
0.01

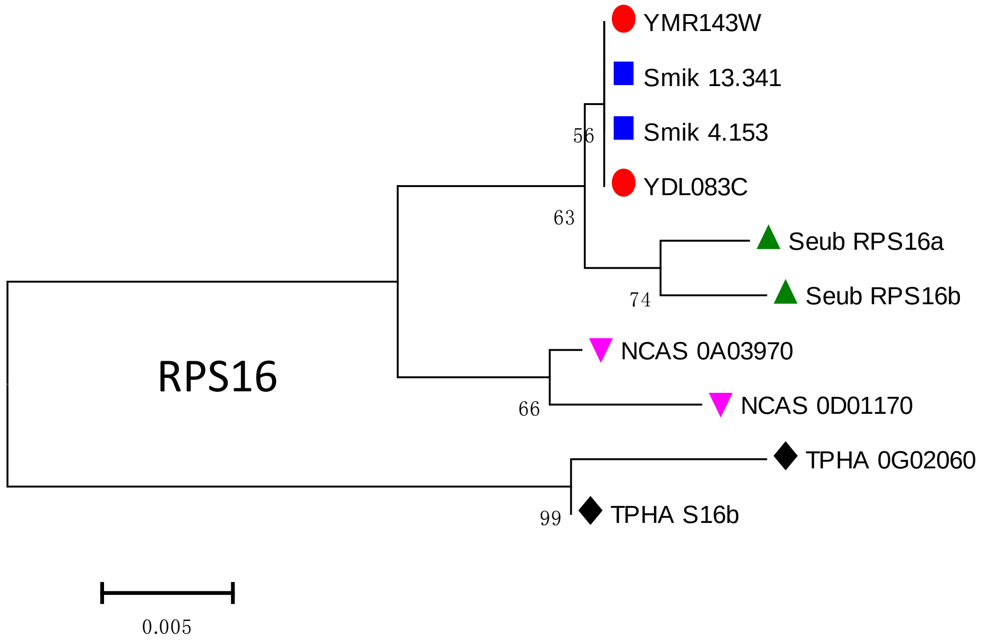
YMR143W
Smik 13.341
Smik 4.153
YDL083C
Seub RPS16a
Seub RPS16b
NCAS 0A03970
NCAS 0D01170
TPHA 0G02060
TPHA S16b
56
63
74
66
99
RPS16
0.005

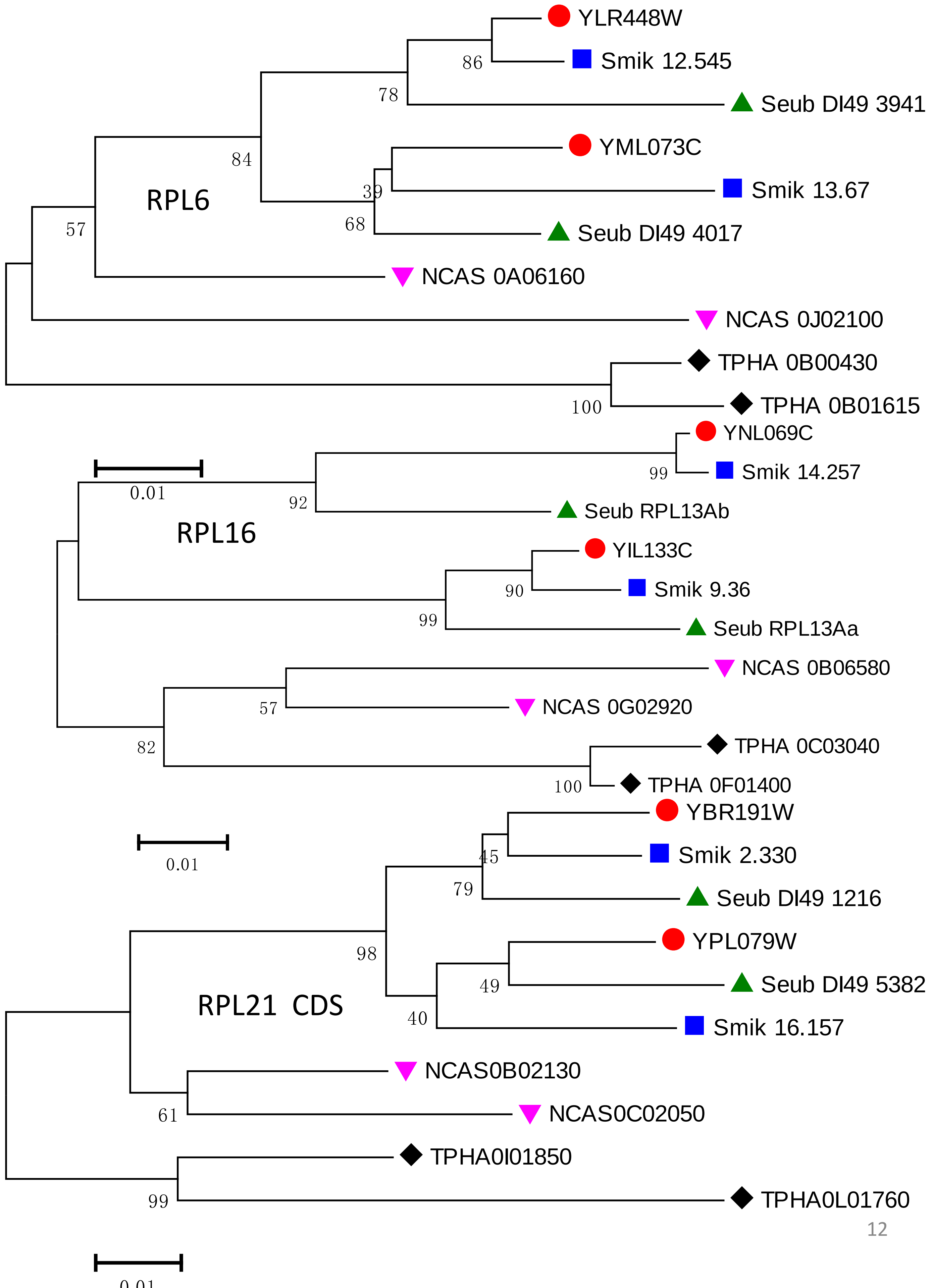
RPL6
YLR448W
Smik 12.545
Seub DI49 3941
YML073C
Smik 13.67
Seub DI49 4017
NCAS 0A06160
NCAS 0J02100
TPHA 0B00430
TPHA 0B01615
86
78
84
39
68
57
100
RPL16
YNL069C
Smik 14.257
Seub RPL13Ab
YIL133C
Smik 9.36
Seub RPL13Aa
NCAS 0B06580
NCAS 0G02920
TPHA 0C03040
TPHA 0F01400
0.01
99
92
90
99
57
82
100
0.01
RPL21 CDS
YBR191W
Smik 2.330
Seub DI49 1216
YPL079W
Seub DI49 5382
Smik 16.157
NCAS0B02130
NCAS0C02050
TPHA0I01850
TPHA0L01760
45
79
98
49
40
61
99
0.01

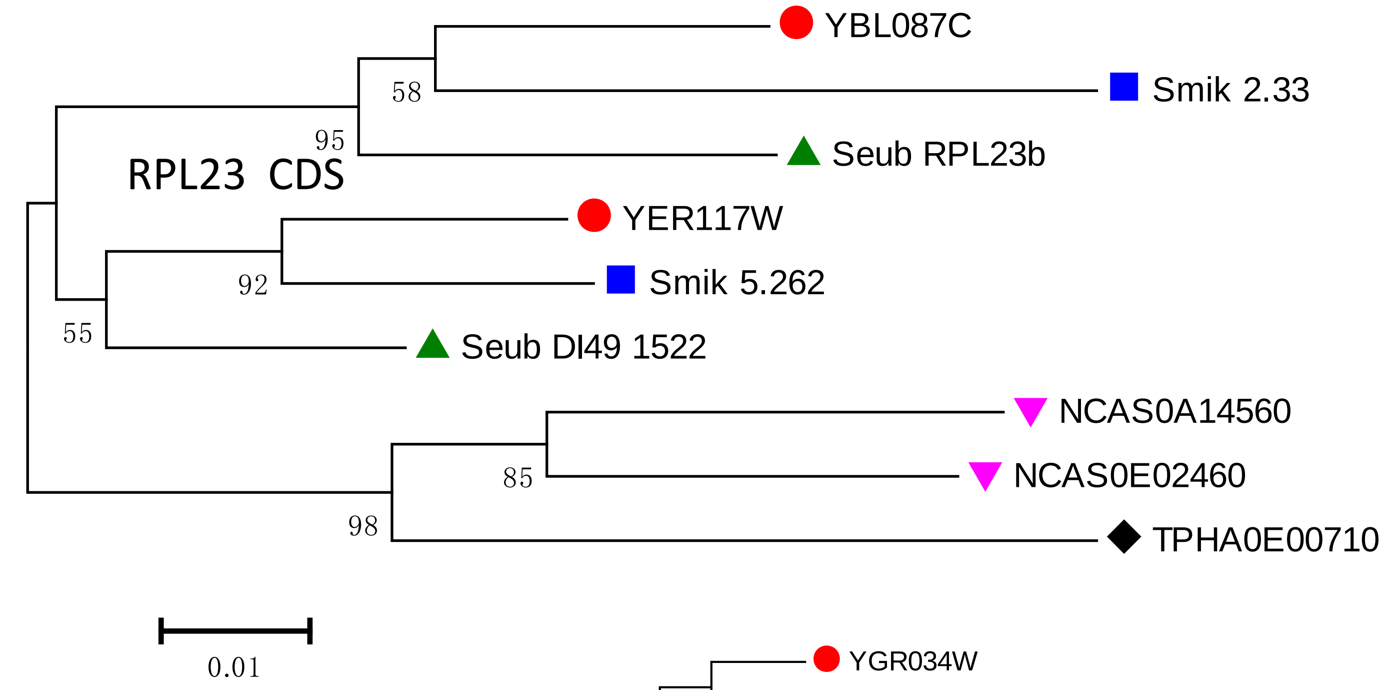
RPL23 CDS
YBL087C
Smik 2.33
Seub RPL23b
YER117W
Smik 5.262
Seub DI49 1522
NCAS0A14560
NCAS0E02460
TPHA0E00710
58
95
92
55
85
98
0.01

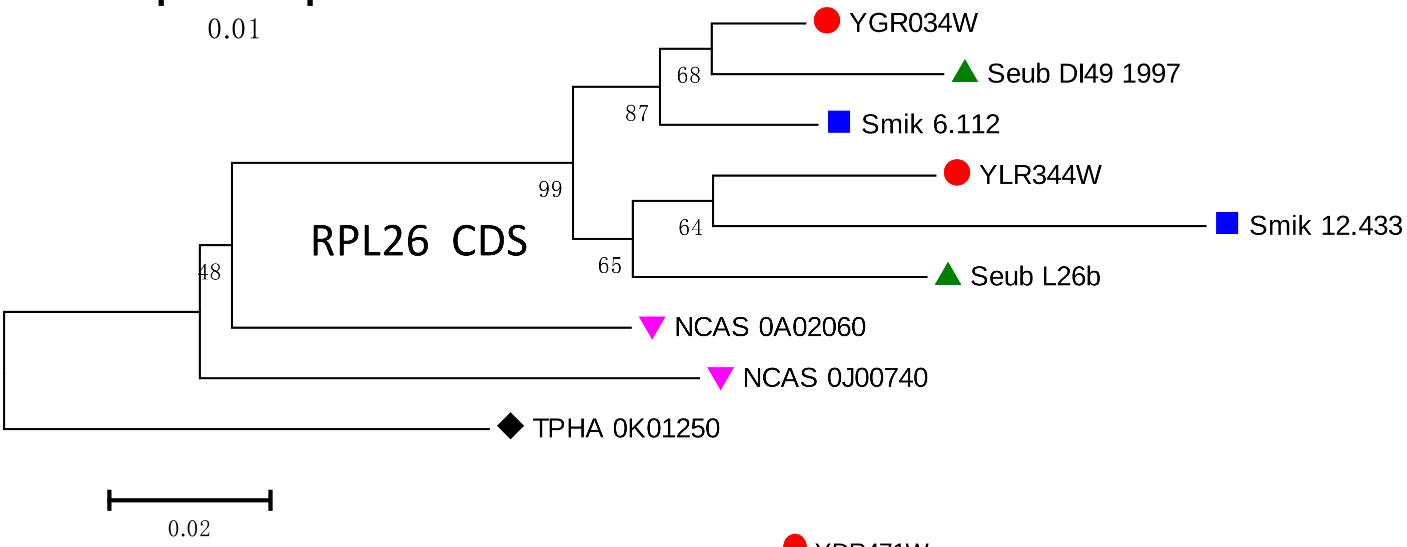
RPL26 CDS
YGR034W
Seub DI49 1997
Smik 6.112
YLR344W
Smik 12.433
Seub L26b
NCAS 0A02060
NCAS 0J00740
TPHA 0K01250
68
87
99
64
65
48
0.02

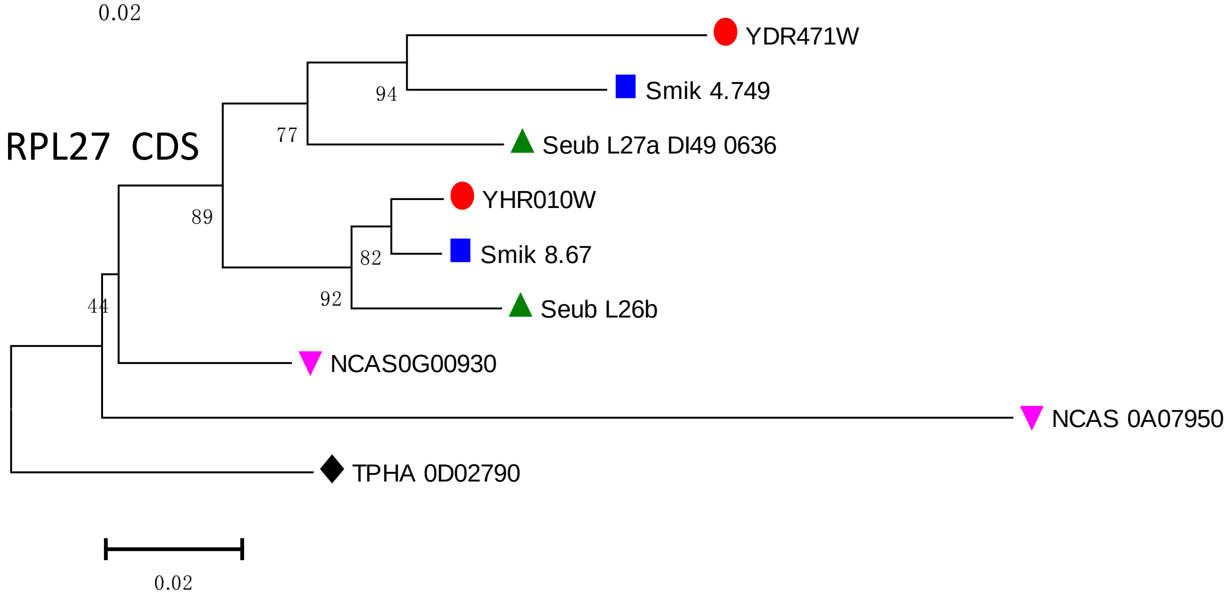
RPL27 CDS
YDR471W
Smik 4.749
Seub L27a DI49 0636
YHR010W
Smik 8.67
Seub L26b
NCAS0G00930
NCAS 0A07950
TPHA 0D02790
94
77
89
82
92
44
0.02

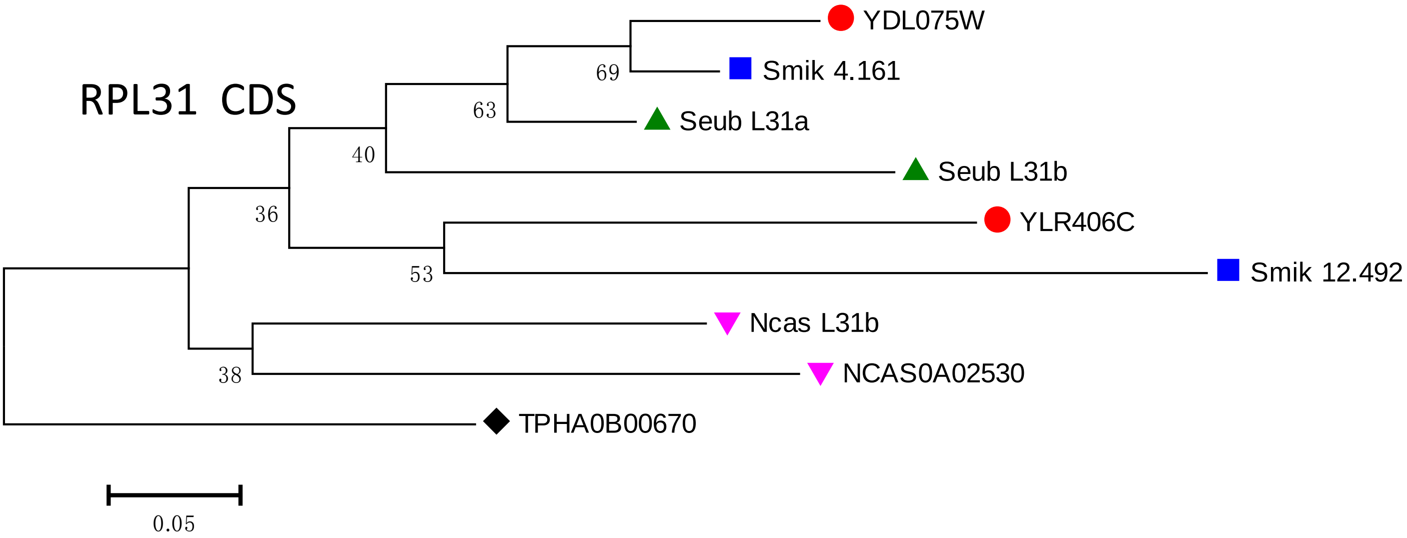
RPL31 CDS
YDL075W
Smik 4.161
Seub L31a
Seub L31b
YLR406C
Smik 12.492
Ncas L31b
NCAS0A02530
TPHA0B00670
69
63
40
36
53
38
0.05

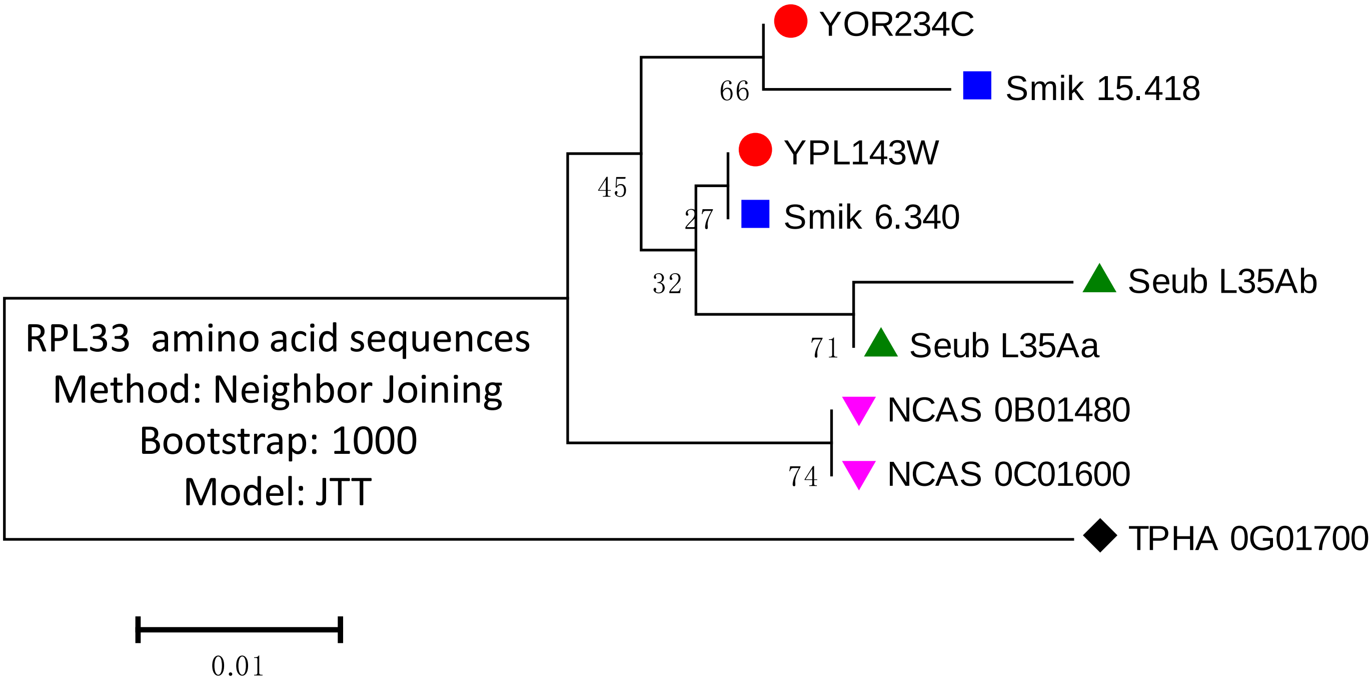
YOR234C
Smik 15.418
YPL143W
Smik 6.340
Seub L35Ab
Seub L35Aa
NCAS 0B01480
NCAS 0C01600
TPHA 0G01700
66
45
27
32
71
74
RPL33 amino acid sequences
Method: Neighbor Joining
Bootstrap: 1000
Model: JTT
0.01

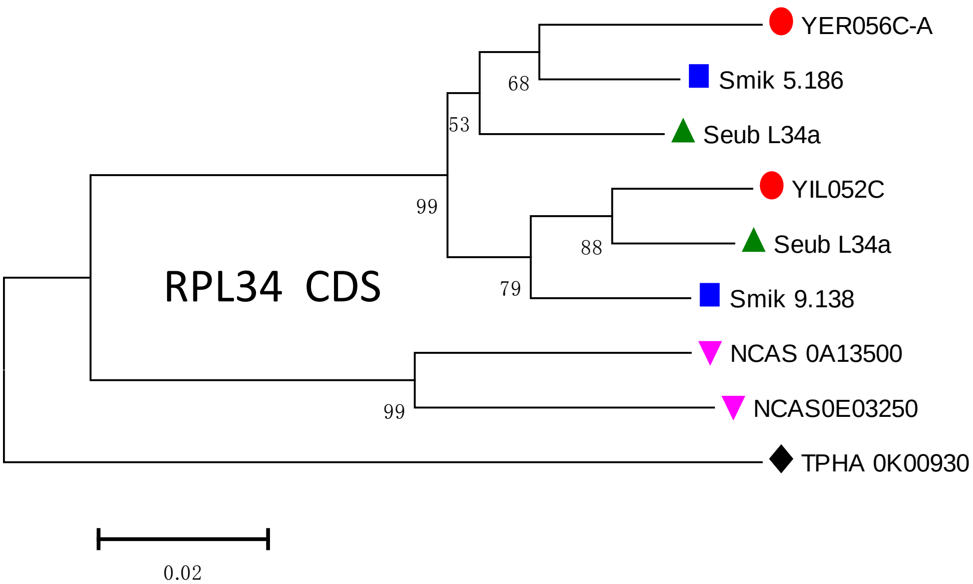
YER056C-A
Smik 5.186
Seub L34a
YIL052C
Seub L34a
Smik 9.138
NCAS 0A13500
NCAS0E03250
TPHA 0K00930
68
53
99
88
79
99
RPL34 CDS
0.02

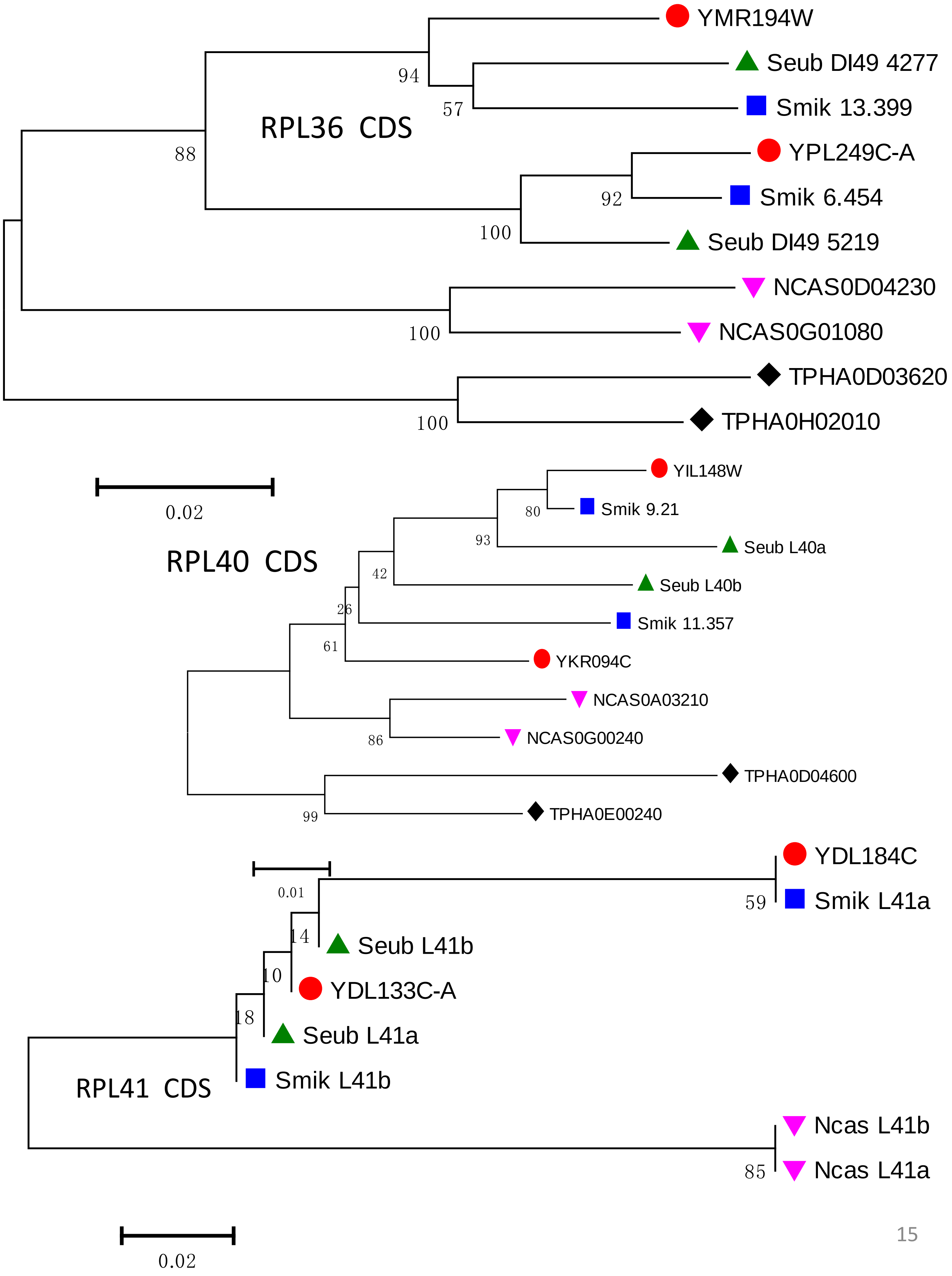
YMR194W
Seub DI49 4277
Smik 13.399
YPL249C-A
Smik 6.454
Seub DI49 5219
NCAS0D04230
NCAS0G01080
TPHA0D03620
TPHA0H02010
RPL36 CDS
94
57
88
92
100
100
100
0.02
YIL148W
Smik 9.21
Seub L40a
Seub L40b
Smik 11.357
YKR094C
NCAS0A03210
NCAS0G00240
TPHA0D04600
TPHA0E00240
RPL40 CDS
80
93
42
26
61
86
99
YDL184C
Smik L41a
Seub L41b
YDL133C-A
Seub L41a
Smik L41b
Ncas L41b
Ncas L41a
RPL41 CDS
0.01
59
14
10
18
85
0.02

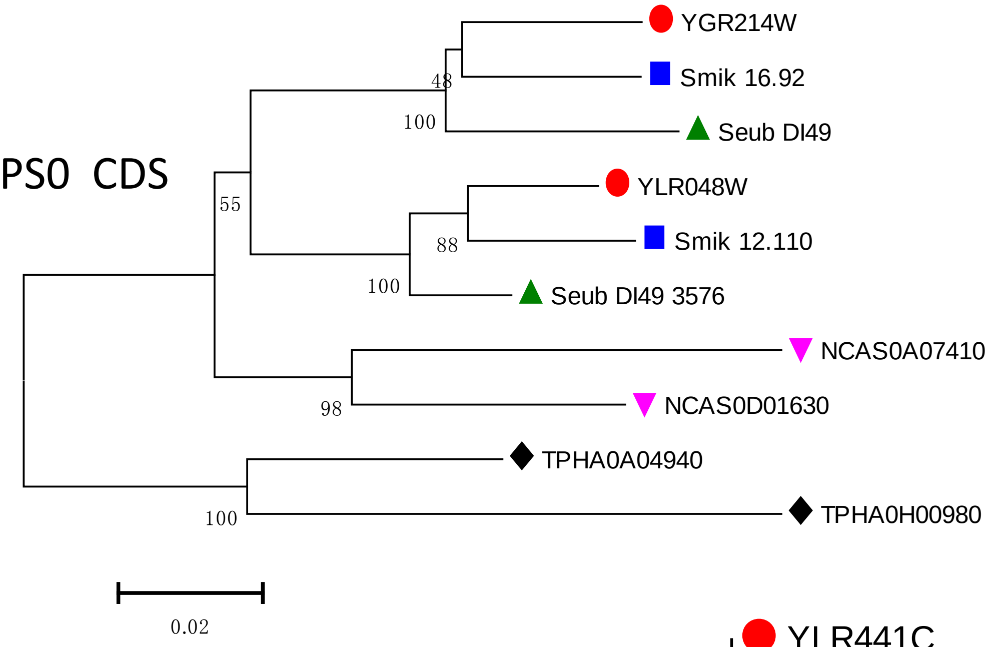


RPS0 CDS

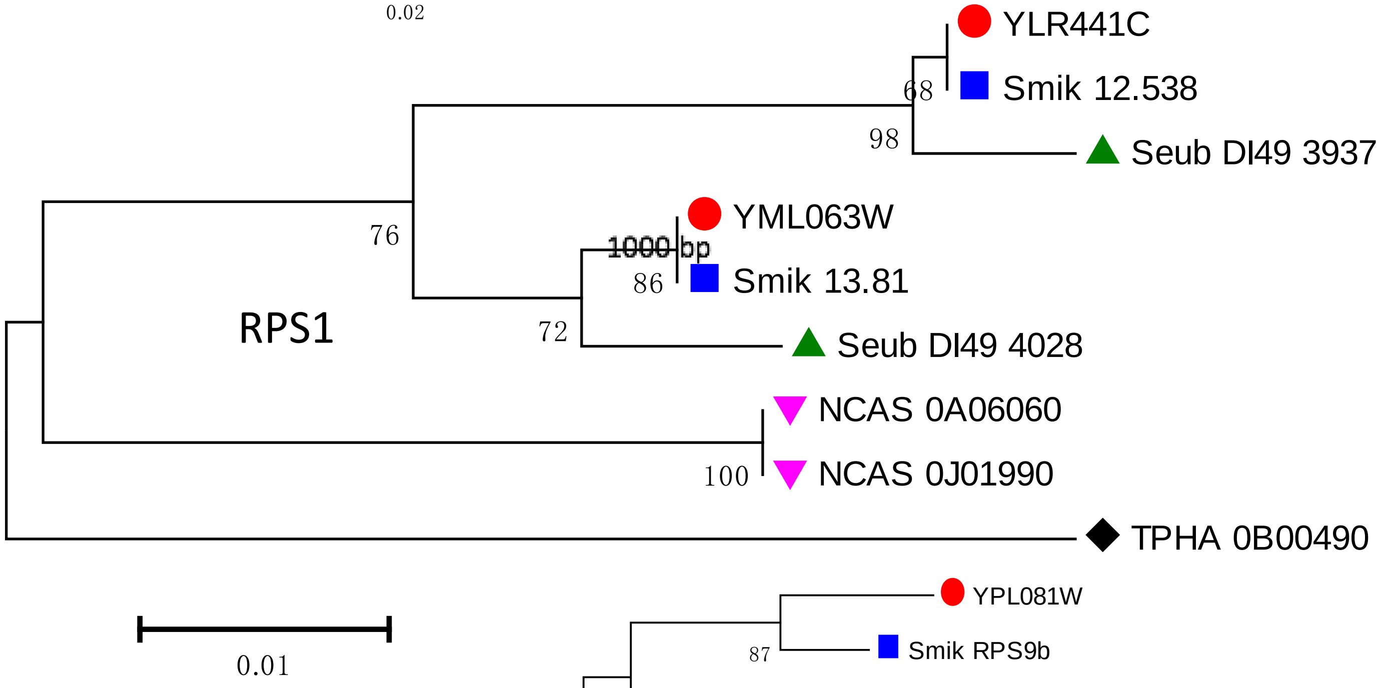


RPS1

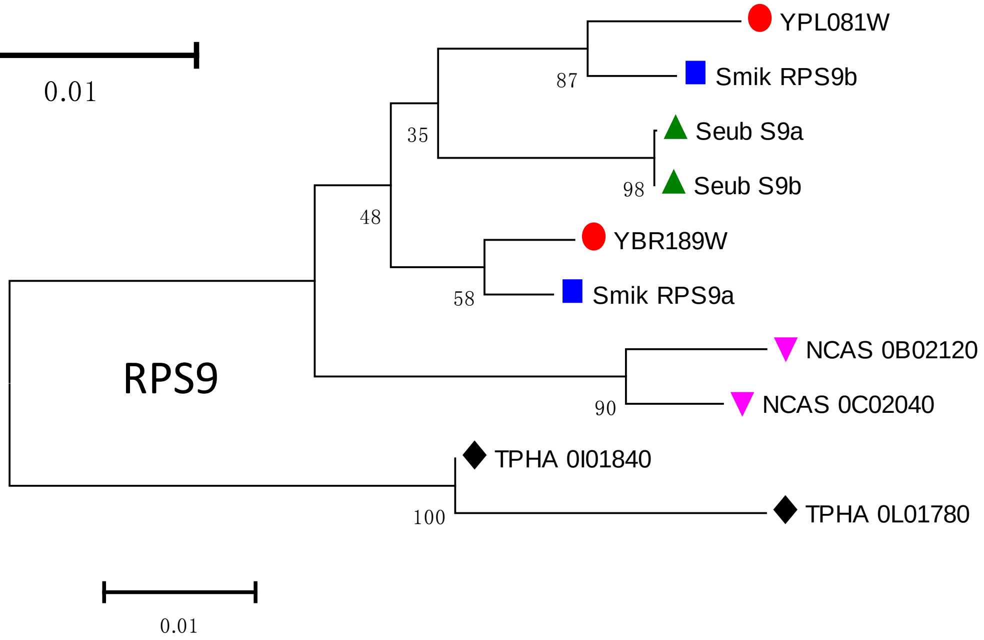


RPS9

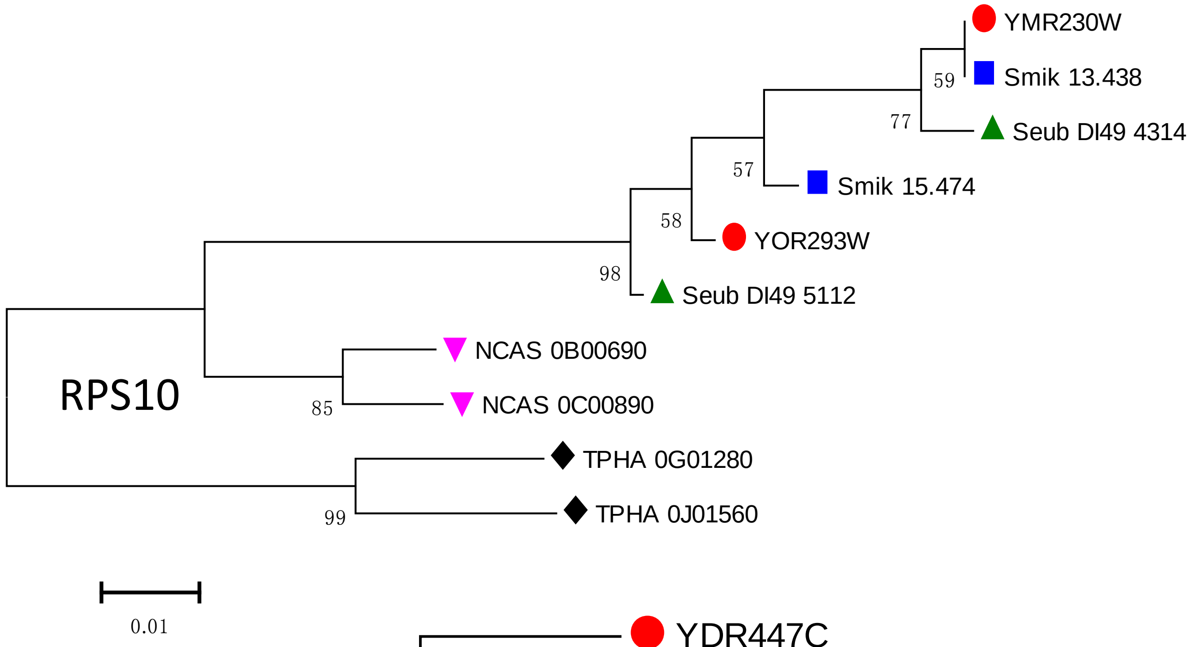
YMR230W
59
Smik 13.438
77
Seub DI49 4314
57
Smik 15.474
58
YOR293W
98
Seub DI49 5112
RPS10
NCAS 0B00690
85
NCAS 0C00890
TPHA 0G01280
99
TPHA 0J01560
0.01

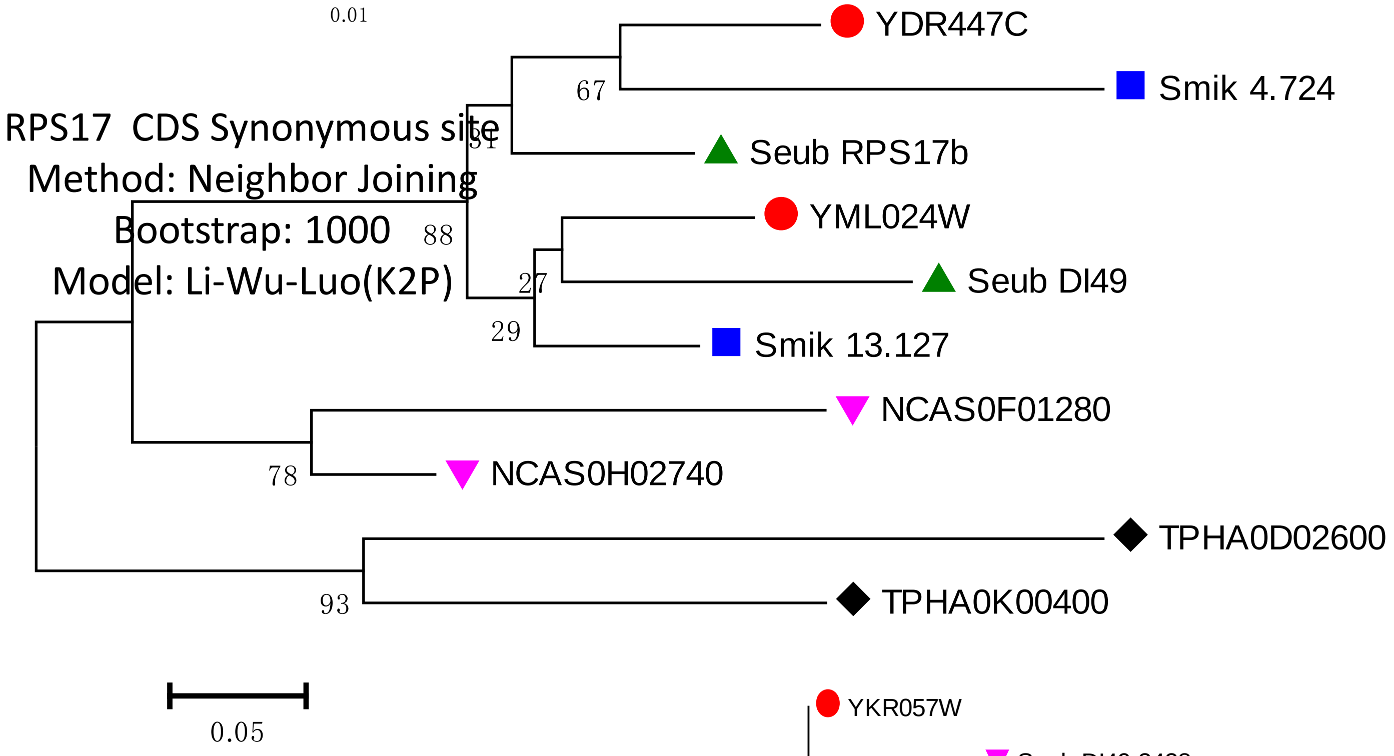
RPS17 CDS Synonymous site
Method: Neighbor Joining
Bootstrap: 1000
Model: Li-Wu-Luo(K2P)
YDR447C
67
Smik 4.724
51
Seub RPS17b
88
YML024W
27
Seub DI49
29
Smik 13.127
NCAS0F01280
78
NCAS0H02740
TPHA0D02600
93
TPHA0K00400
0.05

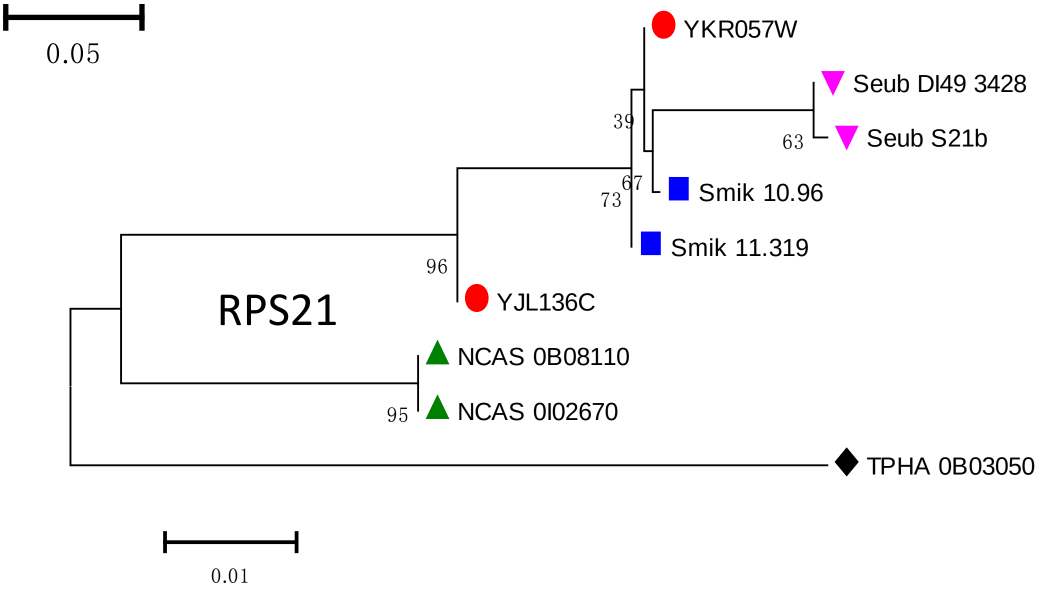
YKR057W
Seub DI49 3428
39
63
Seub S21b
67
73
Smik 10.96
Smik 11.319
96
RPS21
YJL136C
NCAS 0B08110
95
NCAS 0I02670
TPHA 0B03050
0.01

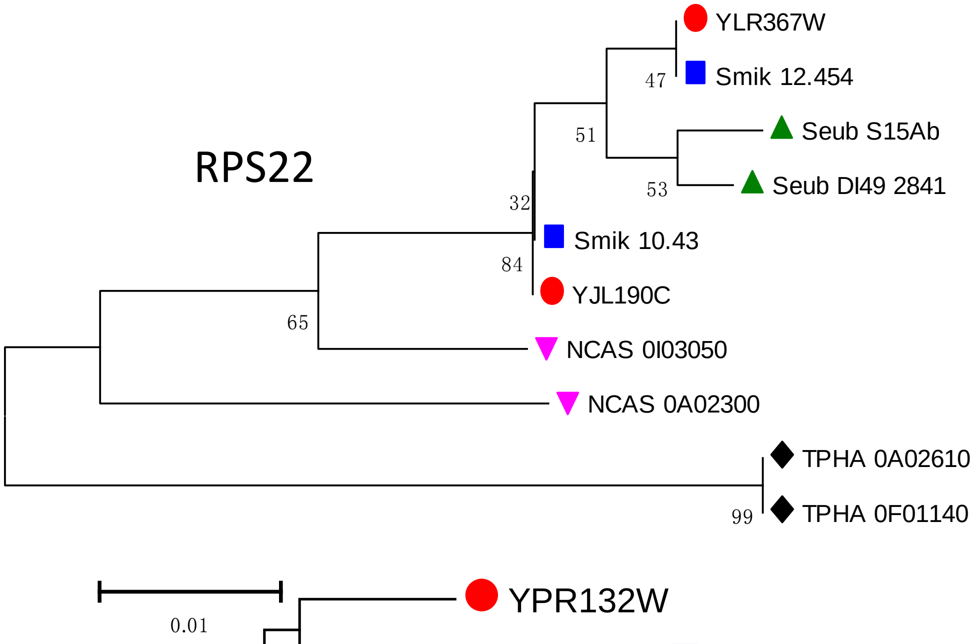

RPS22
YLR367W
Smik 12.454
Seub S15Ab
Seub DI49 2841
Smik 10.43
YJL190C
NCAS 0I03050
NCAS 0A02300
TPHA 0A02610
TPHA 0F01140
47
51
53
32
84
65
99
0.01


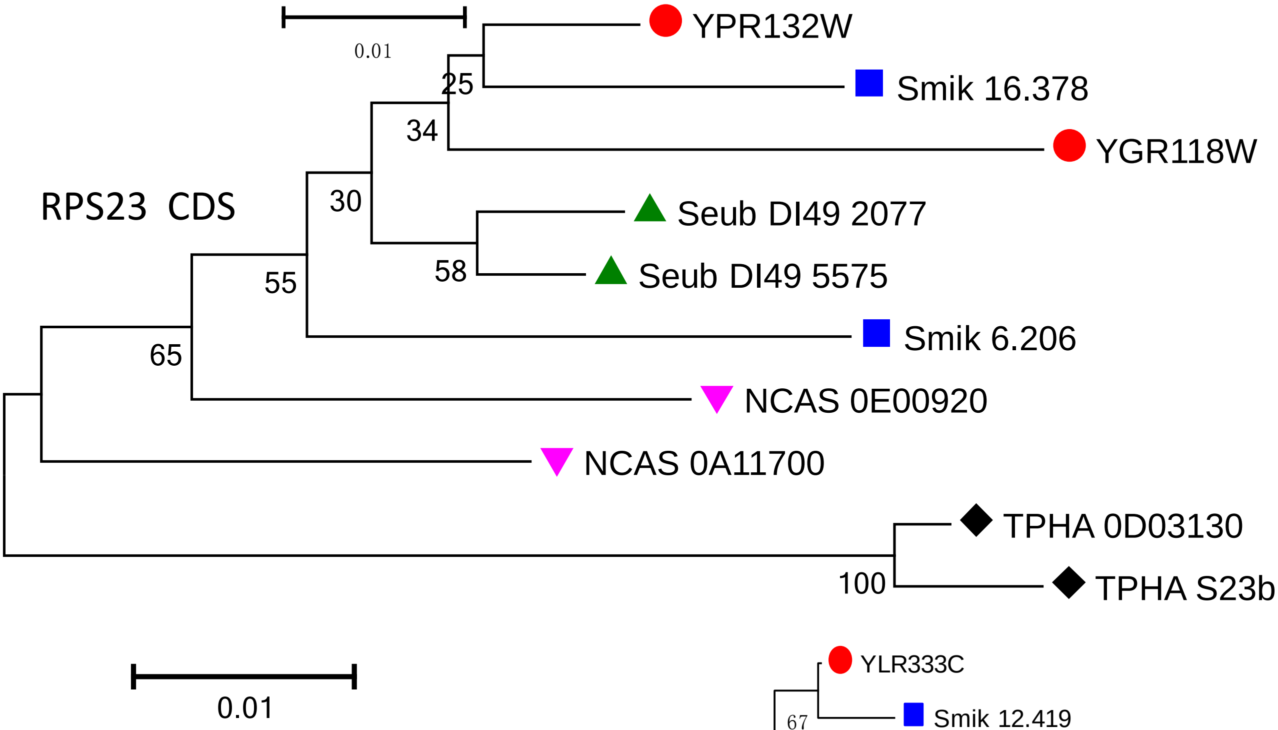

RPS23 CDS
YPR132W
Smik 16.378
YGR118W
Seub DI49 2077
Seub DI49 5575
Smik 6.206
NCAS 0E00920
NCAS 0A11700
TPHA 0D03130
TPHA S23b
25
34
30
58
55
65
100
0.01


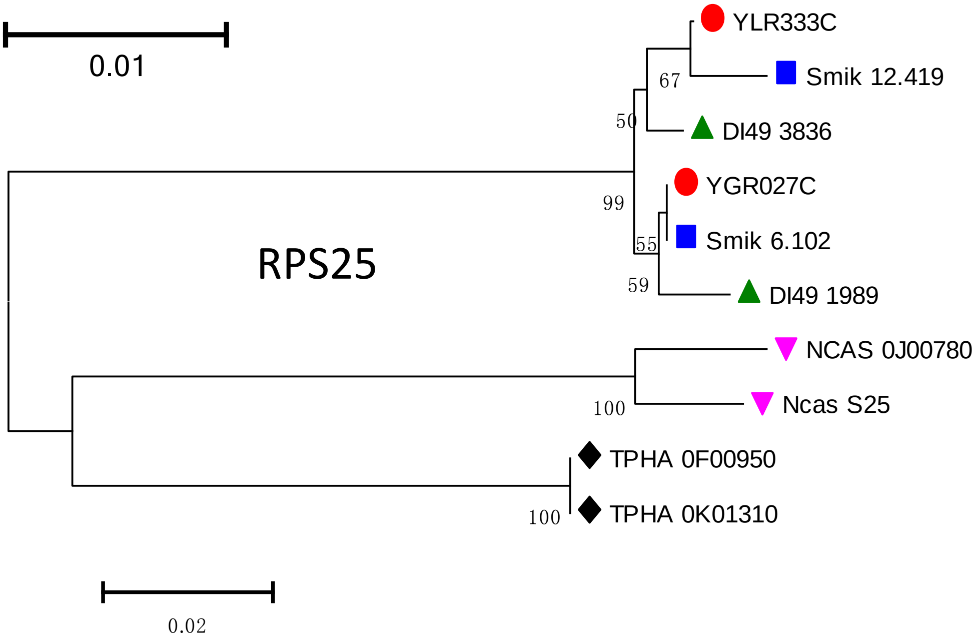

RPS25
YLR333C
Smik 12.419
DI49 3836
YGR027C
Smik 6.102
DI49 1989
NCAS 0J00780
Ncas S25
TPHA 0F00950
TPHA 0K01310
67
50
99
55
59
100
100
0.02

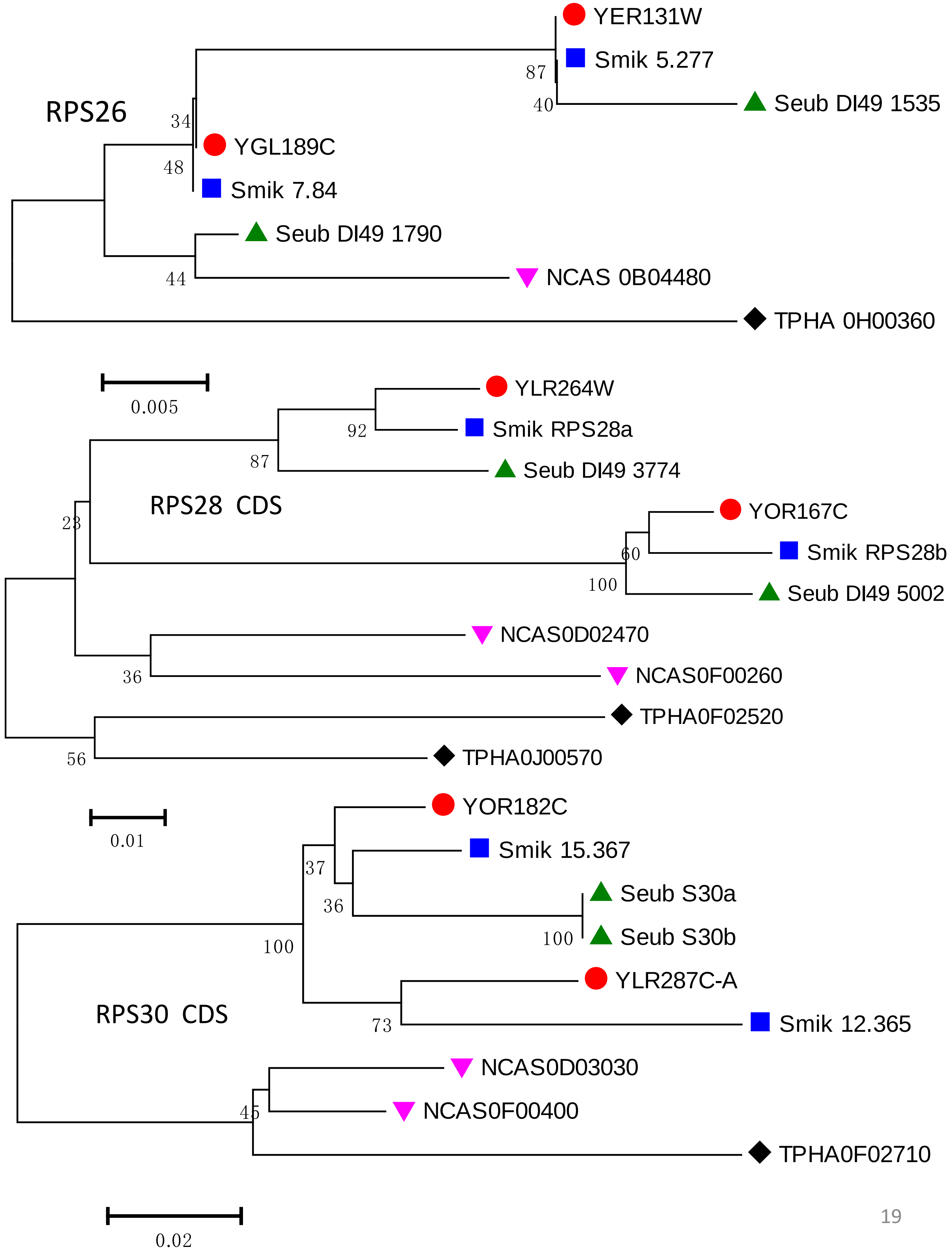
RPS26
YER131W
Smik 5.277
Seub DI49 1535
87
40
34
48
YGL189C
Smik 7.84
Seub DI49 1790
44
NCAS 0B04480
TPHA 0H00360
0.005
RPS28 CDS
YLR264W
Smik RPS28a
Seub DI49 3774
92
87
23
YOR167C
Smik RPS28b
Seub DI49 5002
60
100
NCAS0D02470
NCAS0F00260
36
TPHA0F02520
TPHA0J00570
56
0.01
RPS30 CDS
YOR182C
Smik 15.367
Seub S30a
Seub S30b
37
36
100
100
YLR287C-A
Smik 12.365
73
NCAS0D03030
NCAS0F00400
45
TPHA0F02710
0.02

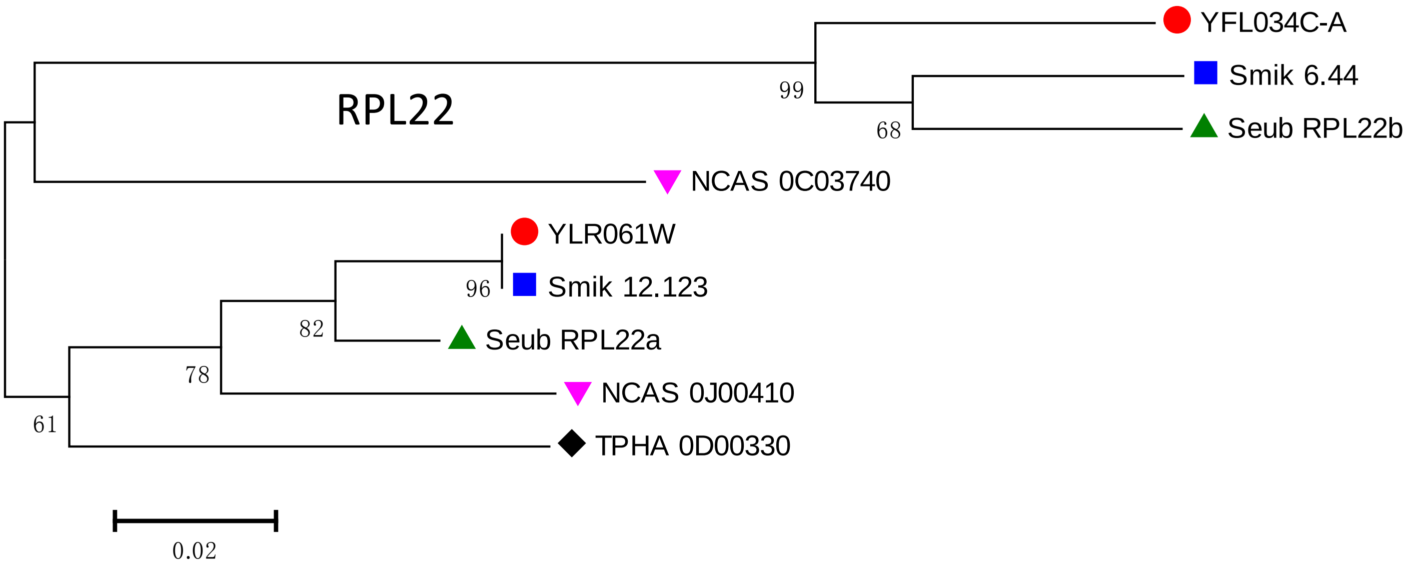
RPL22
YFL034C-A
Smik 6.44
Seub RPL22b
NCAS 0C03740
YLR061W
Smik 12.123
Seub RPL22a
NCAS 0J00410
TPHA 0D00330
99
68
96
82
78
61
0.02

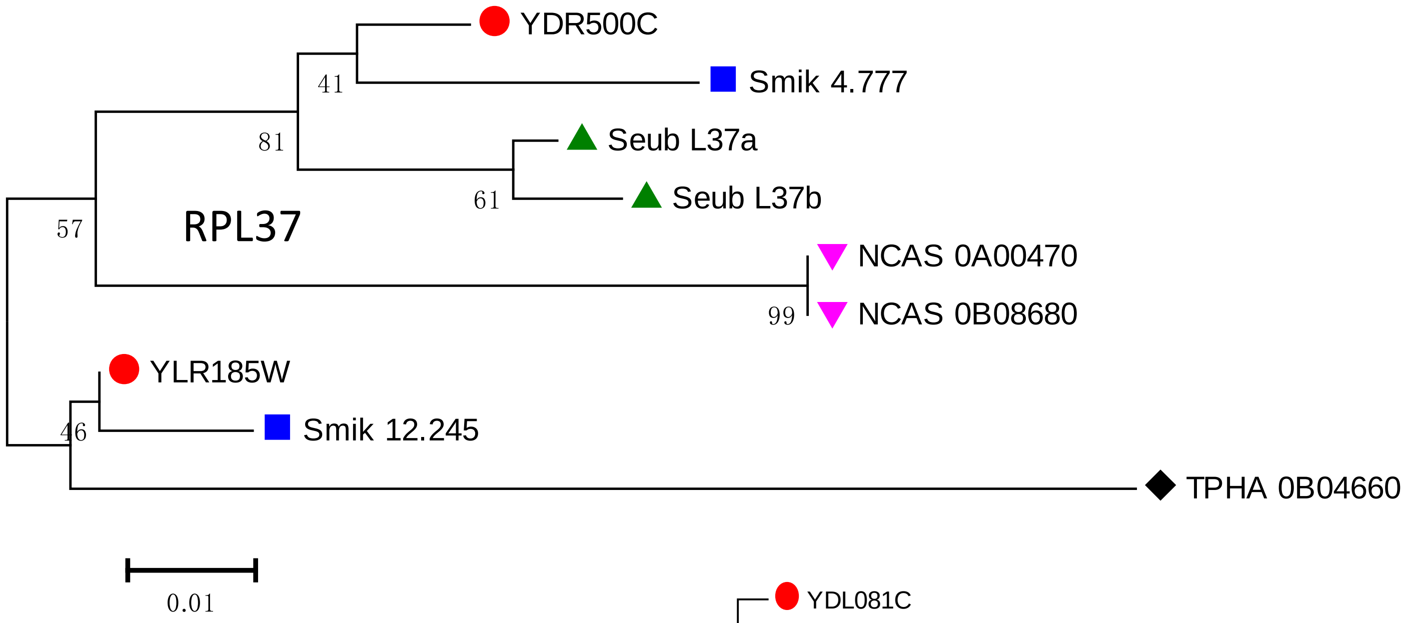
RPL37
YDR500C
Smik 4.777
Seub L37a
Seub L37b
NCAS 0A00470
NCAS 0B08680
YLR185W
Smik 12.245
TPHA 0B04660
41
81
61
57
99
46
0.01

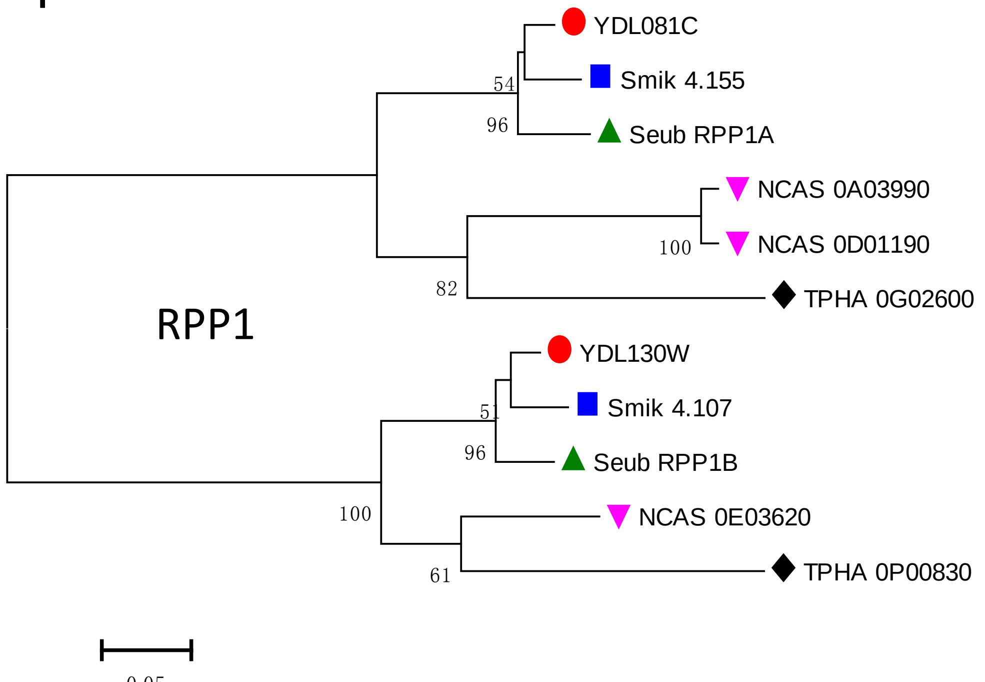
RPP1
YDL081C
Smik 4.155
Seub RPP1A
NCAS 0A03990
NCAS 0D01190
TPHA 0G02600
YDL130W
Smik 4.107
Seub RPP1B
NCAS 0E03620
TPHA 0P00830
54
96
100
82
51
96
100
61

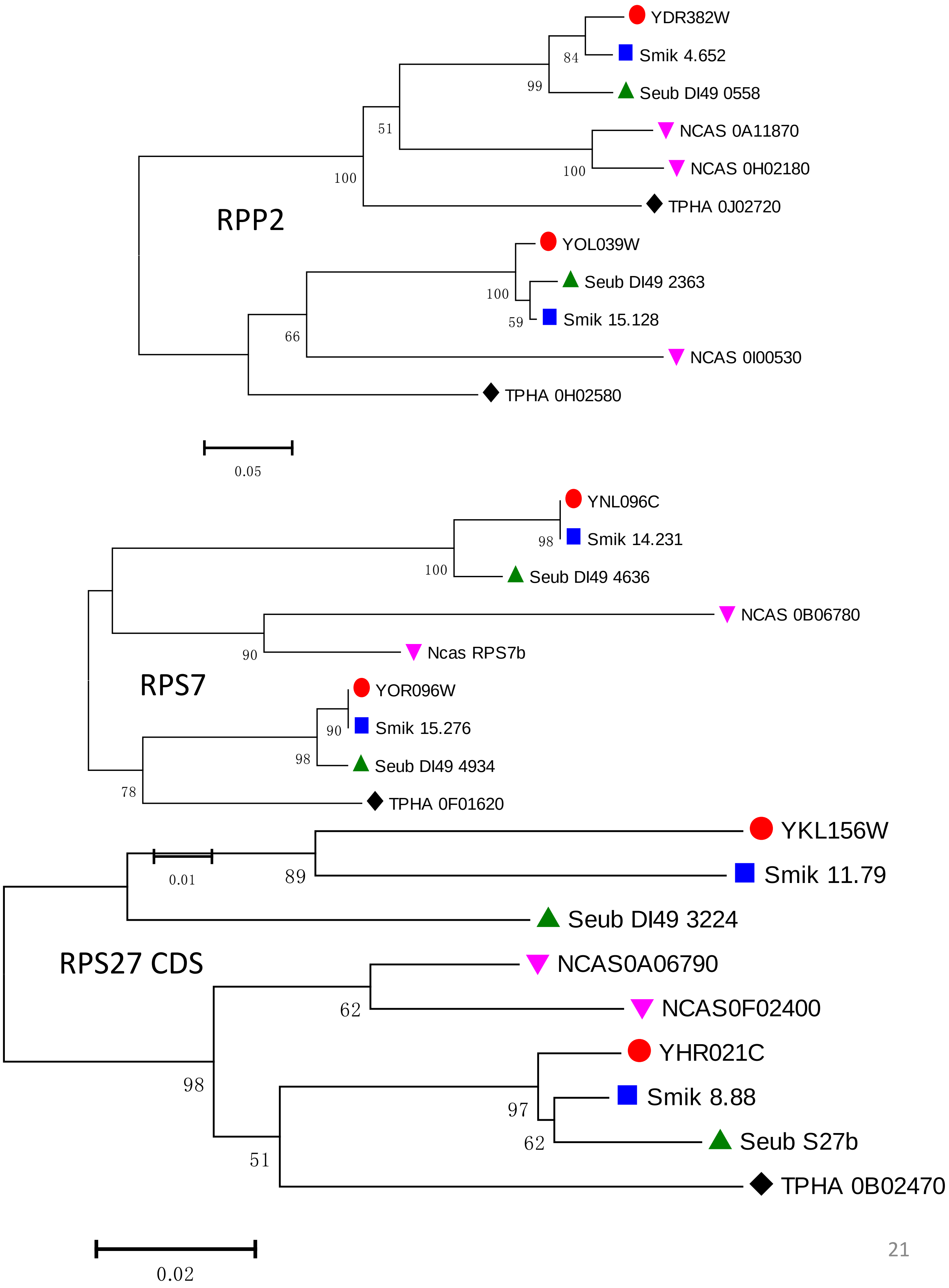
RPP2
YDR382W
Smik 4.652
Seub DI49 0558
NCAS 0A11870
NCAS 0H02180
TPHA 0J02720
YOL039W
Seub DI49 2363
Smik 15.128
NCAS 0I00530
TPHA 0H02580
84
99
51
100
100
100
59
66
0.05
RPS7
YNL096C
Smik 14.231
Seub DI49 4636
NCAS 0B06780
Ncas RPS7b
YOR096W
Smik 15.276
Seub DI49 4934
TPHA 0F01620
98
100
90
90
98
78
RPS27 CDS
YKL156W
Smik 11.79
Seub DI49 3224
NCAS0A06790
NCAS0F02400
YHR021C
Smik 8.88
Seub S27b
TPHA 0B02470
0.01
89
62
98
97
62
51
0.02

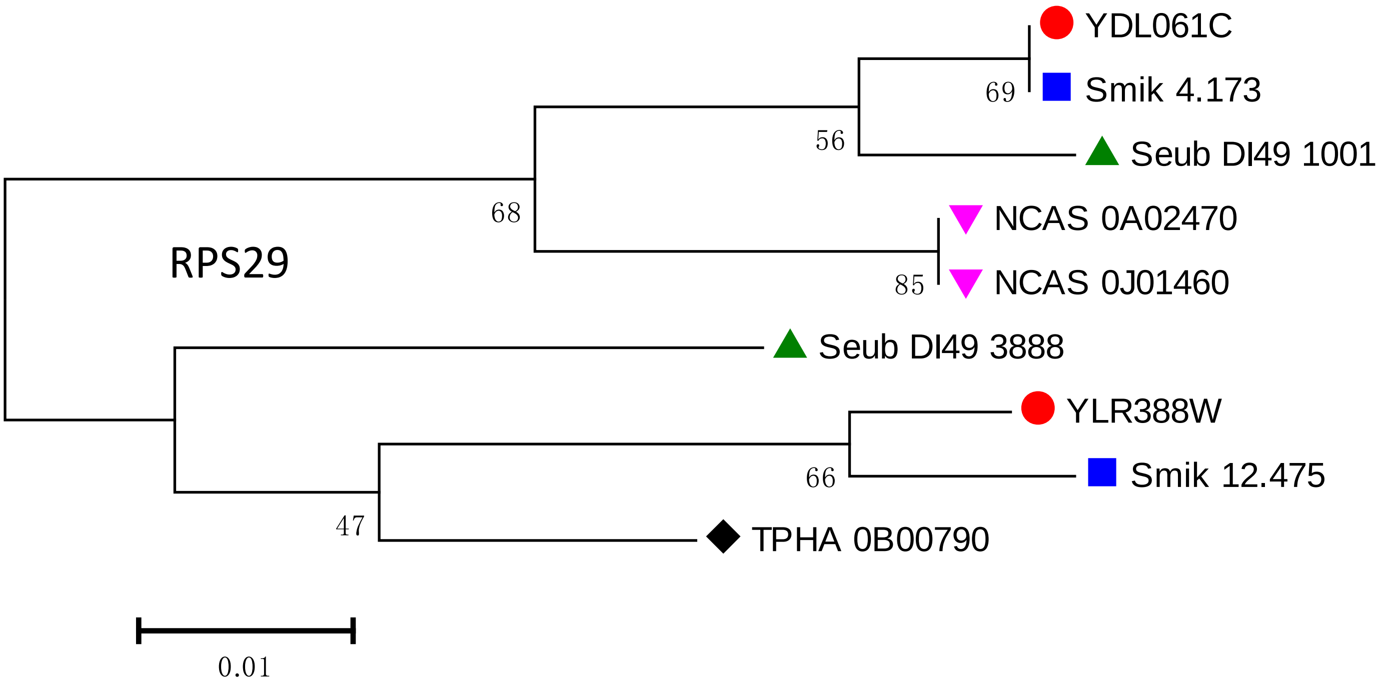
YDL061C
Smik 4.173
Seub DI49 1001
NCAS 0A02470
NCAS 0J01460
Seub DI49 3888
YLR388W
Smik 12.475
TPHA 0B00790
RPS29
69
56
68
85
66
47
0.01
