## Supplemental File 3 for "Parallel Concerted Evolution of Ribosomal Protein Genes in Fungi and Its Adaptive Significance"

**Supplementary File 3: Phylogenetic trees of RP families in the four fission yeast species.**

| Gene family | Category | Page | Gene family | Category | Page |
|---|---|---|---|---|---|
| RPL1 | A | 2 | RPS22 | A | 15 |
| RPL2 | A | 2 | RPS25 | A | 15 |
| RPL3 | A | 2 | RPS30 | A | 15 |
| RPL4 | A | 3 | RPS31 | A | 16 |
| RPL5 | A | 3 | RPP1 | A | 16 |
| RPL9 | A | 3 | RPP2 | A | 16 |
| RPL10 | A | 4 | RPL7 | B | 17 |
| RPL11 | A | 4 | RPL33 | B | 17 |
| RPL12 | A | 4 | RPL34 | B | 17 |
| RPL15 | A | 5 | RPL38 | B | 18 |
| RPL16 | A | 5 | RPL41 | B | 18 |
| RPL17 | A | 5 | RPS0 | B | 18 |
| RPL18 | A | 6 | RPS1 | B | 19 |
| RPL19 | A | 6 | RPS17 | B | 19 |
| RPL20 | A | 6 | RPS26 | B | 19 |
| RPL21 | A | 7 | RPL30 | C | 20 |
| RPL23 | A | 7 | RPS5 | C | 20 |
| RPL24 | A | 7 | RPS12 | C | 20 |
| RPL25 | A | 8 | RPS28 | C | 21 |
| RPL27 | A | 8 | | | |
| RPL28 | A | 8 | | | |
| RPL32 | A | 9 | | | |
| RPL36 | A | 9 | | | |
| RPL37 | A | 9 | | | |
| RPL40 | A | 10 | | | |
| RPL43 | A | 10 | | | |
| RPS4 | A | 10 | | | |
| RPS6 | A | 11 | | | |
| RPS8 | A | 11 | | | |
| RPS9 | A | 11 | | | |
| RPS10 | A | 12 | | | |
| RPS11 | A | 12 | | | |
| RPS14 | A | 12 | | | |
| RPS15 | A | 13 | | | |
| RPS16 | A | 13 | | | |
| RPS18 | A | 13 | | | |
| RPS19 | A | 14 | | | |
| RPS22 | A | 14 | | | |
| RPS23 | A | 14 | | | |

*The phylogenetic trees were built based on amino acid sequences of four WGD budding yeast species, unless specially specified. Each tree was reconstructed by Neighbor-Joining method using Poisson correction with 1000 Bootstrap replicates. For some trees, if the phylogenetic tree topology is not resolved due to high degree of sequence similarity, we used their CDS sequences to build their phylogenetic tree.

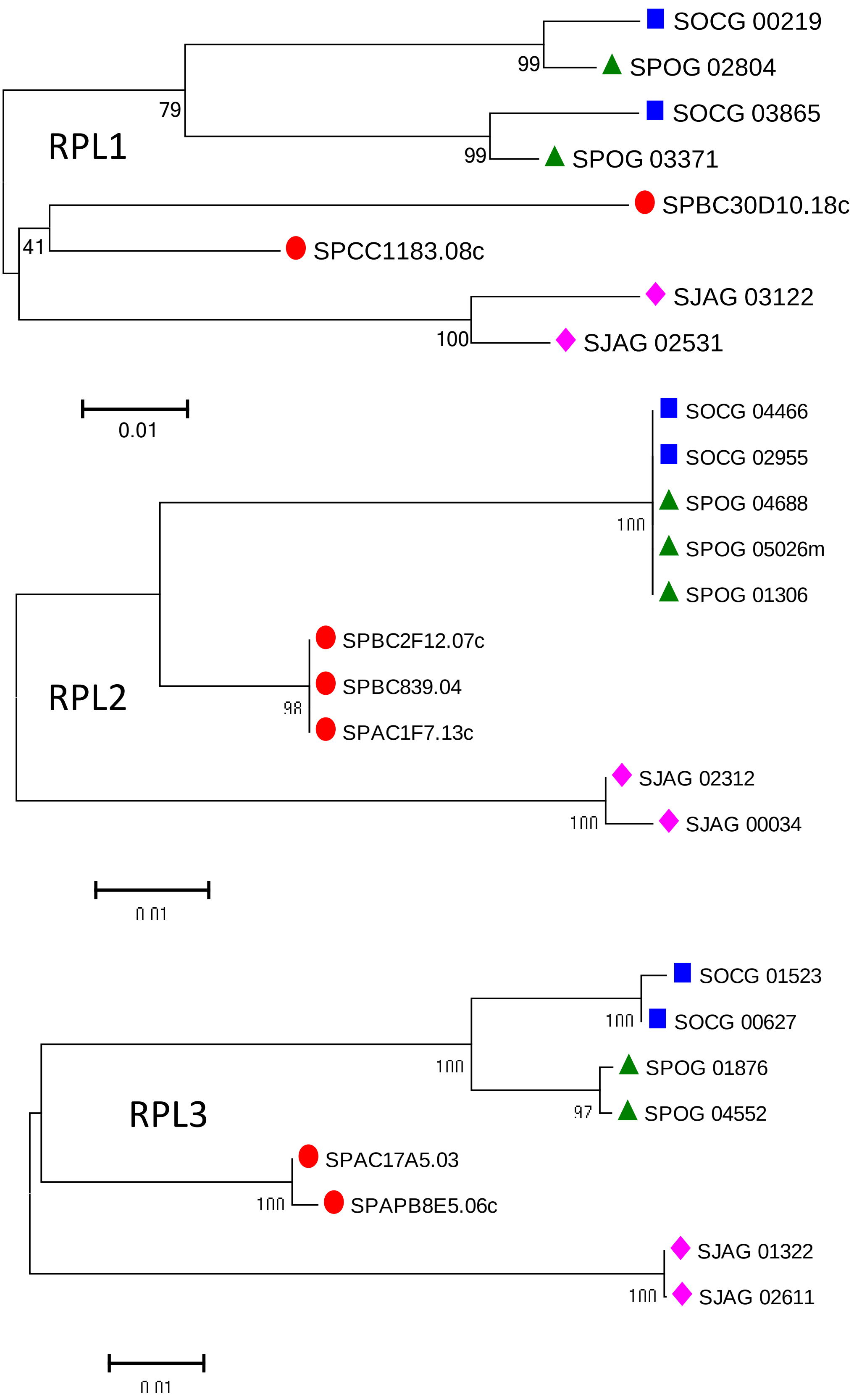
RPL1
SOCG 00219
SPOG 02804
SOCG 03865
SPOG 03371
SPBC30D10.18c
SPCC1183.08c
SJAG 03122
SJAG 02531
99
79
99
41
100
0.01
RPL2
SOCG 04466
SOCG 02955
SPOG 04688
SPOG 05026m
SPOG 01306
SPBC2F12.07c
SPBC839.04
SPAC1F7.13c
SJAG 02312
SJAG 00034
100
98
100
0.01
RPL3
SOCG 01523
SOCG 00627
SPOG 01876
SPOG 04552
SPAC17A5.03
SPAPB8E5.06c
SJAG 01322
SJAG 02611
100
100
97
100
100
0.01

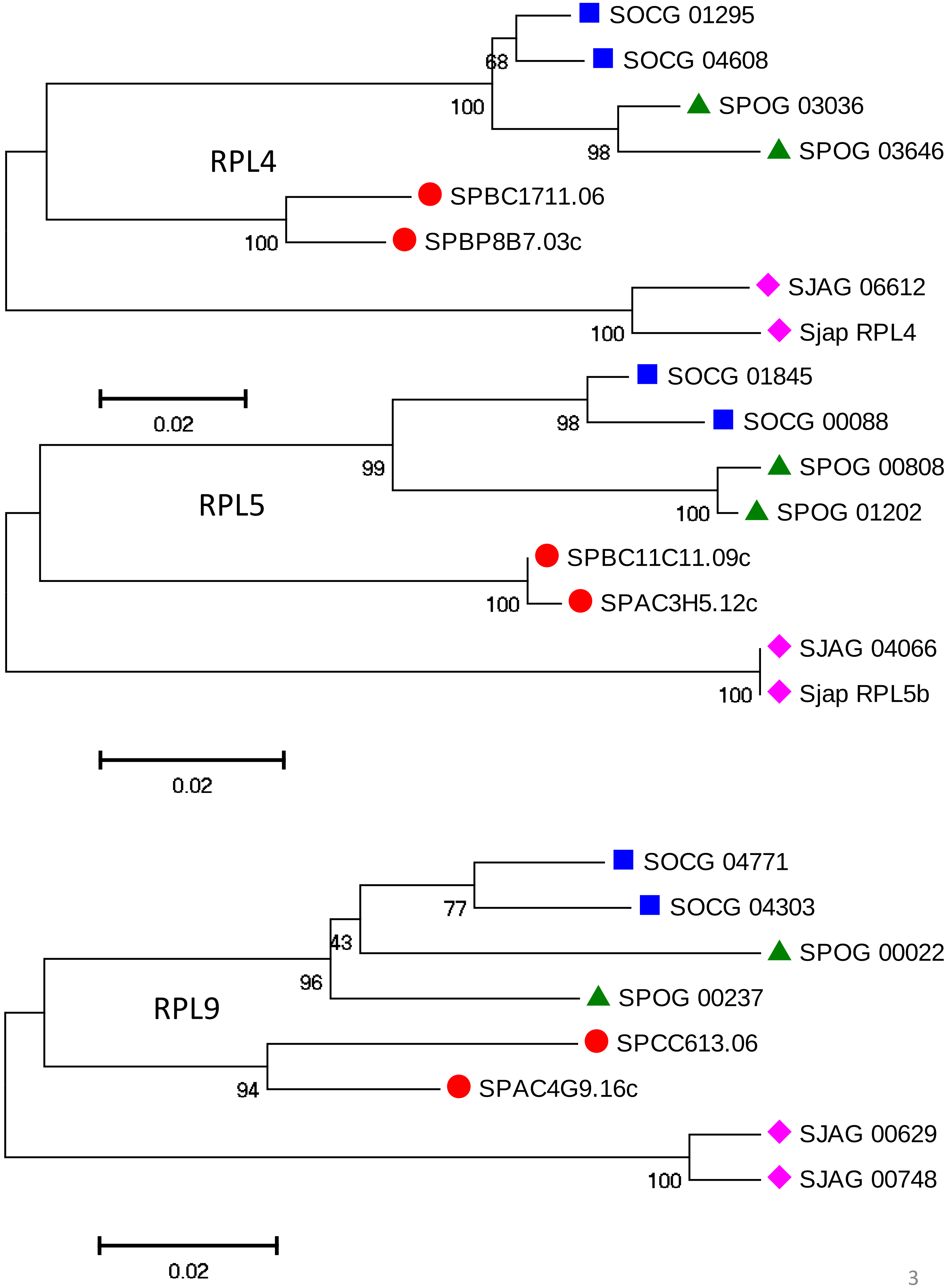

SOCG 01295
68
SOCG 04608
100
SPOG 03036
RPL4
98
SPOG 03646
SPBC1711.06
100
SPBP8B7.03c
SJAG 06612
100
Sjap RPL4
SOCG 01845
0.02
98
SOCG 00088
99
SPOG 00808
RPL5
100
SPOG 01202
SPBC11C11.09c
100
SPAC3H5.12c
SJAG 04066
100
Sjap RPL5b
0.02
SOCG 04771
77
SOCG 04303
43
SPOG 00022
96
SPOG 00237
RPL9
SPCC613.06
94
SPAC4G9.16c
SJAG 00629
100
SJAG 00748
0.02

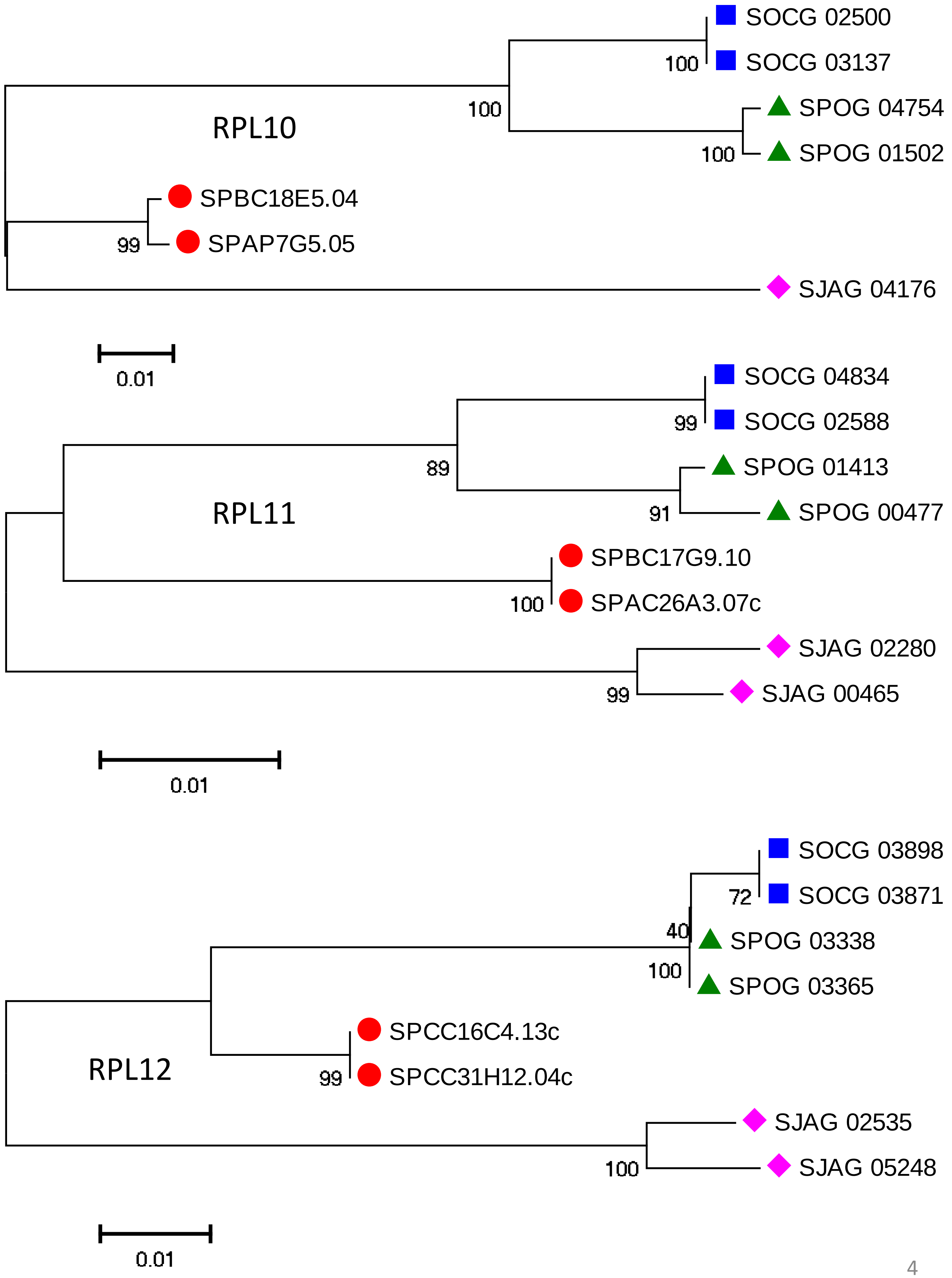
RPL10
SOCG 02500
SOCG 03137
SPOG 04754
SPOG 01502
SPBC18E5.04
SPAP7G5.05
SJAG 04176
100
100
100
99
0.01
RPL11
SOCG 04834
SOCG 02588
SPOG 01413
SPOG 00477
SPBC17G9.10
SPAC26A3.07c
SJAG 02280
SJAG 00465
99
89
91
100
99
0.01
RPL12
SOCG 03898
SOCG 03871
SPOG 03338
SPOG 03365
SPCC16C4.13c
SPCC31H12.04c
SJAG 02535
SJAG 05248
72
40
100
99
100
0.01

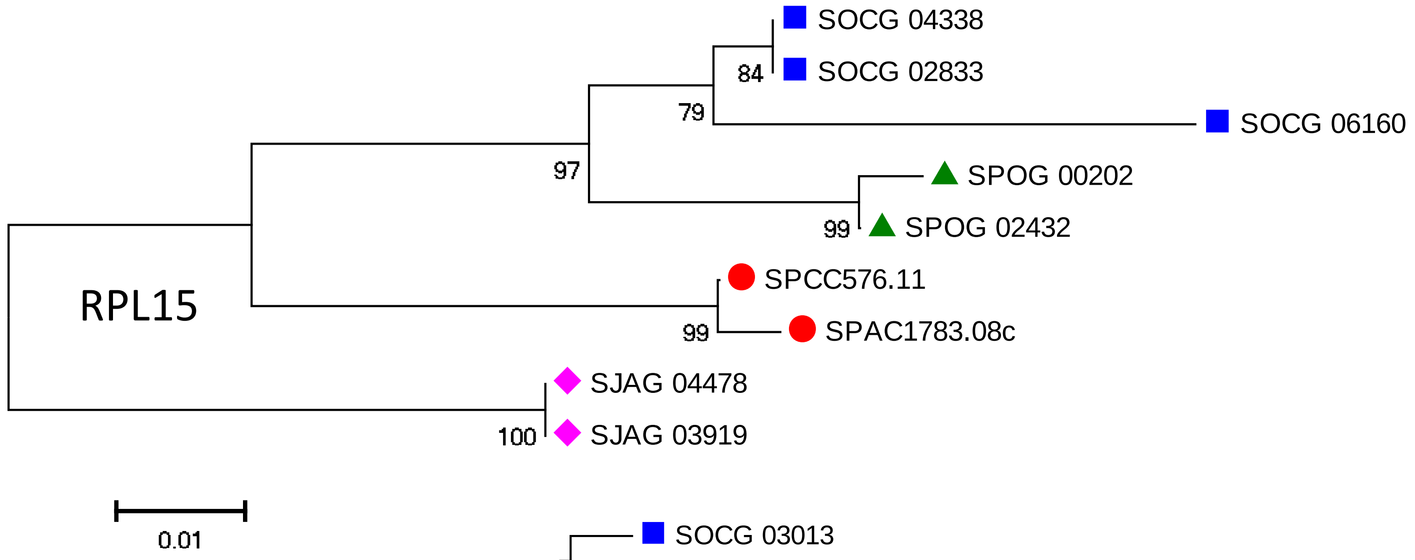

SOCG 04338
SOCG 02833
SOCG 06160
SPOG 00202
SPOG 02432
SPCC576.11
SPAC1783.08c
SJAG 04478
SJAG 03919
RPL15
84
79
97
99
99
100
0.01


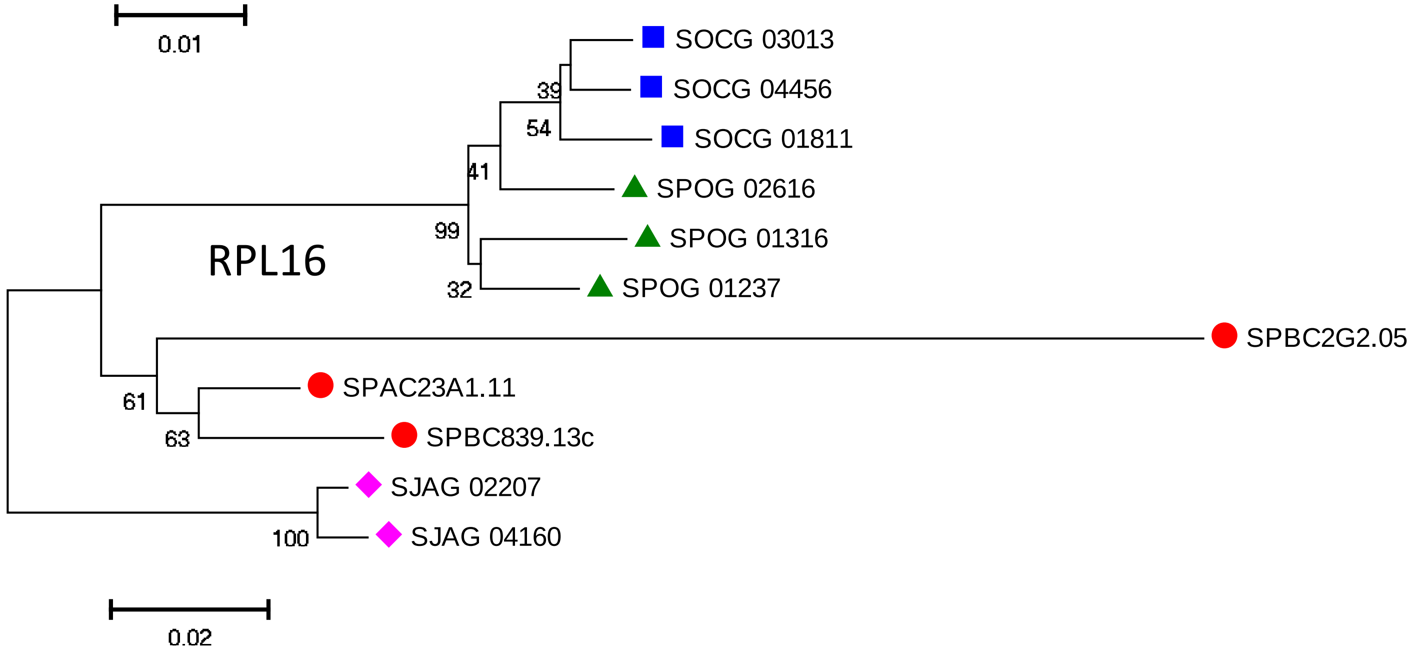

SOCG 03013
SOCG 04456
SOCG 01811
SPOG 02616
SPOG 01316
SPOG 01237
SPBC2G2.05
SPAC23A1.11
SPBC839.13c
SJAG 02207
SJAG 04160
RPL16
39
54
41
99
32
61
63
100
0.02


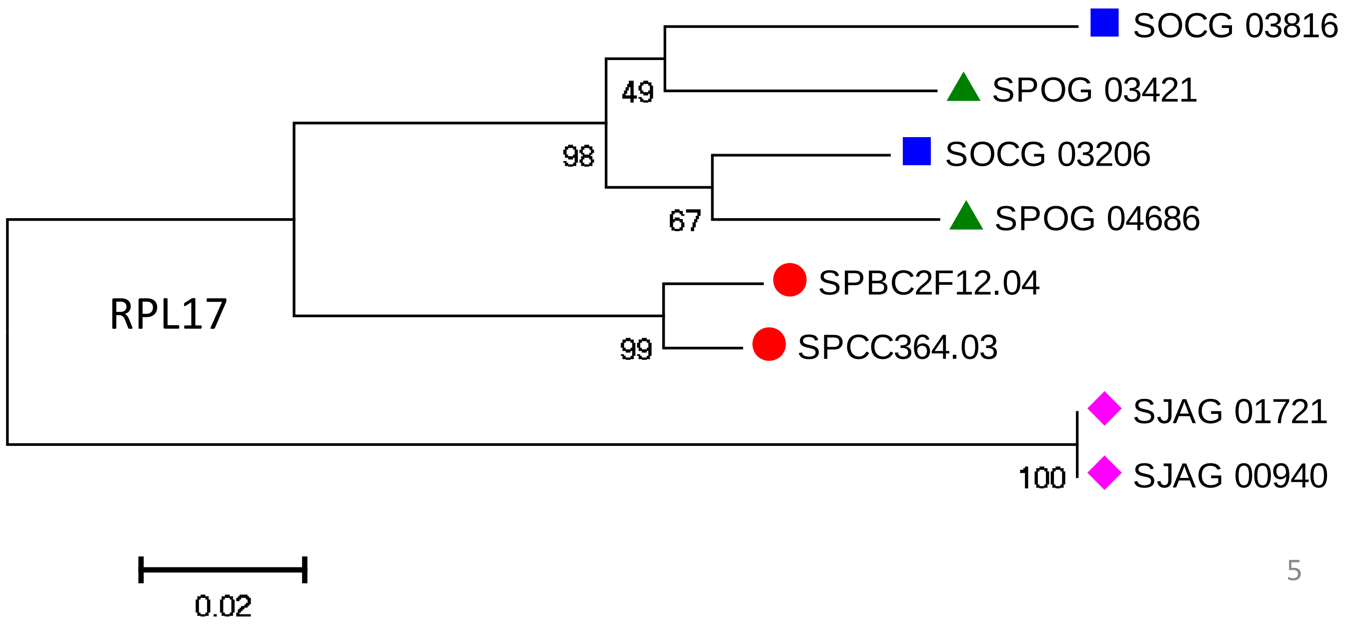

SOCG 03816
SPOG 03421
SOCG 03206
SPOG 04686
SPBC2F12.04
SPCC364.03
SJAG 01721
SJAG 00940
RPL17
49
98
67
99
100
0.02

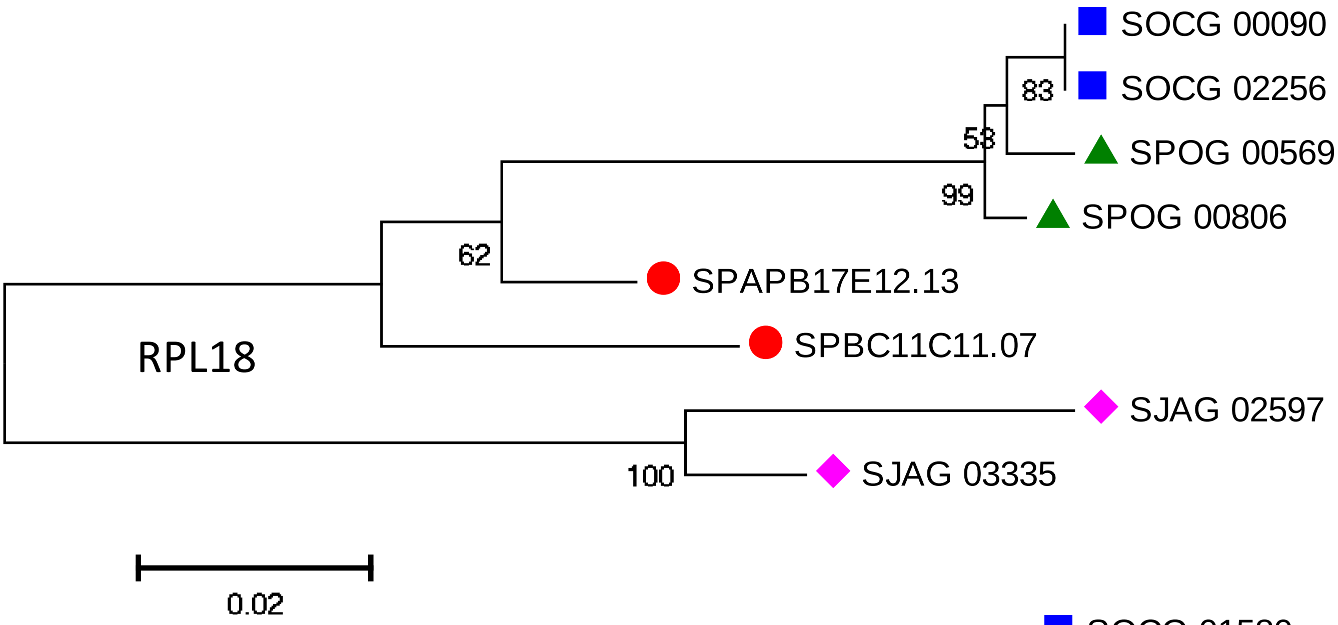
SOCG 00090
SOCG 02256
SPOG 00569
SPOG 00806
SPAPB17E12.13
SPBC11C11.07
SJAG 02597
SJAG 03335
RPL18
83
58
99
62
100
0.02

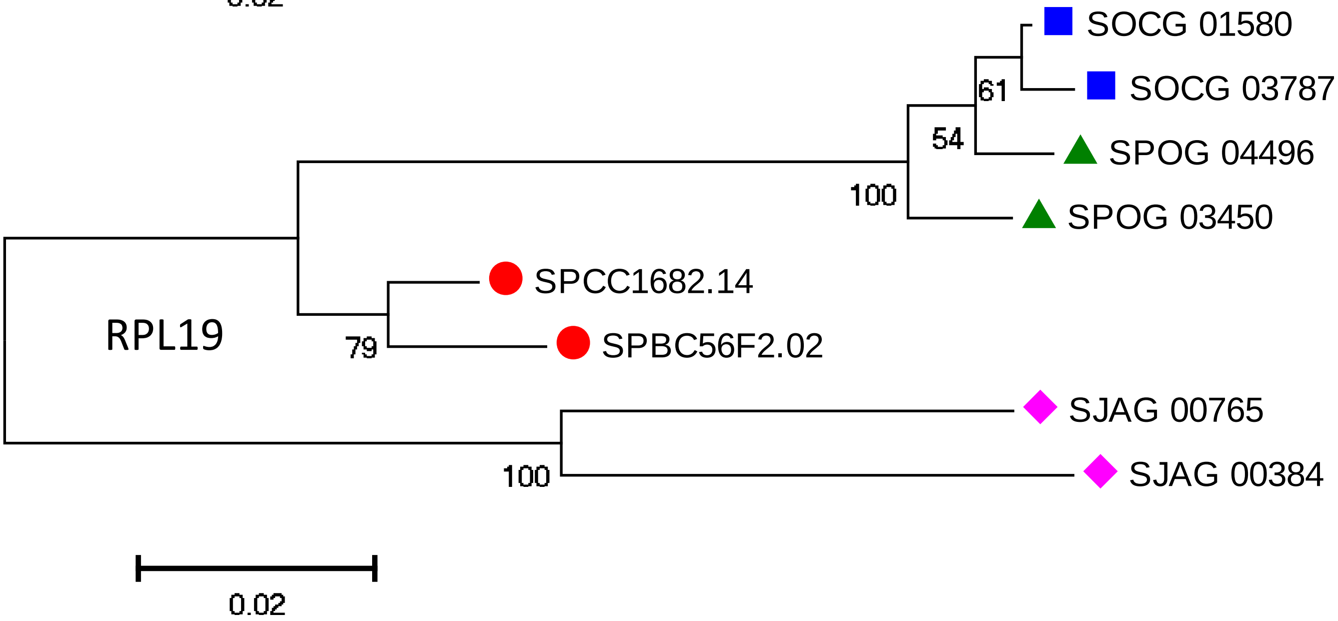
SOCG 01580
SOCG 03787
SPOG 04496
SPOG 03450
SPCC1682.14
SPBC56F2.02
SJAG 00765
SJAG 00384
RPL19
61
54
100
79
100
0.02

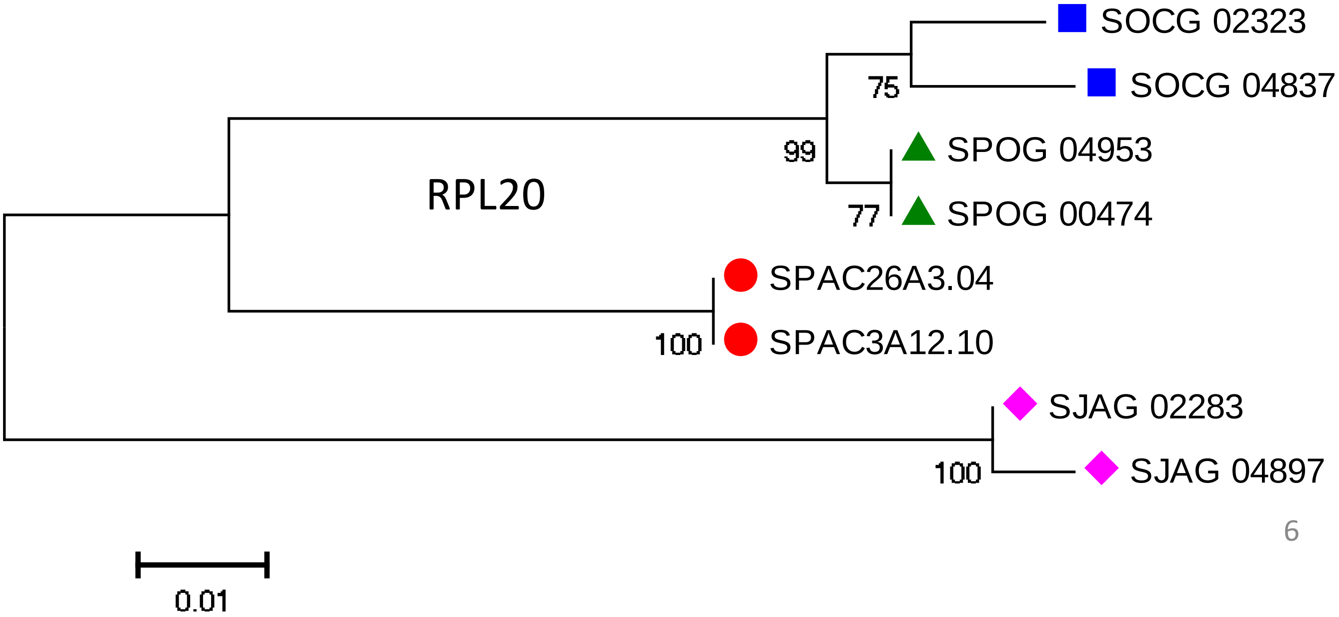
SOCG 02323
SOCG 04837
SPOG 04953
SPOG 00474
SPAC26A3.04
SPAC3A12.10
SJAG 02283
SJAG 04897
RPL20
75
99
77
100
100
0.01

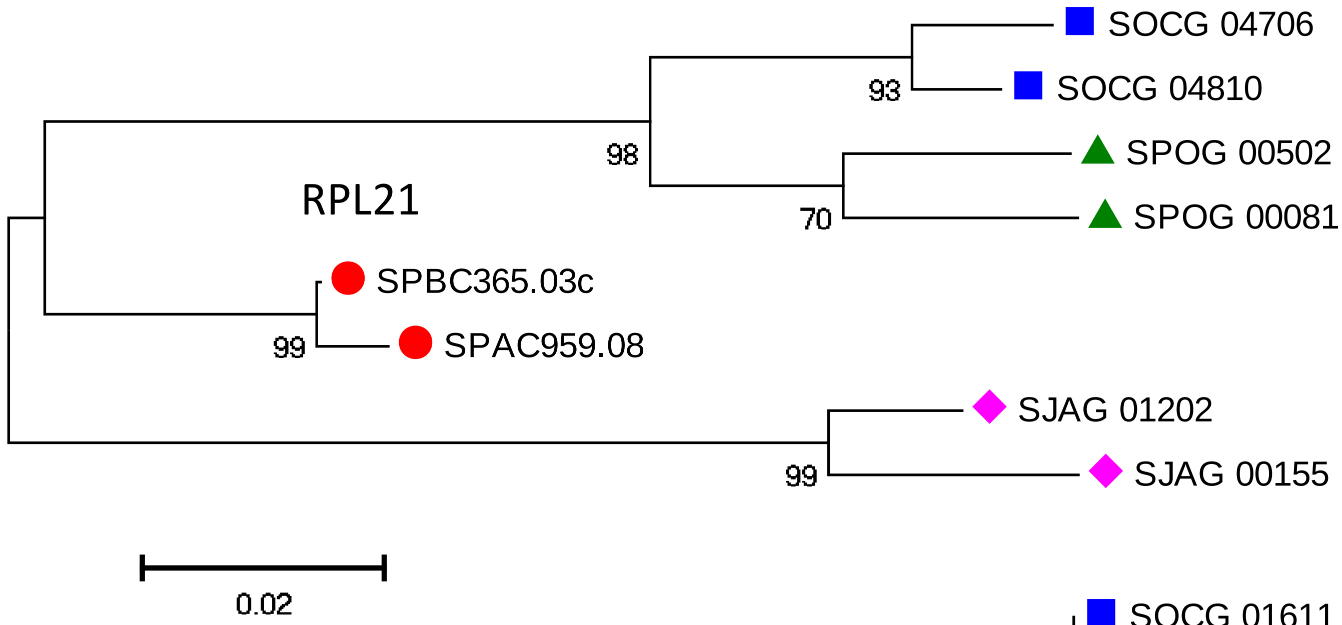
RPL21
SOCG 04706
SOCG 04810
SPOG 00502
SPOG 00081
SPBC365.03c
SPAC959.08
SJAG 01202
SJAG 00155
93
98
70
99
99
0.02

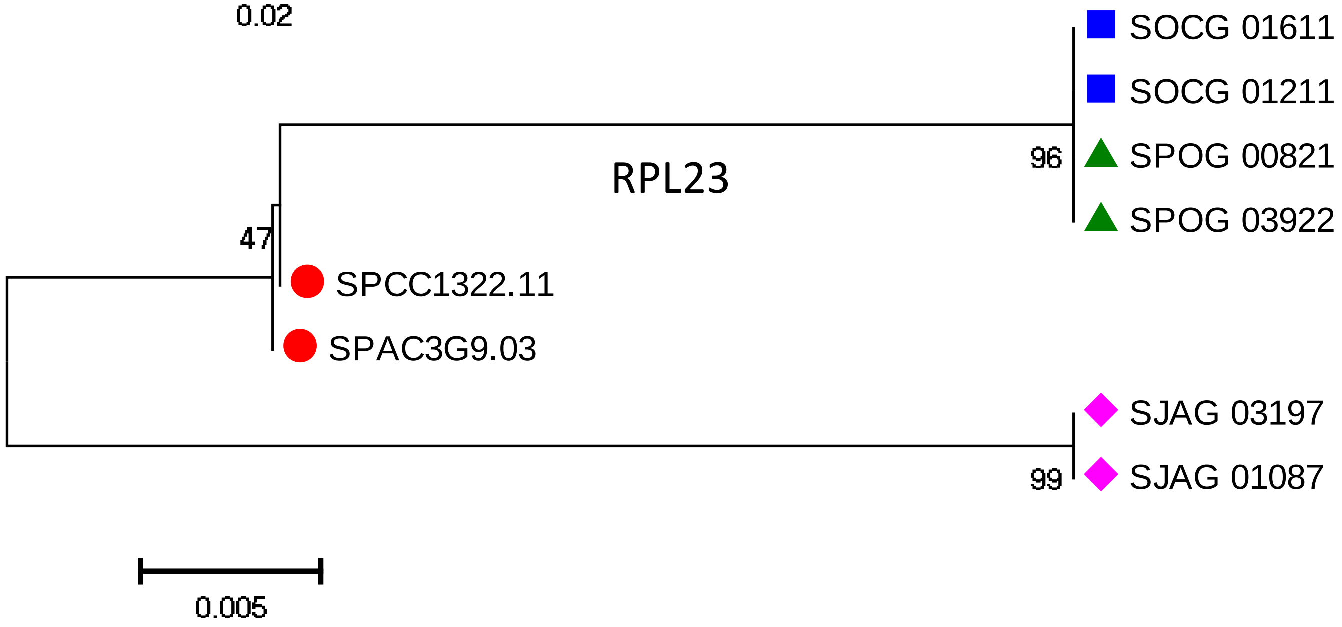
RPL23
SOCG 01611
SOCG 01211
SPOG 00821
SPOG 03922
SPCC1322.11
SPAC3G9.03
SJAG 03197
SJAG 01087
96
47
99
0.005

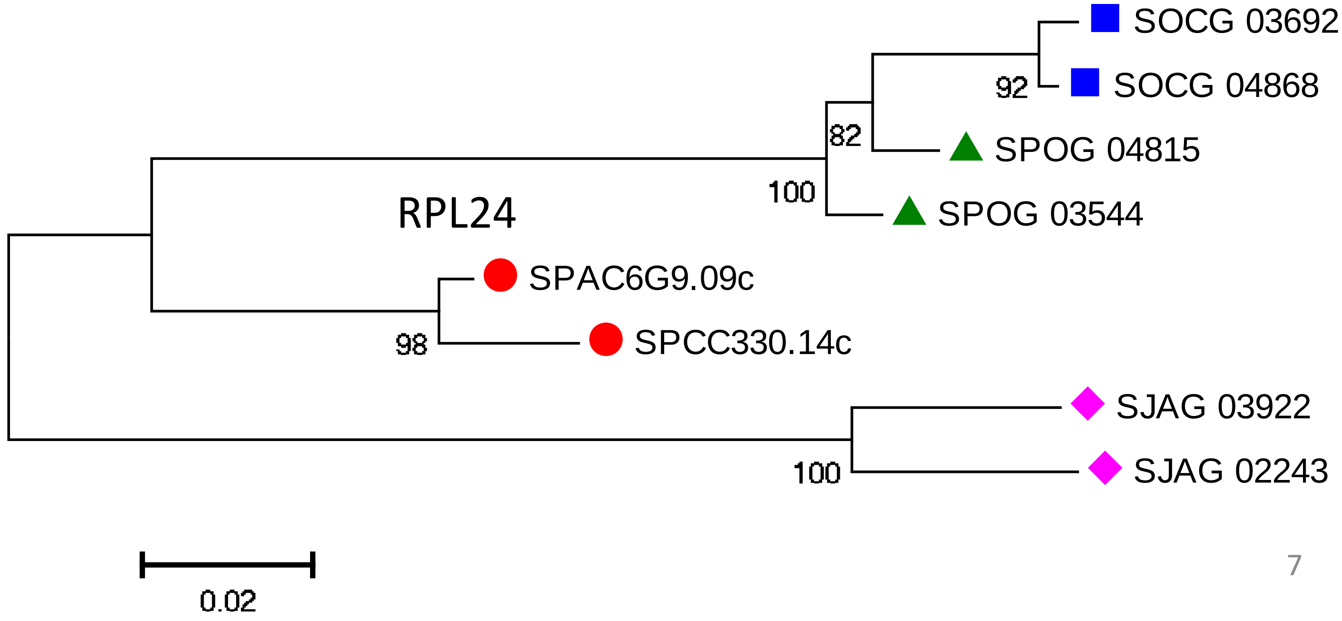
RPL24
SOCG 03692
SOCG 04868
SPOG 04815
SPOG 03544
SPAC6G9.09c
SPCC330.14c
SJAG 03922
SJAG 02243
92
82
100
98
100
0.02

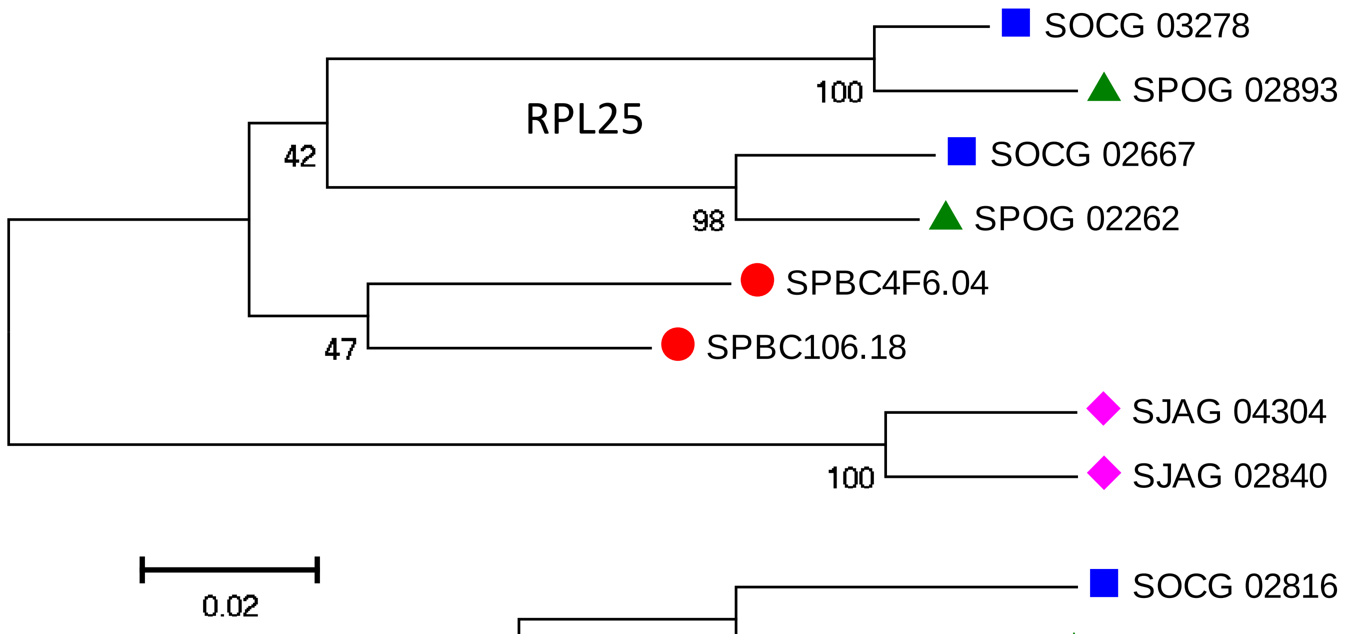
RPL25
SOCG 03278
SPOG 02893
100
SOCG 02667
SPOG 02262
98
42
SPBC4F6.04
SPBC106.18
47
SJAG 04304
SJAG 02840
100
0.02

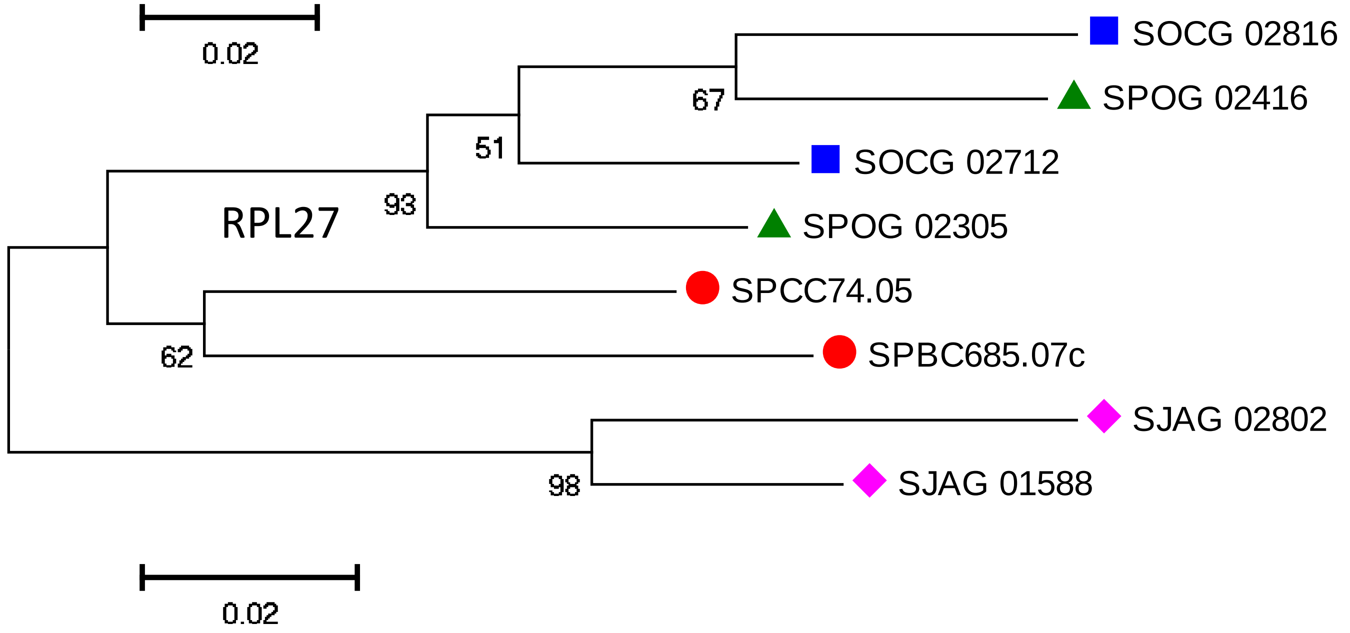
RPL27
SOCG 02816
SPOG 02416
67
SOCG 02712
51
SPOG 02305
93
SPCC74.05
SPBC685.07c
62
SJAG 02802
SJAG 01588
98
0.02

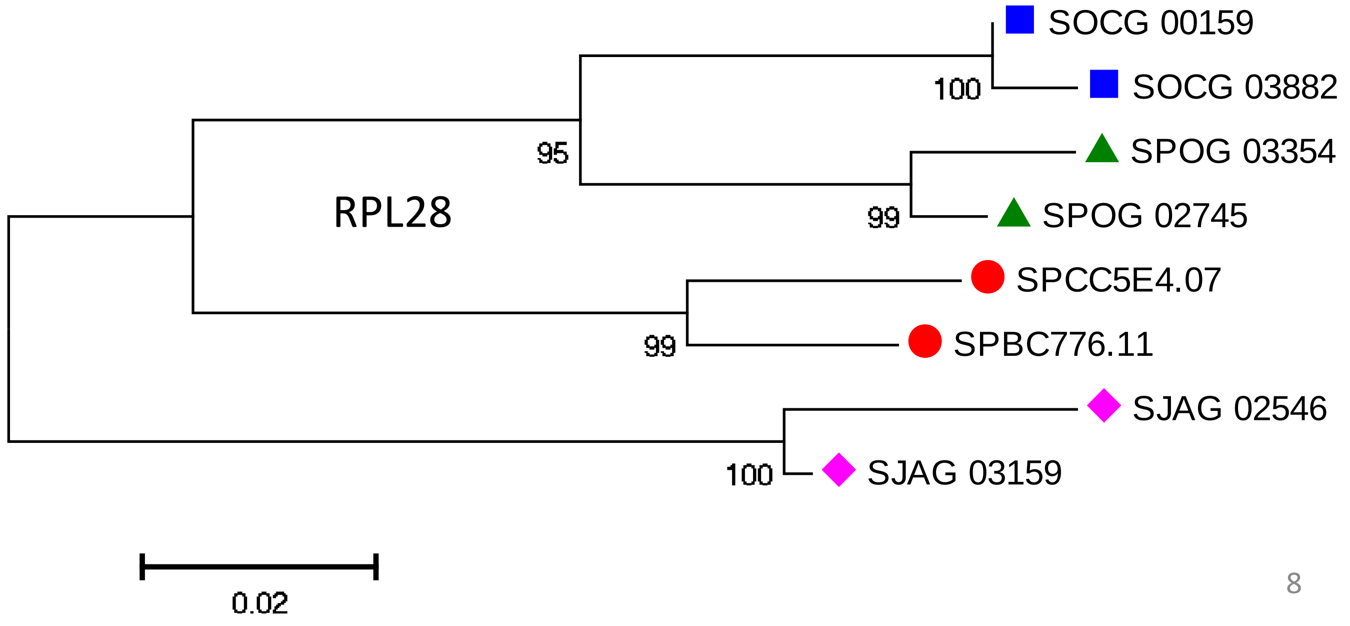
RPL28
SOCG 00159
SOCG 03882
100
SPOG 03354
SPOG 02745
99
95
SPCC5E4.07
SPBC776.11
99
SJAG 02546
SJAG 03159
100
0.02

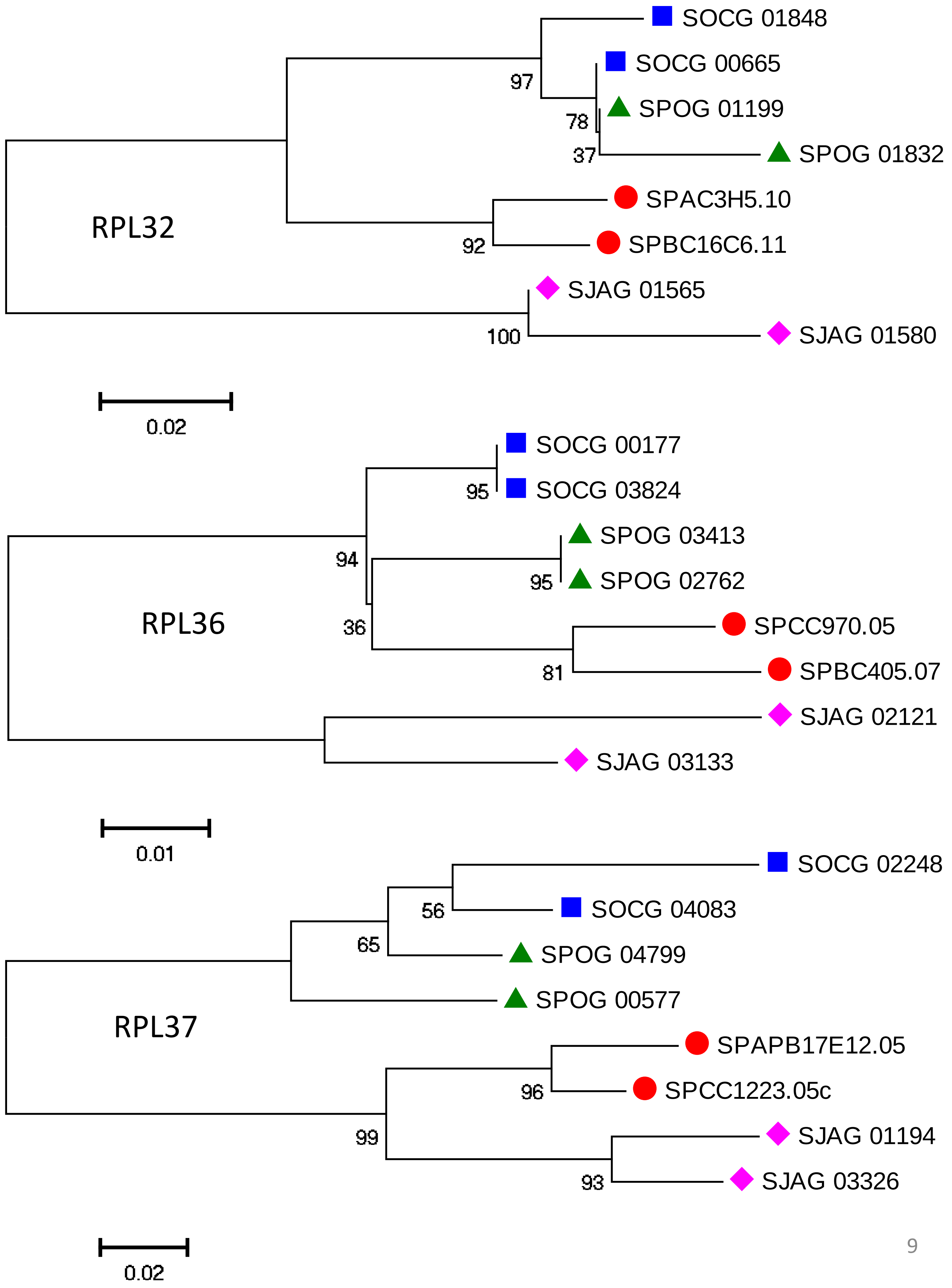
SOCG 01848
SOCG 00665
SPOG 01199
SPOG 01832
97
78
37
SPAC3H5.10
SPBC16C6.11
92
RPL32
SJAG 01565
SJAG 01580
100
0.02
SOCG 00177
SOCG 03824
95
SPOG 03413
SPOG 02762
95
94
36
SPCC970.05
SPBC405.07
81
RPL36
SJAG 02121
SJAG 03133
0.01
SOCG 02248
SOCG 04083
56
SPOG 04799
65
SPOG 00577
RPL37
SPAPB17E12.05
SPCC1223.05c
96
99
SJAG 01194
SJAG 03326
93
0.02

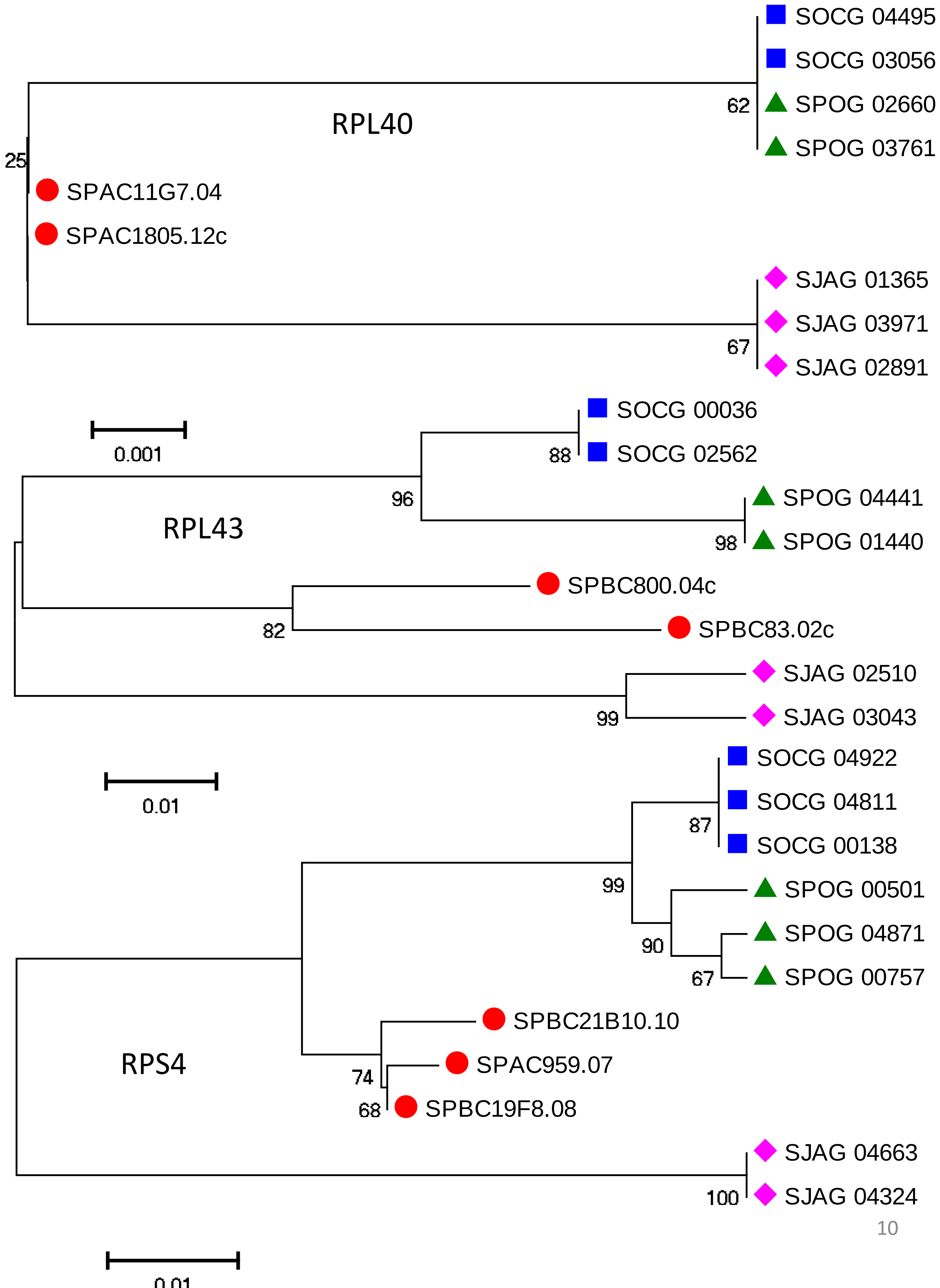
RPL40
SOCG 04495
SOCG 03056
SPOG 02660
SPOG 03761
62
25
SPAC11G7.04
SPAC1805.12c
SJAG 01365
SJAG 03971
SJAG 02891
67
0.001
RPL43
SOCG 00036
SOCG 02562
88
96
SPOG 04441
SPOG 01440
98
SPBC800.04c
SPBC83.02c
82
SJAG 02510
SJAG 03043
99
0.01
RPS4
SOCG 04922
SOCG 04811
SOCG 00138
87
99
SPOG 00501
SPOG 04871
SPOG 00757
90
67
SPBC21B10.10
SPAC959.07
SPBC19F8.08
74
68
SJAG 04663
SJAG 04324
100
0.01

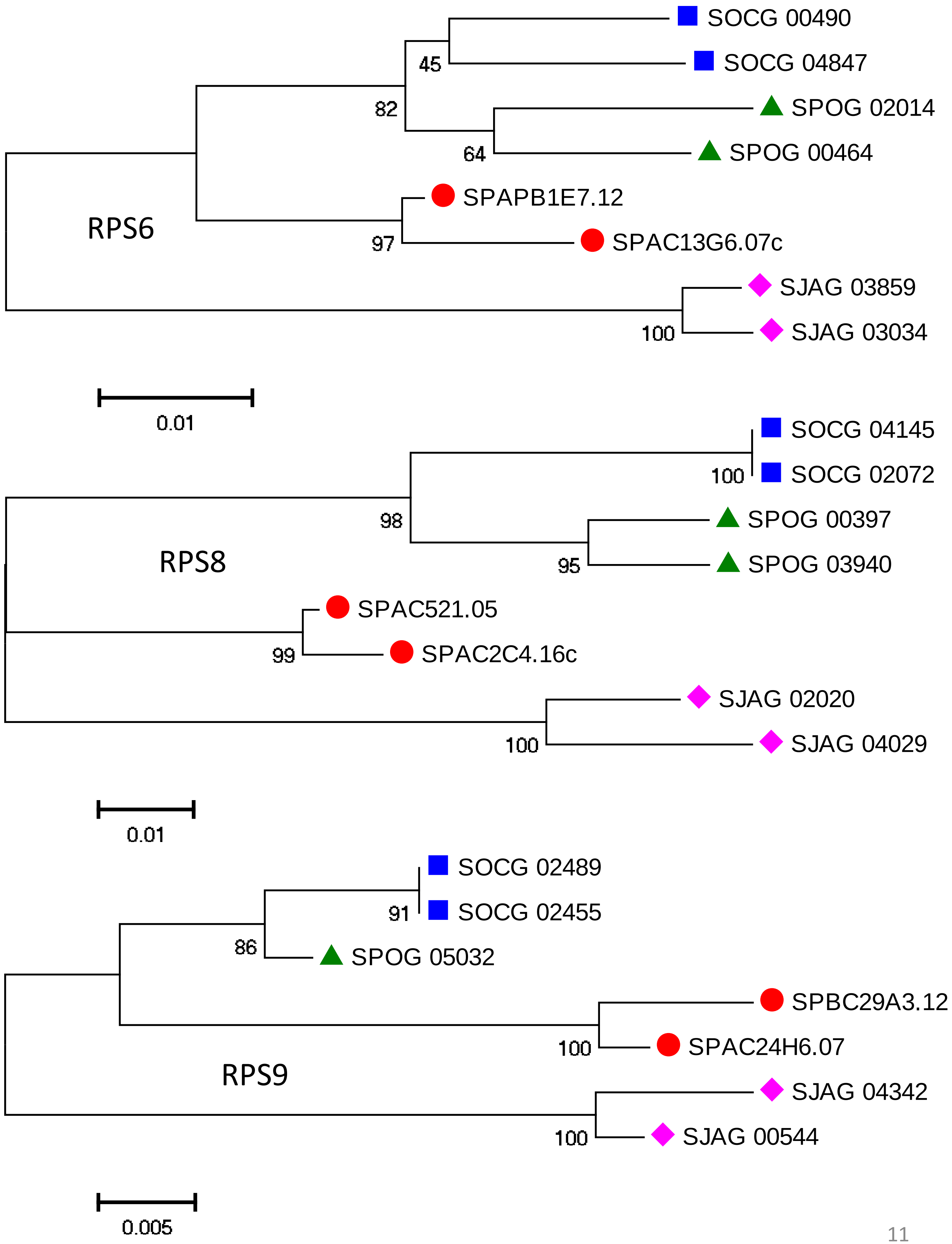
SOCG 00490
SOCG 04847
SPOG 02014
SPOG 00464
SPAPB1E7.12
SPAC13G6.07c
SJAG 03859
SJAG 03034
45
82
64
97
100
RPS6
0.01
SOCG 04145
SOCG 02072
SPOG 00397
SPOG 03940
SPAC521.05
SPAC2C4.16c
SJAG 02020
SJAG 04029
100
98
95
99
100
RPS8
0.01
SOCG 02489
SOCG 02455
SPOG 05032
SPBC29A3.12
SPAC24H6.07
SJAG 04342
SJAG 00544
91
86
100
100
RPS9
0.005

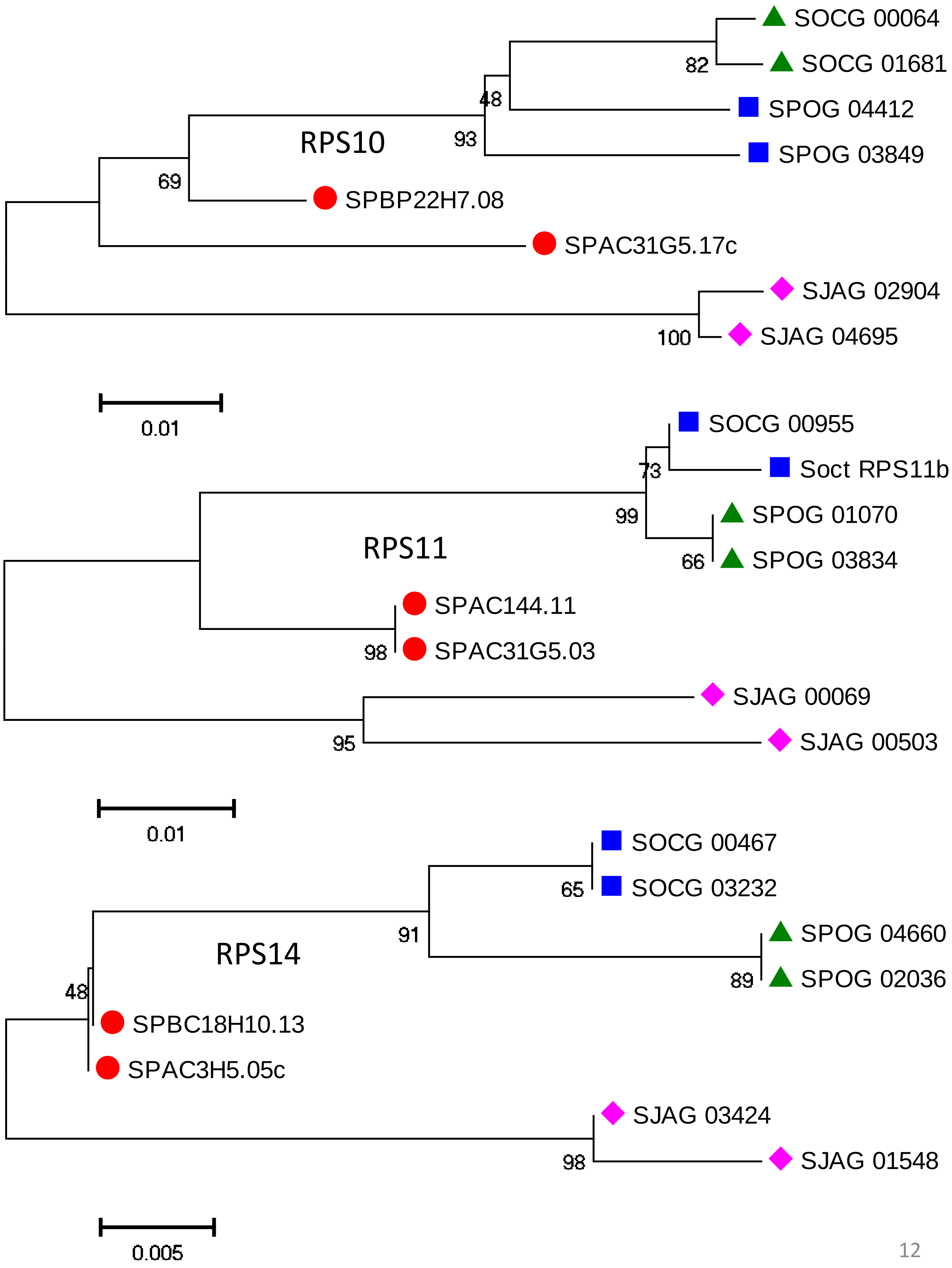
SOCG 00064
SOCG 01681
82
48
SPOG 04412
93
SPOG 03849
RPS10
69
SPBP22H7.08
SPAC31G5.17c
SJAG 02904
100
SJAG 04695
0.01
SOCG 00955
73
Soct RPS11b
99
SPOG 01070
RPS11
66
SPOG 03834
SPAC144.11
98
SPAC31G5.03
SJAG 00069
95
SJAG 00503
0.01
SOCG 00467
65
SOCG 03232
91
SPOG 04660
RPS14
89
SPOG 02036
48
SPBC18H10.13
SPAC3H5.05c
SJAG 03424
98
SJAG 01548
0.005

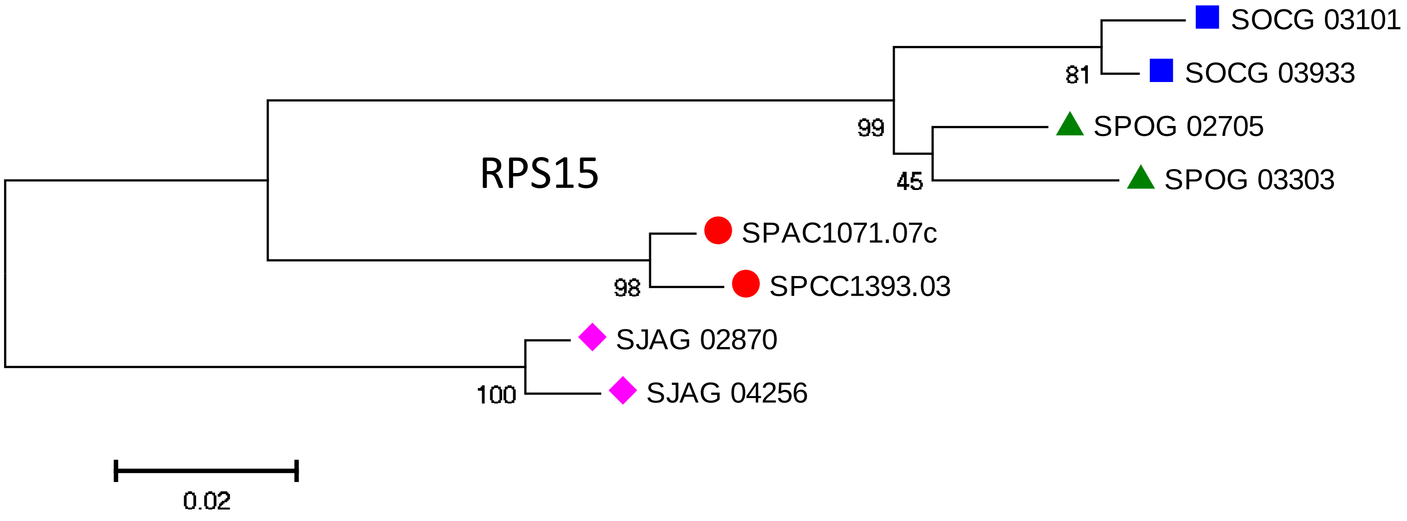
SOCG 03101
SOCG 03933
81
SPOG 02705
SPOG 03303
99
45
RPS15
SPAC1071.07c
SPCC1393.03
98
SJAG 02870
SJAG 04256
100
0.02

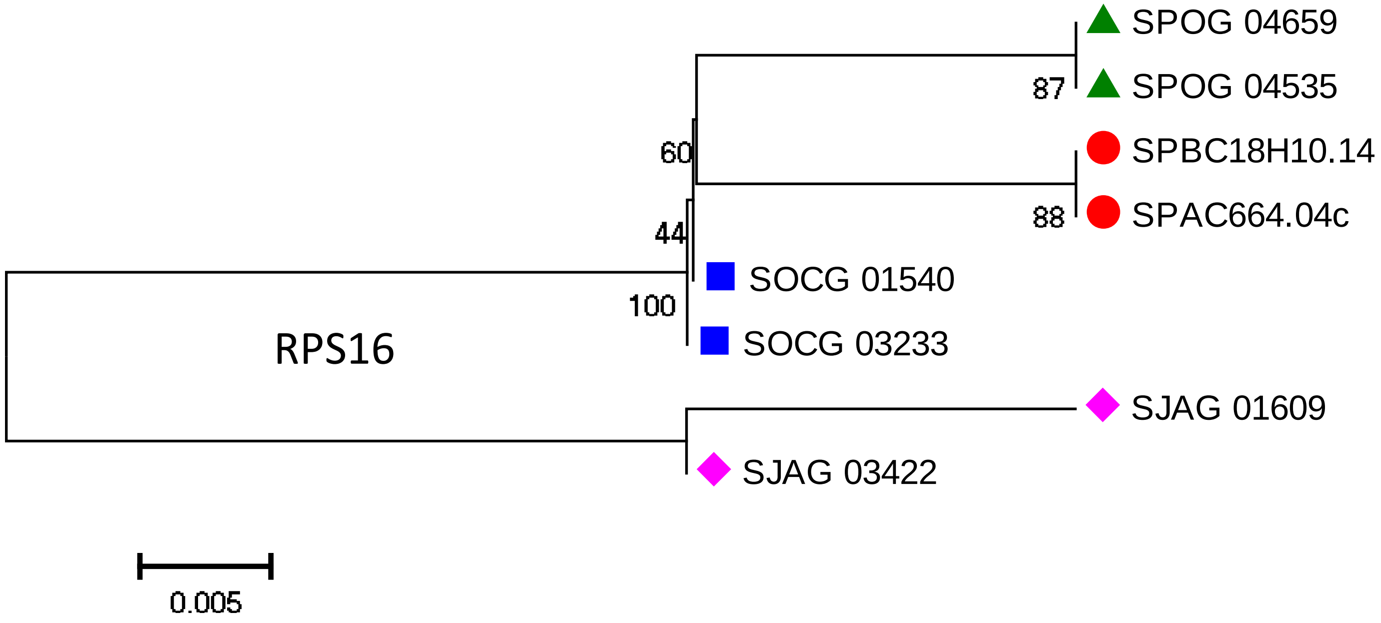
SPOG 04659
SPOG 04535
87
SPBC18H10.14
SPAC664.04c
88
60
44
SOCG 01540
SOCG 03233
100
RPS16
SJAG 01609
SJAG 03422
0.005

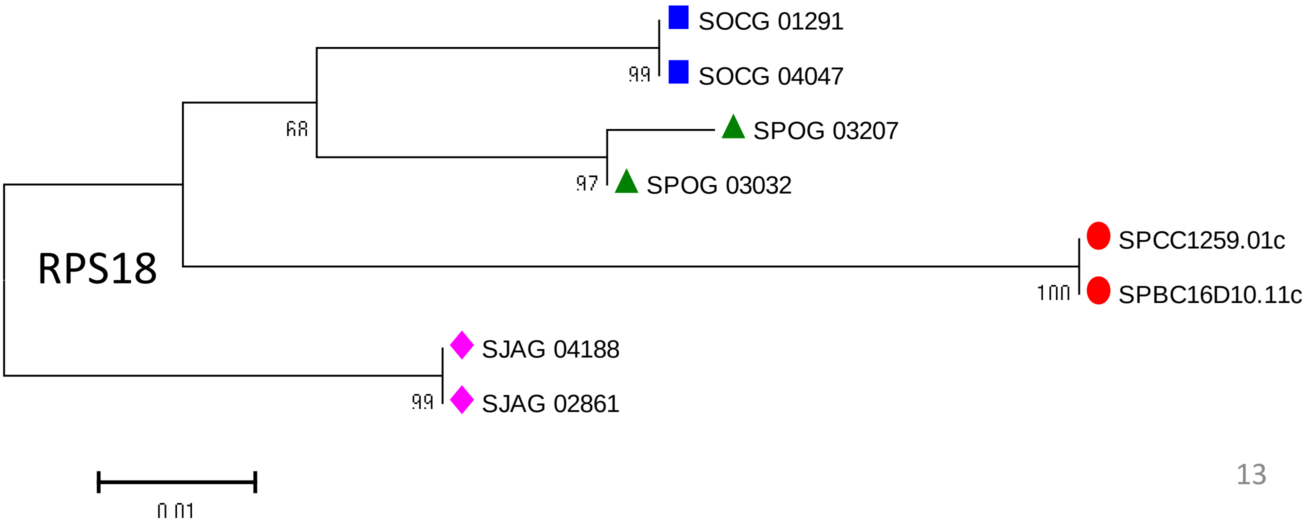
SOCG 01291
SOCG 04047
99
68
SPOG 03207
SPOG 03032
97
RPS18
SPCC1259.01c
SPBC16D10.11c
100
SJAG 04188
SJAG 02861
99
0.01

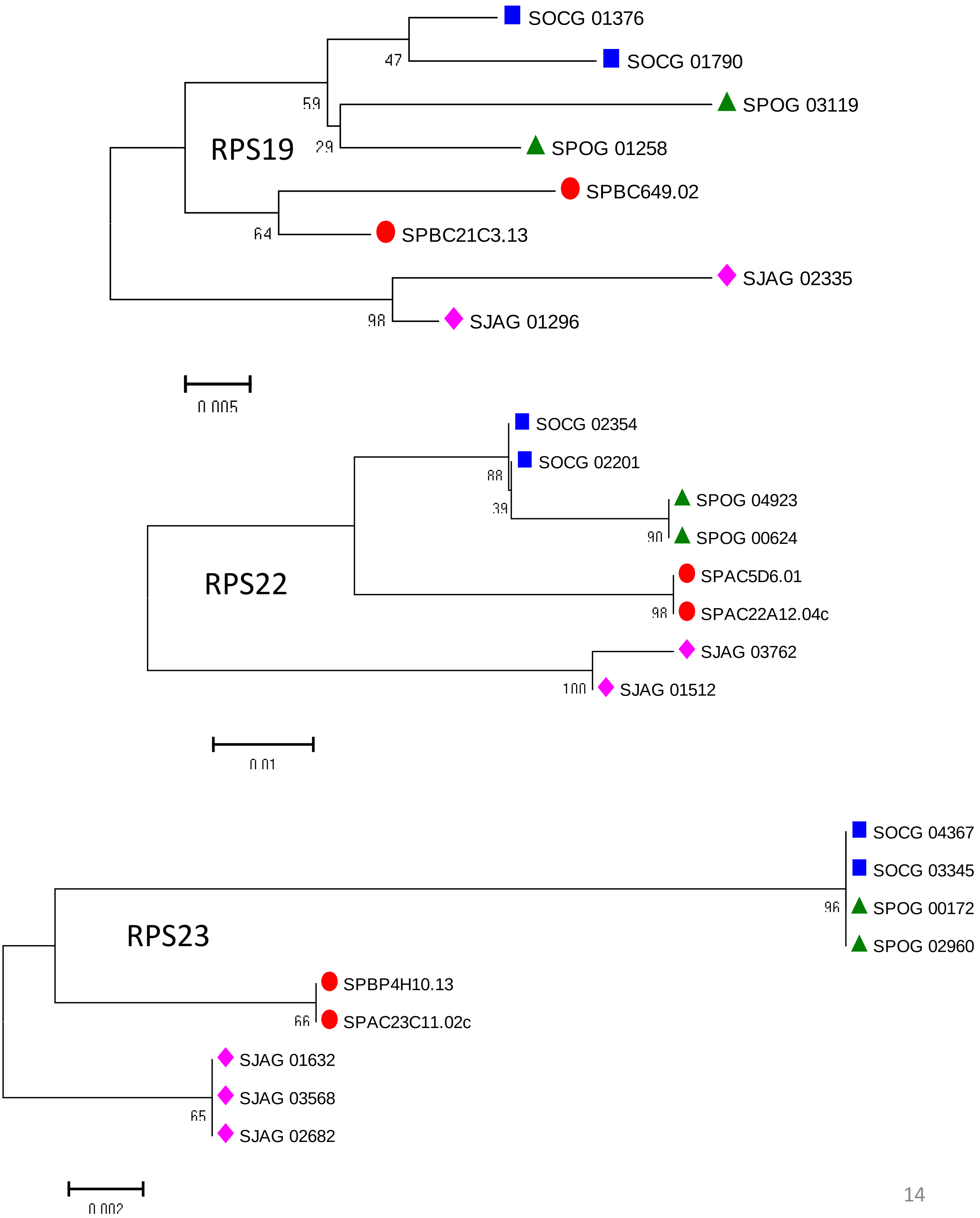
RPS19
SOCG 01376
SOCG 01790
SPOG 03119
SPOG 01258
SPBC649.02
SPBC21C3.13
SJAG 02335
SJAG 01296
47
59
29
64
98
0.005
RPS22
SOCG 02354
SOCG 02201
SPOG 04923
SPOG 00624
SPAC5D6.01
SPAC22A12.04c
SJAG 03762
SJAG 01512
88
39
90
98
100
0.01
RPS23
SOCG 04367
SOCG 03345
SPOG 00172
SPOG 02960
SPBP4H10.13
SPAC23C11.02c
SJAG 01632
SJAG 03568
SJAG 02682
96
66
65
0.002

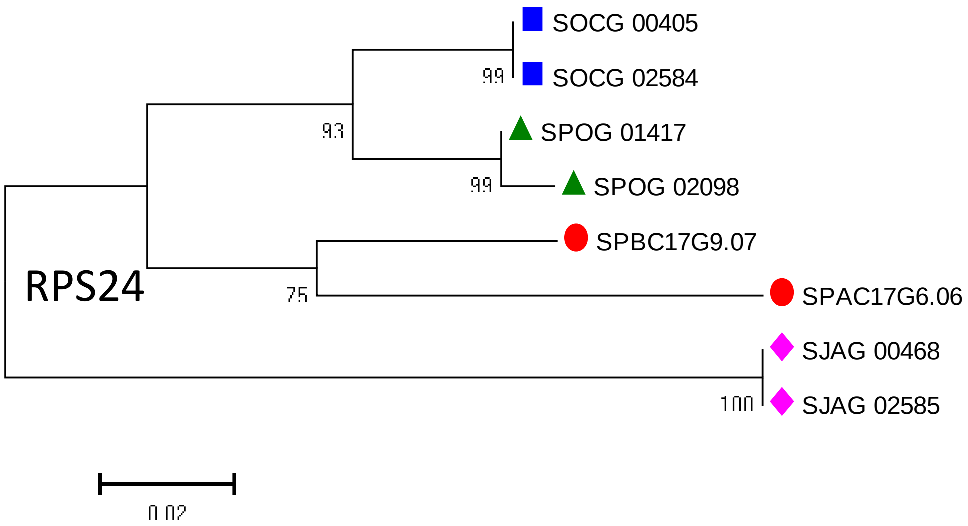
SOCG 00405
SOCG 02584
99
93
SPOG 01417
SPOG 02098
99
SPBC17G9.07
SPAC17G6.06
75
RPS24
SJAG 00468
SJAG 02585
100
0.02

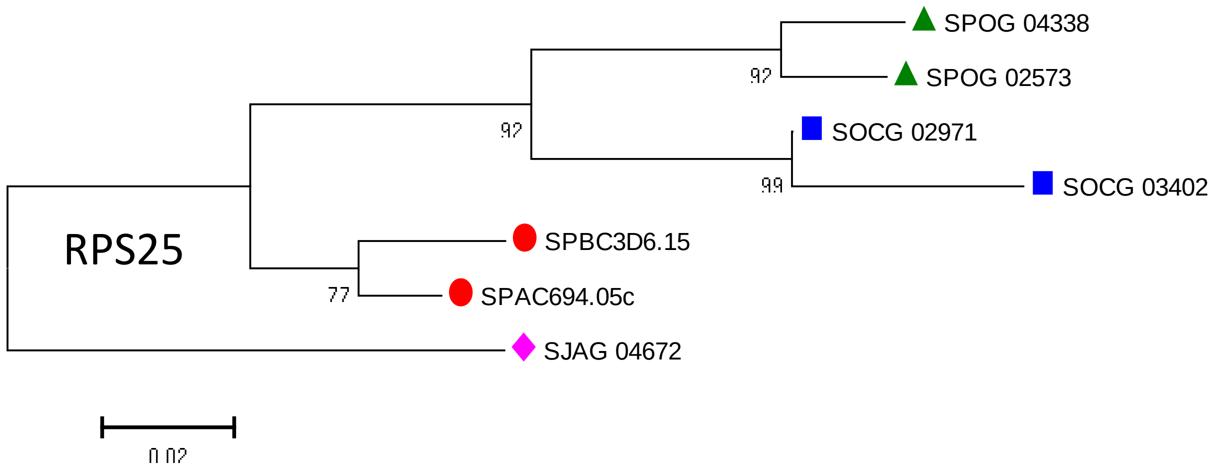
SPOG 04338
SPOG 02573
92
92
SOCG 02971
SOCG 03402
99
RPS25
SPBC3D6.15
SPAC694.05c
77
SJAG 04672
0.02

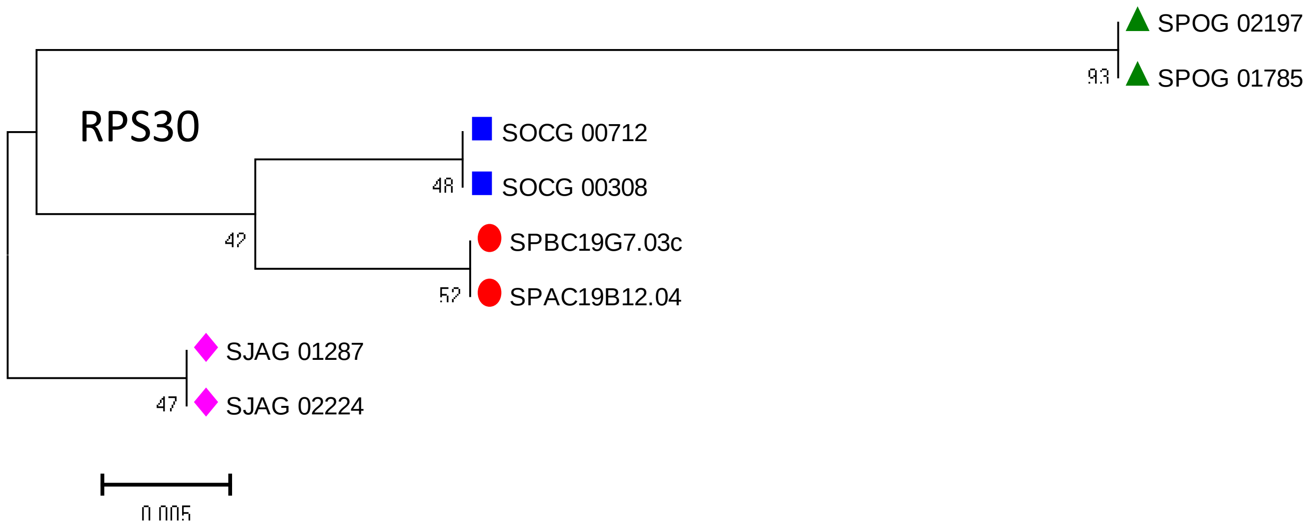
SPOG 02197
SPOG 01785
93
RPS30
SOCG 00712
SOCG 00308
48
42
SPBC19G7.03c
SPAC19B12.04
52
SJAG 01287
SJAG 02224
47
0.005

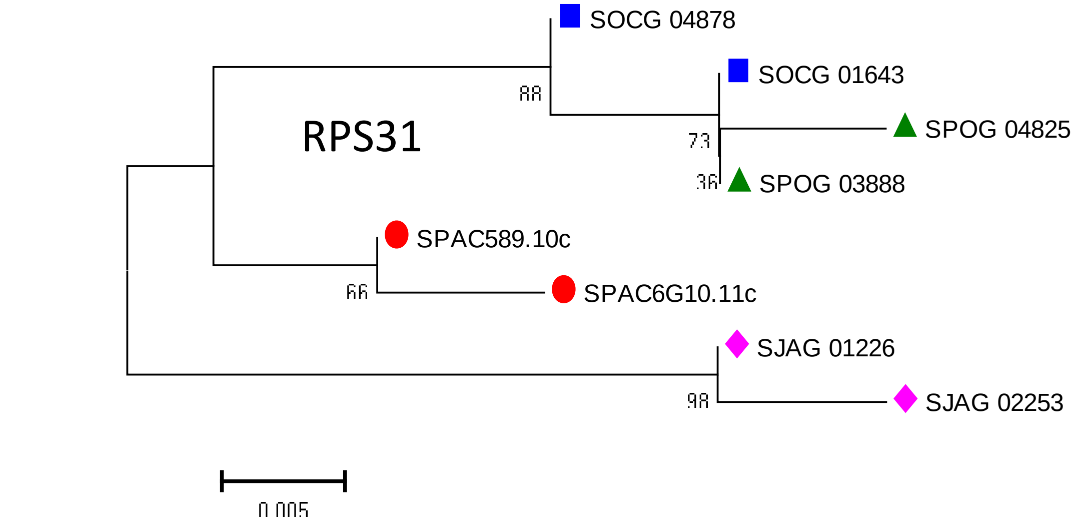
RPS31
SOCG 04878
SOCG 01643
SPOG 04825
SPOG 03888
SPAC589.10c
SPAC6G10.11c
SJAG 01226
SJAG 02253
88
73
36
66
98
0.005

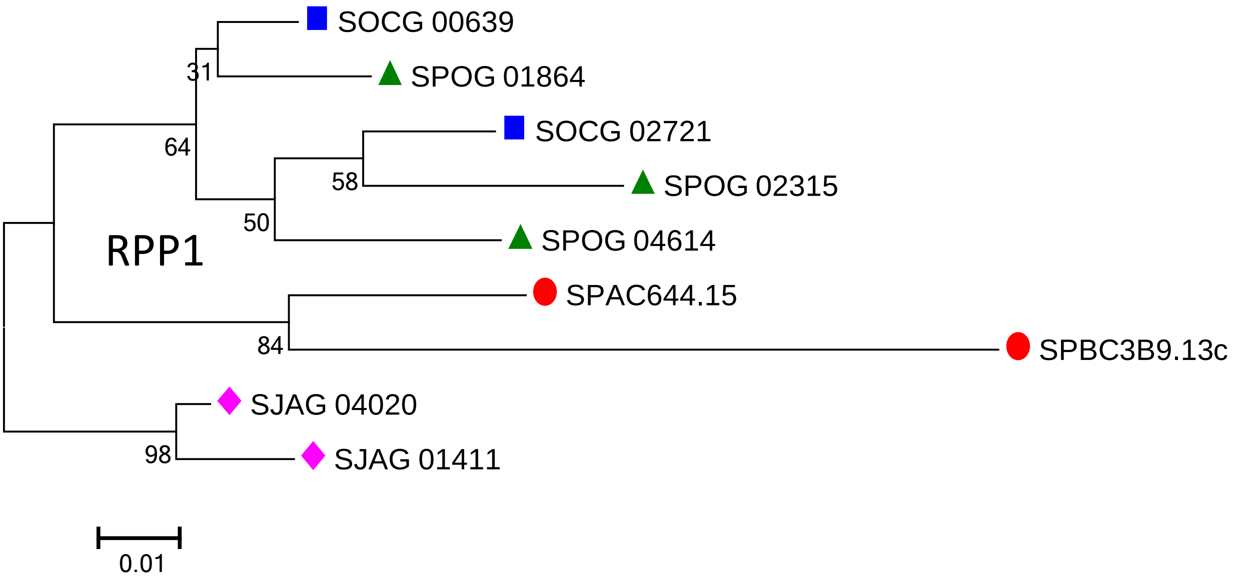
RPP1
SOCG 00639
SPOG 01864
SOCG 02721
SPOG 02315
SPOG 04614
SPAC644.15
SPBC3B9.13c
SJAG 04020
SJAG 01411
31
64
58
50
84
98
0.01

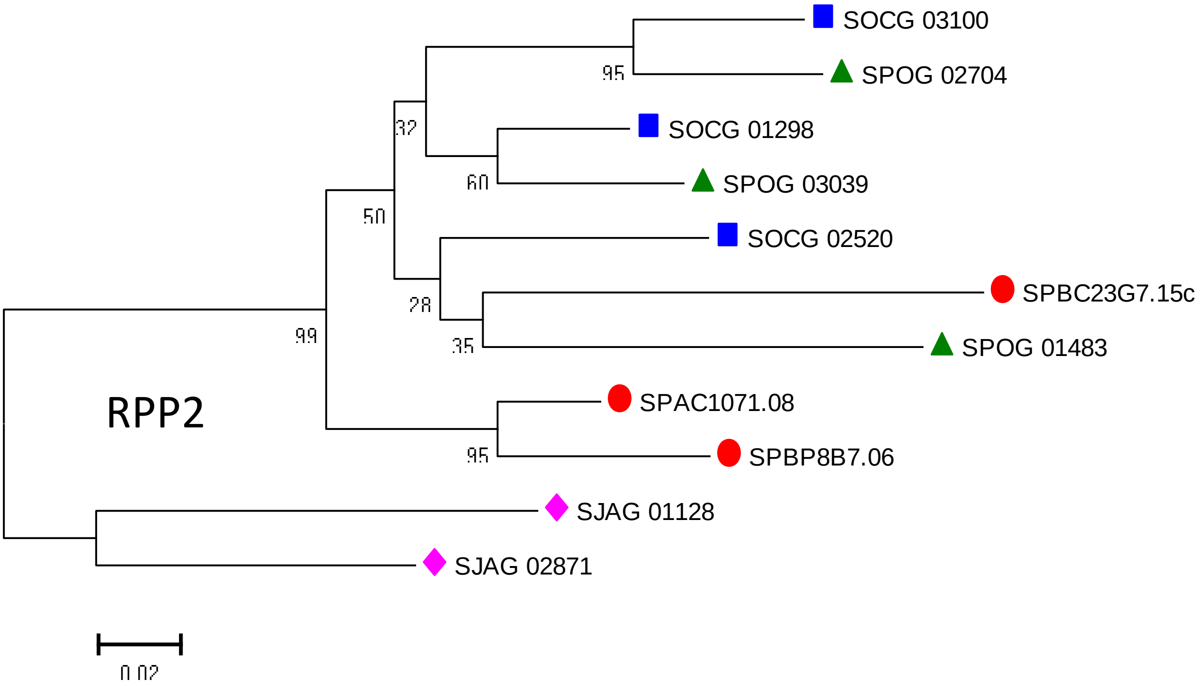
RPP2
SOCG 03100
SPOG 02704
SOCG 01298
SPOG 03039
SOCG 02520
SPBC23G7.15c
SPOG 01483
SPAC1071.08
SPBP8B7.06
SJAG 01128
SJAG 02871
95
32
60
50
28
35
99
95
0.02

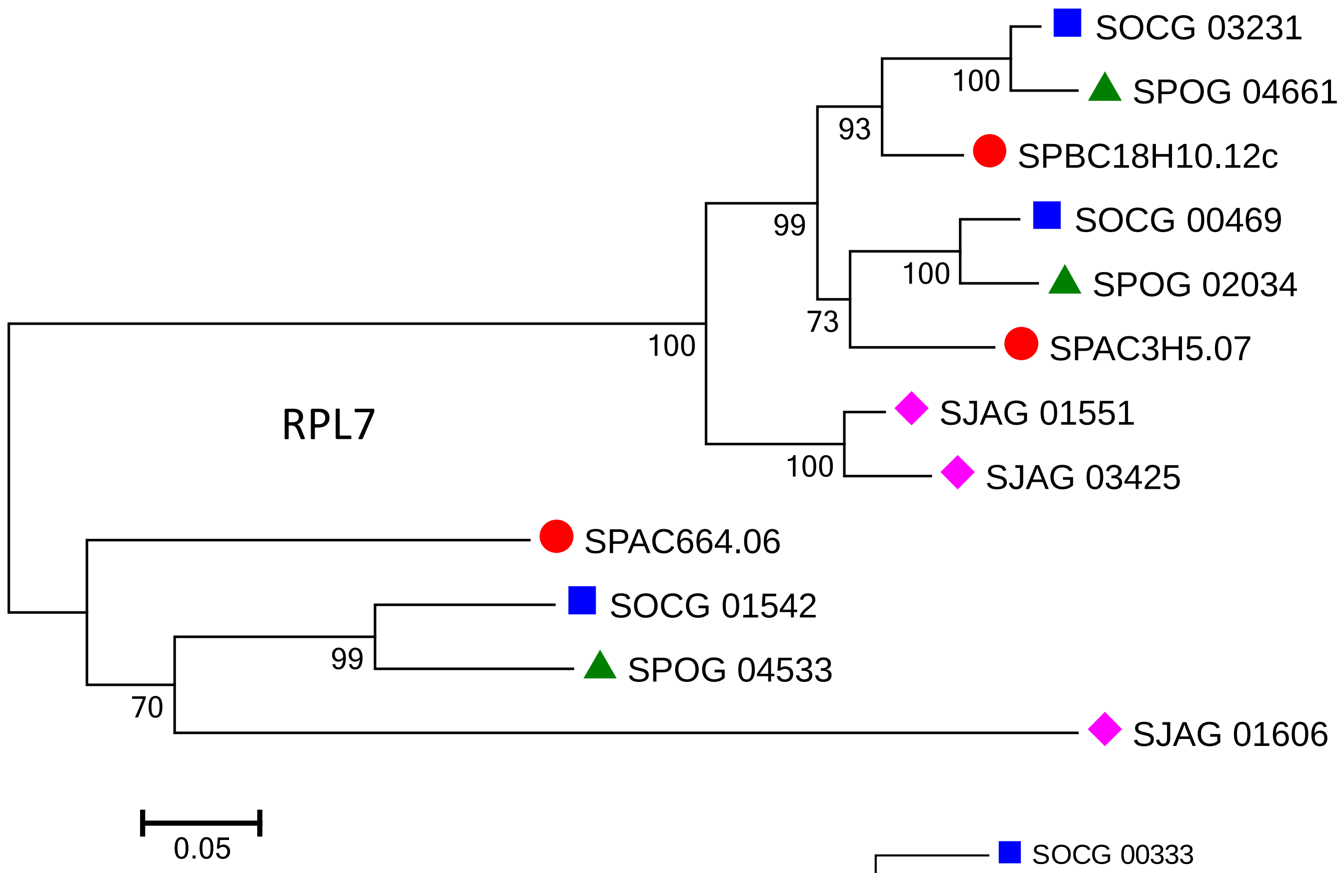
RPL7
SOCG 03231
SPOG 04661
SPBC18H10.12c
SOCG 00469
SPOG 02034
SPAC3H5.07
SJAG 01551
SJAG 03425
SPAC664.06
SOCG 01542
SPOG 04533
SJAG 01606
100
93
99
100
73
100
100
99
70
0.05

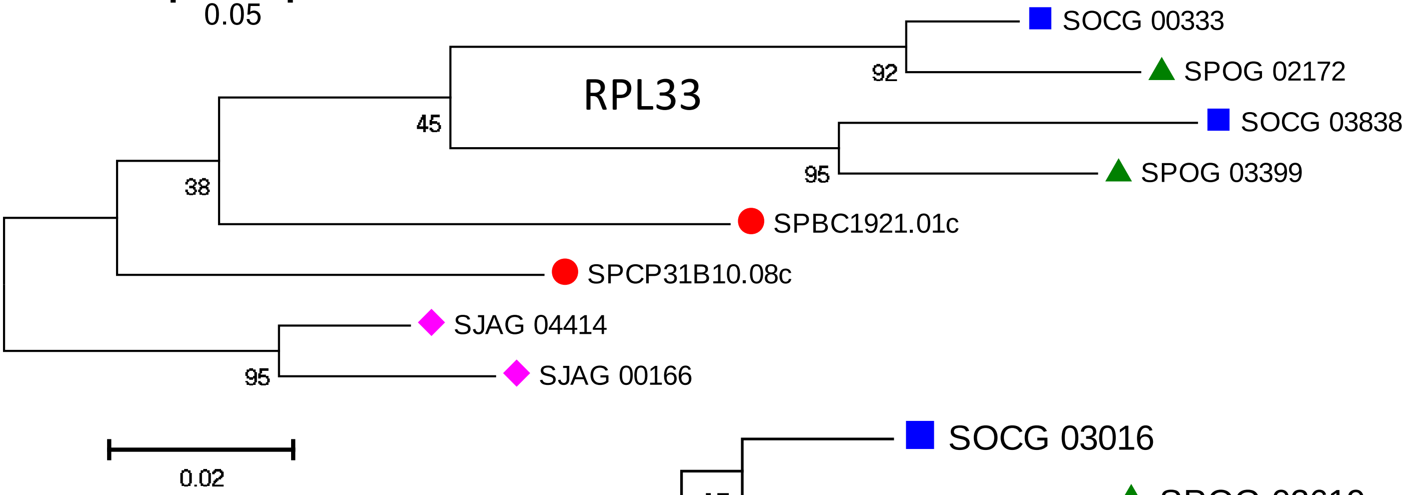
RPL33
SOCG 00333
SPOG 02172
SOCG 03838
SPOG 03399
SPBC1921.01c
SPCP31B10.08c
SJAG 04414
SJAG 00166
92
45
95
38
95
0.02

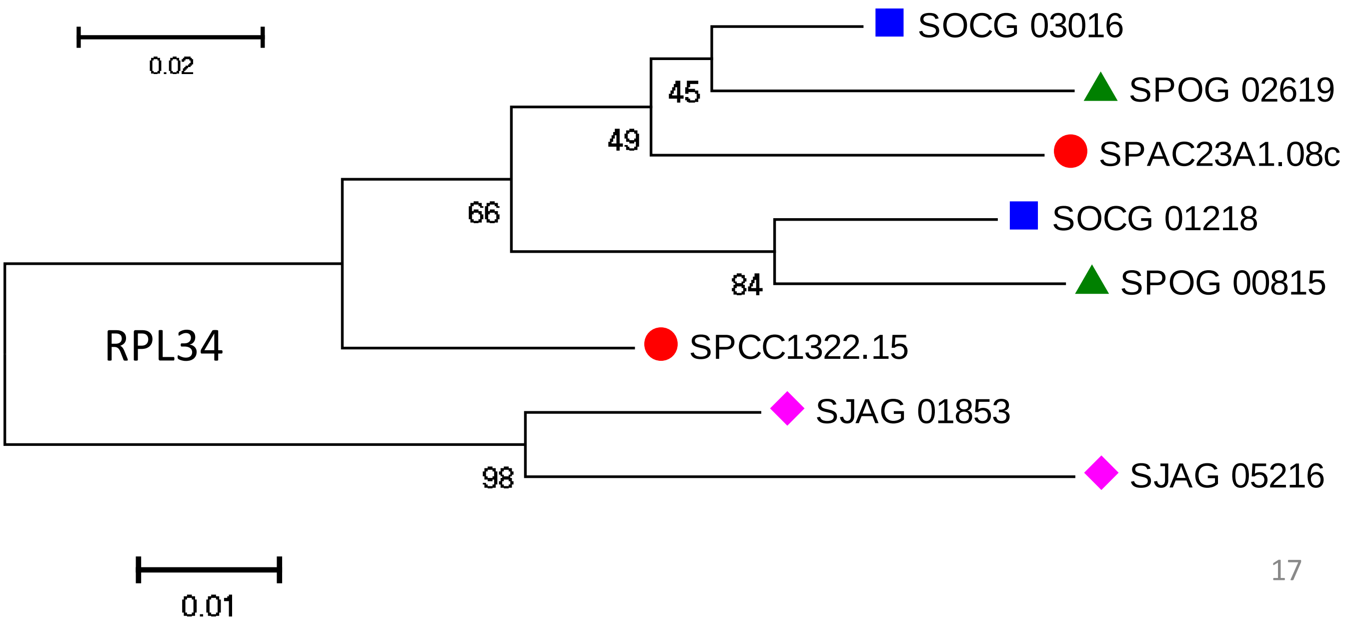
RPL34
SOCG 03016
SPOG 02619
SPAC23A1.08c
SOCG 01218
SPOG 00815
SPCC1322.15
SJAG 01853
SJAG 05216
45
49
66
84
98
0.01

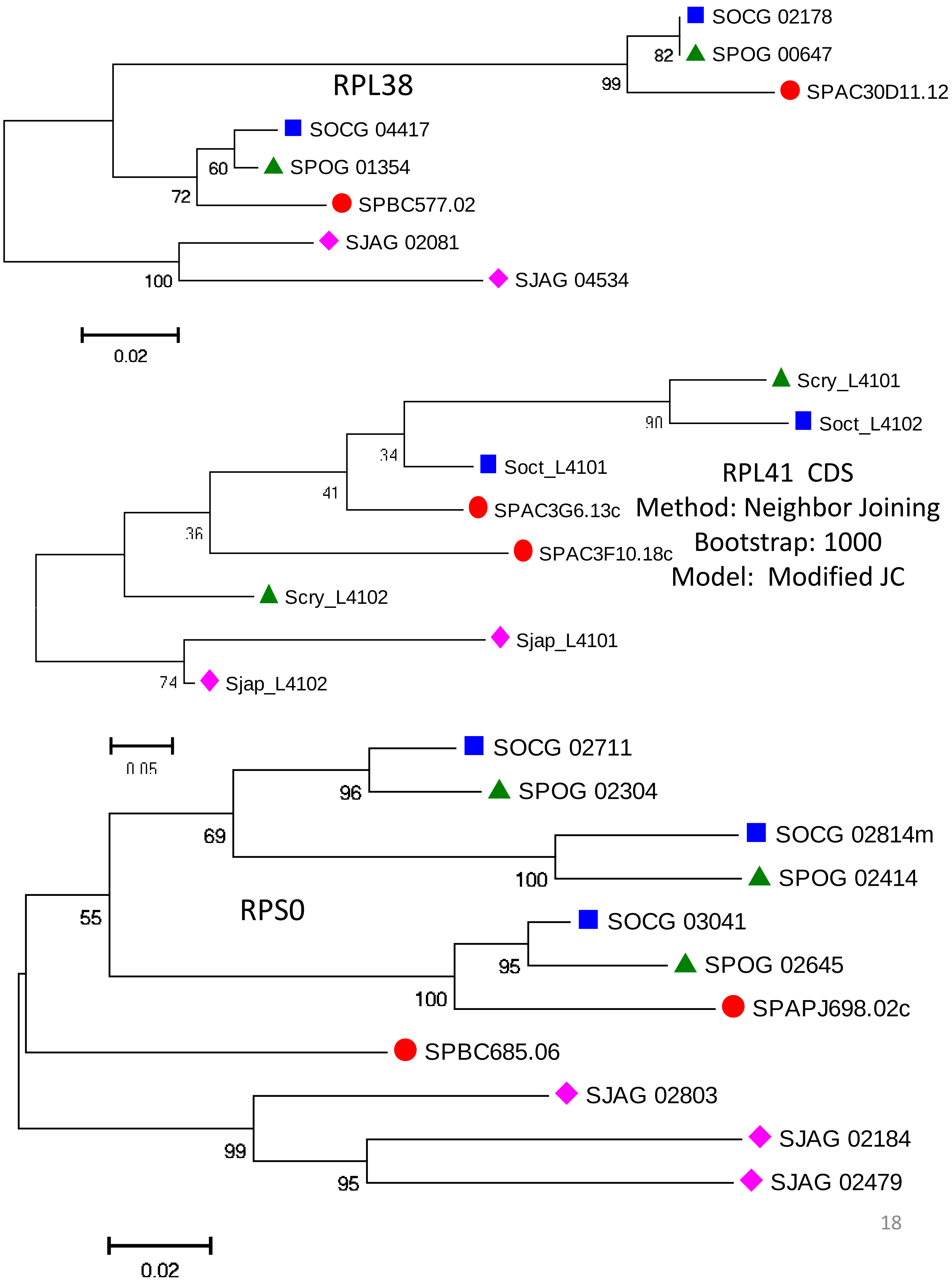

SOCG 02178
SPOG 00647
82
99
SPAC30D11.12
RPL38
SOCG 04417
SPOG 01354
60
72
SPBC577.02
SJAG 02081
SJAG 04534
100
0.02
Scry_L4101
Soct_L4102
90
34
Soct_L4101
RPL41 CDS
41
SPAC3G6.13c
Method: Neighbor Joining
36
SPAC3F10.18c
Bootstrap: 1000
Model: Modified JC
Scry_L4102
Sjap_L4101
74
Sjap_L4102
0.05
SOCG 02711
SPOG 02304
96
69
SOCG 02814m
SPOG 02414
100
55
RPS0
SOCG 03041
SPOG 02645
95
100
SPAPJ698.02c
SPBC685.06
SJAG 02803
SJAG 02184
99
95
SJAG 02479
0.02

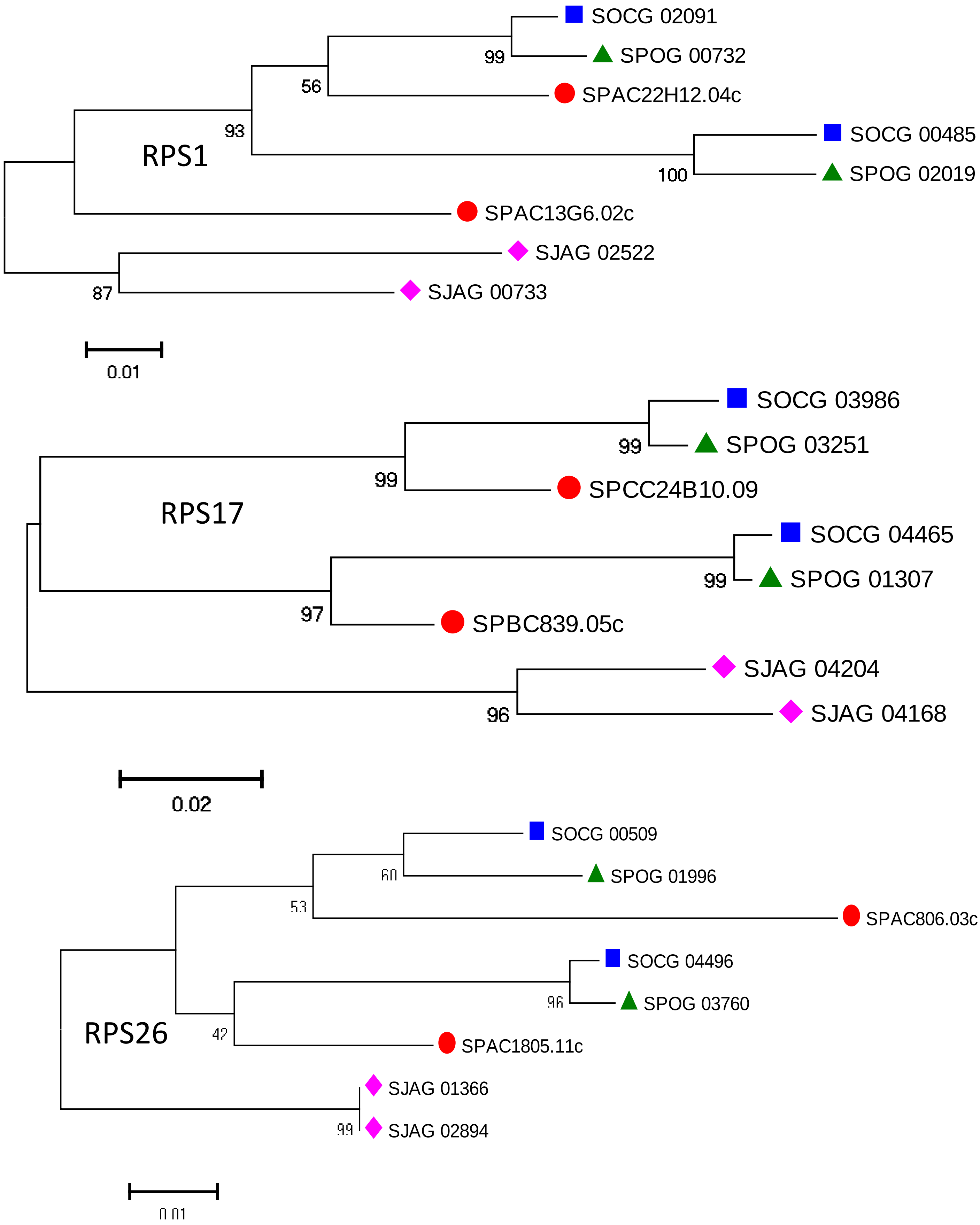

SOCG 02091
SPOG 00732
99
SPAC22H12.04c
56
93
RPS1
SOCG 00485
SPOG 02019
100
SPAC13G6.02c
SJAG 02522
87
SJAG 00733
0.01
SOCG 03986
SPOG 03251
99
99
SPCC24B10.09
RPS17
SOCG 04465
SPOG 01307
99
97
SPBC839.05c
SJAG 04204
96
SJAG 04168
0.02
SOCG 00509
SPOG 01996
60
53
SPAC806.03c
SOCG 04496
SPOG 03760
96
RPS26
42
SPAC1805.11c
SJAG 01366
99
SJAG 02894
0.01

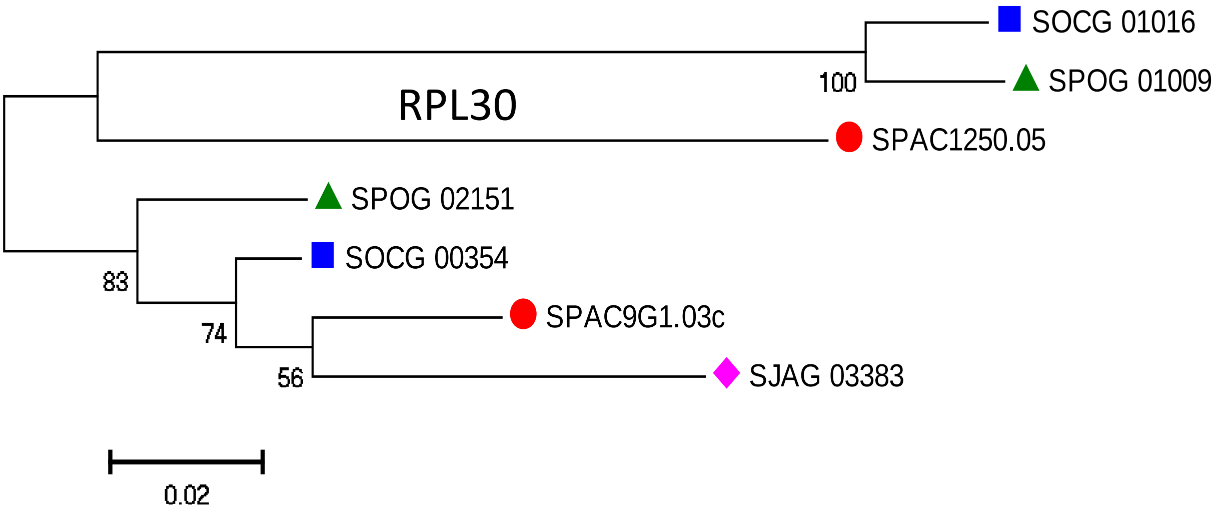
RPL30
SOCG 01016
SPOG 01009
100
SPAC1250.05
SPOG 02151
SOCG 00354
83
74
SPAC9G1.03c
56
SJAG 03383
0.02

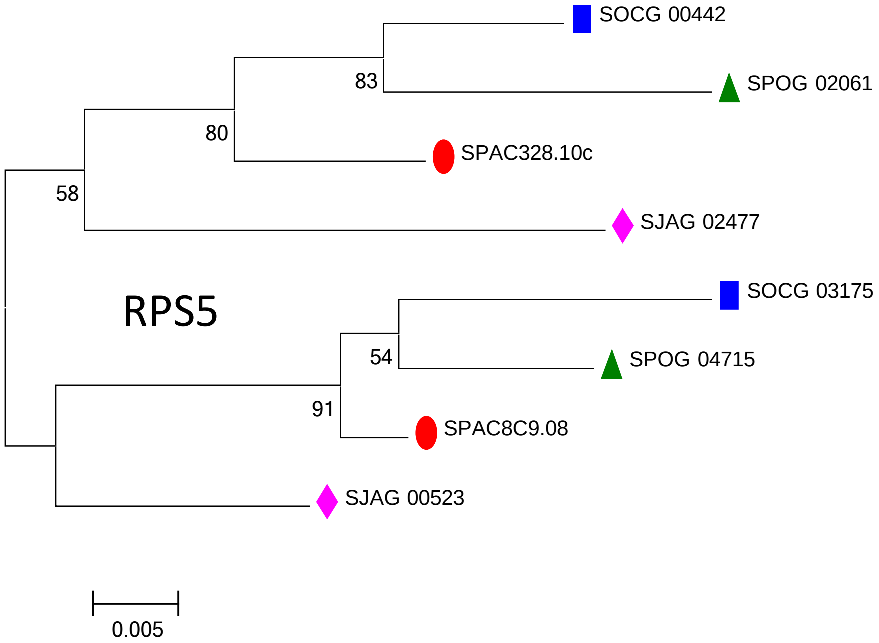
RPS5
SOCG 00442
83
SPOG 02061
80
SPAC328.10c
58
SJAG 02477
SOCG 03175
54
SPOG 04715
91
SPAC8C9.08
SJAG 00523
0.005

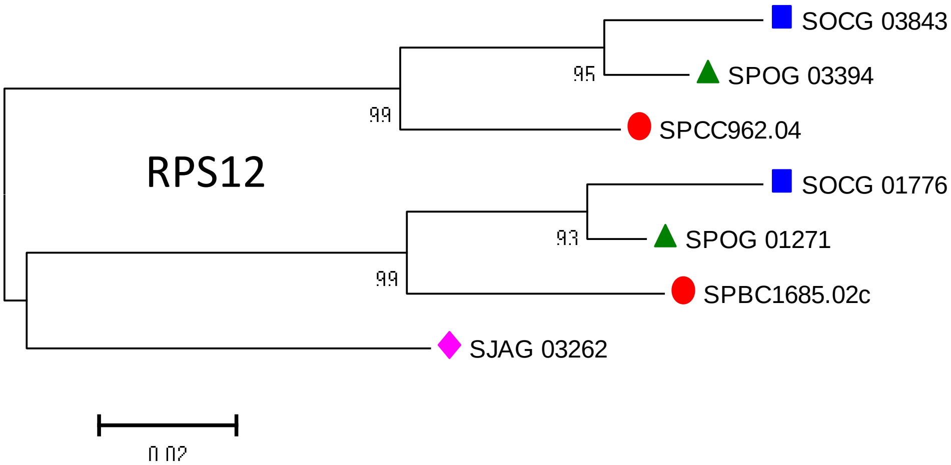
RPS12
SOCG 03843
95
SPOG 03394
99
SPCC962.04
SOCG 01776
93
SPOG 01271
99
SPBC1685.02c
SJAG 03262
0.02

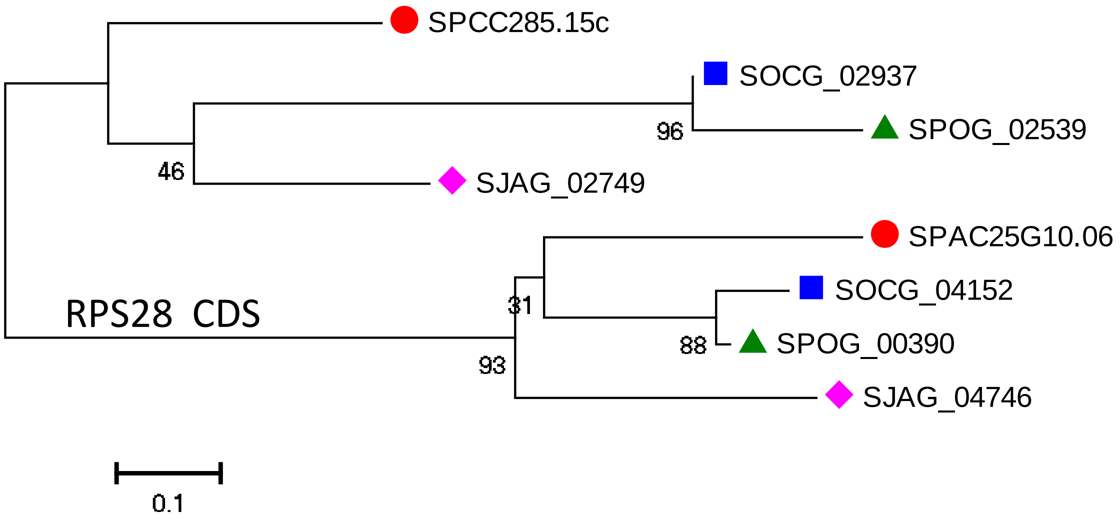
SPCC285.15c
SOCG_02937
96
SPOG_02539
46
SJAG_02749
SPAC25G10.06
SOCG_04152
31
RPS28 CDS
88
SPOG_00390
93
SJAG_04746
0.1
