## Supplemental File 4 for "Parallel Concerted Evolution of Ribosomal Protein Genes in Fungi and Its Adaptive Significance"

**Supplementary File 4: Phylogenetic trees of RP families in the four Mucoromycotina species.**

| Gene family | Category | Page | Gene family | Category | Page |
|---|---|---|---|---|---|
| RPL2 | A | 2 | RPS5 | A | 15 |
| RPL3 | A | 2 | RPS6 | A | 15 |
| RPL4 | A | 2 | RPS8 | A | 15 |
| RPL5 | A | 3 | RPS9 | A | 16 |
| RPL7 | A | 3 | RPS13 | A | 16 |
| RPL8 | A | 3 | RPS14 | A | 16 |
| RPL9 | A | 4 | RPS15 | A | 17 |
| RPL10 | A | 4 | RPS18 | A | 17 |
| RPL11 | A | 4 | RPS19 | A | 17 |
| RPL12 | A | 5 | RPS21 | A | 18 |
| RPL13 | A | 5 | RPS22 | A | 18 |
| RPL14 | A | 5 | RPS23 | A | 18 |
| RPL15 | A | 6 | RPS25 | A | 19 |
| RPL16 | A | 6 | RPS26 | A | 19 |
| RPL17 | A | 6 | RPS27 | A | 19 |
| RPL18 | A | 7 | RPS28 | A | 20 |
| RPL21 | A | 7 | RPS29 | A | 20 |
| RPL23 | A | 7 | RPS30 | A | 20 |
| RPL24 | A | 8 | RPL1 | B | 21 |
| RPL25 | A | 8 | RPL6 | B | 21 |
| RPL26 | A | 8 | RPL19 | B | 21 |
| RPL27 | A | 9 | RPL20 | B | 22 |
| RPL28 | A | 9 | RPL22 | B | 22 |
| RPL28e | A | 9 | RPL31 | B | 22 |
| RPL29 | A | 10 | RPL38 | B | 23 |
| RPL30 | A | 10 | RPL39 | B | 23 |
| RPL32 | A | 10 | RPL43 | B | 23 |
| RPL33 | A | 11 | RPS7 | B | 24 |
| RPL34 | A | 11 | RPS10 | B | 24 |
| RPL35 | A | 11 | RPS12 | B | 24 |
| RPL36 | A | 12 | RPS16 | B | 25 |
| RPL37 | A | 12 | RPS17 | B | 25 |
| RPL40 | A | 12 | RPS24 | B | 25 |
| RPL42 | A | 13 | RPS31 | B | 26 |
| RPP1 | A | 13 | RPP2 | B | 26 |
| RPS1 | A | 13 | RPS0 | C | 27 |
| RPS2 | A | 14 | RPS20 | C | 27 |
| RPS3 | A | 14 | | | |
| RPS4 | A | 14 | | | |

*The phylogenetic trees were built based on amino acid sequences of four WGD budding yeast species, unless specially specified. Each tree was reconstructed by Neighbor-Joining method using Poisson correction with 1000 Bootstrap replicates. For some trees, if the phylogenetic tree topology is not resolved due to high degree of sequence similarity, we used their CDS sequences to build their phylogenetic tree.

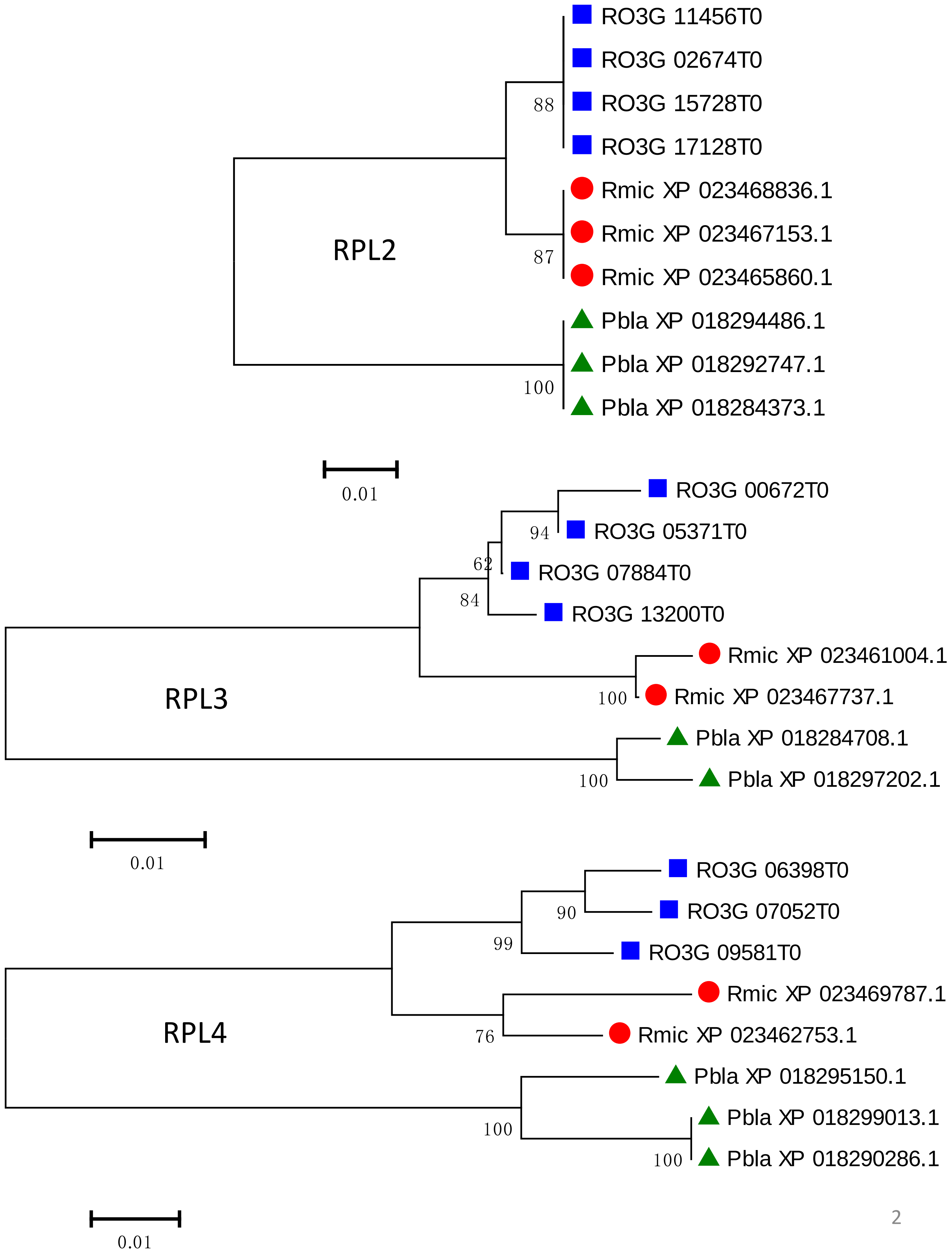
RO3G 11456T0
RO3G 02674T0
RO3G 15728T0
RO3G 17128T0
88
RPL2
Rmic XP 023468836.1
Rmic XP 023467153.1
Rmic XP 023465860.1
87
Pbla XP 018294486.1
Pbla XP 018292747.1
Pbla XP 018284373.1
100
0.01
RO3G 00672T0
RO3G 05371T0
94
RO3G 07884T0
62
84
RO3G 13200T0
Rmic XP 023461004.1
Rmic XP 023467737.1
100
RPL3
Pbla XP 018284708.1
Pbla XP 018297202.1
100
0.01
RO3G 06398T0
RO3G 07052T0
90
99
RO3G 09581T0
Rmic XP 023469787.1
RPL4
76
Rmic XP 023462753.1
Pbla XP 018295150.1
Pbla XP 018299013.1
100
Pbla XP 018290286.1
100
0.01

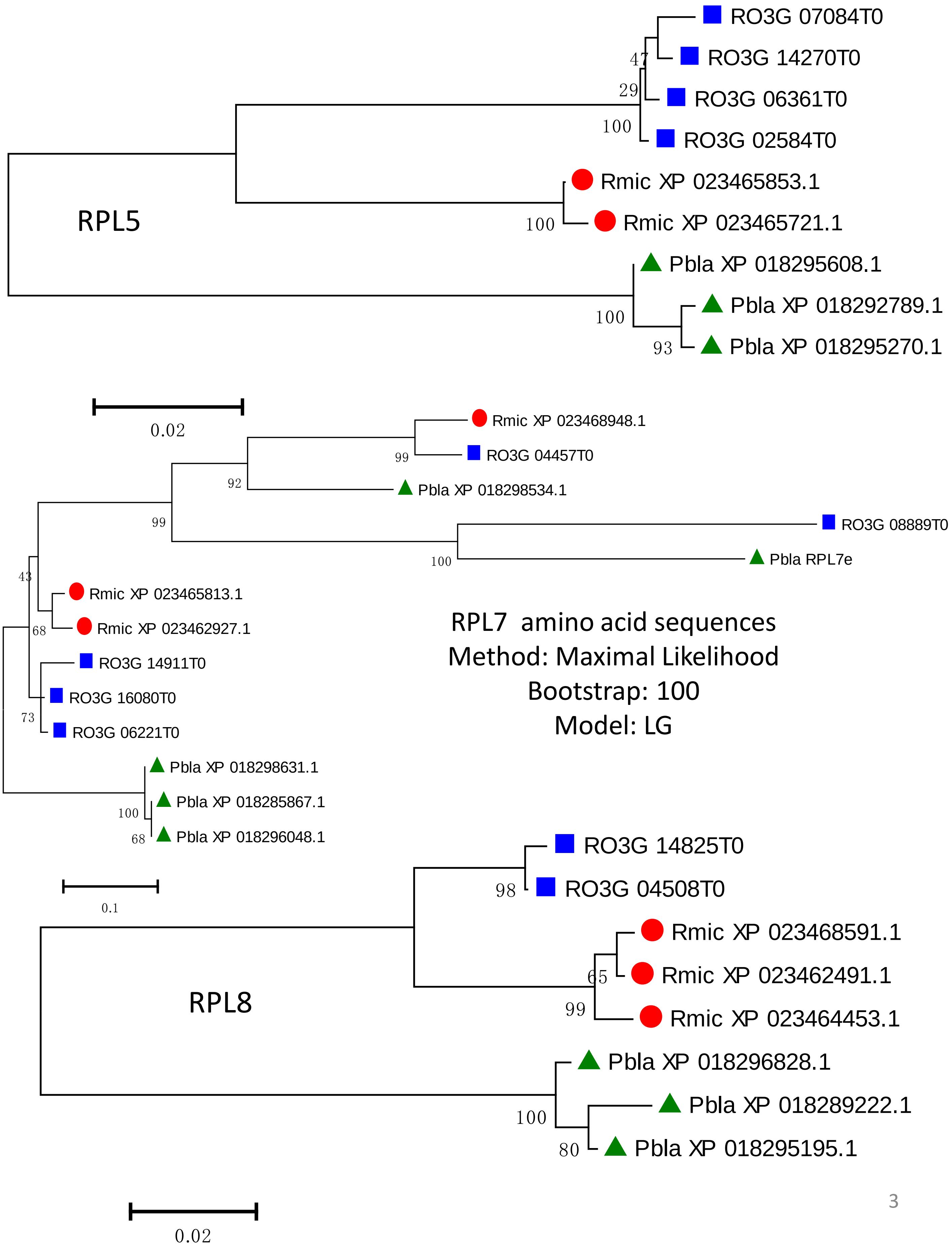
RO3G 07084T0
RO3G 14270T0
RO3G 06361T0
RO3G 02584T0
Rmic XP 023465853.1
Rmic XP 023465721.1
Pbla XP 018295608.1
Pbla XP 018292789.1
Pbla XP 018295270.1
47
29
100
100
100
93
RPL5
0.02
Rmic XP 023468948.1
RO3G 04457T0
Pbla XP 018298534.1
RO3G 08889T0
Pbla RPL7e
Rmic XP 023465813.1
Rmic XP 023462927.1
RO3G 14911T0
RO3G 16080T0
RO3G 06221T0
Pbla XP 018298631.1
Pbla XP 018285867.1
Pbla XP 018296048.1
99
92
99
100
43
68
73
100
68
RPL7 amino acid sequences
Method: Maximal Likelihood
Bootstrap: 100
Model: LG
0.1
RO3G 14825T0
RO3G 04508T0
Rmic XP 023468591.1
Rmic XP 023462491.1
Rmic XP 023464453.1
Pbla XP 018296828.1
Pbla XP 018289222.1
Pbla XP 018295195.1
98
65
99
100
80
RPL8
0.02

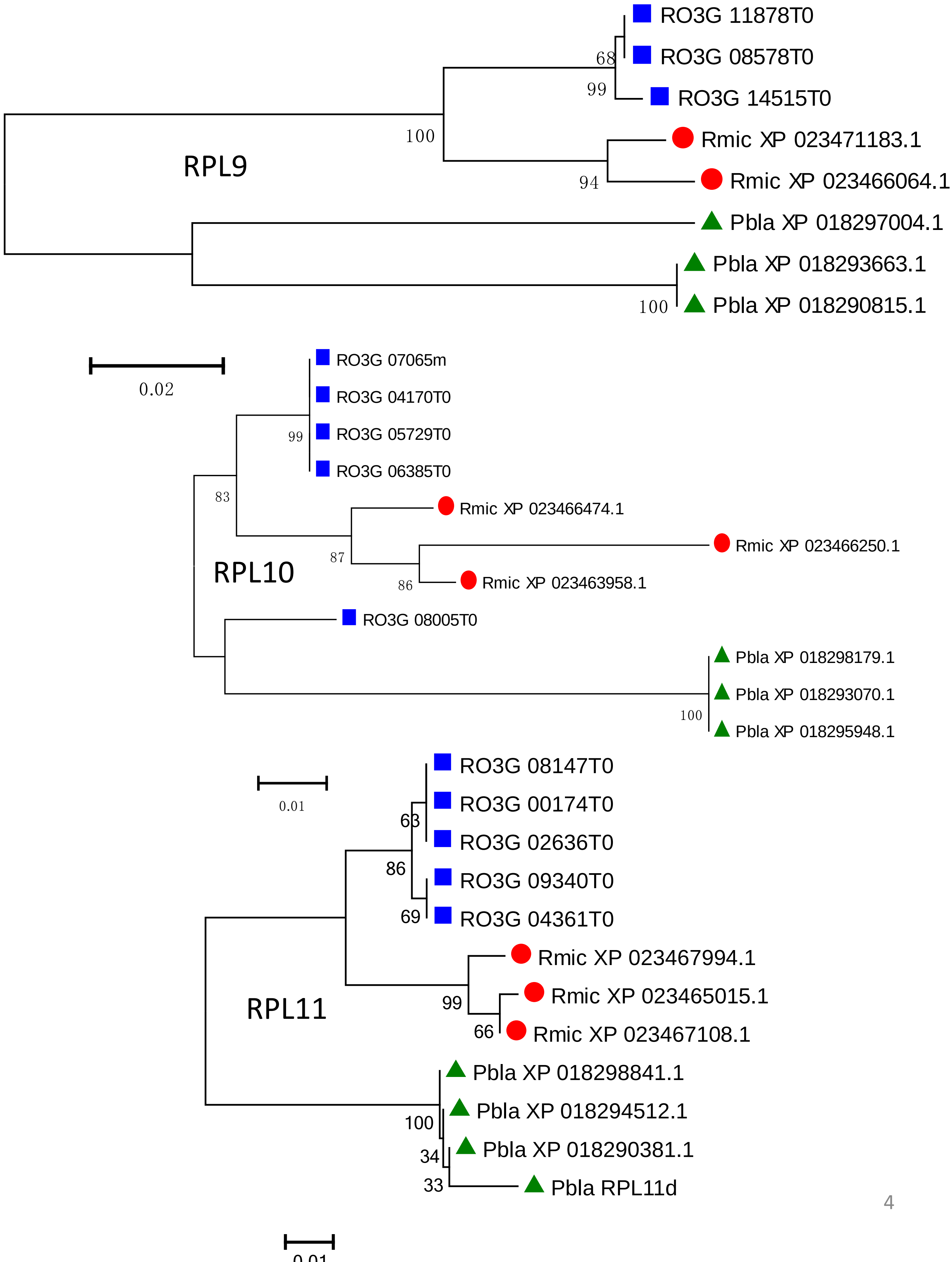
RPL9
RO3G 11878T0
RO3G 08578T0
RO3G 14515T0
Rmic XP 023471183.1
Rmic XP 023466064.1
Pbla XP 018297004.1
Pbla XP 018293663.1
Pbla XP 018290815.1
68
99
100
94
100
0.02
RPL10
RO3G 07065m
RO3G 04170T0
RO3G 05729T0
RO3G 06385T0
Rmic XP 023466474.1
Rmic XP 023466250.1
Rmic XP 023463958.1
RO3G 08005T0
Pbla XP 018298179.1
Pbla XP 018293070.1
Pbla XP 018295948.1
99
83
87
86
100
0.01
RPL11
RO3G 08147T0
RO3G 00174T0
RO3G 02636T0
RO3G 09340T0
RO3G 04361T0
Rmic XP 023467994.1
Rmic XP 023465015.1
Rmic XP 023467108.1
Pbla XP 018298841.1
Pbla XP 018294512.1
Pbla XP 018290381.1
Pbla RPL11d
63
86
69
99
66
100
34
33
0.01

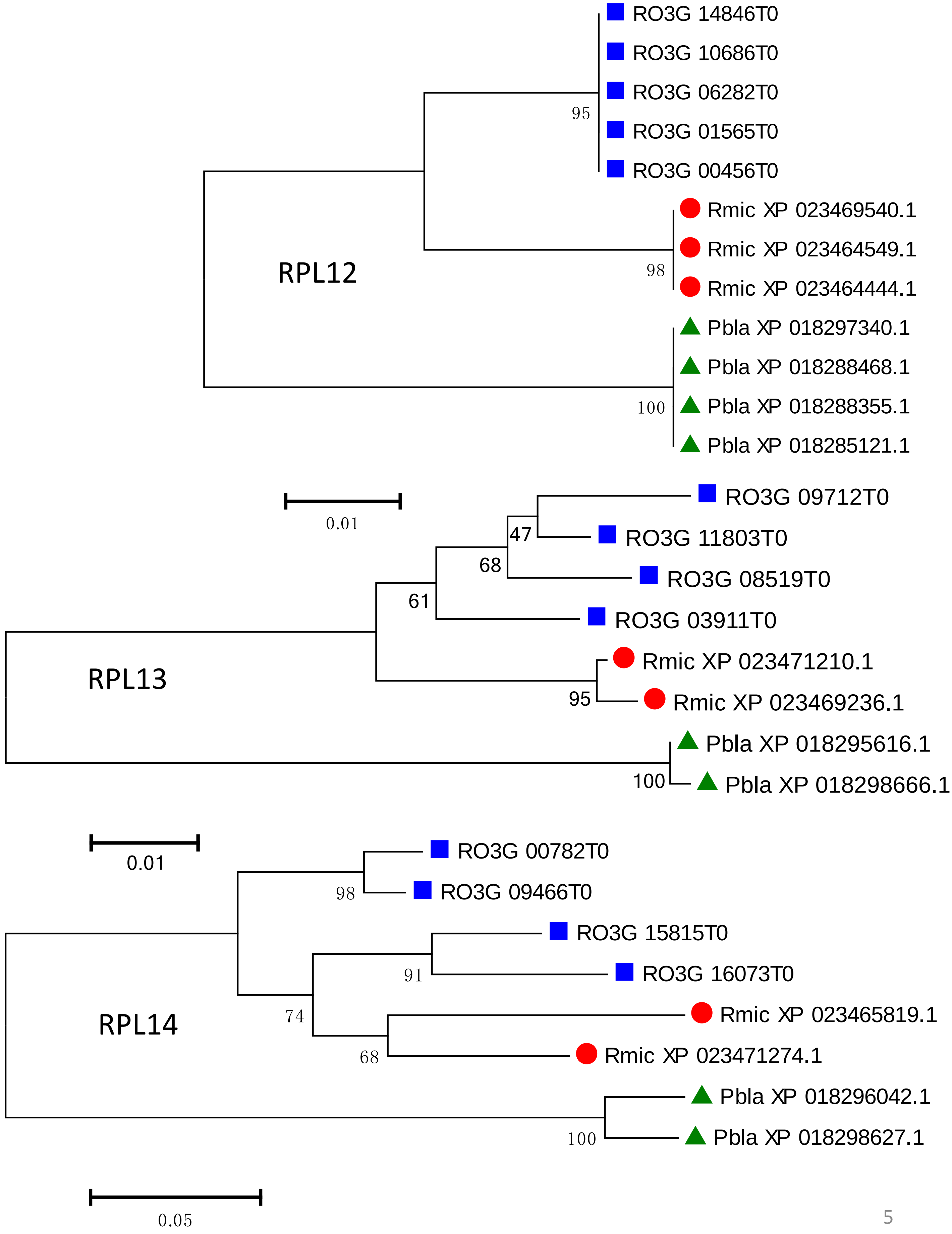
RPL12
RO3G 14846T0
RO3G 10686T0
RO3G 06282T0
RO3G 01565T0
RO3G 00456T0
Rmic XP 023469540.1
Rmic XP 023464549.1
Rmic XP 023464444.1
Pbla XP 018297340.1
Pbla XP 018288468.1
Pbla XP 018288355.1
Pbla XP 018285121.1
95
98
100
0.01
RPL13
RO3G 09712T0
RO3G 11803T0
RO3G 08519T0
RO3G 03911T0
Rmic XP 023471210.1
Rmic XP 023469236.1
Pbla XP 018295616.1
Pbla XP 018298666.1
47
68
61
95
100
0.01
RPL14
RO3G 00782T0
RO3G 09466T0
RO3G 15815T0
RO3G 16073T0
Rmic XP 023465819.1
Rmic XP 023471274.1
Pbla XP 018296042.1
Pbla XP 018298627.1
98
91
74
68
100
0.05

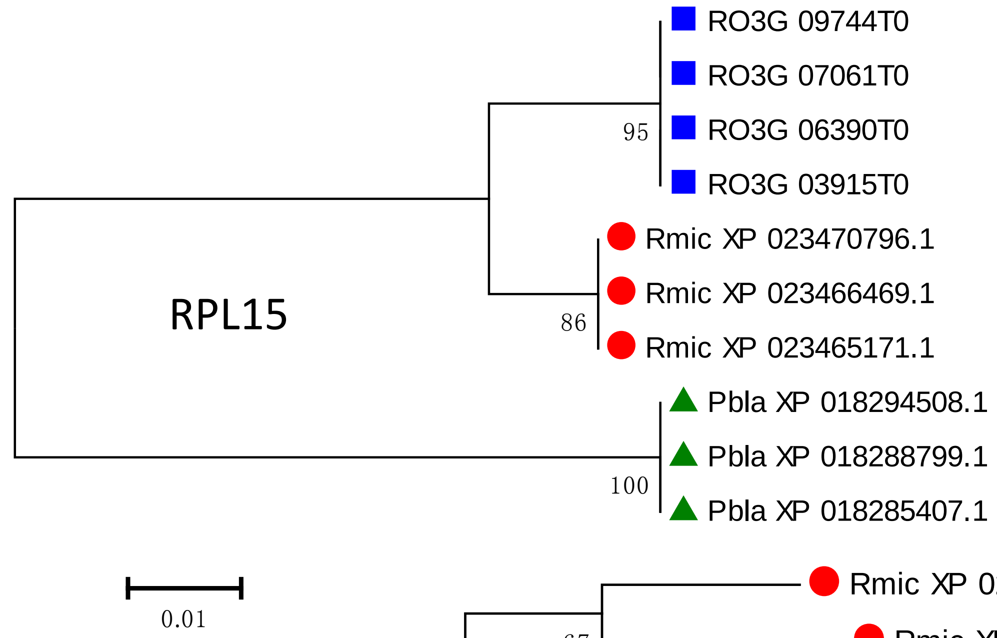
RO3G 09744T0
RO3G 07061T0
RO3G 06390T0
RO3G 03915T0
95
Rmic XP 023470796.1
Rmic XP 023466469.1
Rmic XP 023465171.1
86
RPL15
Pbla XP 018294508.1
Pbla XP 018288799.1
Pbla XP 018285407.1
100
0.01

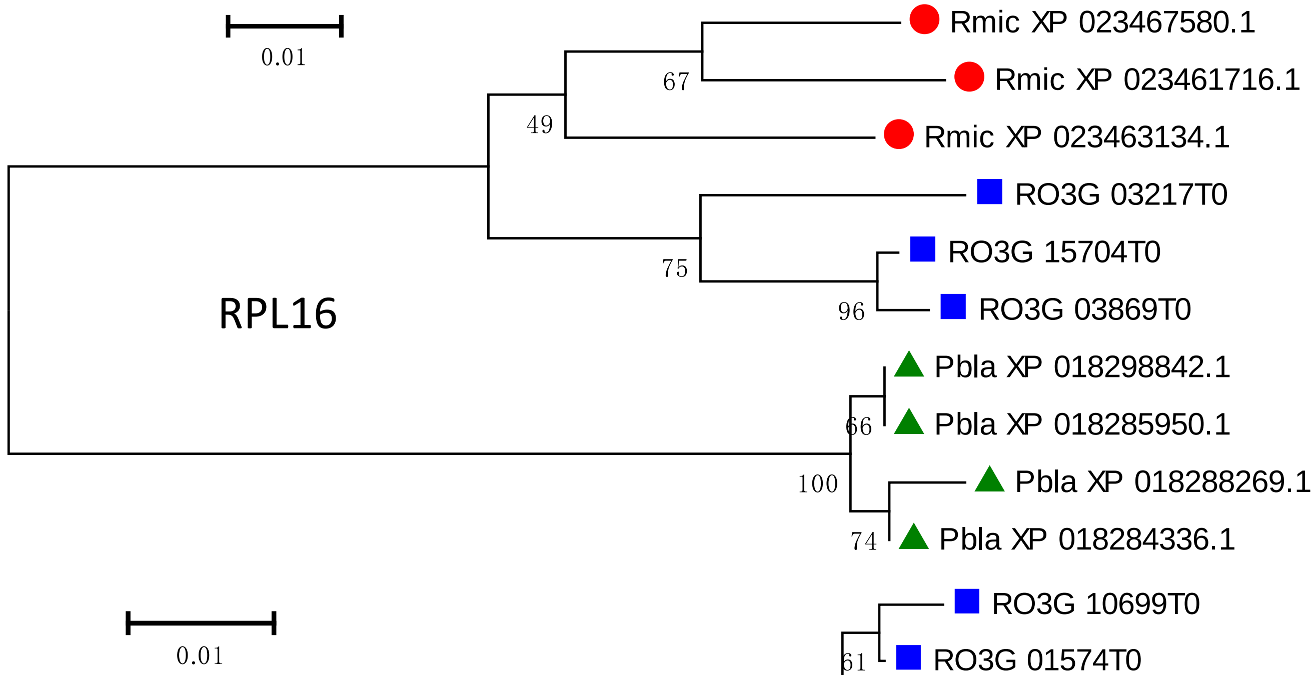
Rmic XP 023467580.1
Rmic XP 023461716.1
67
49
Rmic XP 023463134.1
RO3G 03217T0
RO3G 15704T0
75
RO3G 03869T0
96
RPL16
Pbla XP 018298842.1
Pbla XP 018285950.1
66
100
Pbla XP 018288269.1
Pbla XP 018284336.1
74
0.01

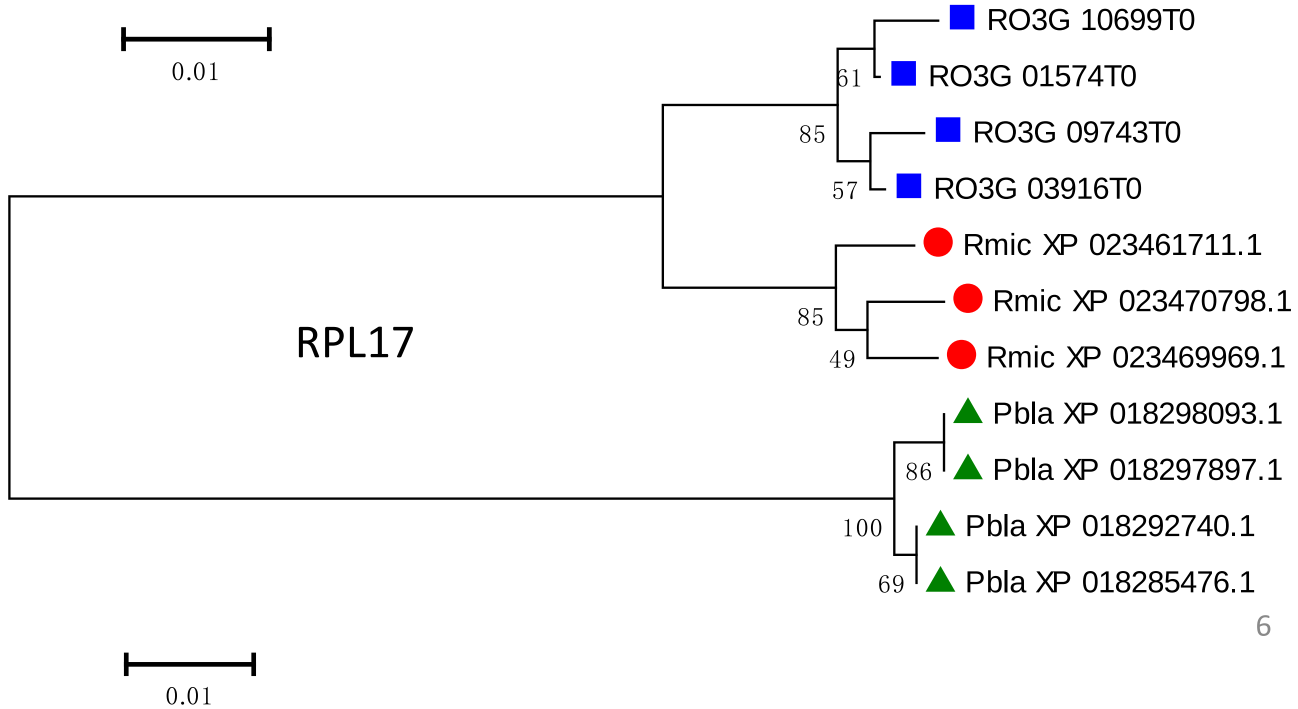
RO3G 10699T0
RO3G 01574T0
61
RO3G 09743T0
85
RO3G 03916T0
57
Rmic XP 023461711.1
Rmic XP 023470798.1
85
RPL17
Rmic XP 023469969.1
49
Pbla XP 018298093.1
Pbla XP 018297897.1
86
Pbla XP 018292740.1
100
Pbla XP 018285476.1
69
0.01

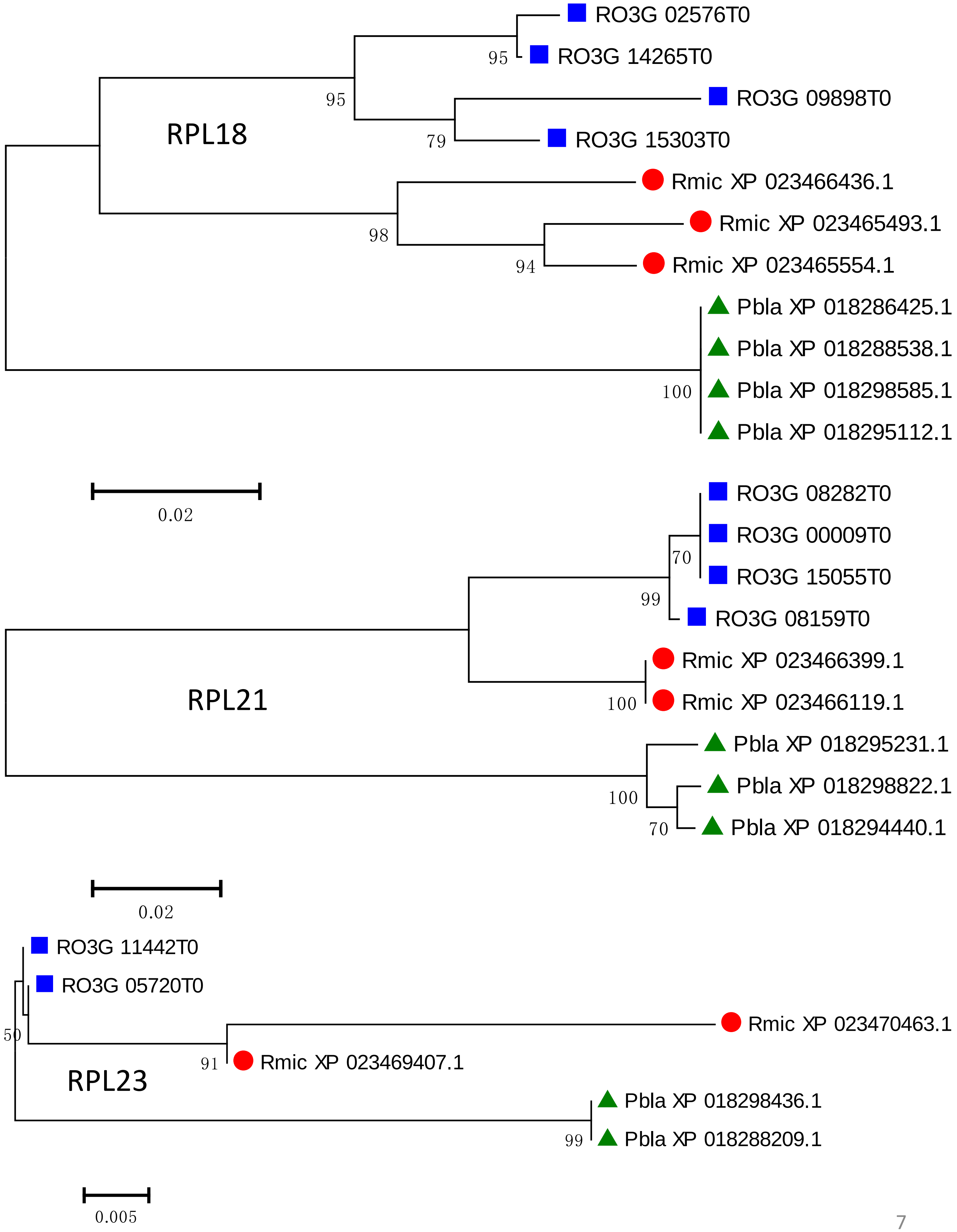
RO3G 02576T0
95
RO3G 14265T0
95
RO3G 09898T0
RPL18
79
RO3G 15303T0
Rmic XP 023466436.1
98
Rmic XP 023465493.1
94
Rmic XP 023465554.1
Pbla XP 018286425.1
Pbla XP 018288538.1
100
Pbla XP 018298585.1
Pbla XP 018295112.1
0.02
RO3G 08282T0
RO3G 00009T0
70
RO3G 15055T0
99
RO3G 08159T0
Rmic XP 023466399.1
RPL21
100
Rmic XP 023466119.1
Pbla XP 018295231.1
100
Pbla XP 018298822.1
70
Pbla XP 018294440.1
0.02
RO3G 11442T0
RO3G 05720T0
Rmic XP 023470463.1
50
91
Rmic XP 023469407.1
RPL23
Pbla XP 018298436.1
99
Pbla XP 018288209.1
0.005

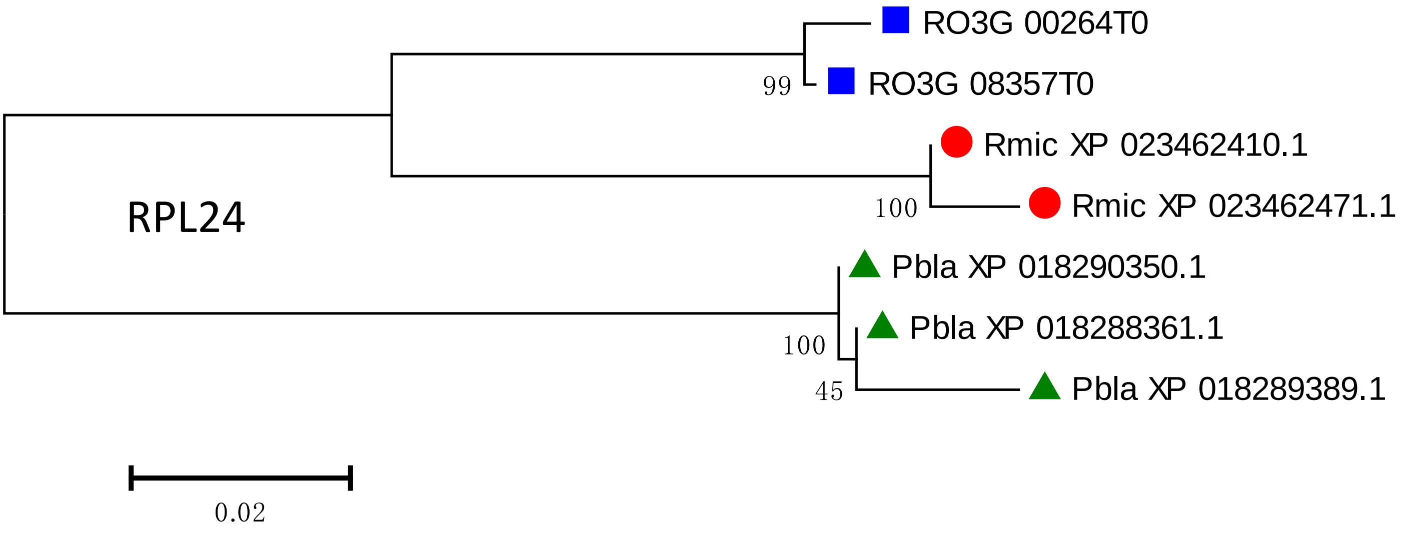
RO3G 00264T0
99
RO3G 08357T0
Rmic XP 023462410.1
100
Rmic XP 023462471.1
RPL24
Pbla XP 018290350.1
Pbla XP 018288361.1
100
45
Pbla XP 018289389.1
0.02

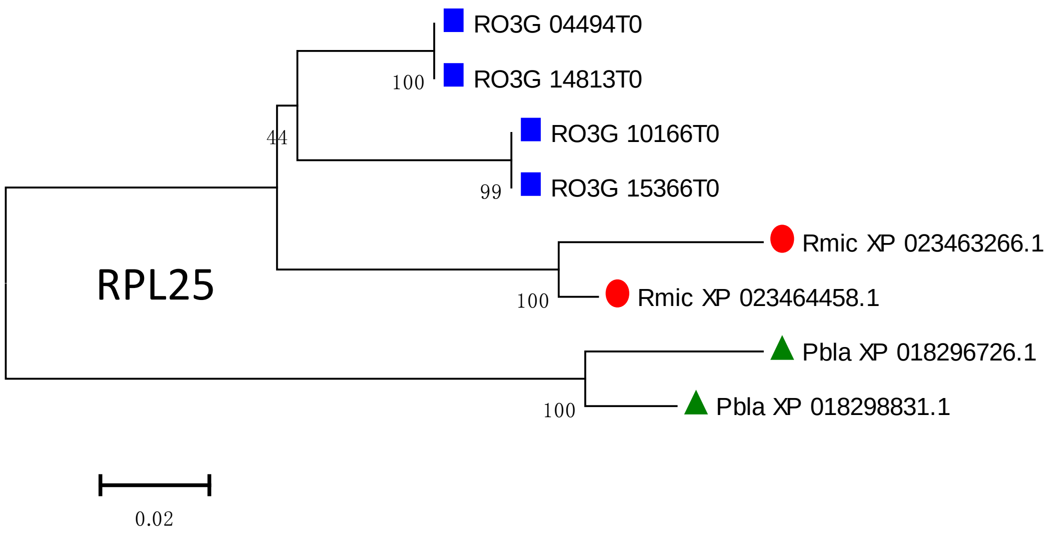
RO3G 04494T0
100
RO3G 14813T0
44
RO3G 10166T0
99
RO3G 15366T0
Rmic XP 023463266.1
RPL25
100
Rmic XP 023464458.1
Pbla XP 018296726.1
100
Pbla XP 018298831.1
0.02

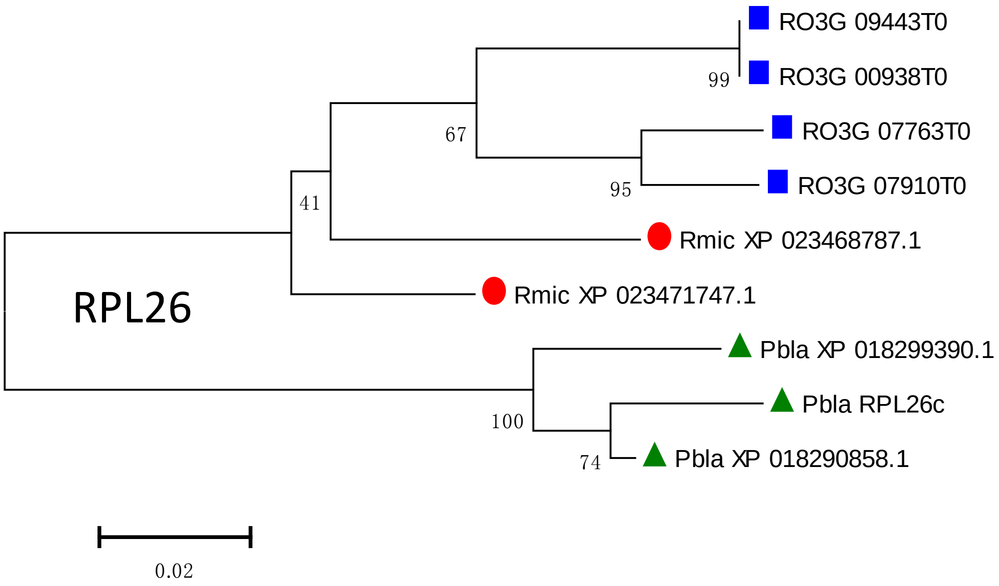
RO3G 09443T0
99
RO3G 00938T0
67
RO3G 07763T0
95
RO3G 07910T0
41
Rmic XP 023468787.1
Rmic XP 023471747.1
RPL26
Pbla XP 018299390.1
Pbla RPL26c
100
74
Pbla XP 018290858.1
0.02

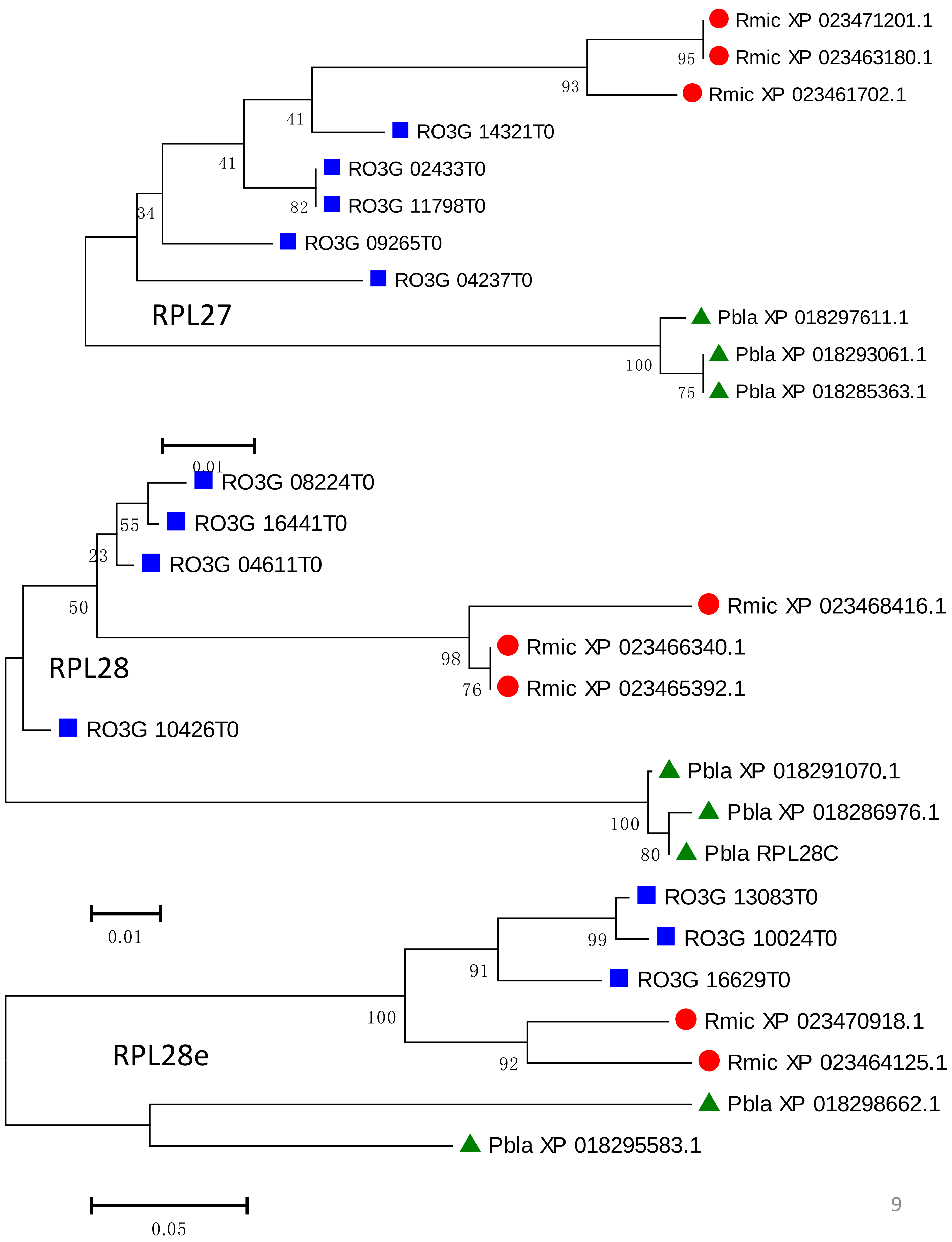
Rmic XP 023471201.1
Rmic XP 023463180.1
Rmic XP 023461702.1
RO3G 14321T0
RO3G 02433T0
RO3G 11798T0
RO3G 09265T0
RO3G 04237T0
RPL27
Pbla XP 018297611.1
Pbla XP 018293061.1
Pbla XP 018285363.1
95
93
41
41
82
34
100
75
0.01
RO3G 08224T0
RO3G 16441T0
RO3G 04611T0
Rmic XP 023468416.1
Rmic XP 023466340.1
Rmic XP 023465392.1
RPL28
RO3G 10426T0
Pbla XP 018291070.1
Pbla XP 018286976.1
Pbla RPL28C
55
23
50
98
76
100
80
0.01
RO3G 13083T0
RO3G 10024T0
RO3G 16629T0
Rmic XP 023470918.1
Rmic XP 023464125.1
Pbla XP 018298662.1
Pbla XP 018295583.1
RPL28e
99
91
100
92
0.05

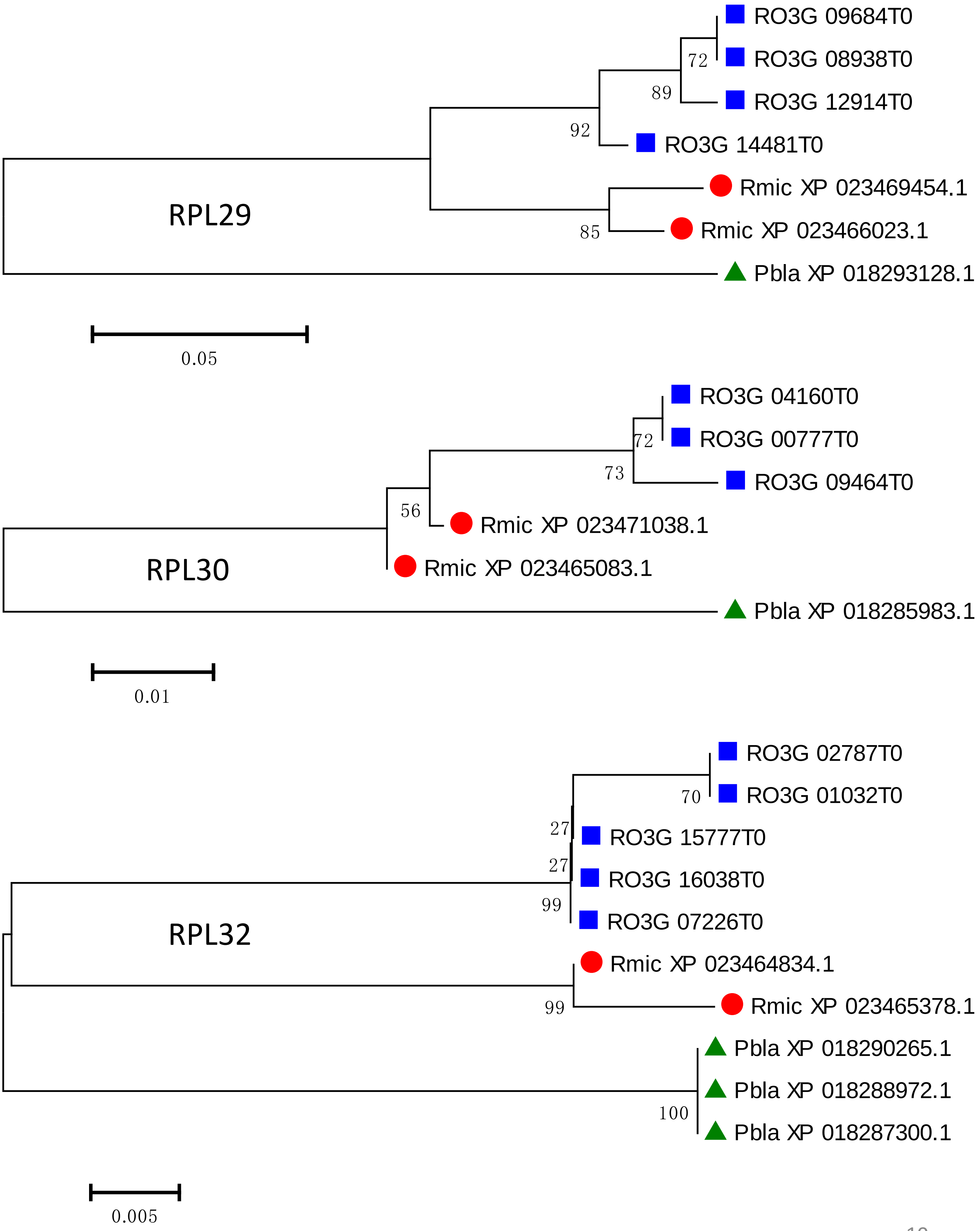
RO3G 09684T0
RO3G 08938T0
RO3G 12914T0
RO3G 14481T0
Rmic XP 023469454.1
Rmic XP 023466023.1
Pbla XP 018293128.1
72
89
92
85
RPL29
0.05
RO3G 04160T0
RO3G 00777T0
RO3G 09464T0
Rmic XP 023471038.1
Rmic XP 023465083.1
Pbla XP 018285983.1
72
73
56
RPL30
0.01
RO3G 02787T0
RO3G 01032T0
RO3G 15777T0
RO3G 16038T0
RO3G 07226T0
Rmic XP 023464834.1
Rmic XP 023465378.1
Pbla XP 018290265.1
Pbla XP 018288972.1
Pbla XP 018287300.1
70
27
27
99
99
100
RPL32
0.005

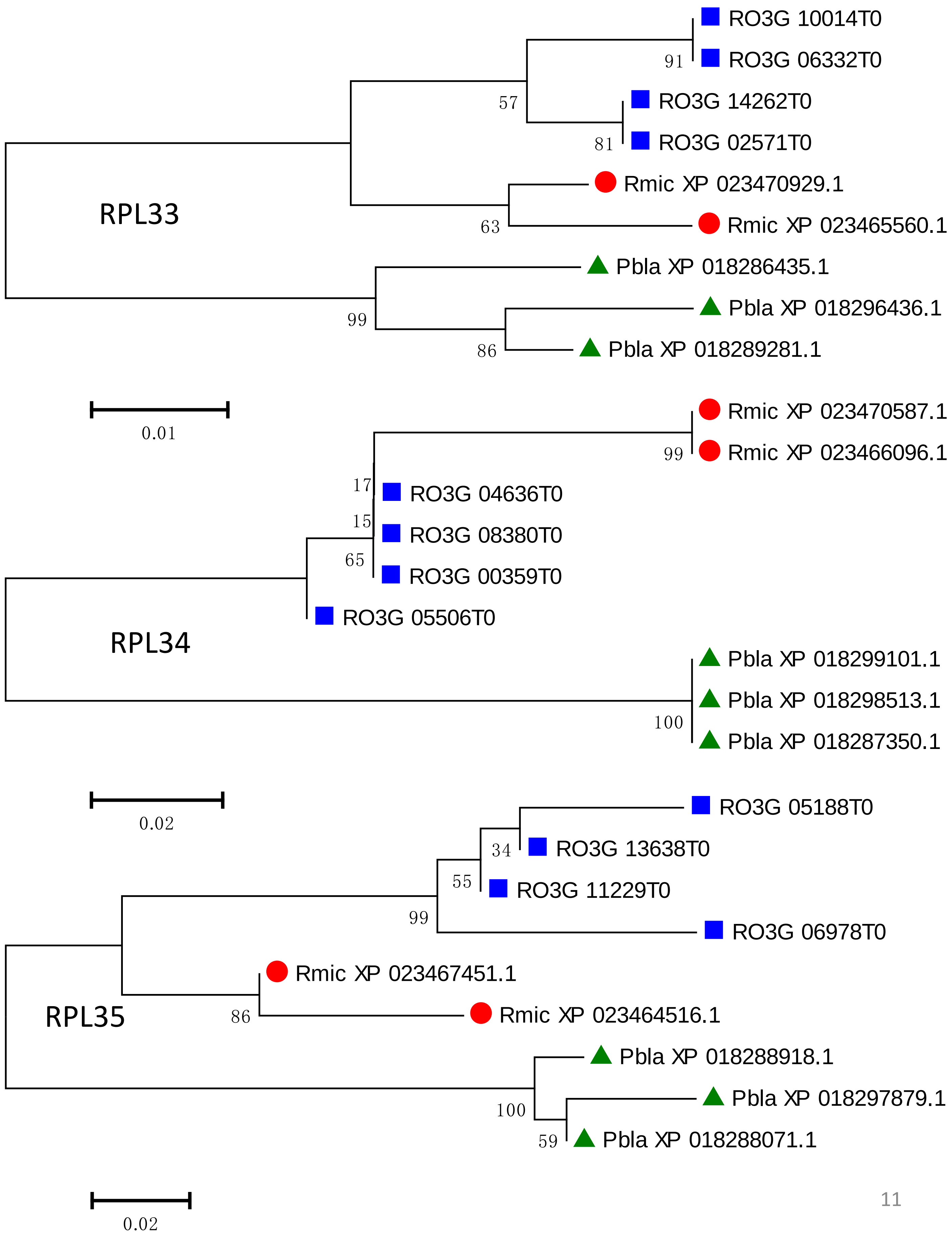
RO3G 10014T0
91
RO3G 06332T0
57
RO3G 14262T0
81
RO3G 02571T0
Rmic XP 023470929.1
RPL33
63
Rmic XP 023465560.1
Pbla XP 018286435.1
99
Pbla XP 018296436.1
86
Pbla XP 018289281.1
0.01
Rmic XP 023470587.1
99
Rmic XP 023466096.1
17
RO3G 04636T0
15
RO3G 08380T0
65
RO3G 00359T0
RO3G 05506T0
RPL34
Pbla XP 018299101.1
Pbla XP 018298513.1
100
Pbla XP 018287350.1
0.02
RO3G 05188T0
34
RO3G 13638T0
55
RO3G 11229T0
99
RO3G 06978T0
Rmic XP 023467451.1
RPL35
86
Rmic XP 023464516.1
Pbla XP 018288918.1
Pbla XP 018297879.1
100
59
Pbla XP 018288071.1
0.02

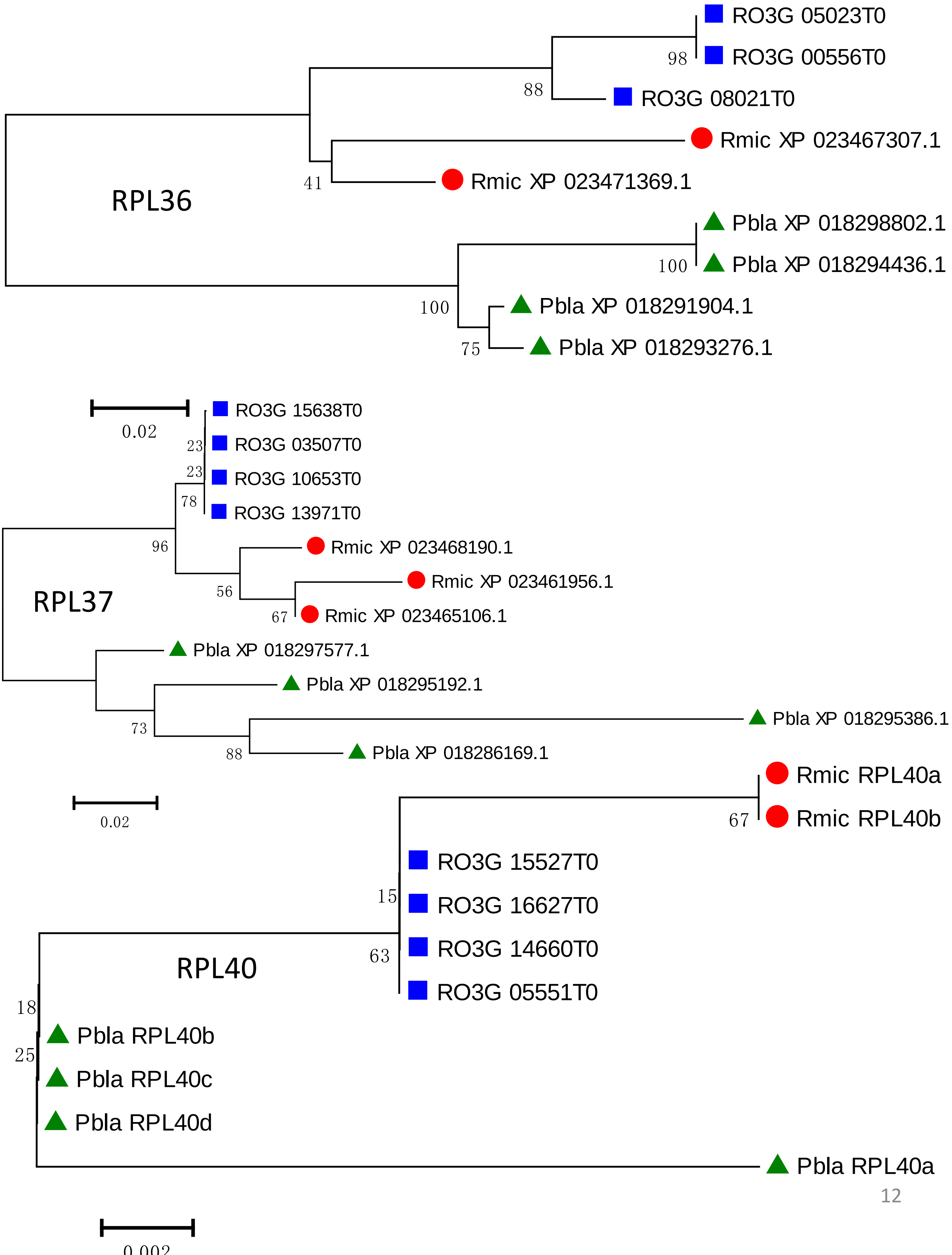
RO3G 05023T0
RO3G 00556T0
98
88
RO3G 08021T0
Rmic XP 023467307.1
41
Rmic XP 023471369.1
RPL36
Pbla XP 018298802.1
100
Pbla XP 018294436.1
100
Pbla XP 018291904.1
75
Pbla XP 018293276.1
0.02
RO3G 15638T0
23
RO3G 03507T0
23
RO3G 10653T0
78
RO3G 13971T0
96
Rmic XP 023468190.1
56
Rmic XP 023461956.1
67
Rmic XP 023465106.1
RPL37
Pbla XP 018297577.1
Pbla XP 018295192.1
73
Pbla XP 018295386.1
88
Pbla XP 018286169.1
0.02
Rmic RPL40a
67
Rmic RPL40b
RO3G 15527T0
15
RO3G 16627T0
63
RO3G 14660T0
RPL40
RO3G 05551T0
18
Pbla RPL40b
25
Pbla RPL40c
Pbla RPL40d
Pbla RPL40a
0.002

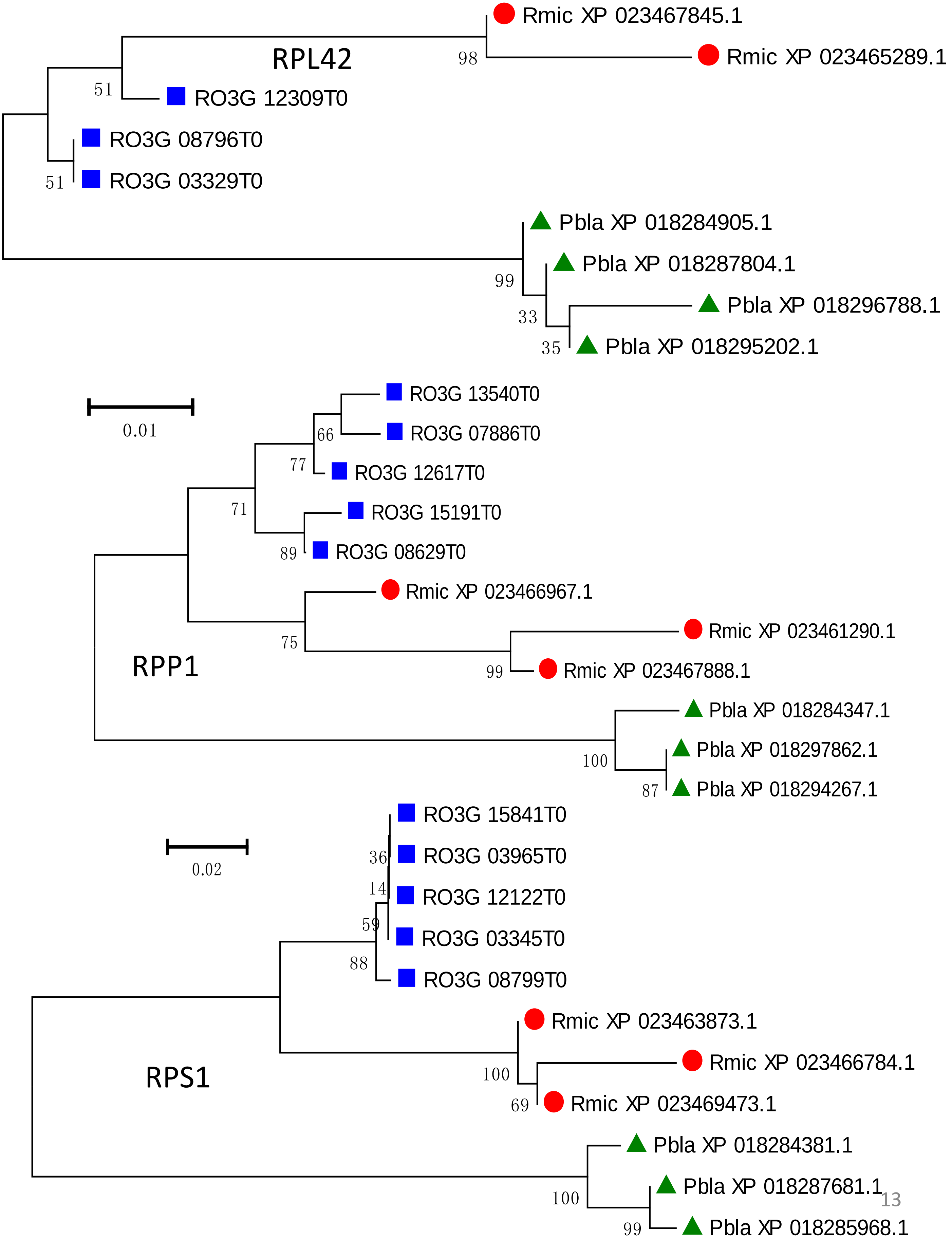
RPL42
Rmic XP 023467845.1
Rmic XP 023465289.1
RO3G 12309T0
RO3G 08796T0
RO3G 03329T0
Pbla XP 018284905.1
Pbla XP 018287804.1
Pbla XP 018296788.1
Pbla XP 018295202.1
98
51
51
99
33
35
0.01
RPP1
RO3G 13540T0
RO3G 07886T0
RO3G 12617T0
RO3G 15191T0
RO3G 08629T0
Rmic XP 023466967.1
Rmic XP 023461290.1
Rmic XP 023467888.1
Pbla XP 018284347.1
Pbla XP 018297862.1
Pbla XP 018294267.1
66
77
71
89
75
99
100
87
0.02
RPS1
RO3G 15841T0
RO3G 03965T0
RO3G 12122T0
RO3G 03345T0
RO3G 08799T0
Rmic XP 023463873.1
Rmic XP 023466784.1
Rmic XP 023469473.1
Pbla XP 018284381.1
Pbla XP 018287681.1
Pbla XP 018285968.1
36
14
59
88
100
69
100
99

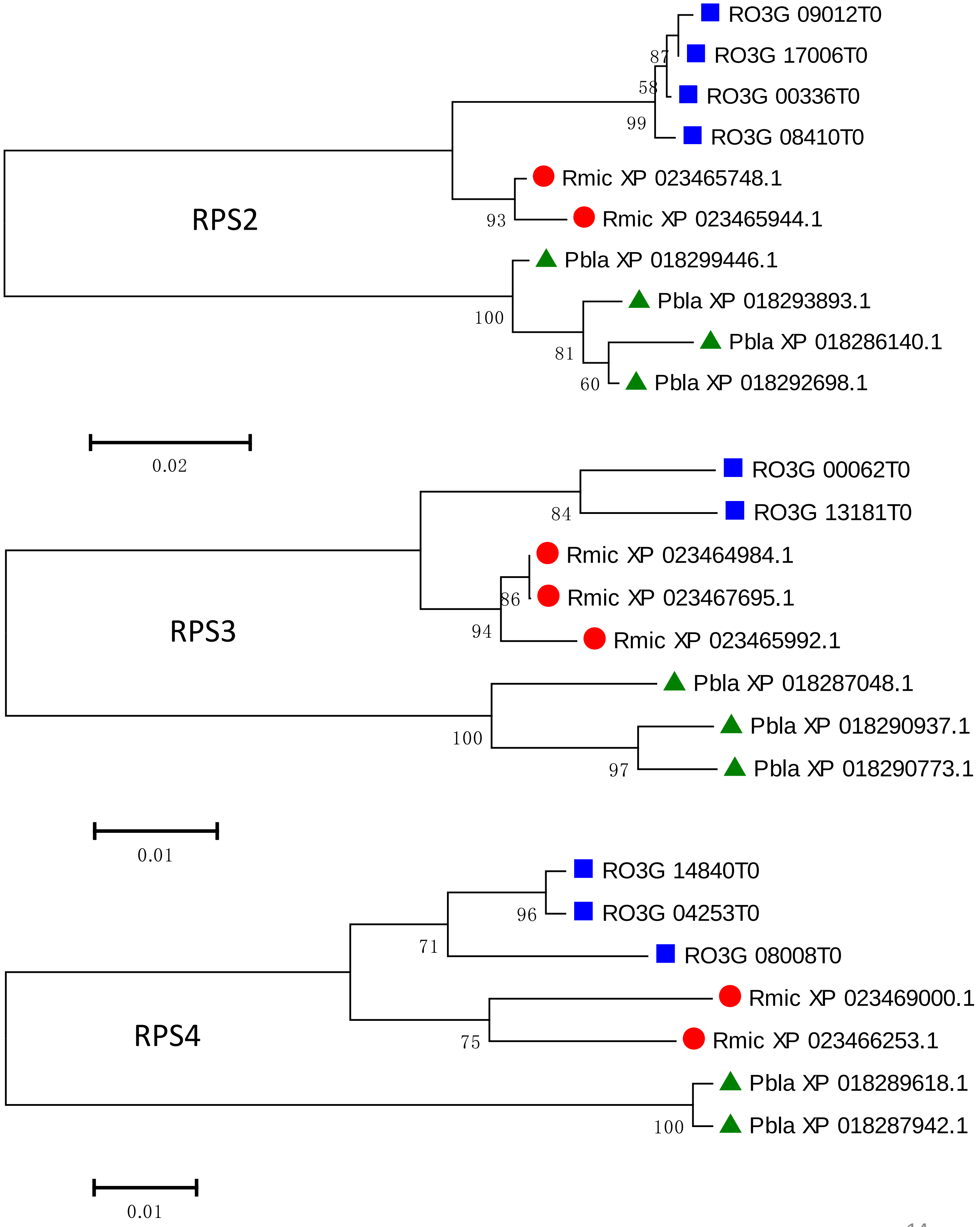
RO3G 09012T0
RO3G 17006T0
RO3G 00336T0
RO3G 08410T0
87
58
99
Rmic XP 023465748.1
Rmic XP 023465944.1
93
RPS2
Pbla XP 018299446.1
Pbla XP 018293893.1
Pbla XP 018286140.1
Pbla XP 018292698.1
100
81
60
0.02
RO3G 00062T0
RO3G 13181T0
84
Rmic XP 023464984.1
Rmic XP 023467695.1
86
94
Rmic XP 023465992.1
RPS3
Pbla XP 018287048.1
Pbla XP 018290937.1
Pbla XP 018290773.1
100
97
0.01
RO3G 14840T0
RO3G 04253T0
96
71
RO3G 08008T0
Rmic XP 023469000.1
Rmic XP 023466253.1
75
RPS4
Pbla XP 018289618.1
Pbla XP 018287942.1
100
0.01

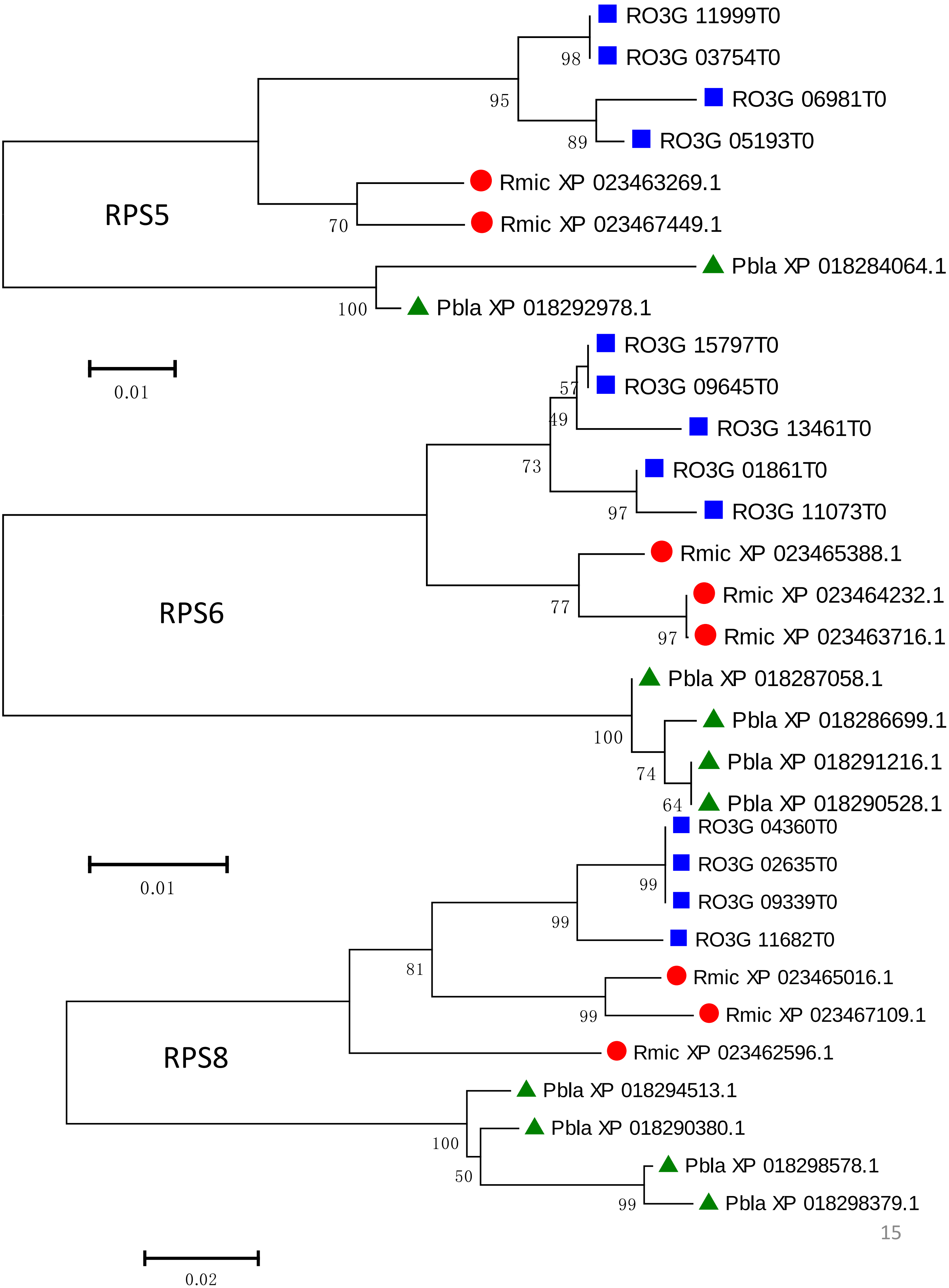
RO3G 11999T0
RO3G 03754T0
RO3G 06981T0
RO3G 05193T0
Rmic XP 023463269.1
Rmic XP 023467449.1
Pbla XP 018284064.1
Pbla XP 018292978.1
98
95
89
70
100
RPS5
0.01
RO3G 15797T0
RO3G 09645T0
RO3G 13461T0
RO3G 01861T0
RO3G 11073T0
Rmic XP 023465388.1
Rmic XP 023464232.1
Rmic XP 023463716.1
Pbla XP 018287058.1
Pbla XP 018286699.1
Pbla XP 018291216.1
Pbla XP 018290528.1
57
49
73
97
77
97
100
74
64
RPS6
0.01
RO3G 04360T0
RO3G 02635T0
RO3G 09339T0
RO3G 11682T0
Rmic XP 023465016.1
Rmic XP 023467109.1
Rmic XP 023462596.1
Pbla XP 018294513.1
Pbla XP 018290380.1
Pbla XP 018298578.1
Pbla XP 018298379.1
99
99
81
99
100
50
99
RPS8
0.02

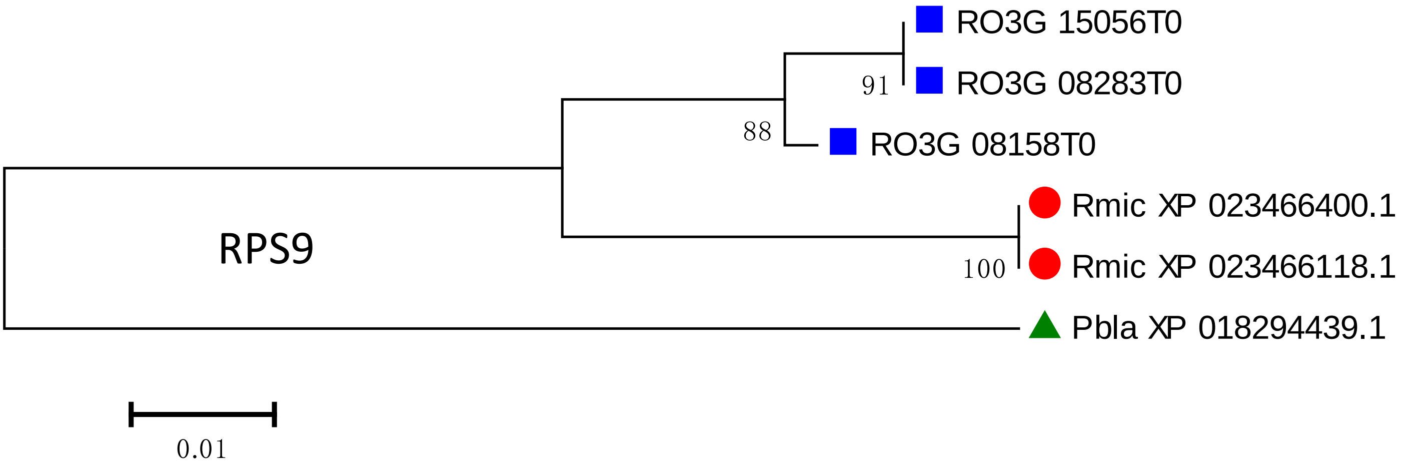
RO3G 15056T0
RO3G 08283T0
91
88
RO3G 08158T0
Rmic XP 023466400.1
Rmic XP 023466118.1
100
RPS9
Pbla XP 018294439.1
0.01

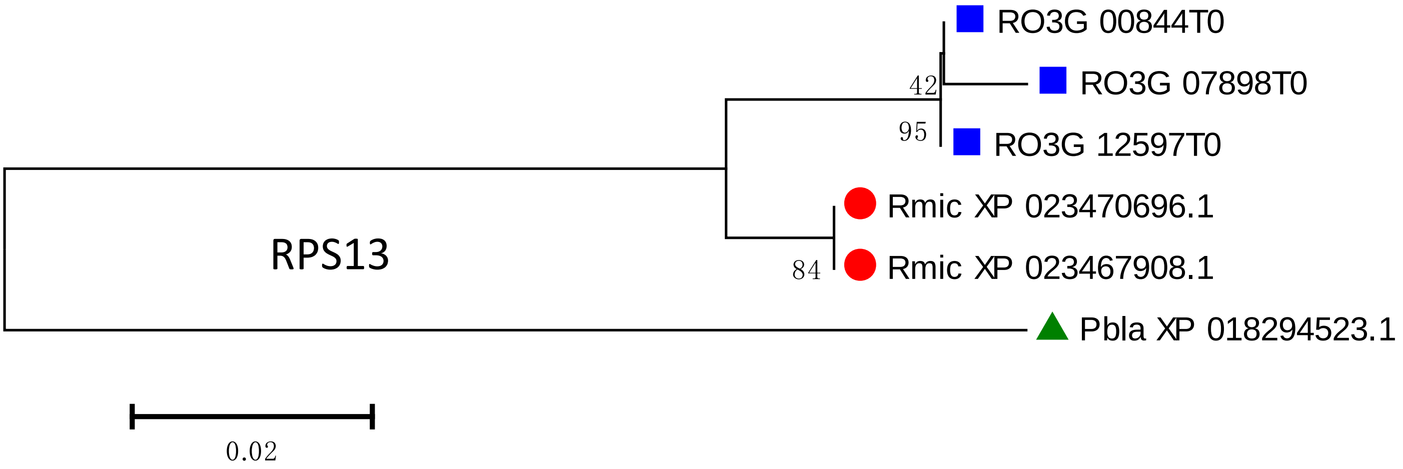
RO3G 00844T0
RO3G 07898T0
42
95
RO3G 12597T0
Rmic XP 023470696.1
Rmic XP 023467908.1
84
RPS13
Pbla XP 018294523.1
0.02

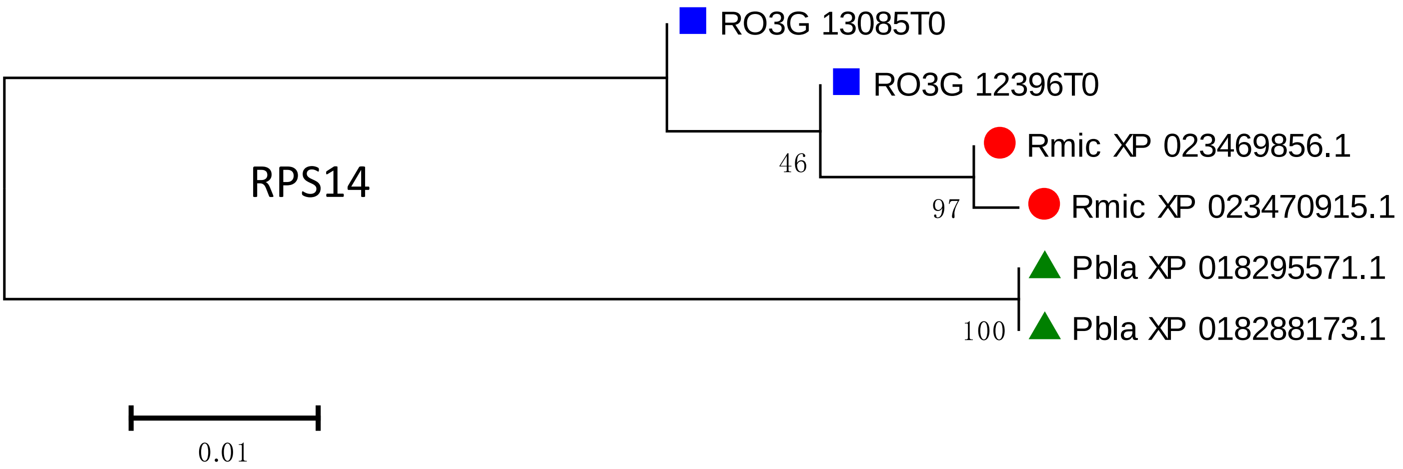
RO3G 13085T0
RO3G 12396T0
Rmic XP 023469856.1
46
RPS14
Rmic XP 023470915.1
97
Pbla XP 018295571.1
Pbla XP 018288173.1
100
0.01

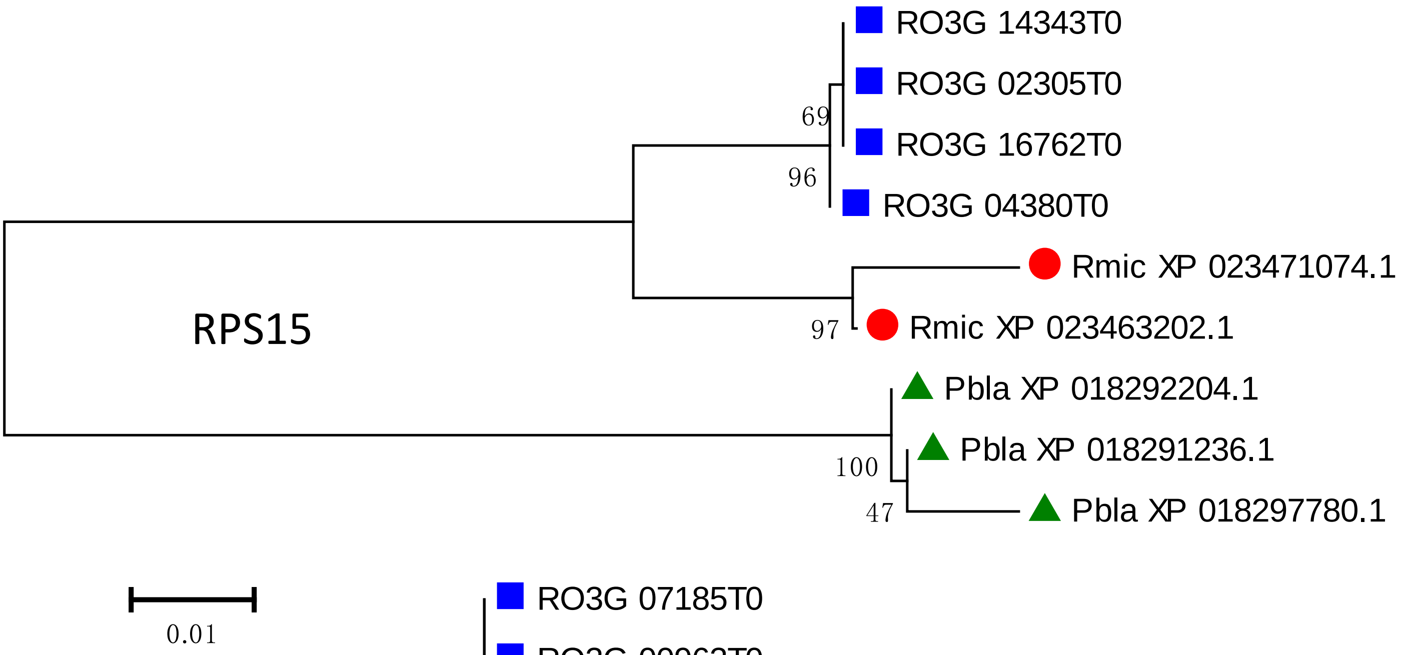
RO3G 14343T0
RO3G 02305T0
69
RO3G 16762T0
96
RO3G 04380T0
Rmic XP 023471074.1
RPS15
97
Rmic XP 023463202.1
Pbla XP 018292204.1
Pbla XP 018291236.1
100
47
Pbla XP 018297780.1
0.01

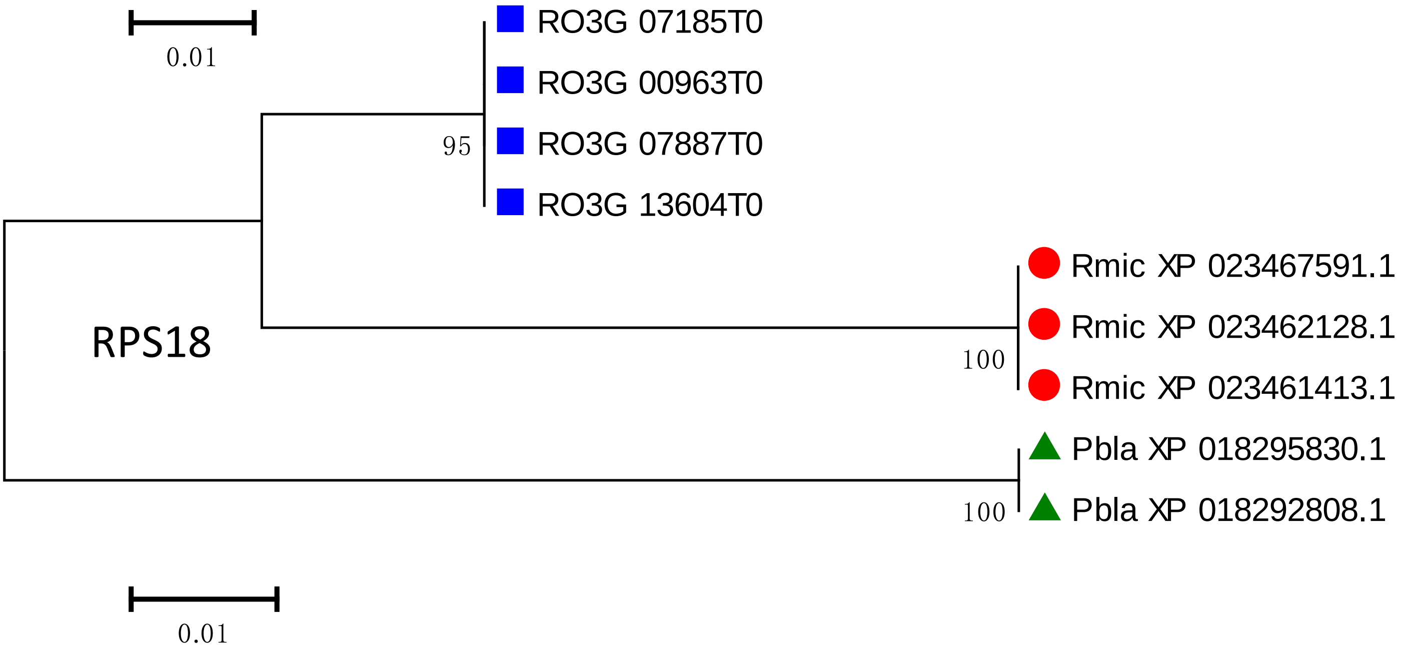
RO3G 07185T0
RO3G 00963T0
95
RO3G 07887T0
RO3G 13604T0
Rmic XP 023467591.1
Rmic XP 023462128.1
RPS18
100
Rmic XP 023461413.1
Pbla XP 018295830.1
100
Pbla XP 018292808.1
0.01

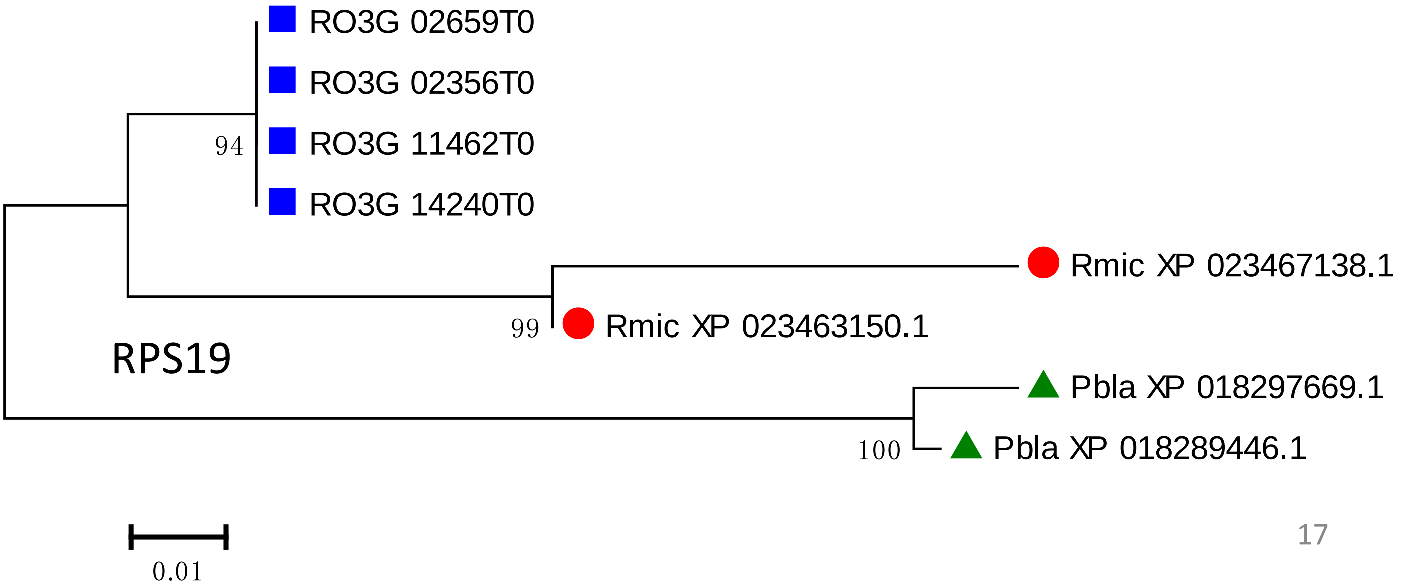
RO3G 02659T0
RO3G 02356T0
94
RO3G 11462T0
RO3G 14240T0
Rmic XP 023467138.1
99
Rmic XP 023463150.1
RPS19
Pbla XP 018297669.1
100
Pbla XP 018289446.1
0.01

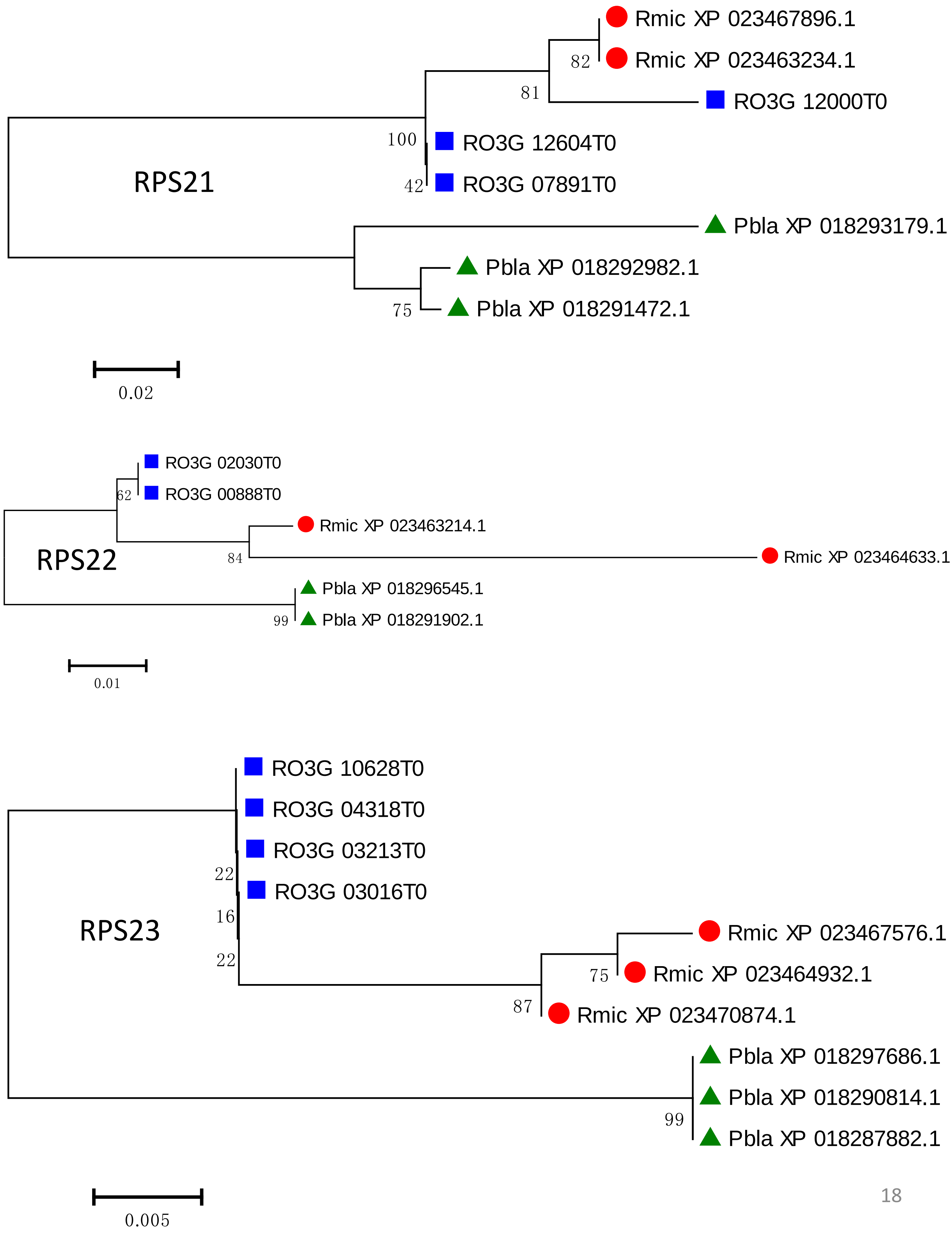

Rmic XP 023467896.1
82
Rmic XP 023463234.1
81
RO3G 12000T0
100
RO3G 12604T0
RPS21
42
RO3G 07891T0
Pbla XP 018293179.1
Pbla XP 018292982.1
75
Pbla XP 018291472.1
0.02
RO3G 02030T0
62
RO3G 00888T0
Rmic XP 023463214.1
RPS22
84
Rmic XP 023464633.1
Pbla XP 018296545.1
99
Pbla XP 018291902.1
0.01
RO3G 10628T0
RO3G 04318T0
RO3G 03213T0
22
RO3G 03016T0
16
RPS23
22
Rmic XP 023467576.1
75
Rmic XP 023464932.1
87
Rmic XP 023470874.1
Pbla XP 018297686.1
Pbla XP 018290814.1
99
Pbla XP 018287882.1
0.005

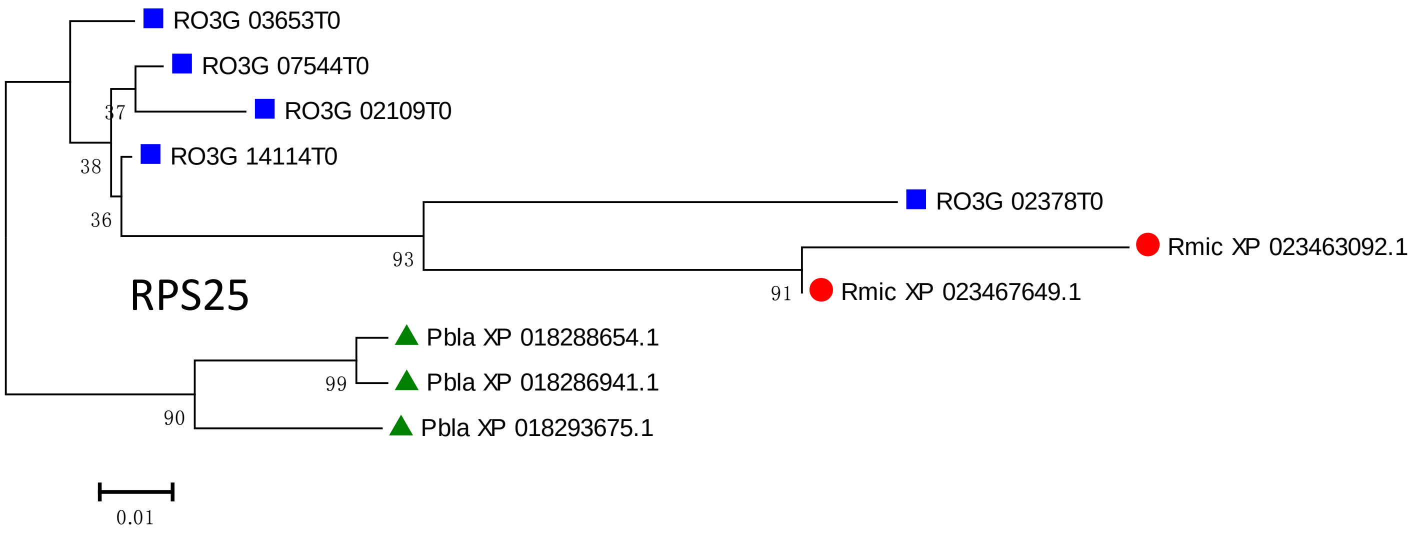
RO3G 03653T0
RO3G 07544T0
RO3G 02109T0
RO3G 14114T0
RO3G 02378T0
Rmic XP 023463092.1
Rmic XP 023467649.1
Pbla XP 018288654.1
Pbla XP 018286941.1
Pbla XP 018293675.1
37
38
36
93
91
99
90
RPS25
0.01

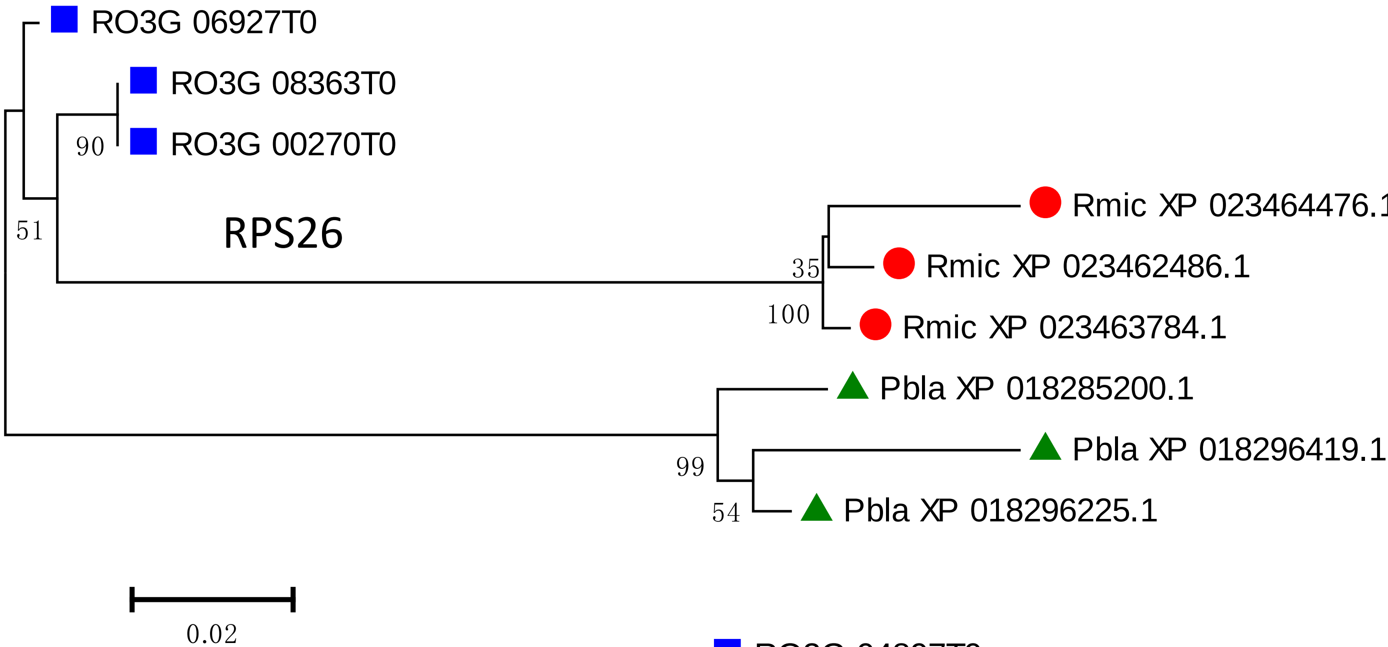
RO3G 06927T0
RO3G 08363T0
RO3G 00270T0
Rmic XP 023464476.
Rmic XP 023462486.1
Rmic XP 023463784.1
Pbla XP 018285200.1
Pbla XP 018296419.1
Pbla XP 018296225.1
90
51
35
100
99
54
RPS26
0.02

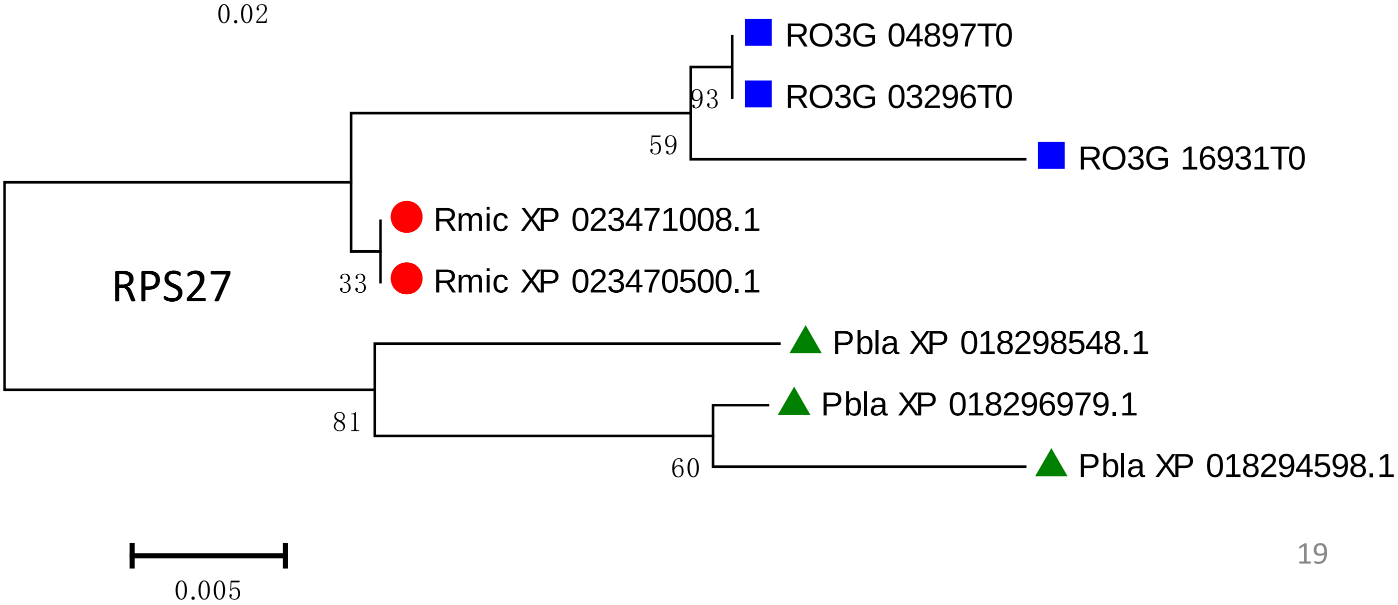
RO3G 04897T0
RO3G 03296T0
RO3G 16931T0
Rmic XP 023471008.1
Rmic XP 023470500.1
Pbla XP 018298548.1
Pbla XP 018296979.1
Pbla XP 018294598.1
93
59
33
81
60
RPS27
0.005

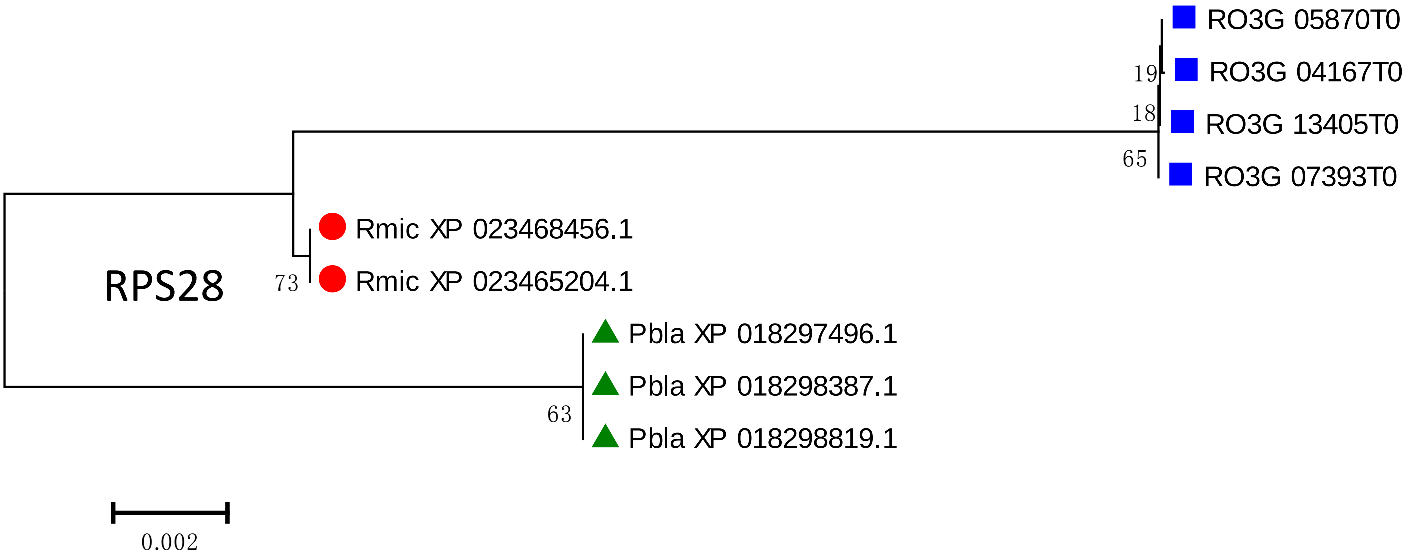
RO3G 05870T0
RO3G 04167T0
RO3G 13405T0
RO3G 07393T0
19
18
65
Rmic XP 023468456.1
Rmic XP 023465204.1
73
RPS28
Pbla XP 018297496.1
Pbla XP 018298387.1
Pbla XP 018298819.1
63
0.002

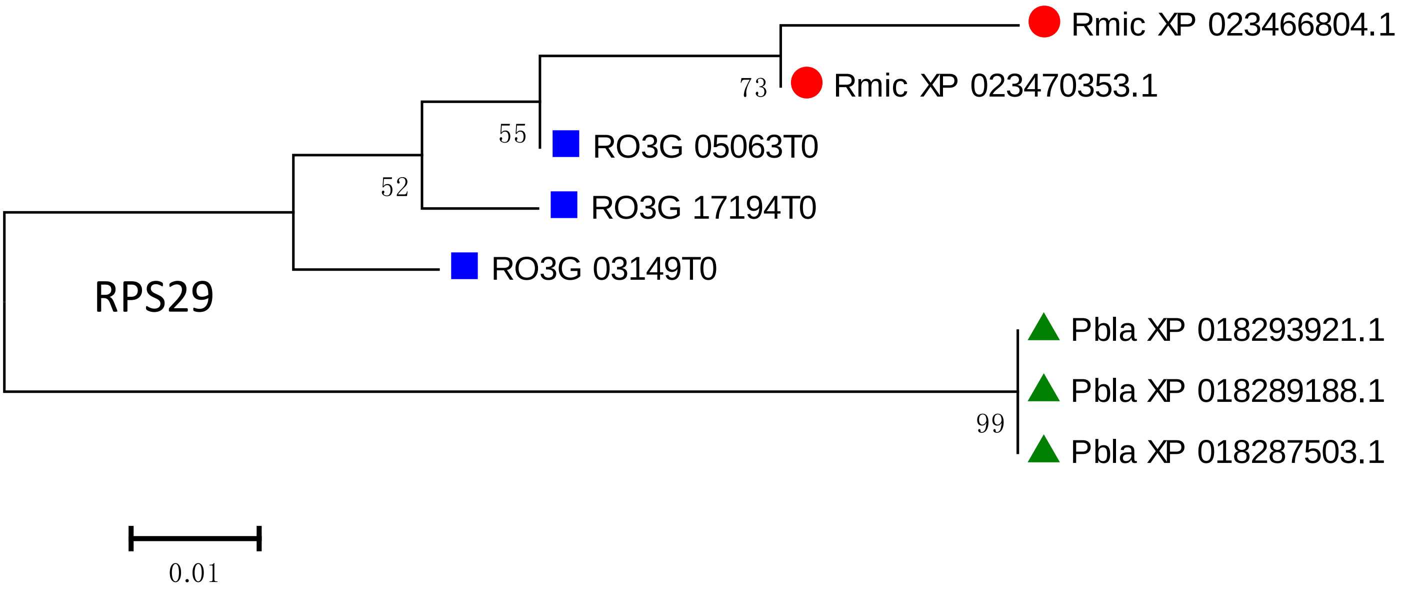
Rmic XP 023466804.1
Rmic XP 023470353.1
73
55
RO3G 05063T0
52
RO3G 17194T0
RO3G 03149T0
RPS29
Pbla XP 018293921.1
Pbla XP 018289188.1
Pbla XP 018287503.1
99
0.01

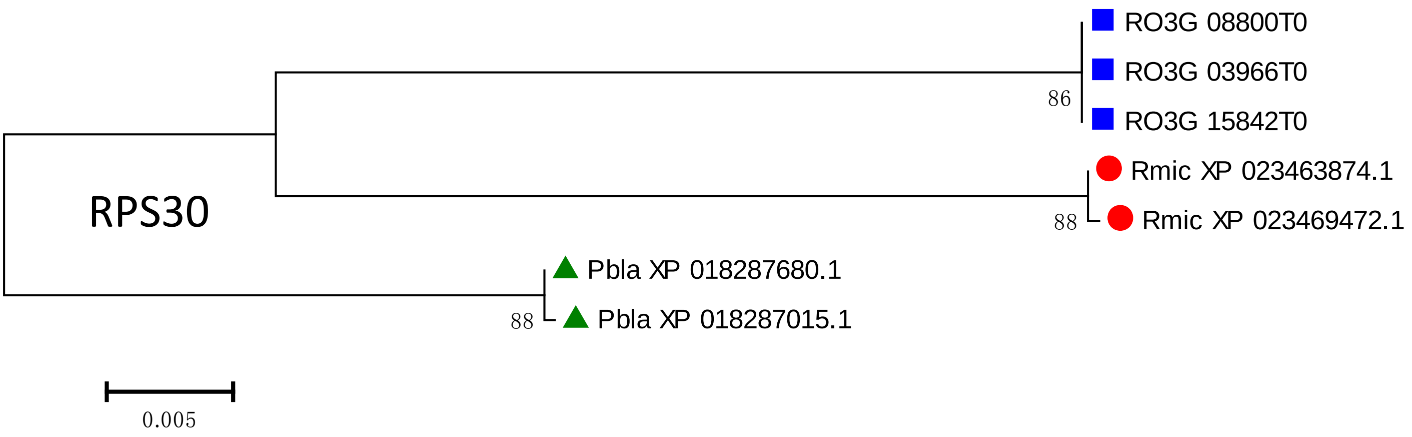
RO3G 08800T0
RO3G 03966T0
RO3G 15842T0
86
Rmic XP 023463874.1
Rmic XP 023469472.1
88
RPS30
Pbla XP 018287680.1
Pbla XP 018287015.1
88
0.005

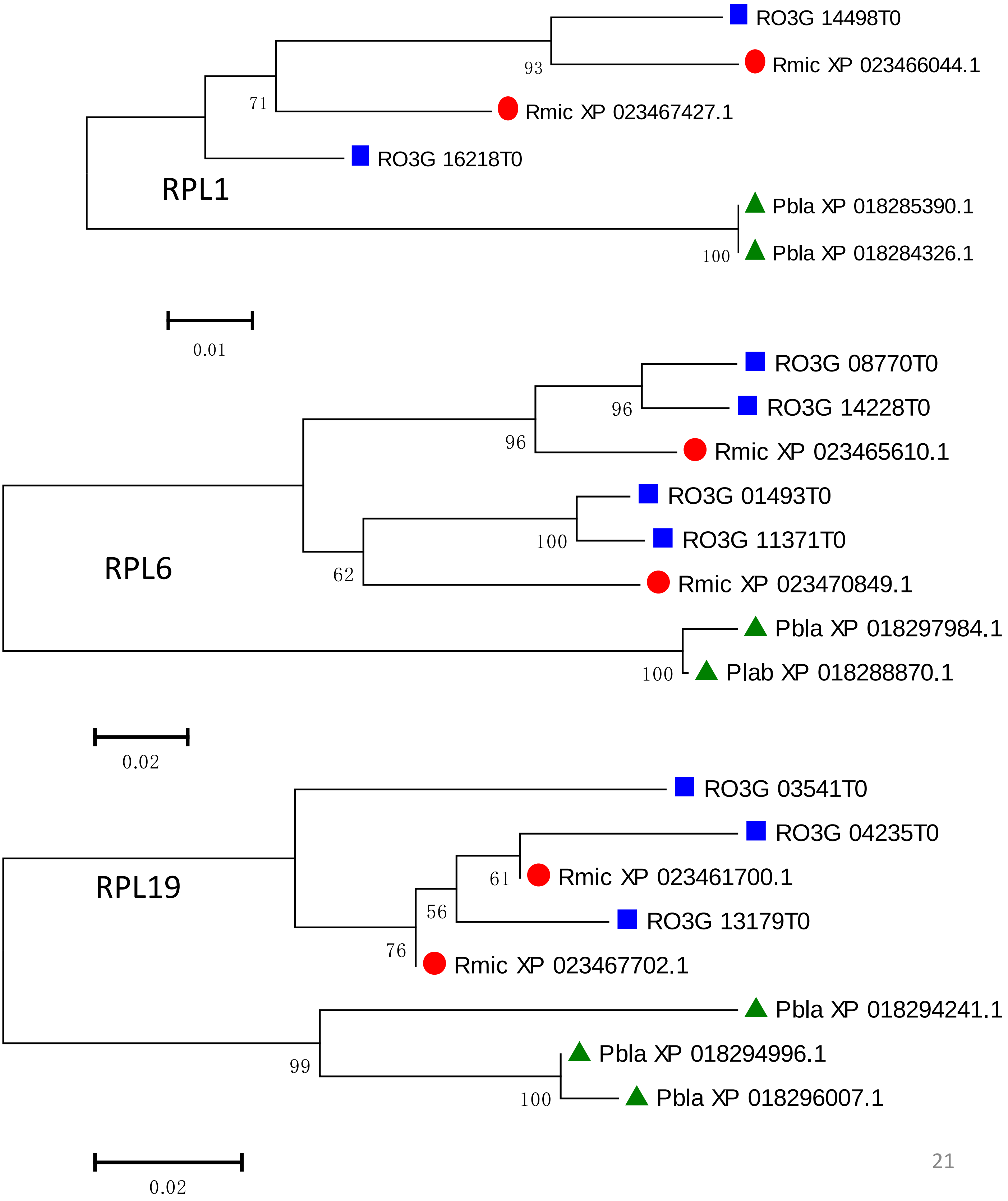
RO3G 14498T0
93
Rmic XP 023466044.1
71
Rmic XP 023467427.1
RO3G 16218T0
RPL1
Pbla XP 018285390.1
100
Pbla XP 018284326.1
0.01
RO3G 08770T0
96
RO3G 14228T0
96
Rmic XP 023465610.1
RO3G 01493T0
100
RO3G 11371T0
RPL6
62
Rmic XP 023470849.1
Pbla XP 018297984.1
100
Plab XP 018288870.1
0.02
RO3G 03541T0
RO3G 04235T0
RPL19
61
Rmic XP 023461700.1
56
RO3G 13179T0
76
Rmic XP 023467702.1
Pbla XP 018294241.1
99
Pbla XP 018294996.1
100
Pbla XP 018296007.1
0.02

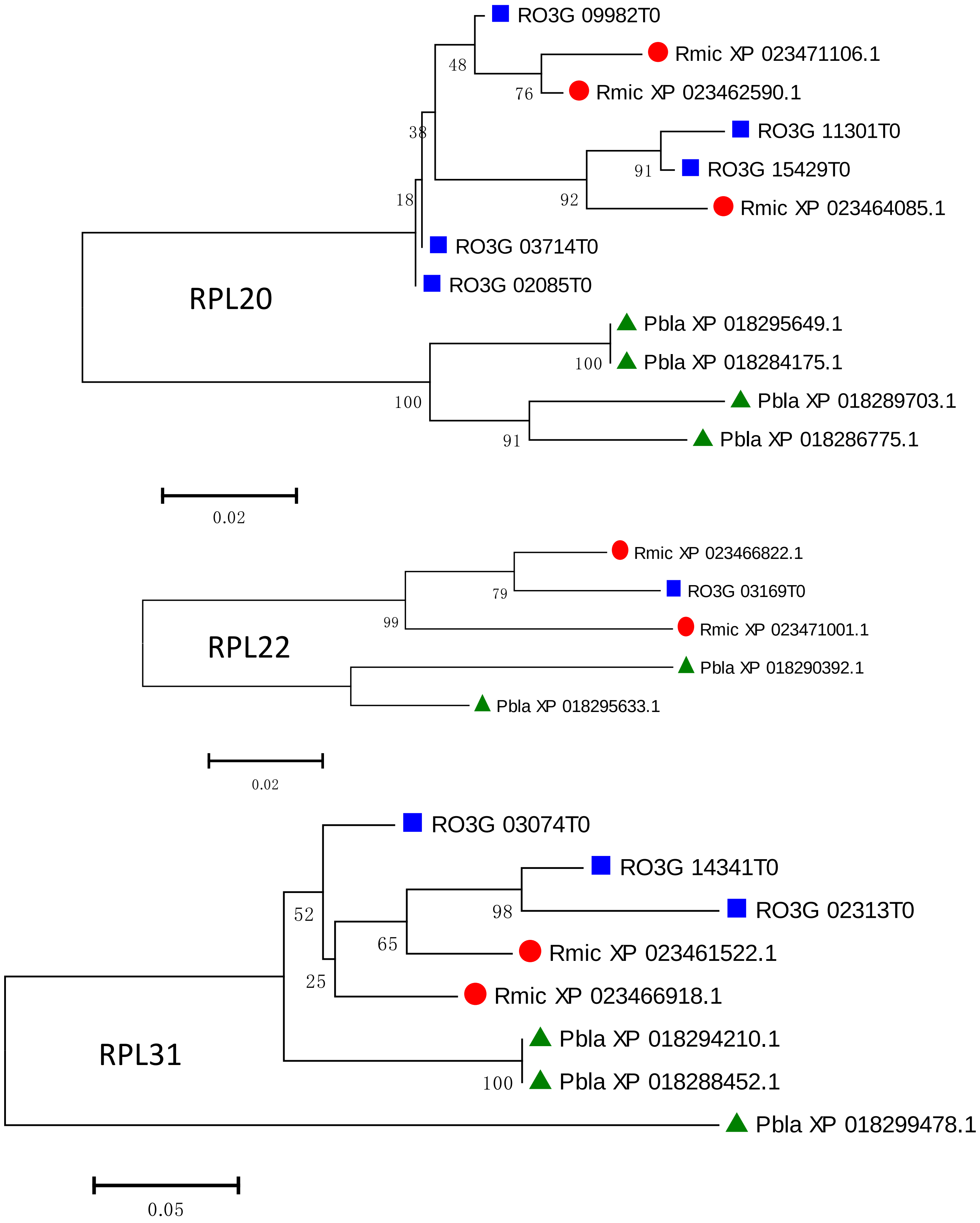
RO3G 09982T0
Rmic XP 023471106.1
48
76
Rmic XP 023462590.1
38
RO3G 11301T0
91
RO3G 15429T0
18
92
Rmic XP 023464085.1
RO3G 03714T0
RPL20
RO3G 02085T0
Pbla XP 018295649.1
100
Pbla XP 018284175.1
100
Pbla XP 018289703.1
91
Pbla XP 018286775.1
0.02
Rmic XP 023466822.1
79
RO3G 03169T0
99
Rmic XP 023471001.1
RPL22
Pbla XP 018290392.1
Pbla XP 018295633.1
0.02
RO3G 03074T0
RO3G 14341T0
52
98
RO3G 02313T0
65
Rmic XP 023461522.1
25
Rmic XP 023466918.1
Pbla XP 018294210.1
RPL31
100
Pbla XP 018288452.1
Pbla XP 018299478.1
0.05

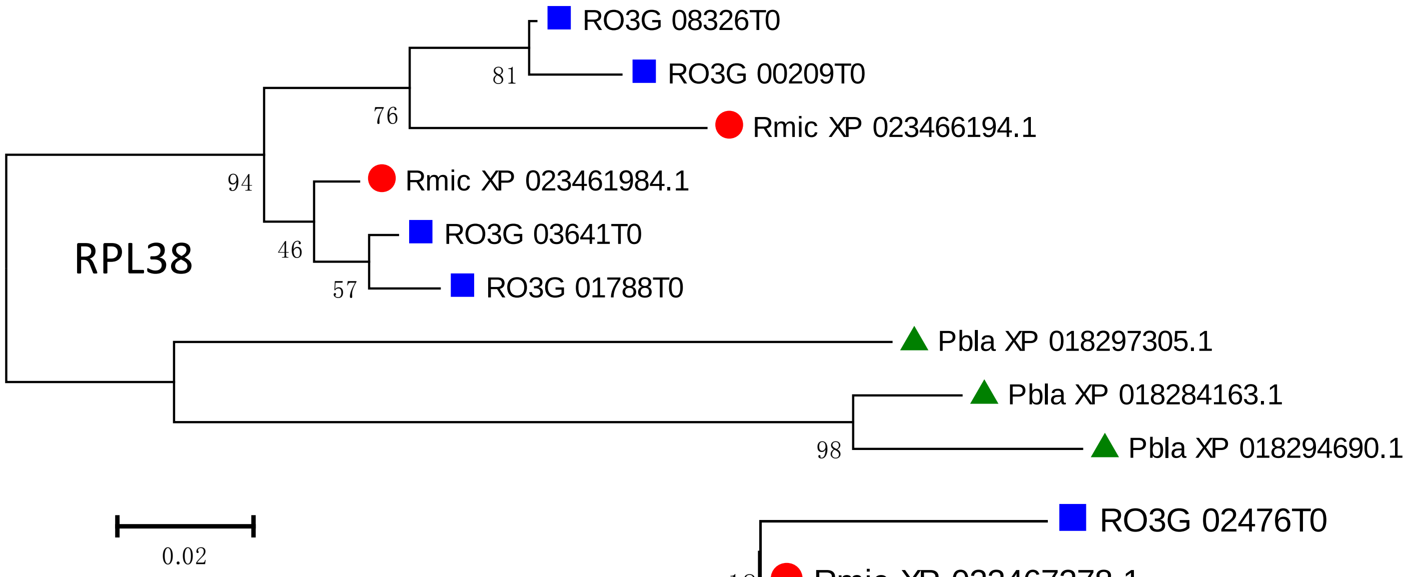
RPL38
RO3G 08326T0
RO3G 00209T0
Rmic XP 023466194.1
Rmic XP 023461984.1
RO3G 03641T0
RO3G 01788T0
Pbla XP 018297305.1
Pbla XP 018284163.1
Pbla XP 018294690.1
81
76
94
46
57
98
0.02

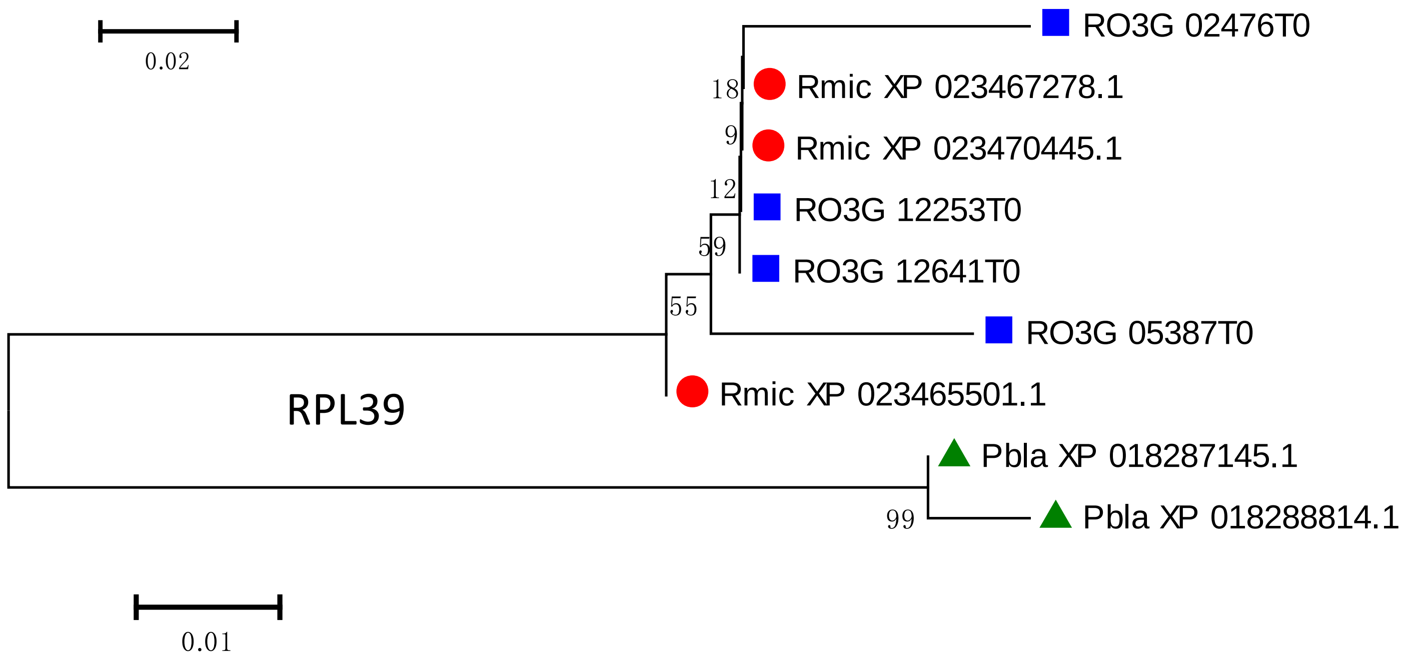
RPL39
RO3G 02476T0
Rmic XP 023467278.1
Rmic XP 023470445.1
RO3G 12253T0
RO3G 12641T0
RO3G 05387T0
Rmic XP 023465501.1
Pbla XP 018287145.1
Pbla XP 018288814.1
18
9
12
59
55
99
0.01

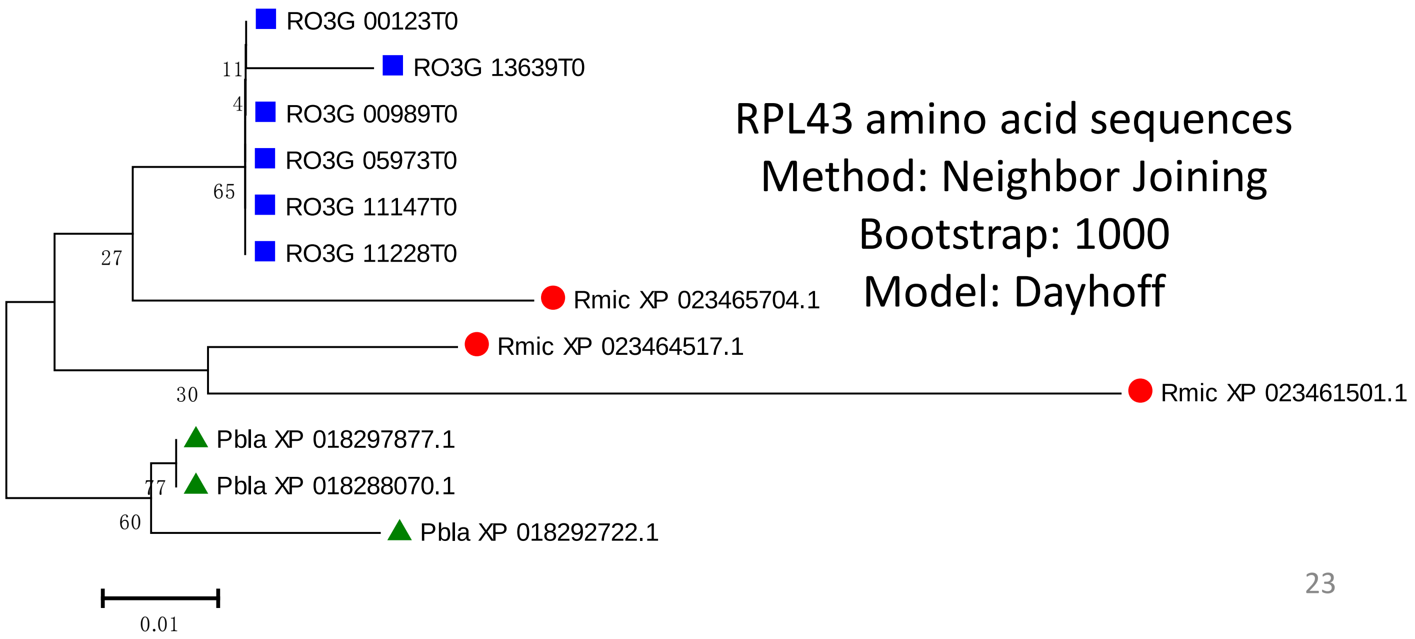
RO3G 00123T0
RO3G 13639T0
RO3G 00989T0
RO3G 05973T0
RO3G 11147T0
RO3G 11228T0
Rmic XP 023465704.1
Rmic XP 023464517.1
Rmic XP 023461501.1
Pbla XP 018297877.1
Pbla XP 018288070.1
Pbla XP 018292722.1
11
4
65
27
30
77
60
0.01
RPL43 amino acid sequences
Method: Neighbor Joining
Bootstrap: 1000
Model: Dayhoff

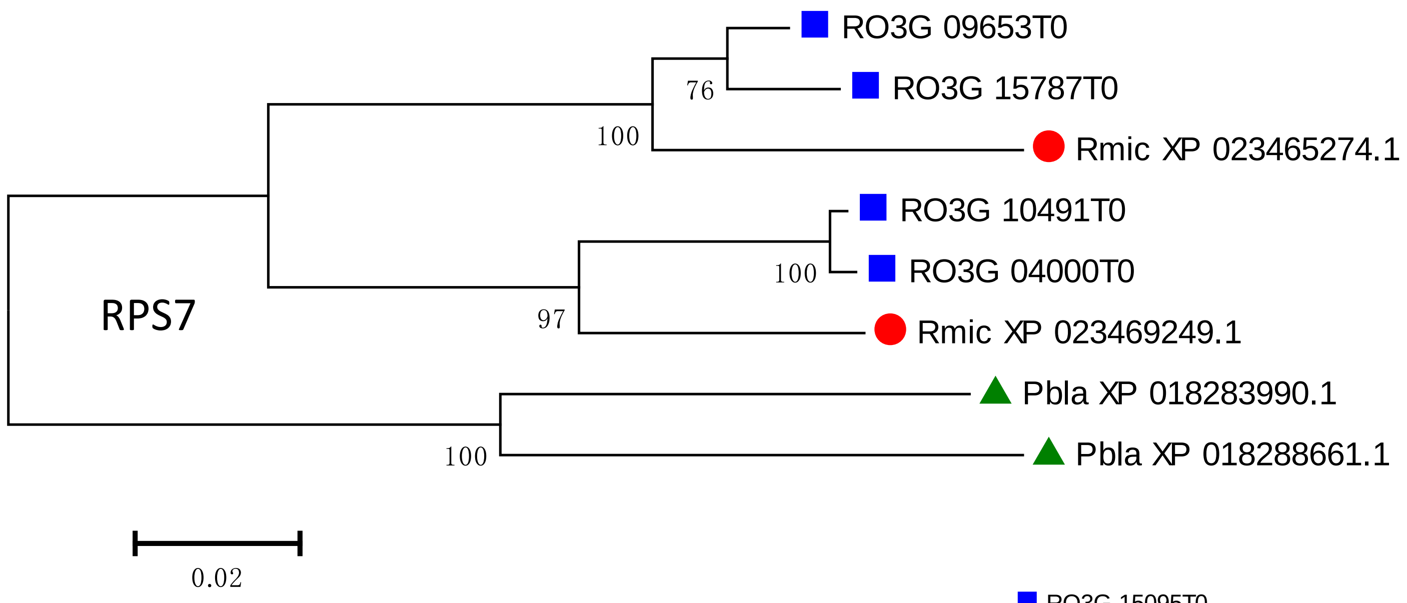
RO3G 09653T0
RO3G 15787T0
Rmic XP 023465274.1
RO3G 10491T0
RO3G 04000T0
Rmic XP 023469249.1
Pbla XP 018283990.1
Pbla XP 018288661.1
76
100
100
97
100
RPS7
0.02

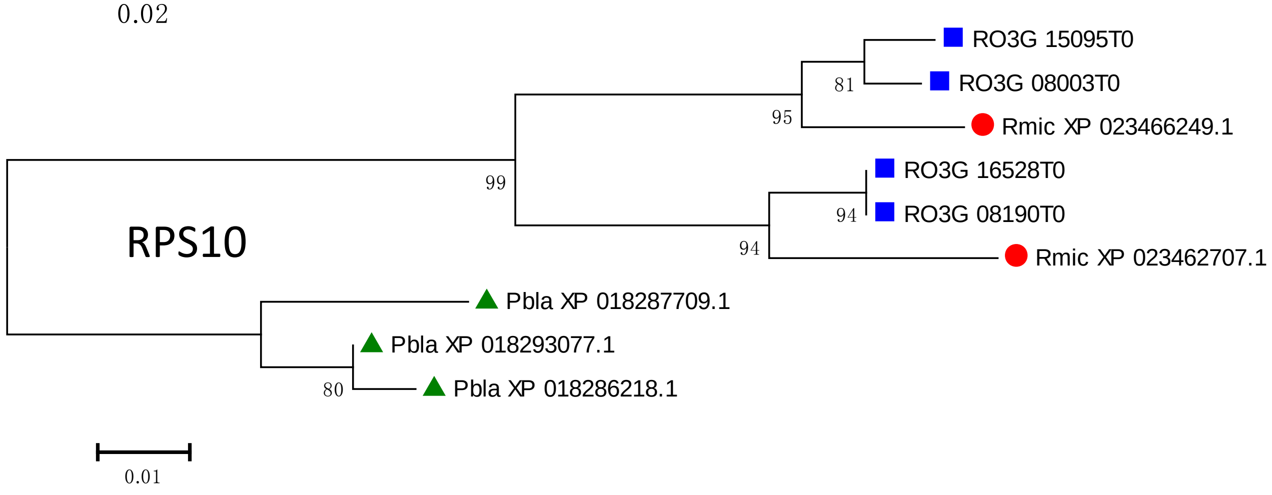
RO3G 15095T0
RO3G 08003T0
Rmic XP 023466249.1
RO3G 16528T0
RO3G 08190T0
Rmic XP 023462707.1
Pbla XP 018287709.1
Pbla XP 018293077.1
Pbla XP 018286218.1
81
95
99
94
94
80
RPS10
0.01

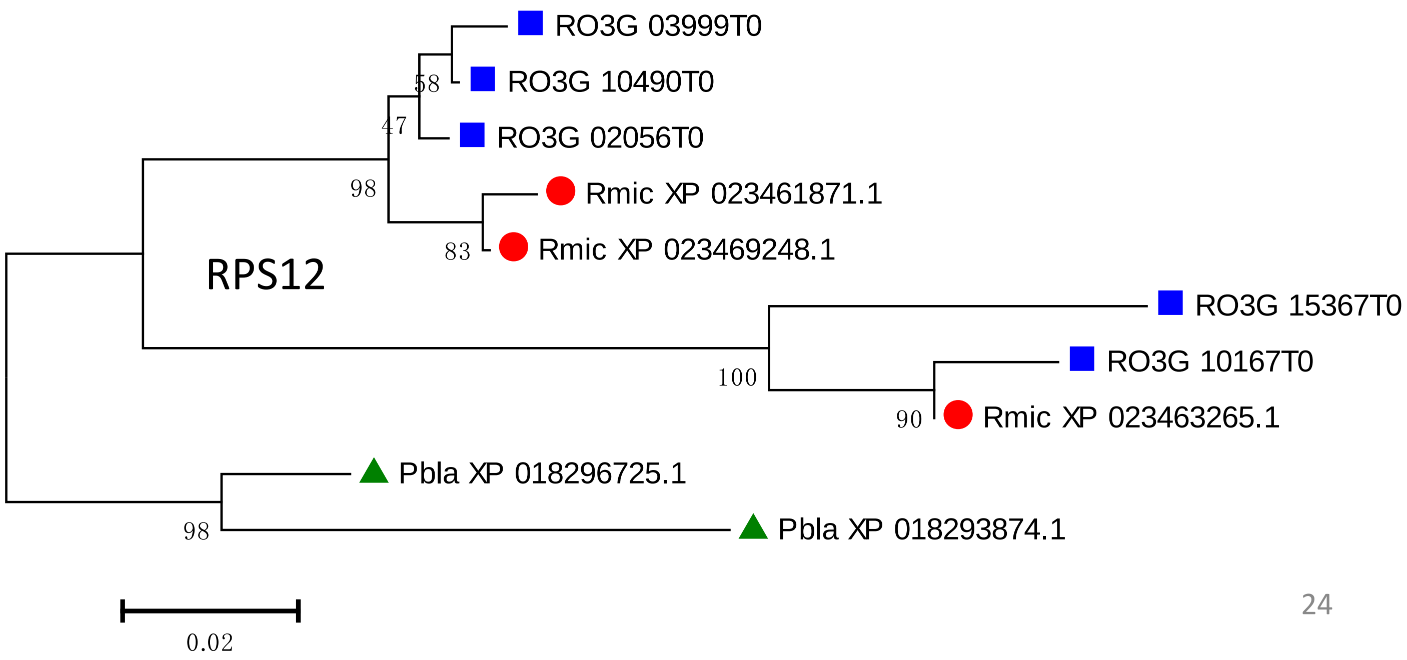
RO3G 03999T0
RO3G 10490T0
RO3G 02056T0
Rmic XP 023461871.1
Rmic XP 023469248.1
RO3G 15367T0
RO3G 10167T0
Rmic XP 023463265.1
Pbla XP 018296725.1
Pbla XP 018293874.1
58
47
98
83
100
90
98
RPS12
0.02

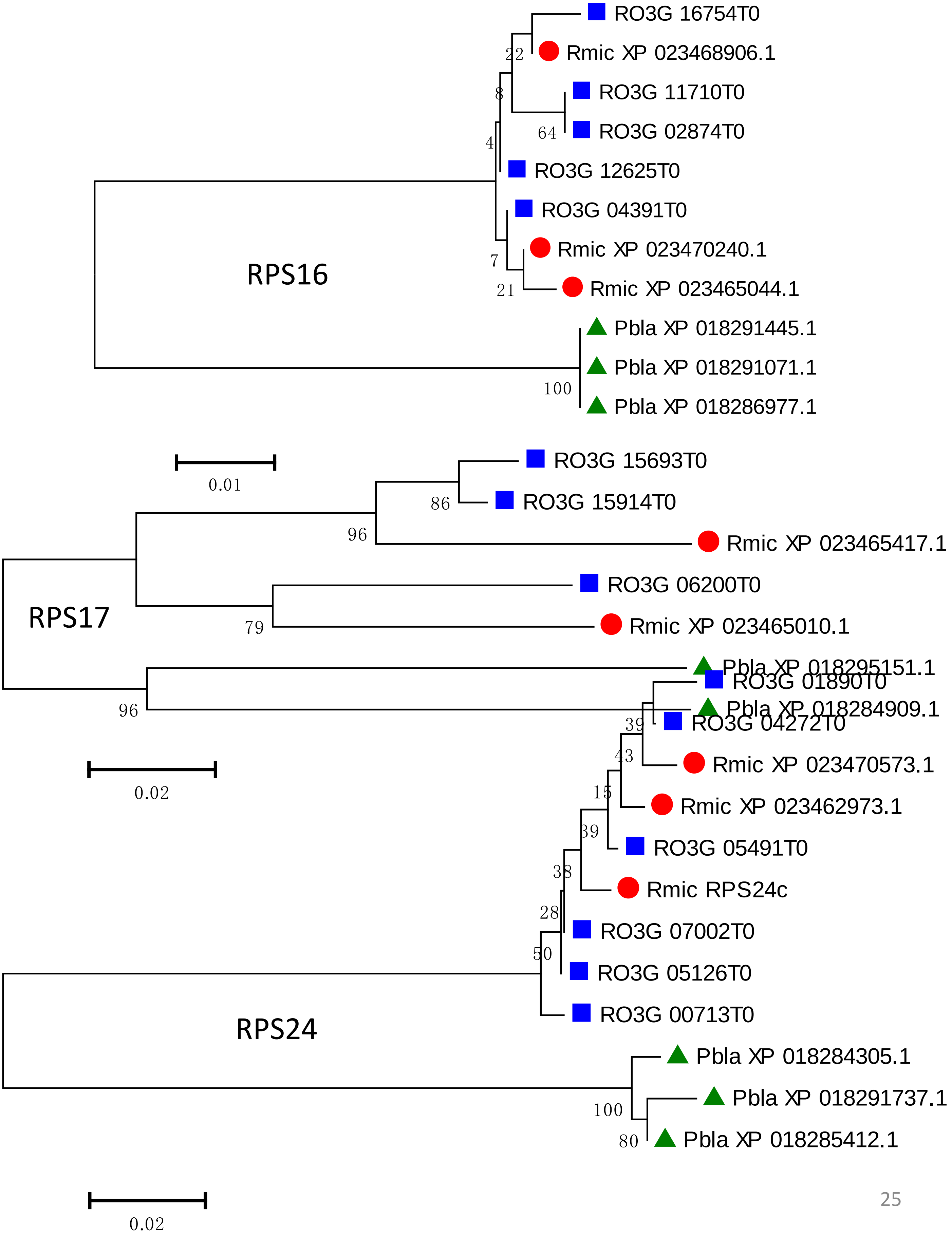
RO3G 16754T0
Rmic XP 023468906.1
RO3G 11710T0
RO3G 02874T0
RO3G 12625T0
RO3G 04391T0
Rmic XP 023470240.1
Rmic XP 023465044.1
Pbla XP 018291445.1
Pbla XP 018291071.1
Pbla XP 018286977.1
22
8
64
4
7
21
100
RPS16
0.01
RO3G 15693T0
RO3G 15914T0
Rmic XP 023465417.1
RO3G 06200T0
Rmic XP 023465010.1
Pbla XP 018295151.1
Pbla XP 018284909.1
86
96
79
96
RPS17
0.02
RO3G 01890T0
RO3G 04272T0
Rmic XP 023470573.1
Rmic XP 023462973.1
RO3G 05491T0
Rmic RPS24c
RO3G 07002T0
RO3G 05126T0
RO3G 00713T0
Pbla XP 018284305.1
Pbla XP 018291737.1
Pbla XP 018285412.1
39
43
15
39
38
28
50
100
80
RPS24
0.02

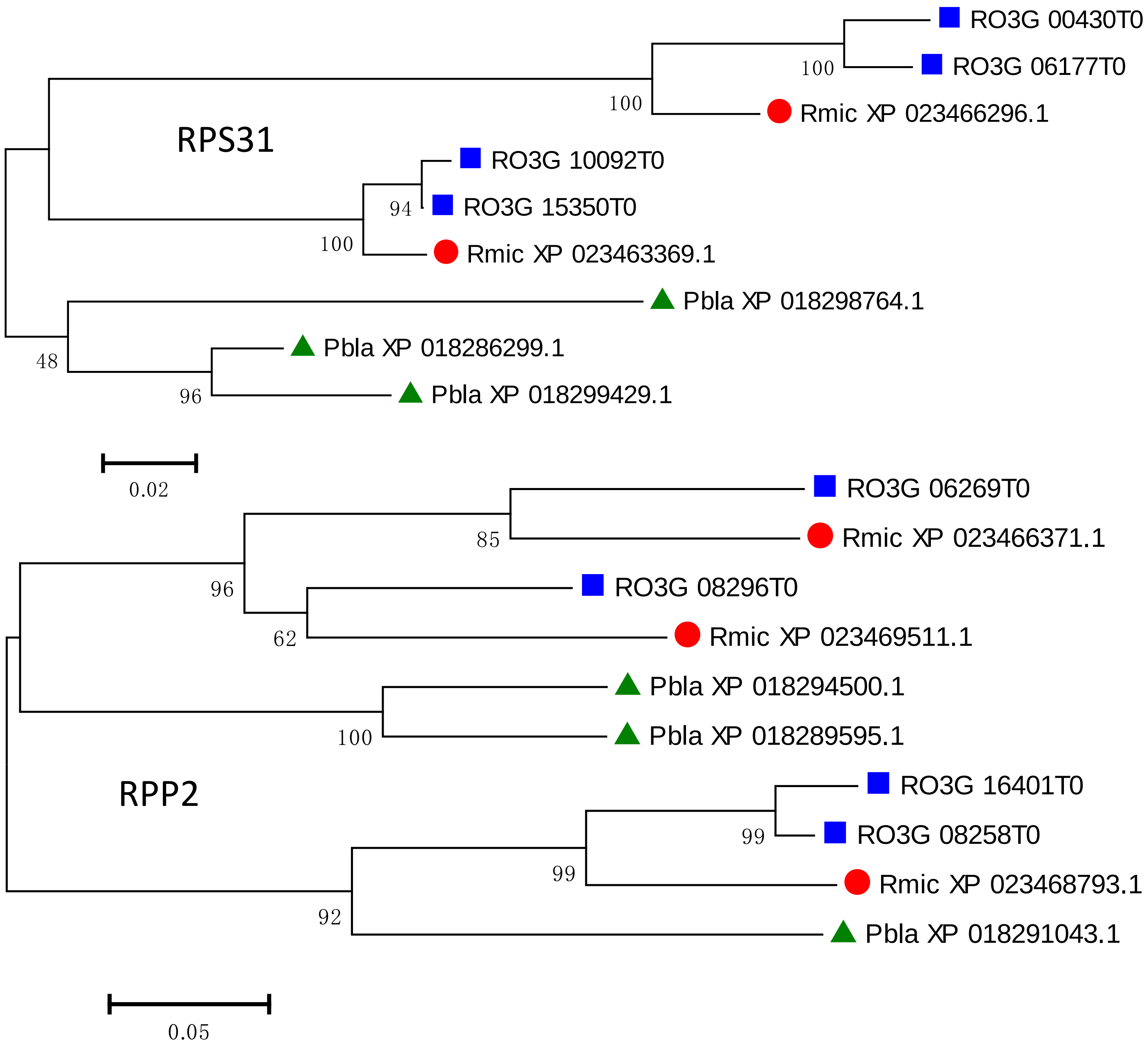

RO3G 00430T0
RO3G 06177T0
Rmic XP 023466296.1
RPS31
RO3G 10092T0
RO3G 15350T0
Rmic XP 023463369.1
Pbla XP 018298764.1
Pbla XP 018286299.1
Pbla XP 018299429.1
100
100
94
100
48
96
0.02
RO3G 06269T0
Rmic XP 023466371.1
RO3G 08296T0
Rmic XP 023469511.1
Pbla XP 018294500.1
Pbla XP 018289595.1
RPP2
RO3G 16401T0
RO3G 08258T0
Rmic XP 023468793.1
Pbla XP 018291043.1
85
96
62
100
99
99
92
0.05

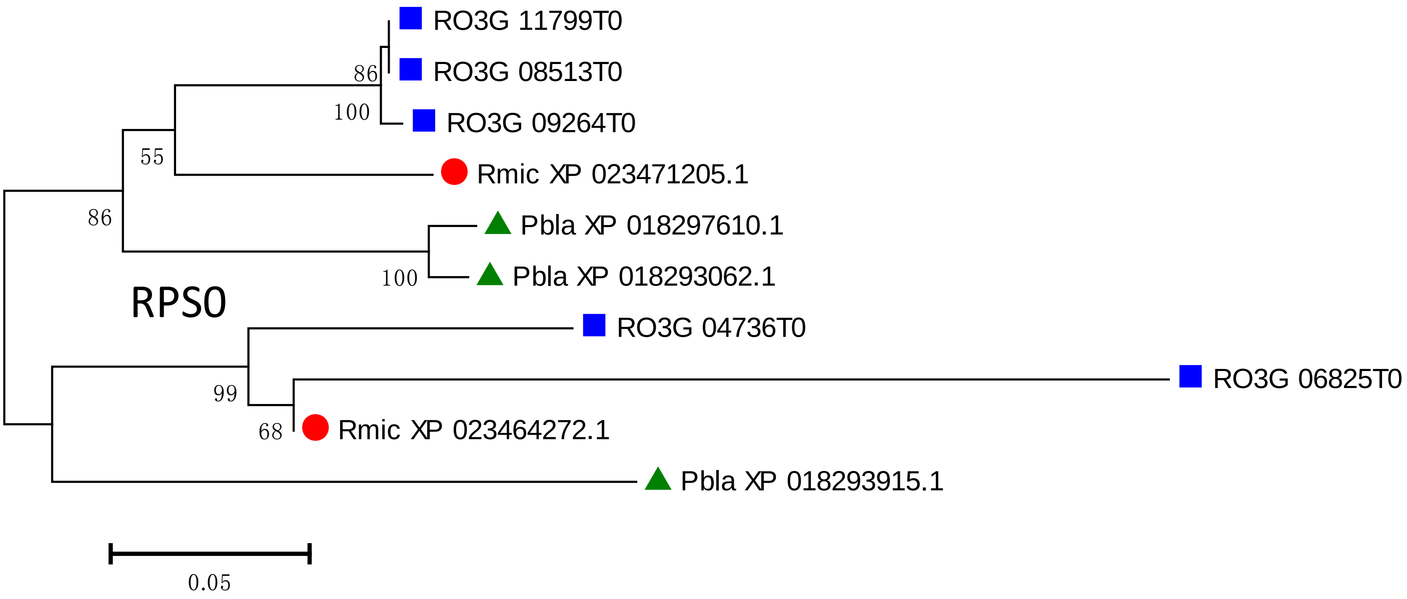
RO3G 11799T0
RO3G 08513T0
RO3G 09264T0
Rmic XP 023471205.1
Pbla XP 018297610.1
Pbla XP 018293062.1
RPS0
RO3G 04736T0
RO3G 06825T0
Rmic XP 023464272.1
Pbla XP 018293915.1
86
100
55
86
100
99
68
0.05

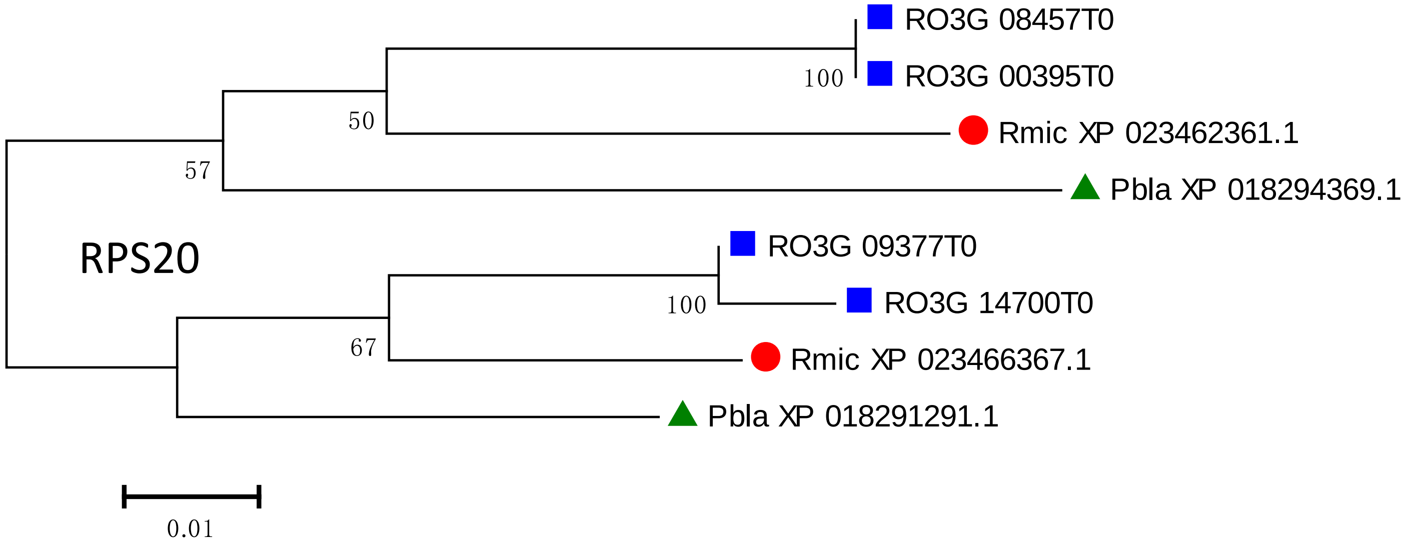
RO3G 08457T0
RO3G 00395T0
Rmic XP 023462361.1
Pbla XP 018294369.1
RPS20
RO3G 09377T0
RO3G 14700T0
Rmic XP 023466367.1
Pbla XP 018291291.1
100
50
57
100
67
0.01
